# Targeted phosphoproteomics of the Ras signaling network reveal regulatory mechanisms mediated by oncogenic KRAS

**DOI:** 10.1101/695460

**Authors:** Anatoly Urisman, Tina L. Yuan, Marena Trinidad, John H. Morris, Shervin Afghani, Juan A. Oses-Prieto, Cayde D. Ritchie, Muhammad S. Zahari, Cyril H. Benes, Alma L. Burlingame, Frank McCormick

## Abstract

**Background:** KRAS mutations are present in up to 30% of lung adenocarcinoma cases and are associated with poor survival. No effective targeted therapy against KRAS is currently available, and novel strategies to counteract oncogenic KRAS signaling are needed.

**Results:** We used targeted proteomics to monitor abundance and site-specific phosphorylation in a network of over 150 upstream and downstream effectors of KRAS signaling in H358 cells (KRAS G12C). We compared patterns of protein regulation following sustained signaling blockade in the RAS/ERK module at two different levels, KRAS and MEK. Network-based analysis demonstrated complex non-linear patterns of regulation with wide-spread crosstalk among diverse subnetworks. Among 85 most regulated proteins in the network, only 12 proteins showed concordant regulation in response to signaling blockade at both KRAS and MEK levels, while the remainder were either specifically regulated in response to KRAS knockdown or MEK inhibition or showed orthogonal regulation in both conditions. Dephosphorylation of DNA methyltransferase 1 (DNMT1) at S714 was identified among the changes unique to KRAS knockdown, and here we elucidate the role of this phosphorylation in KRAS-dependent transcriptional silencing of tumor suppressor genes.

**Conclusions:** Network-based analysis of the Ras signaling has shown complex non-linear patterns of regulation with wide-spread crosstalk among diverse subnetworks. Our work illustrates a targeted proteomics approach to functional interrogation of complex signaling networks focused on identification of readily testable hypotheses. These methods are widely applicable to diverse questions in tumor biology and other signaling paradigms.

## Background

Activating mutations in KRAS are present in approximately one third of all lung adenocarcinoma cases [1, 2]. KRAS-mutant tumors are associated with dismal survival and poor response to therapy [3, 4]. These so-called “driver” mutations often result in sustained activation of the RAS/ERK signaling pathway, which promotes cell proliferation and blocks cell death [5]. Despite much effort over the last couple of decades, no effective targeted therapy for KRAS has been developed [6]. Targeted therapies directed at the kinases downstream of KRAS, including BRAF, MEK and ERK, have demonstrated effectiveness in the treatment of melanoma with BRAF driver mutations [7]. However, so far, these compounds have not produced clinically significant effects in KRAS-mutant tumors [8]. This lack of efficacy suggests that signaling mediated by mutant KRAS has effects that go beyond the mere constitutive activation of the RAS/ERK signaling module. Consistent with this notion, several mechanisms of feedback within the RAS/ERK module and crosstalk with other signaling cascades have been described, including activation of receptor tyrosine kinases and PI3K/AKT signaling by MEK inhibitors [9], feedback activation of MEK by first-generation MEK inhibitors [10], and ERK activation of mTORC1/4E-BP [11].

Understanding mechanisms of signal modulation, rewiring, and crosstalk within complex signaling networks requires simultaneous measurements of protein abundance and phosphorylation (and likely other post-translational modifications) at numerous sites in dozens if not hundreds of proteins. Such measurements are not possible with conventional immuno-detection methodologies, as phosphosite-specific antibodies are not available for the great majority of phosphorylation (and other PTM) sites that have been identified in high throughput proteomic screens. However, recent advances in targeted proteomics have made possible the goal of simultaneous abundance and phosphosite-specific quantitation.

We used targeted proteomics based on selected reaction monitoring (SRM) to monitor abundance and phosphorylation of approximately 150 proteins involved in RAS signaling, herein referred to as *RAS signaling network*. We specifically focused on the question of how signaling mediated by mutant KRAS might be different from that of simple constitutive activation of downstream signaling through MEK/ERK. To this end, we compared signaling within the RAS network in H358 lung adenocarcinoma cells (KRAS G12C mutant) following blockade of signaling through the RAS/ERK module at two different levels – at the level of KRAS using siRNA against KRAS (si-KRAS), and at the level of MEK using selumetinib (AZD6244), a specific MEK1/2 inhibitor (MEK-i).

## Results

### Experimental Approach

We defined the RAS signaling network as a set of approximately 300 human proteins (Additional File 1) known to be involved in signal transduction upstream and downstream of the RAS/ERK module. In order to develop SRM methods to quantify abundance (unmodified peptides) and site-specific phosphorylation (phospho-peptides) from these proteins, we first conducted “deep” proteome discovery of unmodified and phosphorylated peptides from a pool of 8 different lung adenocarcinoma cell lines (Additional File 2). The discovery effort produced high resolution spectra of >100,000 unmodified peptides from 10,372 proteins and >81,000 phospho-peptides from 5,208 proteins, with a total of 12,760 proteins represented (the dataset is further described in the Methods and is freely available for download). The obtained reference spectra and corresponding retention times were used to schedule and refine the performance of the SRM methods as previously described [12].

The SRM workflow we employed (Supplementary Figure S1) started with trypsin protein digests of whole cell lysates, which were subjected to titanium dioxide (TiO_2_) enrichment. The TiO_2_ eluate (containing mostly phospho-peptides) and the flow-through (containing mostly unmodified peptides) were analyzed using two separate SRM method sets targeting phospho-peptides and unmodified peptides, respectively. The measured SRM intensities were used to quantify site-specific phosphorylation of 114 proteins and protein abundance of 143 proteins (a total of 159 proteins) in the RAS network. Among these, 50 proteins had both abundance and phosphorylation measurements allowing us to estimate the net changes in phosphorylation at particular sites that are independent of concurrent changes in abundance.

Using the outlined SRM approach, we compared signaling patterns in the RAS network in H358 cells under three biological states: (1) following siRNA knockdown of KRAS (si-KRAS); (2) following inhibition of MEK by selumetinib (MEK-i); and (3) in mock-treated control (NC). The cells were assayed at peak activity of each treatment (70 hrs after si-KRAS and 48 hrs after MEK-i) in order to focus on the stable “rewired” states of signaling achieved as the result of these perturbations.

### Statistically significant changes in abundance and phosphorylation

We first sought to identify statistically significant changes in abundance and phosphorylation introduced by si-KRAS and MEK-i treatments compared to negative control (NC, siNegCon+DMSO). For this purpose, the SRM intensities for all monitored peptides in si-KRAS and MEK-i conditions were compared to NC using log2 ratios, which were normalized by setting the medians of the log2 ratios in the individual conditions to 0. Three replicate measurements for each peptide were aggregated using the median of the normalized log2 ratios. To calculate protein abundance changes in a given protein, we used the median log2 ratio of all available unmodified peptides for that protein. Phospho-peptide log2 ratios were not aggregated at the protein level and were considered as individual observations based on the assumption that each phospho-peptide is an independent site of regulation and may show differential phosphorylation. When both abundance and phospho-peptide measurements were available in a given protein, the phospho-peptide log2 ratios were adjusted by the abundance log2 ratio of that protein in order to highlight the net change in phosphorylation that is independent of the abundance.

Overall, changes in phosphorylation in response to si-KRAS and MEK-i were more pronounced than the changes in abundance, as judged by the number of proteins and peptides with statistically significant changes (Figure 1). Neither abundance nor phosphorylation measurements demonstrated significant bias toward up- or down-regulation in either condition, implying that the signaling shift following si-KRAS as well as MEK-i involves both positive and negative adjustments in both abundance and phosphorylation. Statistical significance of the observed changes was determined using a two-tailed t-test as described in the Methods. Measurements of abundance and site-specific phosphorylation for each measured protein are plotted in Additional File 3. Statistically significant abundance and phosphorylation changes produced by si-KRAS compared directly to MEK-i response (log2 ratio of si-KRAS/MEK-i) are shown in Supplementary Figure S2). False discovery rate (FDR) of our SRM measurements was estimated at 8% using a set of representative synthetic isotope labeled peptides (see Methods and Supplementary Figure S3).

**Figure 1.**
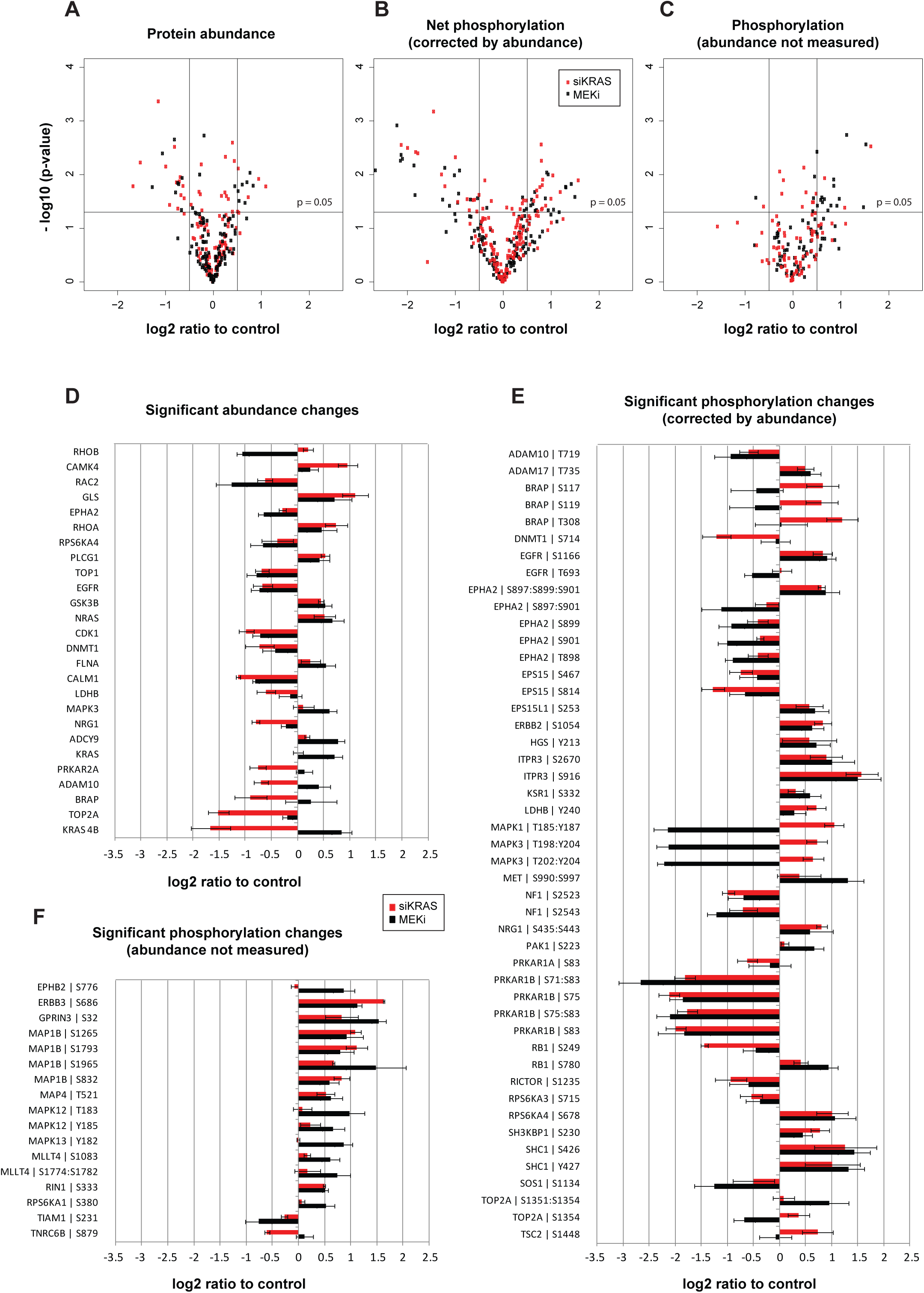
Most significant abundance and phosphorylation changes. Statistical significance threshold was set at p-value <0.05 and log2 change of >0.5 for both si-KRAS (red) and MEK-i (black) conditions. (**A**) Volcano plot of abundance measurements for each protein reflects log2 ratios of p-value-weighed medians of multiple peptides. (**B**) Volcano plot of median log2 ratios of individual phosphorylated peptides corrected by protein abundance. (**C**) Volcano plot of median log2 ratios of individual phosphorylated peptides for which corresponding protein abundance measurements could not be measured. (**D**) Individual statistically significant abundance changes corresponding to plot A. (**E, F**) Individual statistically significant phosphopeptide changes corresponding to plots B and C, respectively. Bars reflect median log2 ratios relative to control and error whiskers – standard deviations.

The most significant expected changes included a decline in KRAS-4B abundance following si-KRAS (over-3-fold relative to NC and over-5-fold relative to MEK-i), and dephosphorylation of MAPK3(ERK1) and MAPK1(ERK2) following MEK-i at the canonical TEY phosphorylation sites targeted by MEK1/2 (almost 5-fold relative to NC and 8-fold relative to si-KRAS). Other observed changes reflecting both known and novel regulatory mechanisms are described below in the context of the entire network.

### Highly regulated proteins in the RAS signaling network

To highlight the proteins with highest degree of regulation in response to si-KRAS and/or MEK-i, with possible combined effects on abundance and phosphorylation, we calculated an aggregate score, *Regulation Index* (RI), for each of the proteins (nodes) in the network (Figure 2). RI was defined as the sum of the absolute values of all log2 abundance and phosphorylation ratios greater than 0.5 (RI = |R_abundance_ > 0.5| + Σ|R_phosphorylation_ > 0.5|). Three RI values were calculated. RI_si-KRAS_ and RI_MEK-i_ highlighted the most regulated proteins as compared to NC, and RI_si-KRAS-MEK-i_ reflected the differential regulation between the two conditions.

**Figure 2.**
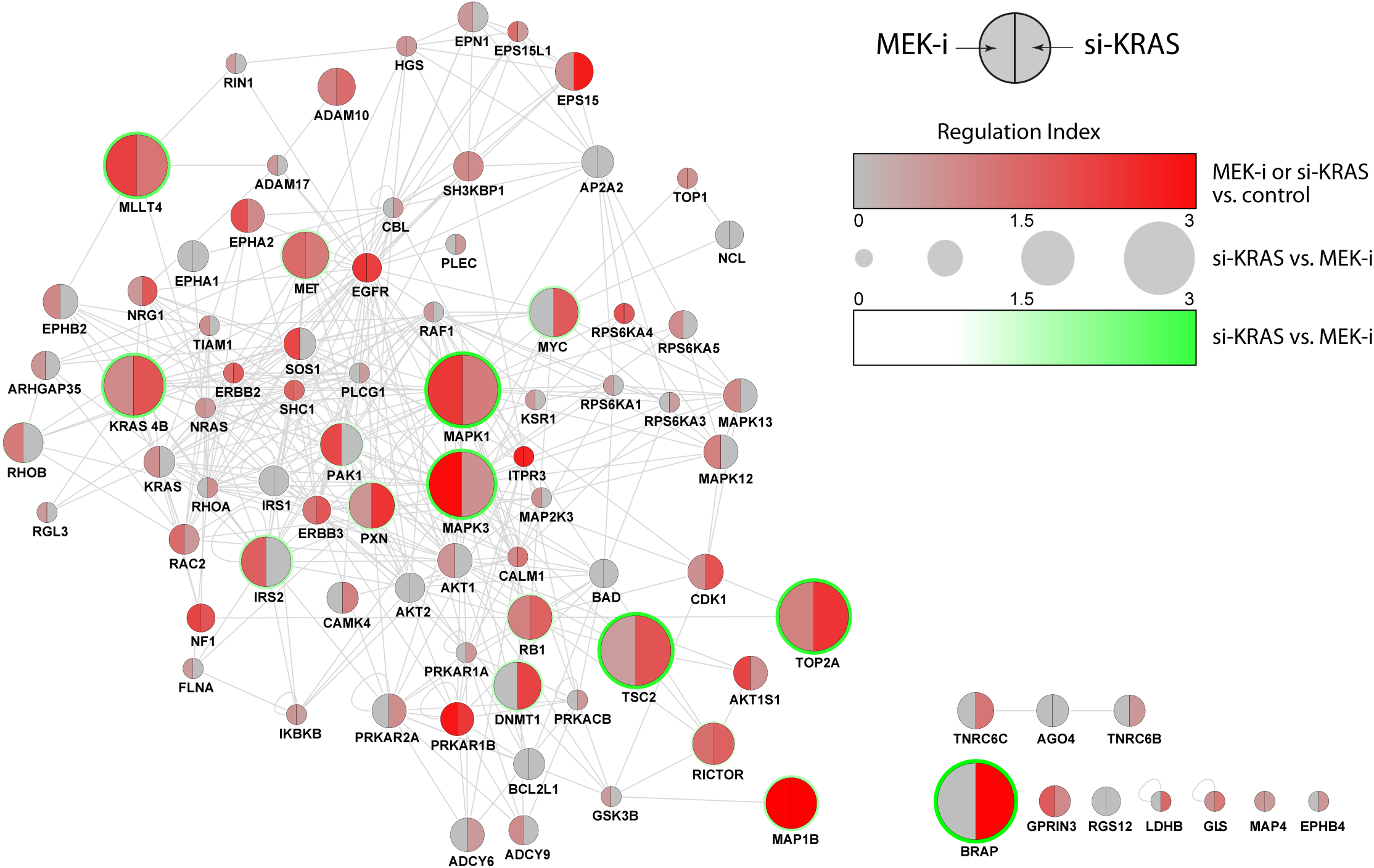
Most regulated proteins in response to si-KRAS and MEK-i. Network connectivity is based on Reactome FI database of functional protein interactions. Regulation of each protein is estimated using an aggregate metric, Regulation Index, representing combined magnitude of abundance and phosphorylation changes in response to MEK-i (left half-node) and si-KRAS (right half-node) relative to the control. Node size reflects the Regulation Index of si-KRAS relative to MEK-i.

We used Reactome-FI database of functional protein interactions [13] to construct the network. All 159 proteins quantified in our experiments were included in the RI calculations, among which 85 proteins had at least one of the three RI scores above the 0.5 threshold, i.e. at least one of the component log2 ratios had to have a magnitude of 0.5 (∼1.5-fold change). Remarkably, 75 of the 85 most regulated proteins selected by this analysis maintained the network connectivity of the larger initial network, supporting their functional proximity to the KRAS and MEK nodes perturbed in the experiments.

The most regulated proteins were not limited or biased to those immediately upstream or downstream of the RAS/ERK module in the conventional sense of the “signaling pathways”, and in fact, network-wide regulation effects were observed. Importantly, regulatory changes in response to MEK-i were not a subset of si-KRAS changes, demonstrating non-linear, distributed, signal transduction patterns that are distinctly different depending on the point of signaling blockade in the RAS/ERK module (KRAS vs. MEK in our system).

Of particular interest were proteins that were differentially regulated by the two conditions. A number of proteins showed regulation patterns that were similar in both MEK-i and si-KRAS conditions (Figure 2; small circles), e.g. RPS6KA4, ITPR3, NF1, and EGFR among others. But the approach also highlighted a set of proteins with different patterns of regulation when comparing the two conditions (Figure 2; large circles), including as expected MAPK1, MAPK3, and KRAS 4B, as well as less expected TSC2, BRAP, TOP2A, MLLT4, MAP1B, IRS2, MET, DNMT1, PXN, and PAK1.

Importantly, this subset of differentially regulated nodes not only included proteins that were regulated in only one of the two conditions and unregulated in the other, but also those that exhibited two distinctly different patterns of regulation in si-KRAS vs MEK-i. To further illustrate this point (Figure 3), the calculated RI values were used to classify each of the 85 most regulated proteins into one of four categories: (i) regulated in response to si-KRAS only (17 proteins); (ii) regulated in response to MEK-i only (21 proteins), (iii) concordantly regulated in response to both conditions (12 proteins); and (iv) orthogonally regulated by the two conditions (35 proteins). Therefore, only 12 of 85 most regulated proteins in the network were coordinately regulated in response to si-KRAS and MEK-i, while the remainder fell into one of three categories of differential regulation (i, ii, iv). These differentially regulated proteins are of particular interest as they represent nodes in the network that are engaged differently by active KRAS as compared to active MEK/ERK, either directly or as a result of compensatory mechanisms in response to ablation of signaling at these two levels in the RAS/ERK module.

**Figure 3.**
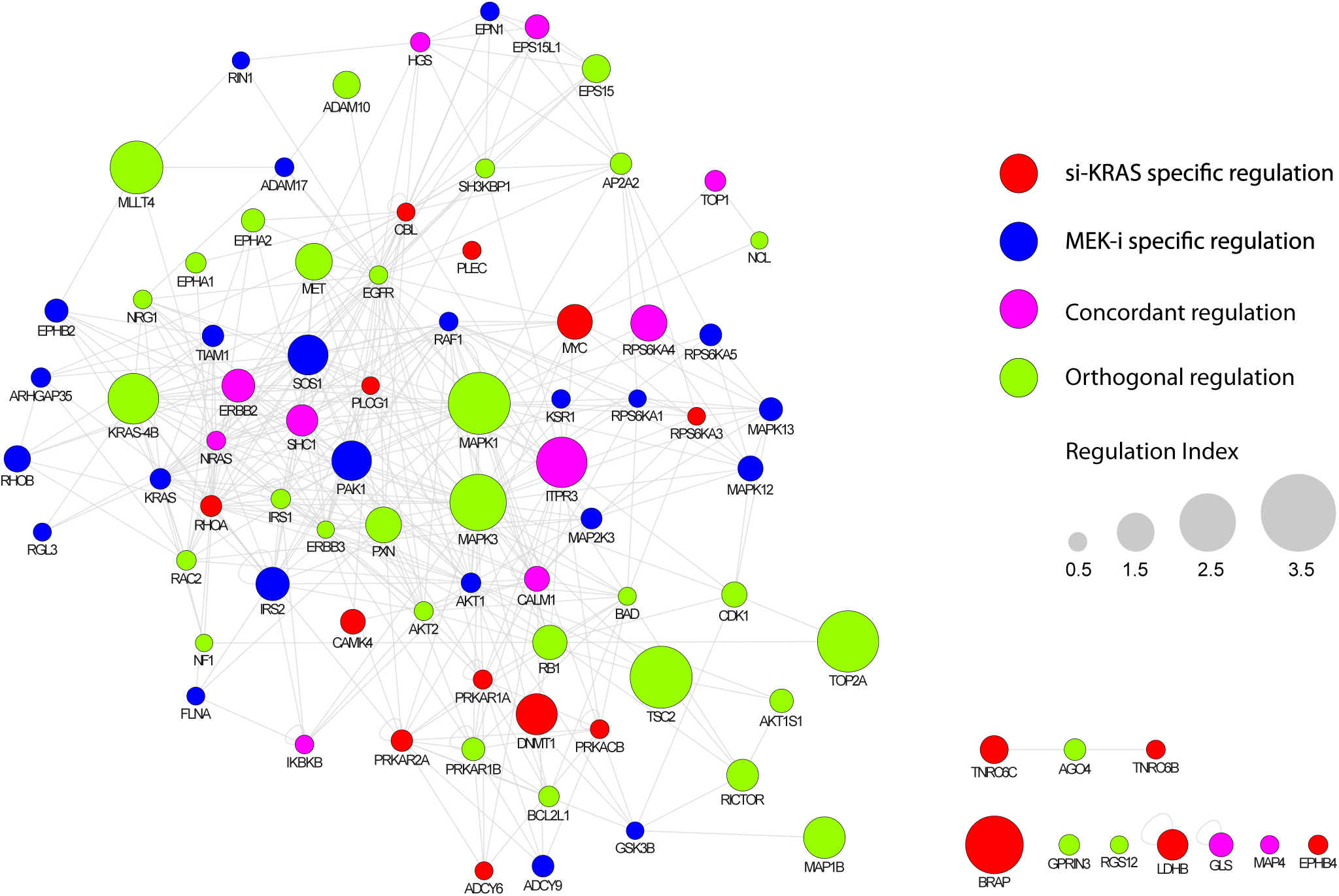
Major patterns of protein regulation in response to si-KRAS and MEK-i. Network connectivity is based on Reactome FI database of functional protein interactions. Based on Regulation Index values, each regulated protein is classified into one of four categories: si-KRAS specific (red), MEK-i specific (blue), concordant (purple), or orthogonal (green). Node size reflects the magnitude of corresponding Regulation Index.

### Patterns of regulation in the RAS signaling network in response to MEK-i and si-KRAS

A detailed network-based view of abundance and phosphorylation changes in the RAS signaling network under MEK-i and si-KRAS conditions (Figure 4) highlights both common patterns of regulation in response to either condition as well as condition-dependent differential regulation. This analysis is based on all SRM measurements obtained in the study (159 proteins), with median log2 ratios visualized independently of their statistical significance.

**Figure 4.**
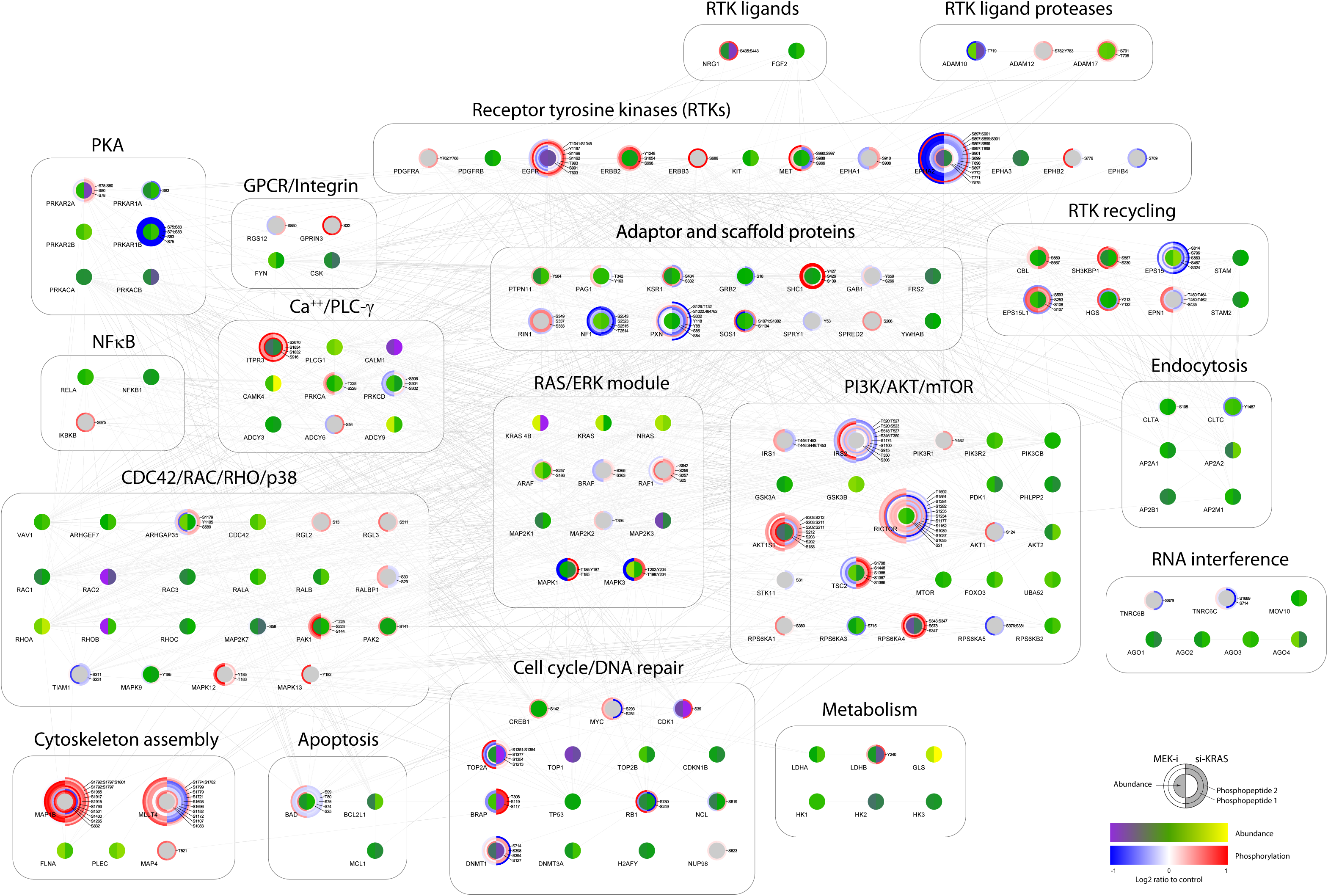
Differential regulation in RAS signaling network in response to si-KRAS and MEK-i. Network connectivity is based on Reactome FI database of functional protein interactions. Protein nodes are grouped into subnetworks according to conventional signaling pathways or common cellular roles. Changes in abundance (node center) and phosphorylation (outer circles) in response to MEK-i (left half-node) and si-KRAS (right half-node) are shown as log2 ratios relative to the control.

Observed patterns of regulation (Figure 4) involved signaling cascades (or subnetworks) outside of the RAS/ERK module, highlighting the complex nature of cross-regulation affecting the entire network. Although changes in signaling were seen in essentially all subnetworks (roughly corresponding to conventional signaling “pathways”, highlighted as groups of proteins in the figure), only a subset of subnetworks showed differential patterns of regulation under si-KRAS and MEK-i. Furthermore, only a subset of proteins and phosphosites within these subnetworks showed differential regulation.

#### RAS/ERK module

As expected, KRAS showed the greatest measured decrease in protein abundance under si-KRAS (Figure 1D and Supplementary Figure S2). Interestingly, the knockdown effect was measured only by a KRAS 4B-specific peptide (over 3 -fold decrease in abundance relative to NC), while a peptide shared by both 4A and 4B isoforms did not show any change likely due to coelution and co-fragmentation of a highly homologous peptide from HRAS and NRAS (with only a single residue difference and a large overlap in the y-series fragment ions used for its detection by SRM, Supplementary Figure S4).. Surprisingly, both peptides documented a near 2-fold increase in KRAS abundance under MEK-i, confirmed by western blotting (Supplementary Figure S5), suggesting that KRAS expression is up-regulated in response to loss of signaling through MEK/ERK.

Canonical MEK target sites in both ERK1 (MAPK3) and ERK2 (MAPK1), as expected, were dephosphorylated almost 5-fold in response to MEK-i, but surprisingly this coincided with a 1.5-fold increase in ERK1 (MAPK3) abundance. This observation suggests a compensatory up-regulation of ERK1 expression in response to loss of MEK activity.

MAP2K3 (MEK3) demonstrated a modest decrease in abundance in response to MEK-i, and several phosphorylation sites in RAF and MEK proteins showed a trend for differential phosphorylation under the two conditions. These included homologous conserved BRAF S365 and RAF1 S259 inhibitory sites phosphorylated by AKT [14, 15], RAF1 S642 inhibitory site phosphorylated by ERK [16], and MAP2K2 (MEK2) T394 inhibitory site phosphorylated by CDK5 [17, 18]. These sites are thought to function by providing negative feedback on RAS/ERK signaling through AKT, ERK, and CDK5, respectively. Although the contribution of these individual phosphorylation events to the composite signaling output of RAS/ERK signaling is unclear, the observed opposing directions in phosphorylation at these sites highlight the combinatorial nature of counter-balance regulation (crosstalk) between the RAS/ERK module and other arms of the wider RAS signaling network.

#### PI3K/AKT/mTOR subnetwork

The PI3K signaling subnetwork showed a complex pattern of regulatory changes, including many sites with differential phosphorylation in response to si-KRAS vs. MEK-i (Figure 4). This included several previously uncharacterized sites in IRS1 and IRS2, PIK3R1, TSC2, AKT1, AKT1S1, and RICTOR. The numerous sites of differential phosphorylation in this subnetwork highlight the combinatorial complexity of cross-regulation and co-dependence of PI3K/AKT/mTOR and RAS/ERK signaling.

Phosphorylation of AKT1 at S124 was approximately 2-fold lower in si-KRAS compared to MEK-i (Supplementary Figure S2). This site near the catalytic domain of AKT1 has been shown to be a target of PKC-zeta (PRKCZ) in vitro and proposed to act as a possible co-activator of AKT1 [19]. In our experiments, phosphorylation at this site was reduced under si-KRAS and increased under MEK-i as compared to the control. These results are consistent with inhibition of AKT1 following si-KRAS treatment and compensatory activation of AKT1 following MEK-i treatment via KRAS-dependent activation of PI3K signaling. Of note, both effects have been previously observed by western blot of phospho-AKT T308 and S473, canonical sites of AKT1 activation [9, 20, 21], which suggests that phosphorylation of S124 parallels that of T308 and S473 and may contribute to AKT1 activation downstream of oncogenic KRAS.

All measured phosphorylation sites in TSC2, a negative regulator of mTOR, appeared to be differentially regulated, with observed increased phosphorylation under si-KRAS and decreased under MEK-i. Among these, only S1488, a previously uncharacterized site, reached statistical significance (Figure 1E). TSC2 is negatively regulated through phosphorylation by AKT, leading to mTORC1 activation. Conversely, AMP kinase (AMPK) positively regulates TSC2 through phosphorylation at S1387 [22] [23] [24], leading to mTORC1 inhibition. Interestingly, we observed concordant regulation of TSC2 and AKT under si-KRAS and MEK-i. si-KRAS increased TSC2 S1387 and decreased AKT1 S124 phosphorylation to effectively shut down both mTOR and AKT, while MEK-i decreased TSC2 S1387 but increased AKT1 S124 phosphorylation to reactivate both mTOR and AKT, perhaps through positive feedback.

S1235 on RICTOR was dephosphorylated under both si-KRAS (2-fold) and MEK-i (1.5-fold) conditions (Figure 1E and Figure 4). It was previously shown to act as a tumor suppressor site, and its phosphorylation correlated with reduced AKT activation, decreased proliferation and lower tumorigenicity of HRAS-transformed mouse embryonic fibroblasts (MEFs) [25]. Our results suggest that unlike other differentially regulated sites in RICTOR, S1235 is dephosphorylated in response to blockade of RAS/ERK signaling at the level of either KRAS or MEK, consistent with a possible AKT feedback activation mechanism at the level of or below MEK. Importantly, increased AKT activity has been correlated with MEK-i resistance in lung adenocarcinoma cell lines and was proposed as a possible mechanism behind MEK-i resistance in patients [26]. The observed dephosphorylation of RICTOR at S1235 under si-KRAS and MEK-i suggests a possible role for this site in the mechanism of compensatory AKT activation in response to RAS/ERK blockade.

A set of uncharacterized sites in IRS1 and IRS2 showed a trend for differential phosphorylation, including statistically significant IRS1 T446:S449:T453 triple-phosphorylated peptide, detection of which increased under MEK-i and decreased under si-KRAS. Additionally, phosphorylation of PRS6KA1 (RSK1) at S380 was significantly higher under MEK-i than under si-KRAS. This autophosphorylation site in RSK1 is dependent on prior phosphorylation by ERK and serves as a docking site for PDK1 required for activation of RSK1 and phosphorylation of its downstream targets [27], including RAPTOR to activate mTORC1 [28]. Biological significance of the observed differential regulation at these sites in response to si-KRAS and MEK-i is unclear and warrants further study.

#### Receptor tyrosine kinases

Regulatory changes were also evident among receptor tyrosine kinases (RTK). EGFR showed a decline in abundance under both si-KRAS and MEK-i conditions, which coincided with phosphorylation changes at multiple sites in proteins that mediate EGFR endocytosis and recycling in a pattern that was similar in the two conditions. Interestingly, EGFR T693-containing peptide and T1041:S1045 double-phosphorylated peptide demonstrated trends for differential regulation under the two conditions. Phosphorylation at T693 has been previously found to be insensitive to gefitinib treatment [29], and phosphorylation at both S1042 and S1045 was sensitive to inhibition by lapatinib [30]. Previously uncharacterized sites in MET (S966 and S988), EPHA1 (S908/S910), EPHB2 (S776), and ERBB3 (S686) were differentially regulated by si-KRAS and MEK-i. However, the functional significance of these findings is unknown.

Of note, abundance of Neuregulin (NRG1), a precursor to the ligand that interacts with ERBB receptors (measured by a peptide in its cytoplasmic C-terminal portion) as well as abundance of ADAM10, a secreted metalloprotease involved in processing of NRG1 to release an N-terminal extracellular ligand peptide capable of activating both AKT and ERK signaling [31], were both reduced in response to si-KRAS but not MEK-i. This suggests a possible feed-forward mechanisms of ADAM10/NRG1 expression downstream of KRAS signaling that does not depend on MEK/ERK activation.

#### Adaptor and scaffold proteins

Proteins in this loosely defined group (Figure 4) did not show significant changes in abundance based on the available measurements. Several proteins demonstrated phosphorylation changes that were similar in the si-KRAS and MEK-i conditions, including NF1, SOS1, SHC1, GRB2, KSR1, SPRY1, SPRED2, and RIN1, suggesting common regulation mechanisms in response to RAS/ERK blockade downstream of MEK. On the other hand, trends for differential phosphorylation were observed in PAG1, GAB1 and PXN. However, these changes were low-amplitude and did not reach statistical significance.

#### Calcium/PLC-gamma subnetwork

Regulatory changes in the Calcium/PLC-gamma signaling subnetwork included over 1.5-fold reduction in abundance of Calmodulin (CALM1) in both si-KRAS and MEK-i conditions. (Figure 1D and Figure 4). Binding of CALM1 to KRAS is known to have an inhibitory effect on KRAS activation [32, 33] by blocking phosphorylation of KRAS at S181, a site important for its activation of PI3K [34]. Conversely, phosphorylation of oncogenic KRAS at S181 has been shown to mislocalize KRAS from the plasma membrane to the mitochondria to induce apoptosis [35]. Furthermore, the interaction between oncogenic KRAS and CALM1 mediates suppression of Wnt/Ca^2+^ signaling contributing to its tumorigenic properties [36]. Although we were unable to measure phosphorylation at the S181 site (due to lack of a suitable tryptic peptide in this lysine-rich C-terminal stretch of KRAS), the observed down-regulation of CALM1 abundance may serve to alleviate the inhibitory effect on KRAS under conditions of RAS/ERK signaling blockade. It could also be indirect evidence that oncogenic KRAS promotes CALM1 expression, which would serve to suppress p-S181 mediated apoptosis and Wnt/Ca^2+^ signaling to promote survival. Evidence of differential regulation in the Calcium/PLC-gamma subnetwork was also apparent, including higher abundance of CAMK4 following si-KRAS but not MEK-i, and differential phosphorylation of several uncharacterized sites in ADCY6, PRKCA and PRKCD under the two conditions.

#### RHO/RAC/CDC42 and p38 signaling subnetworks

RHOB abundance decreased just over 2-fold under MEK-i (Figures 1D and Supplementary Figure S2), while a minimal (not statistically significant) abundance increase was observed under si-KRAS. This finding is consistent with previously observed increase in RHOB abundance in HeLa cells treated with si-KRAS [37]. It has also been reported that over-expression of HRAS in 3T3 cells leads to transcriptional down-regulation of RHOB that can be rescued by co-expression of dominant negative forms of PI3K and AKT1 but not dominant negative MEK1/2, implicating PI3K/AKT activation in negative regulation of RHOB transcription [38]. Therefore, the decrease in RHOB abundance observed in our experiments might be explained by a compensatory activation of AKT1 downstream of KRAS following MEK-i treatment.

Two p38 kinases, MAPK12 and MAPK13, showed a 2-fold increase in phosphorylation following MEK-i treatment at T183 and Y182, respectively, while si-KRAS did not produce a significant change (Figure 1F and Supplementary Figure S2). These are the canonical TGY activating sites phosphorylated by alternative mitogen-activated kinase kinases MEK3 and MEK6 [39]. Similar activation of p38, as measured by p38 phospho-specific antibody, has been previously observed in HeLa cells treated with MEK-i (PD98059) and correlated with apoptosis through a caspase-dependent mechanism [40].

Phosphorylation of PAK1 at S223 (or S225) was increased by less than two-fold under MEK-i, compared to no change under si-KRAS and no significant abundance changes in either condition. Phosphorylation at S223, by casein kinase II (CK2) based on in-vitro evidence, is essential for PAK1 activity and results in autophosphorylation of PAK1 at S144 and conversion of inactive PAK1 dimer to active monomer [41]. Consistent with this model, phosphorylation of S144 paralleled that of S223 in our experiments, with a greater increase in S144 phosphorylation observed under MEK-i compared to si-KRAS.

Additional changes included down-regulation of abundance of RAC2 (not RAC1 or RAC3) under both MEK-i and si-KRAS conditions. Trends for differential phosphorylation were observed on thus far uncharacterized sites in TIAM1, ARHGAP35, and RALBP1.

#### Protein kinase A (PKA)

PKA regulatory subunits PRKAR1A, PRKAR1B, and PRKAR2A displayed distinctly different patterns of regulation. Only PRKAR2A was differentially regulated at both abundance and phosphorylation levels. Its abundance decreased under si-KRAS and not MEK-i, while two uncharacterized phosphorylation sites, S78 and S80 located in an N-terminal insertion region missing in both PRKAR1A and PRKAR1B, demonstrated increased phosphorylation in response to si-KRAS only.

#### Metabolic enzymes

Two metabolic enzymes, mitochondrial glutaminase kidney isoform (GLS) and lactate dehydrogenase B (LDHB), also showed changes in regulation. GLS abundance increased over 2-fold under si-KRAS and approximately 1.5-fold under MEK-i (Figure 1D and Figure 4). Mutant KRAS transformed pancreatic ductal adenocarcinoma (PDAC) cells have been shown to be dependent on anabolic utilization of glutamine through conversion to glutamate by GLS, and have been shown to be sensitive to GLS inhibitors in vitro [42, 43]. The observed up-regulation of GLS in response to si-KRAS, and to a lesser extent MEK-i. supports the notion of glutaminolysis-dependence for the generation of biosynthetic precursors and NADPH, particularly under the stress of RAS/ERK blockade. LDHB, a key glycolytic enzyme, on the other hand, demonstrated a decrease in abundance in response to si-KRAS and not MEK-i. A prior study has shown that LDHB over-expression correlates with KRAS mutation and/or amplification status and poor patient outcomes in lung adenocarcinoma, consistent with the high glycolytic activity of KRAS mutant cancers [44]. The observed down-regulation of LDHB in response to si-KRAS suggests that LDHB expression may be driven by KRAS activation. Alternatively, the increase in GLS and concomitant decrease in LDHB may reflect a switch in metabolism from glycolysis to mitochondrial respiration upon ablation of oncogenic KRAS signaling [45].

#### RB, p53, cell cycle control, and apoptosis

RB S249 site showed a 2.5-fold decrease in phosphorylation under si-KRAS compared to a 1.5-fold decrease under MEK-i, while RB abundance was not changed significantly. This site is thought to be a target of several different cyclin-dependent kinases and when phosphorylated interferes with binding of EID1 coincident with G0 exit and cell cycle progression; phosphorylation at S249 may be necessary but not sufficient for G0/G1 transition [46, 47]. The observed preferential decrease in phosphorylation of RB at S249 under si-KRAS compared to MEK-i suggests that phosphorylation at this site is more dependent on signaling through KRAS (and possibly mutant KRAS) than activation of MEK/ERK alone.

Phosphorylation of MYC at S281 was reduced over 3-fold by si-KRAS, which was accompanied by an increase in MYC abundance as measured by western blotting (Supplementary Figure S5), while no significant changes were observed in response to MEK-i. It has been shown that dual phosphorylation of S281 by PDK1 and S279 by PKA leads to cooperative stabilization of MYC [48]. It has also been demonstrated that inactivation of MYC by a dominant negative form (Omomyc) leads to both a block in tumor formation and regression of established tumors in LSL-KRAS-G12D mouse lung cancer model [49]. Our results suggest that oncogenic KRAS signaling may exert an effect on the activity and/or stability of MYC via S281 phosphorylation that is independent of MEK/ERK activation.

While abundance of CDK1 was reduced almost two-fold in both MEK-i and si-KRAS conditions, phosphorylation at S39 was differentially increased under si-KRAS and reduced under MEK-i. This site is homologous to CDK2 T39, which has been shown to be a target of AKT; its phosphorylation enhances binding of Cyclin A and facilitates nuclear-cytoplasmic shuttling of CDK2/Cyclin A complex, promoting G2/M transition [50]. We note that the observed differential phosphorylation of CDK1 S39 was inversely correlated with phosphorylation of AKT at S124 (see above), again consistent with inhibition of AKT1 downstream of KRAS under si-KRAS and with compensatory activation of AKT1 under MEK-i.

While abundance of topoisomerase 1 (TOP1) was reduced in both si-KRAS and MEK-i conditions, topoisomerase 2A (TOP2A) abundance was differentially reduced under si-KRAS and not MEK-i. Additionally, several phosphorylation sites in TOP2A showed evidence of differential phosphorylation, including S1354 single phosphorylation that was reduced 1.5-fold and was accompanied by a 2-fold increase in S1351:S1354 double phosphorylation observed only under MEK-i treatment, suggesting possible conversion to the double-phosphorylated state in this region of TOP2A in response to MEK-i. While S1354 is thought to be a cell cycle-dependent target of CDK1 implicated in G2/M transition [51], S1351 site has not been functionally characterized, and neither have been S1213 and S1377 sites which also showed a trend for differential phosphorylation in our experiments.

BRCA1-associated protein (BRAP) abundance was down-regulated almost 2-fold under si-KRAS and not MEK-i. This coincided with differential phosphorylation of S117/S119 (almost 2-fold increase in phosphorylation under si-KRAS and 1.5-fold decrease under MEK-i) and T308 (over 2-fold increase under si-KRAS and no change under MEK-i), none of which have been functionally characterized. BRAP is a tumor suppressor gene, which negatively regulates G1/S transition and is mutated or deleted in many malignancies, including lung and breast cancer [52]. Importantly, BRAP has also been shown to block activation of RAS/ERK signaling at the level of MEK activation by RAF [53]. Our data suggest that the identified sites of differential phosphorylation in BRAP may be mechanistically involved in its cell cycle control and/or modulation of RAF signaling functions, which are in turn differentially regulated by KRAS vs MEK/ERK activation.

At least one of the measured phosphorylation sites in BAD, a proapoptotic protein, showed a trend for differential regulation. Phosphorylation of BAD S99 was increased under MEK-i and reduced under si-KRAS. The site is a known target of AKT, phosphorylation of which causes BAD to bind 14-3-3 gamma (YWHAG) and dissociate from mitochondria blocking its proapoptotic action [54]. The observed differential phosphorylation of BAD S99 is again consistent with inhibition of AKT under si-KRAS and compensatory activation of AKT under MEK-i.

DNA cytosine-5 methyl transferase 1 (DNMT1), in addition to the differentially phosphorylated S714 site described in detail below, showed abundance reduction that was more significant under si-KRAS than MEK-i, and S127, S394 and S398 sites also demonstrated trends for differential phosphorylation under the two conditions, consistent with KRAS-specific DNMT1 regulation that is distinct from mere activation of MEK/ERK.

#### Other proteins

Differential regulation via changes in phosphorylation affected other functionally diverse proteins in the network, including RGS12 at S850, nucleolin (NCL) at S619, MLLT4 at S1083 and S1696, and MAP1B triple-phosphorylation at S1792:S1797:S1801. Although functional significance of these sites will require further study, differential phosphorylation under si-KRAS and MEK-i conditions suggests that functional states of these proteins are likely different in response to activation of KRAS as compared to activation of MEK/ERK alone.

Proteins involved in RNA interference (RNAi) were included as controls but revealed phosphorylation changes that suggest involvement in the mechanism of RNAi. TNRC6B S879 and TNRC6C S714 showed deferential dephosphorylation under si-KRAS and not MEK-i. The two proteins are required for miRNA-dependent translational repression and mRNA degradation, and these homologous sites are located within a conserved N-terminal repeat region (motif II) that is required for interaction with Argonaute family of proteins [55]. The findings raise the possibility that phosphorylation at these sites may mediate docking of Argonaute to TNRC6 proteins.

### KRAS-dependent phosphorylation of DNMT1 at S714 leads to transcriptional silencing of cell cycle genes

Reduction of phosphorylation at S714 in DNA cytosine-5 methyltransferase 1 (DNMT1), a previously uncharacterized site, following si-KRAS treatment (Figure 1E and Supplementary Figure S2) was among the most significant observations in our dataset. DNMT1 is known to transcriptionally silence tumor suppressor genes in KRAS-dependent manner, including cell cycle inhibitors, pro-apoptotic and differentiation genes [56]. We hypothesized that in KRAS-mutant cancer cells, phosphorylation at S714 promotes DNMT1 activity and gene silencing, while dephosphorylation inactivates DNMT1, leading to transcriptional de-repression and growth inhibition. We expressed a non-phosphorylatable S714A or a phospho-mimetic S714D mutant of DNMT1 in H358 cells and measured expression of a panel of DNMT1 target genes (Figure 5A). DNMT1 S714A-expressing cells exhibited increased expression of target genes, suggesting that the S714A mutant has reduced methyl transferase activity (Supplementary Figure S6A). Non-DNMT1 target genes, such as KRAS, were not affected. The DNMT1 S714D mutant did not alter target gene expression compared to cells overexpressing wild-type DNMT1 or GFP, suggesting that most DNMT1 in H358 cells is in the active form.

**Figure 5.**
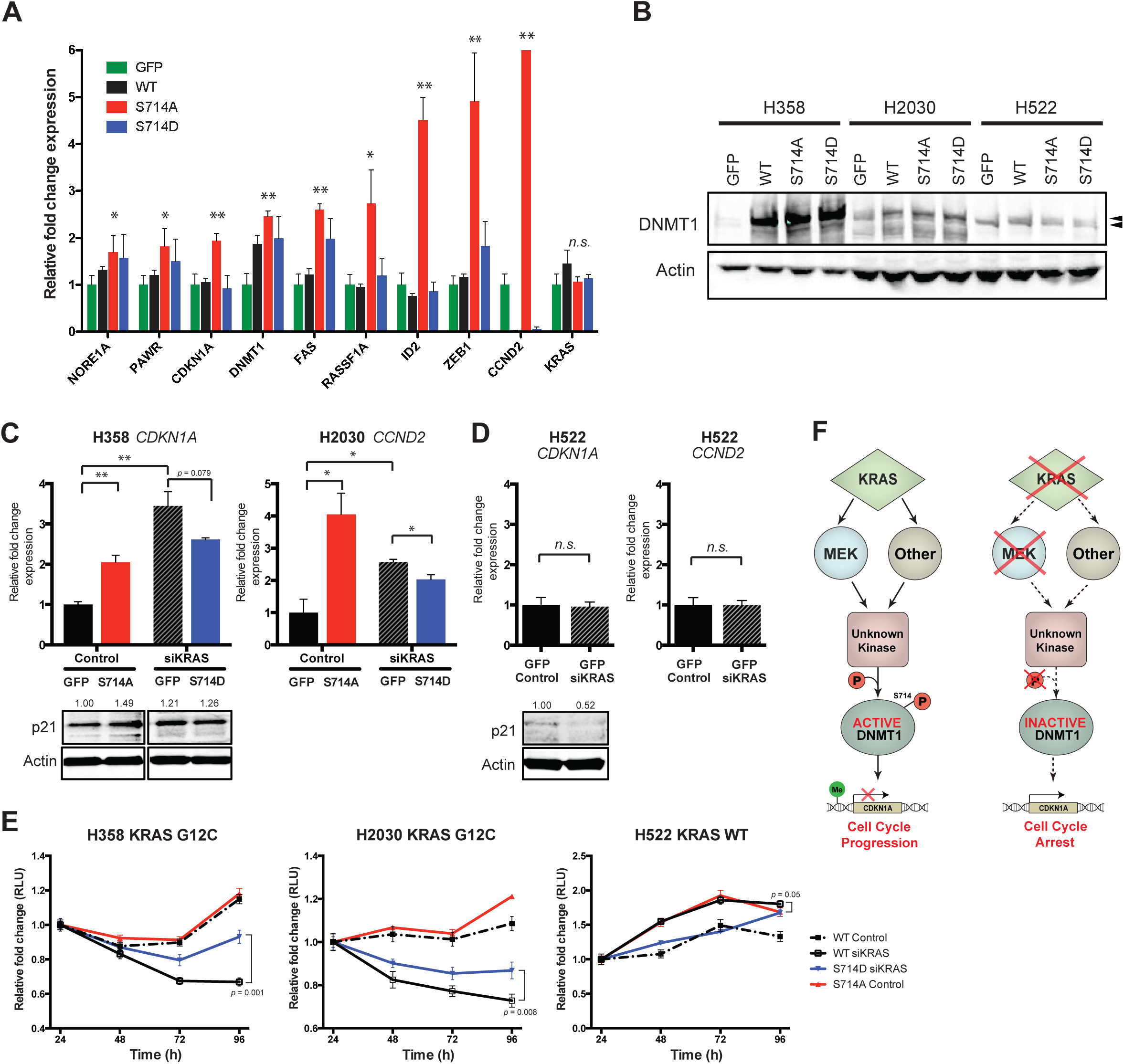
KRAS modulates phosphorylation of DNMT1 at S714. Oncogenic KRAS promotes phosphorylation of DNMT1 S714 to epigenetically silence cell cycle inhibitors. (**A**) mRNA levels of several DNMT1 target genes were measured by qPCR in asynchronously growing H358 cells stably expressing GFP or wild-type, S714A or S714D DNMT1. *p*-values (unpaired t-test) between GFP and S714A lines are shown. n.s. = *p* ≥ 0.05; * = *p* ≤ 0.01; ** = *p* ≤ 0.001. (**B**) GFP or myc-tagged wild-type or mutant DNMT1 was stably expressed in H358, H2030 and H522 NSCLC lines. Arrows point to endogenous (lower) and myc-tagged (upper) DNMT1. (**C**) *CDKN1A* or *CCND2* were upregulated upon KRAS knockdown in H358 or H2030 cells, respectively (black striped bars). The S714A mutant mimicked this upregulation in KRAS replete conditions (red bars) and the S714D mutant slightly suppressed this upregulation in KRAS knockdown conditions (blue bars). p21 protein levels are shown with actin-normalized quantitation (ImageJ). Cyclin D2 protein levels were not consistently detectable. (**D**) In KRAS wild-type H522 cells, KRAS knockdown did not modulate *CDKN1A* or *CCND2* levels. (**E**) Effects on cell proliferation by DNMT1 mutants were most prominent under high confluence. Cells were plated at 75% confluence and proliferation was monitored over 96h. KRAS knockdown significantly inhibited growth in KRAS mutant lines, but not in the KRAS wild-type line. S714A expression was not sufficient to mimic this inhibition in KRAS replete conditions, but S714D expression partially rescued growth inhibition in KRAS knockdown conditions. *p*-values (unpaired t-test) between WT siKRAS and S714D siKRAS at 96h are shown. (**F**) Model of DNMT1-dependent transcriptional silencing of tumor suppressor genes by oncogenic KRAS signaling.

Given that many KRAS-mutant cell lines undergo profound cell cycle arrest upon KRAS knockdown, we investigated if transcriptional down-regulation of cell cycle inhibitory genes *CDKN1A* and/or *CCND2* via DNMT1 phosphorylation was an important part of the oncogenic KRAS program. We expressed wild-type and mutant forms of DNMT1 in H358 (KRAS G12C; TP53 mutant), H2030 (KRAS G12C; TP53 mutant) and H522 (KRAS wild-type; TP53 mutant) lung adenocarcinoma lines (Figure 5B). In KRAS mutant lines (Figure 5C), but not in the KRAS wild-type line (Figure 5D), KRAS knockdown induced *CDKN1A* or *CCND2* upregulation and reduced cell proliferation (Figure 5E).

We then tested if the DNMT1 S714A mutant could mimic these effects in KRAS replete conditions or if the S714D mutant could rescue these effects in KRAS knockdown conditions. S714A expression resulted in an induction of CDKN1A or CCND2, but was not sufficient to induce growth inhibition (Figure 5C,E; red bars and lines). However, S714D expression moderately suppressed CDKN1A or CCND2 levels and was sufficient to partially rescue growth under KRAS knockdown conditions (Figure 5C,E; blue bars and lines). This suggests that proliferation in KRAS mutant NSCLC lines can be partially attributed to promoting DNMT1 phosphorylation, which represses the expression of cell cycle inhibitors. This phenomenon may not be restricted to lung adenocarcinomas, as KRAS mutant ovarian cancer cell lines were recently shown to exhibit selective sensitivity to decitabine, a DNMT1/3a/3b inhibitor [57].

Though we did not pursue the mechanism of DNMT1 phosphorylation, we observed similar rates of growth inhibition and mRNA modulation upon KRAS knockdown at 10% serum and 0% serum (Supplementary Figure S6B), suggesting that oncogenic KRAS rather than exogenous factors in serum are stimulating a yet unknown kinase. Additionally, consistent with the differential SRM results (Figure 2E), AZD6244 treatment did not reliably upregulate target genes (Supplementary Figure S6C) or modulate proliferation upon expression of DNMT1 mutants (Supplementary Figure S6D). This further suggests that DNMT1 phosphorylation is driven by a MEK-independent function of oncogenic KRAS.

## Discussion

The main goal of our study was to identify novel regulatory mechanisms mediated by oncogenic KRAS signaling in lung adenocarcinoma cells. Our targeted proteomic approach directed at the RAS signaling network of over 150 upstream and downstream effectors of KRAS has allowed identification of multiple points of differential regulation in response to RAS/ERK signaling blockade at the level of KRAS (by si-KRAS) vs at the level of MEK (by MEK-i). The observed changes in abundance and phosphorylation in many key proteins within the network highlighted the non-linear and network-wide nature of regulation in response to the two tested perturbations in RAS/ERK signaling. In addition to shared patterns of regulation (presumably due to the common block in signaling through MEK/ERK), majority of the regulated proteins in the network (73 of 85) showed evidence of either KRAS-specific or MEK-specific or distinct dual regulation. Importantly, regulation in response to MEK-i was not a simple subset of regulatory responses to si-KRAS. The identified sites of differential phosphorylation are particularly important as a subset of candidate sites uniquely regulated by oncogenic KRAS.

In this study, we focused our attention on a previously uncharacterized DNMT1 S714 site, which showed statistically significant dephosphorylation in response to si-KRAS and not MEK-i. This observation affirms the notion that KRAS has numerous effectors outside of the canonical RAF/MEK/ERK pathway, which are not affected by MEK inhibition but contribute significantly to the oncogenic program driven by KRAS. We show that oncogenic KRAS promotes the phosphorylation of DNMT1 S714 site, which results in transcriptional repression of genes involved in cell cycle inhibition. The S714 site is immediately adjacent to a stretch of acidic residues within the autoinhibitory CXXC-BAH1 linker in DNMT1, which interferes with binding of the DNA template [58]. Therefore, we predict that S714 phosphorylation activates DNMT1 activity by shifting the autoinhibitory linker away from the DNA binding groove to promote DNA binding. As expected, based on the observed si-KRAS specific S714 dephosphorylation, growth inhibition induced by si-KRAS was more pronounced than that induced by MEK inhibition. Such observations favor the argument that a KRAS inhibitor may be more effective than a potent inhibitor of RAF/MEK/ERK or any other single downstream effector.

In addition to DNMT1, a large number of interesting observations was made in both previously known and more importantly thus far uncharacterized phosphorylation sites, including those in extensively-studies proteins in the RAS signaling network. Many of these observations provide hypotheses that are readily testable by conventional molecular techniques, as illustrated above by the DNMT1 experiments. We are confident that the specific observations regarding possible mechanisms of regulation in the RAS network presented in this study provide a good starting point for further mechanistic studies of oncogenic KRAS signaling.

Several prior phosphoproteomic studies examined oncogenic KRAS-driven signaling in lung cancer. The most notable of these used stable isotope labeling techniques (SILAC and iTRAQ) for proteome-scale analysis of phosphorylation in oncogenic KRAS-driven lung adenocarcinoma cell lines [59] and patient-derived tumors [60]. These studies have identified candidate phosphorylation sites that might be involved in oncogenic KRAS signaling, some of which are also profiled in the current work. In contrast to these studies, we separated phosphosite discovery from functional analysis of their significance in KRAS signaling. Following a proteome-scale discovery of phosphosites in a panel of lung adenocarcinoma cell lines, we focused our functional analysis on the RAS signaling network, i.e. a set of previously known upstream and downstream effectors of KRAS signaling. The targeted nature of this analysis coupled with the specific conditions designed to capture the differences in KRAS-mediated vs. MEK-mediated signaling has provided a comprehensive view of regulatory changes in the network. In addition, the use of SRM-based targeted proteomics allowed reliable label-free quantitation and bypassed the need for offline sample fractionation prior to MS (the main disadvantage of SILAC and iTRAQ approaches). Finally, the SRM methods developed in the study are available for download and may be useful to anyone interested in targeted interrogation of the proteins in the RAS network (either by SRM or PRM).

## Conclusions

It is becoming increasingly clear that understanding the mechanisms of regulation in complex signaling networks that include hundreds of proteins requires simultaneous measurements of phosphorylation (and ideally other PTMs) at hundreds to thousands of sites, in addition to protein abundance. This task, not attainable by conventional antibody-based methodologies, has been a goal of multiple proteomic studies. However, achieving reliable quantitative MS measurements using data-dependent acquisition (shotgun) modalities has proven challenging, particularly for low-abundance proteins and low-stoichiometry PTM sites [61, 62]. By taking advantage of SRM-based targeted proteomics, the work presented here makes a significant leap forward. It illustrates how a question-driven perturbation of the RAS signaling network coupled with as-comprehensive-as-possible targeted monitoring of site-specific phosphorylation and protein abundance leads to identification of mechanistically relevant sites of signaling regulation in the network. We foresee the application of network-based signaling studies such as ours to a wide variety of questions in diverse biological systems, including preclinical models of cancer as well as clinical evaluation of patient tumor samples in precision medicine workflows.

## Methods

### Cell culture, siRNA and MEKi treatments

H358, H2030 and H522 cells were grown in RPMI 1640 Medium (ATCC modification) (Gibco) supplemented with FBS and penicillin and streptomycin. DNMT1 mutant lentiviral constructs were generated using QuikChange site-directed mutagenesis (Agilent) according to the manufacturer’s instructions. Stable cell lines expressing exogenous DNMT1 were generated using lentiviral infection. siRNA transfections were performed at a concentration of 5nM using Lipofectamine RNAiMAX (Invitrogen) according to the manufacturer’s instructions. The siKRAS sequences are provided in Additional File 4 [63] and Qiagen AllStars Negative Control siRNA was used in control samples. AZD6244 was used at a concentration of 1uM (protein and qPCR) or 500nM (growth curves). Protein and RNA were harvested 70h after siRNA transfection (∼48h after target knockdown) and 48h after AZD6244 treatment. To generate growth curves, cells were transfected or treated in 96-well format at 50-70% cell density and grown in RPMI 1640+10% FBS or RPMI 1640+0% FBS. At each time point, cell viability was measured using CellTiter-Glo Luminescent Cell Viability Assay (Promega) in at least 3 technical replicates. Statistical significance was determined by unpaired t-tests, where n.s. = *p* ≥ 0.05; * = *p* ≤ 0.01; ** = *p* ≤ 0.001.

### Western blotting

Lysates for western blotting were collected in 1% NP-40 lysis buffer (20 mM Tris pH 7.4, 150 mM NaCl, 1% NP-40, 1 mM EDTA, 1 mM EGTA, 10% glycerol, 50 mM NaF, 5 mM NaPPi, 1 mM PMSF, 0.5 mM DTT, 200 mM Na3VO4, Sigma protease and phosphatase inhibitor solutions). Lysate was cleared by centrifugation before analysis by SDS-PAGE. Antibodies used include: Human KRAS (Sigma, WH0003845M1); DNMT1 (Cell Signaling, 5032); p21 (BD Pharmingen, 554228); and Beta-actin (Sigma, clone AC-74); pERK1/2 (Cell Signaling, 9101); Rb (Santa Cruz, sc-50); GAPDH (Santa Cruz, sc-47724); c-Myc (Santa Cruz, sc-40); and EPHA2 (Cell Signaling, 12927). Western blot quantitation was performed with ImageJ software [64].

### Cloning of DNMT1 mutants

A TrueORF clone of DNMT1 cDNA, transcript variant 1, was obtained from OriGene (RC226414L1). Site-specific mutagenesis to generate S713D and S713A mutants was performed by PCR-directed mutagenesis as previously described [65]. Wildtype and mutant ORFs were shuttled into Gateway pENTR3C vector (Invitrogen, 11817-012) followed by pLenti-CMV-Puro DEST vector (Addgene, w118-1). The plasmids were propagated in DH5α bacterial cells (Life Technologies). The constructs were verified by sequencing. PCR primers for site-specific mutagenesis are listed in Additional File 4.

### Quantitative RT-PCR

Total RNA was isolated using Trizol reagent and purified with Qiagen RNeasy kit. cDNA was generated using Superscript First-Strand (Invitrogen) synthesis and real-time PCR was performed using SYBR Green PCR Master Mix (Applied Biosystems). Primers for DNMT1 target genes are listed in Additional File 4. All experiments were conducted with at least 3 technical replicates. Statistical significance was determined by unpaired t-tests, where n.s. = *p* ≥ 0.05; * = *p* ≤ 0.01; ** = *p* ≤ 0.001.

### Protein extraction and tryptic digests

For mass spectrometry cells were collected by scraping, brief centrifugation and snap-freezing in liquid nitrogen. Frozen cell pellets containing approximately 1×10e8 cells were resuspended in 0.5 ml of TRIzol reagent (Thermo Fisher Scientific) and protein extraction was carried out according to the standard manufacturer’s protocol. Briefly, 0.2 ml of chloroform was added per 1 ml of TRIzol. Precipitated protein pellets were washed with ethanol and vacuum-dried briefly. The pellets were reconstituted in 300 μL of 8 M urea, 0.2 % Zwittergent 3-16 (Santa Cruz Biotechnology), and 50 mM Tris-HCl, pH 8.0. Twelve μL of 100 mM DTT were added and samples vortexed for 30 min at room temperature. The pellets were dissolved by microtip soniction (30 sec on/30 sec off on ice x3). Eighteen μL of 100 mM iodoacetamide was added, followed by water bath sonication for 30 min at room temperature. Two μL of resuspended protein sample were quantified using Pierce BCA protein assay kit (Thermo Fisher Scientific) according to the standard manufacturer’s protocol. Sample was diluted by adding 1.2 mL of 50 mM Tris-HCl, 1 mM CaCl2 (pH 8.0), and pH was adjusted to 8.0. Sequencing grade trypsin (Promega) was added at a ratio of 1:50 (trypsin/protein), and the samples were incubated with gentle agitation at 37°C overnight. Approximately 10 μL of formic acid per 1 mL of sample was added to adjust pH to ∼3.5. Digested peptides were desalted with Sep-Pak C18 cartridges (Waters) according to the manufacturer’s protocol, dried to remove solvent under vacuum centrifugation, and stored at -80°C for further use.

### Titanium dioxide phosphopeptide enrichment

Phosphopeptide enrichment was performed as previously described [66]. Briefly, 4 mg of trypsin digested peptides were separately processed on an AKTA Purifier UPLC system (GE Healthcare) fitted with an in-house column packed with 5-μm TiO2 beads (GL Sciences, Tokyo, Japan). The eluates containing mostly phosphopeptides (typically >80%) were pooled and desalted with Millipore C18 ZipTips (Sigma-Aldrich) according to the manufacturer’s protocol. Similarly, flow-through fractions were pooled and desalted with Sep-Pak C18 cartridges (Waters) according to the manufacturer’s protocol. Phosphopeptide amounts in the eluates were estimated at 1% of input material. Both samples where dried to remove solvent under vacuum centrifugation, resuspended in 0.1% formic acid at 0.1 ug per μL, and stored at −20°C until MS analysis.

### SRM mass spectrometry

SRM was performed on Waters nanoAcquity UPLC system (Waters) online with AB Sciex Qtrap 5500 mass spectrometer (AB Sciex). Digested peptide samples were loaded at 0.5 µg per injection and separated using two-buffer reverse phase chromatography (Buffer A = 0.1% formic acid in water; Buffer B = 0.1% formic acid in acetonitrile). The flow was equilibrated at 3% B, and peptides were loaded on nanoACQUITY UPLC Symmetry C18 trap 180 µm x 20 mm, 100Å, 5 µm, column (Waters) for 2 minutes at 5 µL/min and then eluted through nanoACQUITY UPLC BEH C18 analytical 100 µm x 100 mm, 130Å, 1.7 µm column (Waters) at 1 µL/min as follows: 3 – 25% B over 70 min; 25 – 50% B over 4 min; 50 – 90% B over 4 min; 90% B for 2 min; and 3% B for 10 min. Qtrap 5500 instrument (AB Sciex) fitted with a Nanospray III source (AB Sciex) was used in MRM mode with the following parameters: positive polarity; Q1 resolution = Unit; Q3 resolution = Unit; ion spray voltage = 3.5 KV. For unscheduled SRM runs, a dwell time of 20 ms was set for each transition. In scheduled SRM runs, a total cycle time of 3.7 – 4.5 sec was used. Collision energies were calculated by Skyline [67] using the standard AB Sciex Qtrap 5500 setting.

### SRM spectral libraries

A combination of spectral libraries was used as a source of reference spectra for the development of SRM methods in this study. This included in-house spectral libraries generated from a large-scale peptide discovery in a set of 8 human lung adenocarcinoma cell lines. Approximately 10e9 cells from each line were used in this effort. The cells were lysed, protein extracted using TRIzol reagent, and digested with trypsin as described above. The digested peptide samples from individual lines were pooled, and the pool was subjected to titanium dioxide phosphopeptide enrichment in 2-mg aliquots as described above. The phosphopeptide enriched eluates and flow-through factions were then pooled, and the resulting two pools were subjected to off-line high-pH reverse phase fractionation into 85 fractions each as previously described [68]. Each fraction was then analyzed by LC-MS/MS on LTQ Orbitrap Velos (Thermo Fisher Scientific). Flow-through factions were analyzed using 2-hr HCD runs, while phosphopeptide fractions were analyzed using 2-hr HCD and CID runs. The obtained spectra were searched with Protein Prospector [69] version 5.10.1 with the default parameters, including trypsin as the protease, up to one allowed missed cleavage site, Carbamidomethyl-C constant modification, default variable modifications with or without “Phospho (STY)”, up to 3 modifications per peptide, 20 ppm precursor mass accuracy, and either 20 ppm (HCD) or 0.6 Da fragment mass accuracy (CID). The resulting high confidence spectra (FDR <1%) were compiled into three corresponding retention-time normalized *blib*-formatted libraries using Protein Prospector. Raw data files, Protein Prospector search results, and the resulting spectral libraries have been uploaded to Proteome Exchange repository [70] via MassIVE. An additional spectral library of predominantly unmodified peptides was constructed from a published large-scale discovery experiment in 11 diverse human cell lines conducted by Geiger, et al. [71]. Raw data from this study (HCD spectra collected on LTQ Orbitrap Velos) were re-searched, and high confidence spectra (FDR <1%) were compiled into a *blib*-formatted library in Protein Prospector using the same parameters as above.

### SRM method development

A list of RAS network target proteins (Additional File 1) was compiled from Reactome pathways database (www.Reactome.org) [72], including pathways R-HSA-5683057.1, R-HSA-177929.1, R-HSA-1489509.1, and R-HSA-1257604.1, and from a custom list of proteins of interest to our group; a set of endogenous target peptides used as sensitivity controls and retention time standards was also included. All target scheduling, SRM method refinement, and SRM data quantitation were performed in Skyline [67]. Initial method for protein abundance quantitation targeted 601 unmodified peptide precursors from 197 proteins, which was split into 22 unscheduled 90-min injections and verified on a pooled sample of TiO_2_ flow-through fractions. In this method, up to 8 peptides per target protein were selected from the spectral libraries, and up to 7 highest intensity y- and b-ion transitions between 350 and 1200 m/z units were included. Similarly, initial method for phosphopeptide quantitation targeted 374 phosphorylated peptide precursors from 122 proteins, which was split into 13 unscheduled injections and verified on a pooled sample of TiO_2_ enriched fractions. This method included all phosphorylated peptides belonging to the target proteins identified in the discovery data. All peaks assigned by Skyline were manually checked for accuracy. In general, only peaks within +/-10 min of predicted retention times, library dot product >0.45, and at least 3 transitions detected were retained. The final protein abundance method targeted 365 peptide precursors from 159 proteins, used +/-15 min detection windows and was split into 4 injections, while the final phosphopeptide method targeted 348 peptide precursors from 114 proteins and was split into 3 injections. Target peptides were checked for uniqueness by BLAST [73] against the SwissProt human proteome database, and only unique peptides were further considered.

A set of 52 unmodified and 50 phosphorylated SIL peptides, which primarily included peptides that demonstrated the largest changes in the experimental runs, were synthesized at JPT technologies (Berlin, Germany) and were examined side-by-side along the endogenously detected targets (Supplementary Figure S3). Forty six of 52 unmodified and 48 of 50 phosphorylated SIL peptides had essentially identical retention times and relative transition intensities to the endogenous counterparts, confirming their specific detection. False discovery rate was estimated as the fraction of the target peptides, whose retention times and/or spectral profiles did not match those of the SIL standards.

### Experimental design and statistical rationale

Each experimental condition (control, MEK-i, and si-KRAS) was evaluated by SRM using three replicate measurements. Each condition was represented by 4 independently treated cultures of H358 cells, which were pooled to adjust for biological variability and to achieve sufficient quantities of starting material for downstream phosphoenrichment, SRM method development, and multi-injection data acquisition runs. Although the lack of true biological repeat measurements is a limitation of the study, this experimental design was justified as the SRM experiments were envisioned to be a hypothesis-generating screen for further functional validation rather than a stand-alone dataset. The SRM experiments did not employ synthetic peptides during data acquisition and thus should be considered a Tier 3 analysis [74].

### Data analysis

Raw data files were imported into Skyline, and all peaks were manually checked for accuracy as above. Total peak areas (sum of individual transitions) were calculated by Skyline and exported for statistical analysis in R (www.R-project.org). In general, we followed a widely used workflow for data analysis in proteomics based on calculating log2 intensity ratios, followed by normalization via centering medians, aggregation of ratios using medians, and significance testing with the Student’s t-test [74–76]. All quantified Peak areas of the experimental conditions (si-KRAS and MEK-i) were converted to log2 ratios relative to control. Replicate runs were normalized by 0-centering the medians of the log2 ratio distributions. For each quantified peptide, an aggregate median ratio, corresponding standard deviation, and two-tailed one-sample t-test p-value (H_0_: μ=0) within replicate runs were calculated. For unmodified peptides, if more than one peptide per protein were quantified, an aggregate ratio was calculated using p-value weighting to give higher consideration to low-variance (i.e. less noisy) peptides [m’ = Σ(m(1-p))/Σ(1-p)]. For proteins with both phosphorylation and protein abundance measurements, measured phosphopeptide log2 ratios were adjusted by the corresponding aggregate protein abundance log2 ratios [log2_Net_ = log2_Phos._ – log2_Abund._]. Statistical significance of the observed changes in protein abundance (Figure 1A) was estimated using two-tailed one-sample t-test (H_0_: μ=0) by treating log2 ratios of all measured peptides within a given protein in all replicates as independent observations. Statistical significance of phosphorylation changes in proteins for which abundance measurements were not available (Figure 1C) was estimated for each phosphorylated peptide using two-tailed one-sample t-test p-value (H_0_: μ=0) as described above. Statistical significance of net (abundance-adjusted) phosphorylation changes (Figure 1B) was estimated using pt() function in R of *t-*distribution, with n=3, df=2, and variance estimated as either the standard deviation of measured abundance ratios or phosphorylation ratios, whichever was higher (i.e. the most conservative estimate to avoid false positives). Tables of calculated abundance and phosphorylation measurements are available as Additional Files 5 and 6, respectively.

Regulation index (RI) for each quantified protein was defined as the sum of the absolute values of the aggregated abundance log2 ratio (if measured) and phosphorylation log2 ratios of all measured phosphopeptides from that protein. Only ratios greater than 0.5 were included. Three different RI indexes were calculated: MEK-i vs control, si-KRAS vs control, and si-KRAS vs MEK-i. Eighty five proteins had non-zero RI in at least one of the three comparisons and were thus considered regulated. An interaction networks of these 85 proteins (Figures 2 and 3) was constructed in Cytoscape v. 3.0.1 [77] using functional interactions annotated in Reactome FI database (www.Reactome.org). The following criteria were used to classify regulated proteins into one of four regulated classes (Figure 3): (i) si-KRAS specific regulation: RI_si-KRAS vs control_ > 0.5 and RI_MEK-i vs control_ < 0.5; (ii) MEK-i specific regulation: RI_MEK-i vs control_ > 0.5 and RI_si-KRAS vs control_ < 0.5; (iii) concordant regulation: RI_MEK-i vs control_ > 0.5 and RI_si-KRAS vs control_ > 0.5 and RI_si-_ KRAS vs MEK-i < 0.5; and (iv) orthogonal regulation: RI_si-KRAS vs MEK-i_ > 0.5. In this figure, the magnitude of regulation was calculated as RIsi-KRAS vs control, RIMEK-i vs control, average of RIMEK-i vs control and RIsi-KRAS vs control, and RI_si-KRAS vs MEK-i_, respectively. A table of the calculated RI values is available as Additional File 7.

Interaction network of all quantified proteins (Figure 4) was constructed in Cytoscape using functional interactions from Reactome FI database. Individual protein nodes were grouped manually into signaling subnetworks or functional categories. Circle plots to visualize abundance and phosphorylation measurements in individual proteins were constructed in Cytoscape using the enhancedGraphics app [78].

## Acknowledgements

We thank Graham Johnson, UCSF, for helpful suggestions on data visualization.

## Availability of data and materials

MS peptide and phosphopeptide discovery data, including raw data files, peaklist files, search engine result files, and generated SRM BLIB-formatted libraries are available via Proteome Exchange and MassIVE public repositories (accessions MSV000082014, MSV000082016, MSV000082018). Skyline documents containing SRM data are available for public access and download on Panorama Web [https://panoramaweb.org/labkey/project/UCSF-MSF/KRAS%20Network/begin.view<u>?</u>]. All relevant calculated data generated during this study are included in this published article and its supplementary information files. Requests for any additional data should be directed to the corresponding author.

## Authors’ contributions

AU, TLY, ALB and FM designed the experiments. AU, TYL, MT, SA, JAOP, CDR and MSZ conducted the experiments. AU, TLY, JHM and CHB analyzed the data. AU and TLY wrote the manuscript. ALB and FM contributed comments on the manuscript.

## Ethics approval and consent to participate

No human subjects or experiments involving animals were used in this study.

## Consent for publication

No human subjects were used in this study.

## Additional files

**File 1** (MS Excel): Initial list of RAS network protein targets.

**File 2** (MS Excel): Lung adenocarcinoma cell lines used in peptide/phosphopeptide discovery.

**File 3** (PDF): Abundance and site-specific phosphorylation in all measured proteins.

**File 4** (MS Excel): PCR primers and siRNA sequences.

**File 5** (MS Excel): Table of calculated abundance values and estimated significance.

**File 6** (MS Excel): Table of calculated phosphorylation values and estimated significance.

**File 7** (MS Excel): Table of calculated regulation index values.

---

**Figure S1.**
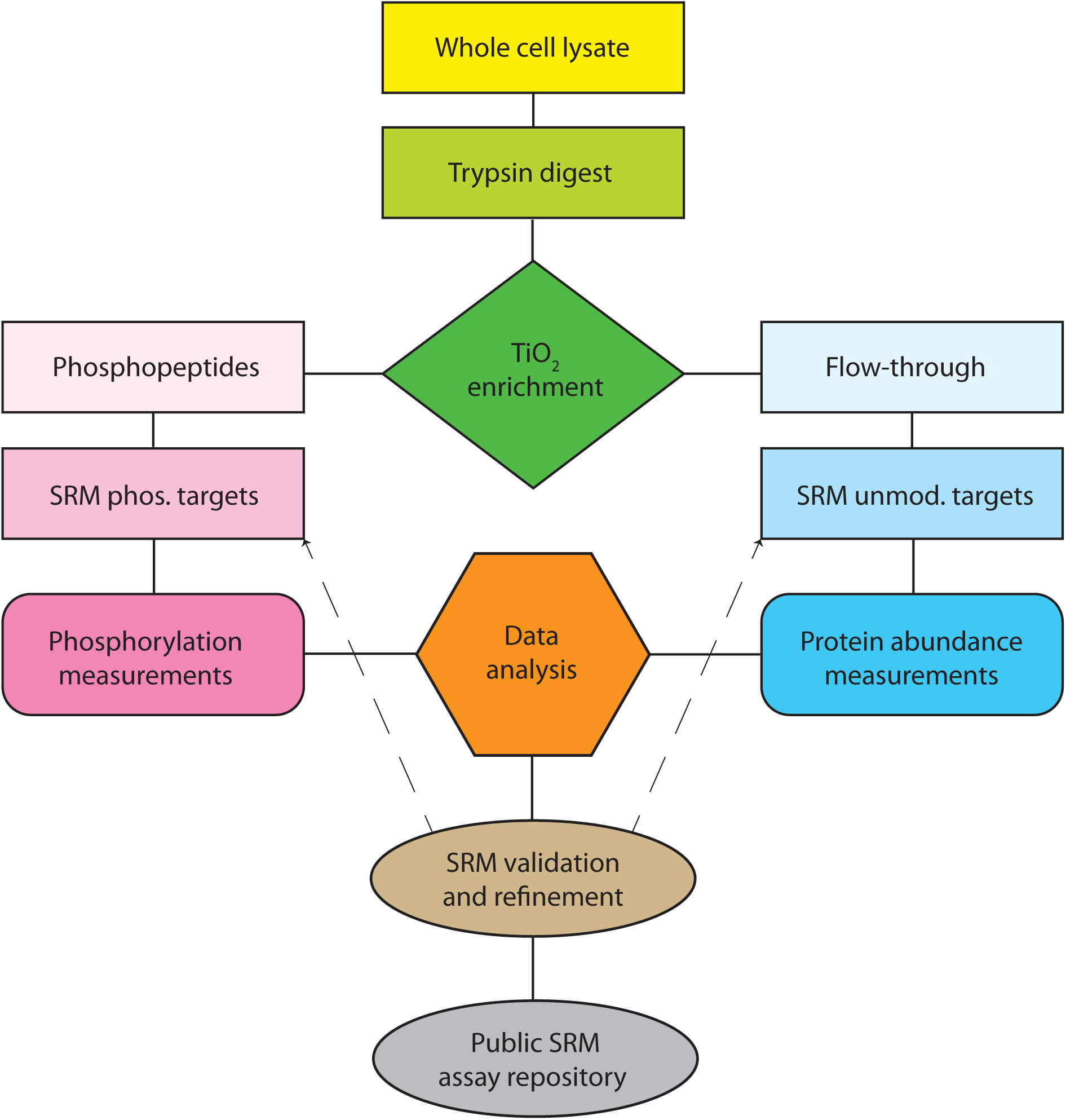
SRM-based targeted phospho-proteomic workflow. Whole cell lysates are trypsinized and subjected to titanium dioxide phosphopeptide affi nity enrichment. Eluate and flow-through fractions are separately analyzed by SRM to quantify target phosphopeptides (site-specific phosphorylation) and unmodified peptides (protein abundance), respectively. Validated SRM methods have been deposited to Panorama for public access.

**Figure S2.**
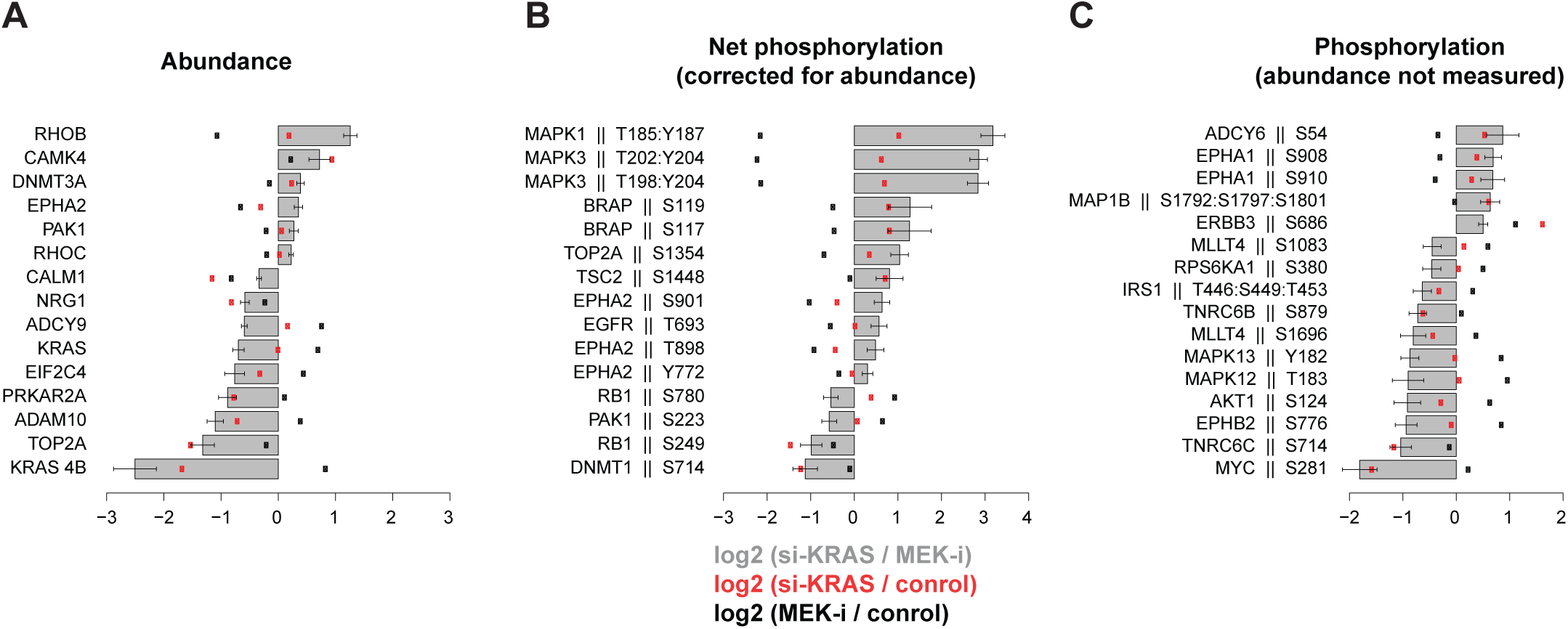
Most significant abundance and phosphorylation changes between si-KRAS and MEK-i conditions. Log2 ratios of si-KRAS vs. MEK-i were calculated (grey bars) and their statistical significance estimated by t-test. The most significant (p-value <0.05 and log2 change of >0.5) are plotted. Ratios of si-KRAS and MEK-i relative to control are shown as red and black dots, respectively. Whiskers reflect standard deviations in replicate measurements. (**A**) Abundance measurements for each protein reflect log2 ratios of p-value-weighed medians of multiple peptides. (**B**) Median log2 ratios of individual phosphorylated peptides corrected by protein abundance. (**C**) Median log2 ratios of individual phosphorylated peptides for which corresponding protein abundance measurements could not be measured.

**Figure S3.**
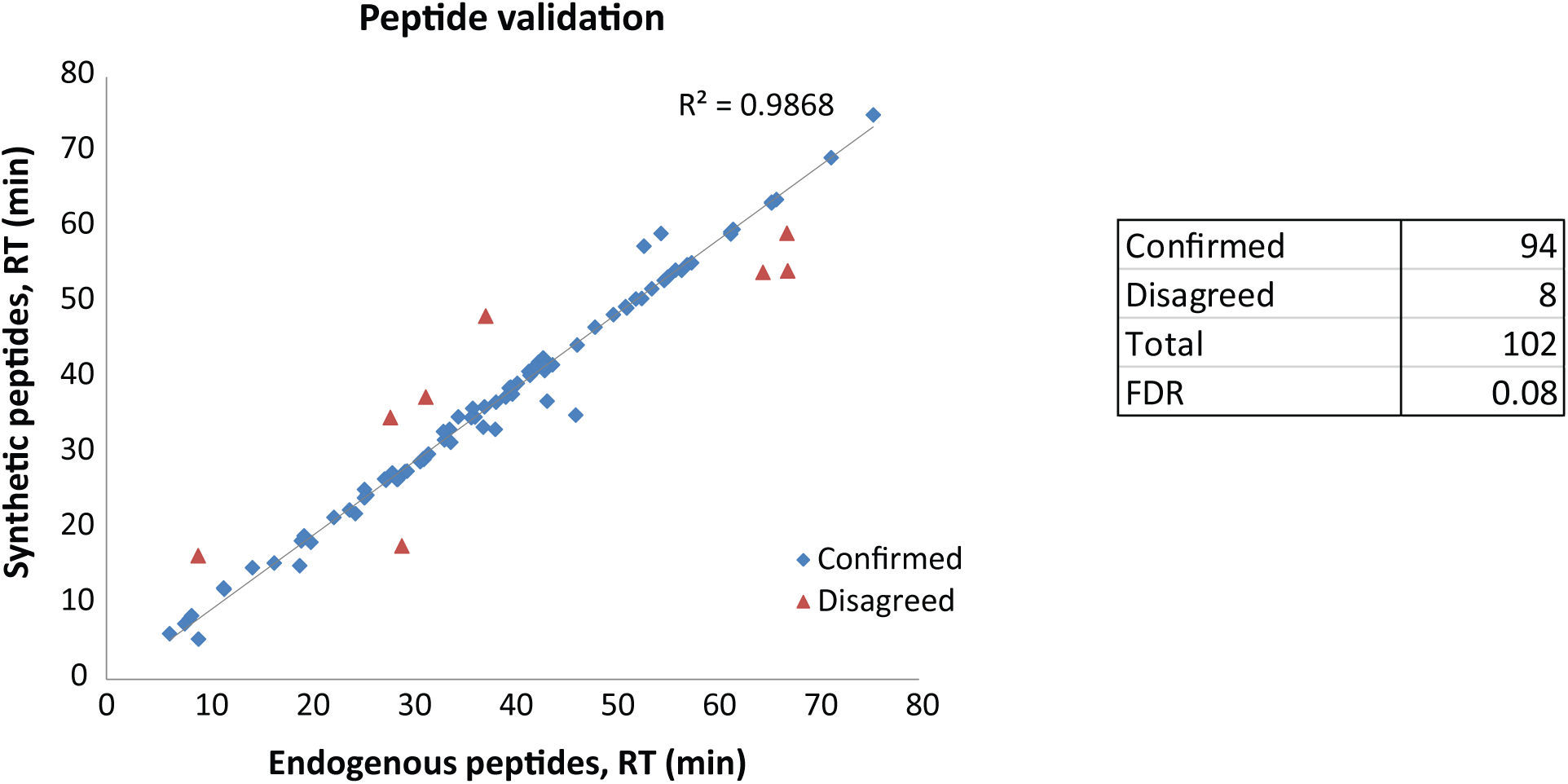
Estimates of false discovery rate due to non-specific SRM peptide detection. A total of 102 isotope labeled synthetic peptides (52 unmodified and 50 phosphorylated), chiefly representing those with the largest changes in the experimental runs, were analyzed by SRM and compared to their endogenous counterparts. Peptides were considered correctly identified (confirmed) if there was close agreement in retention times and relative transition intensities.

**Figure S4.**
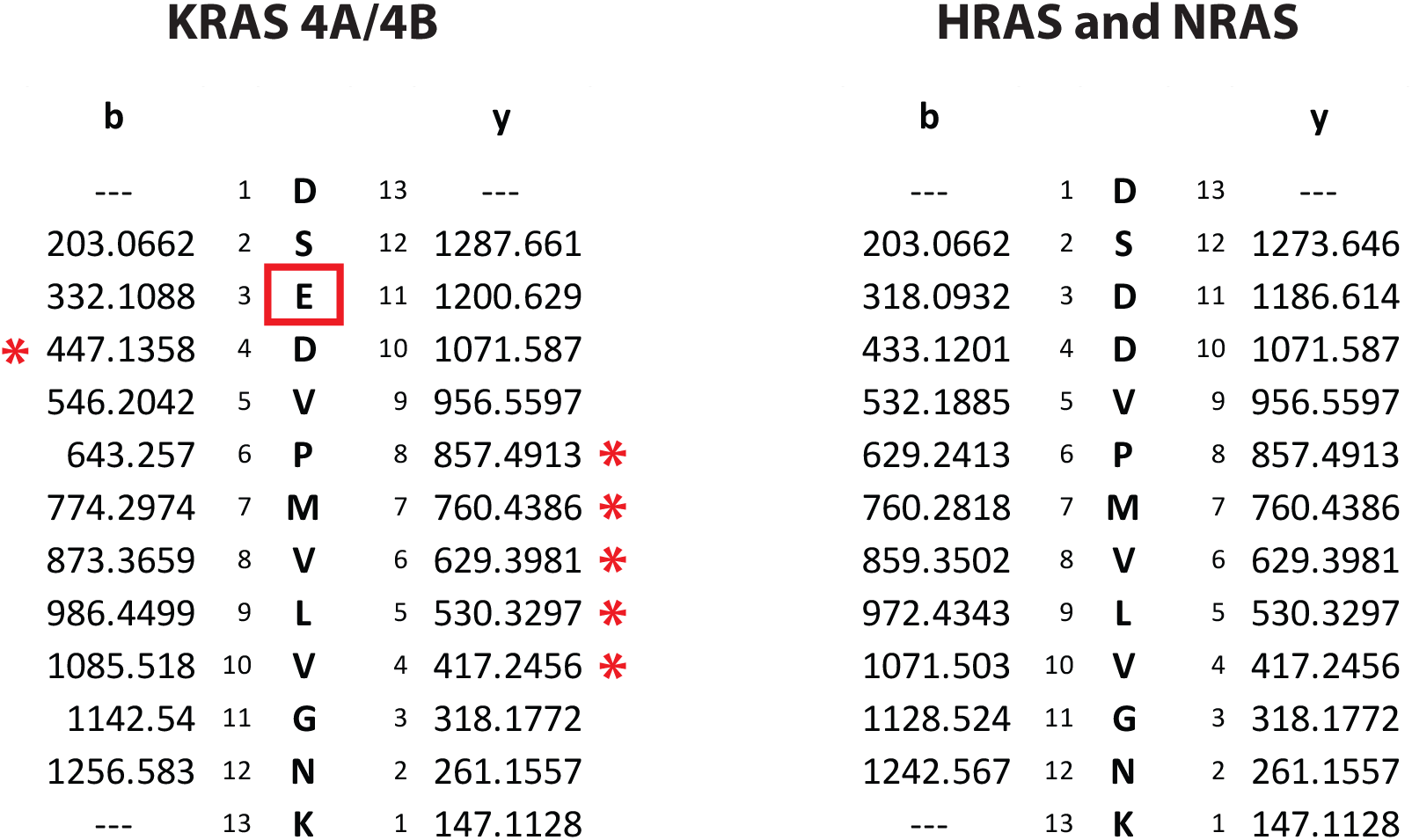
KRAS peptide used to detect both 4A and 4B isoforms shares all but one residue (red box) with a homologous peptide in HRAS and NRAS. Predicted y and b fragment ion series are shown. Red asterisks denote ions monitored by SRM.

**Figure S5.**
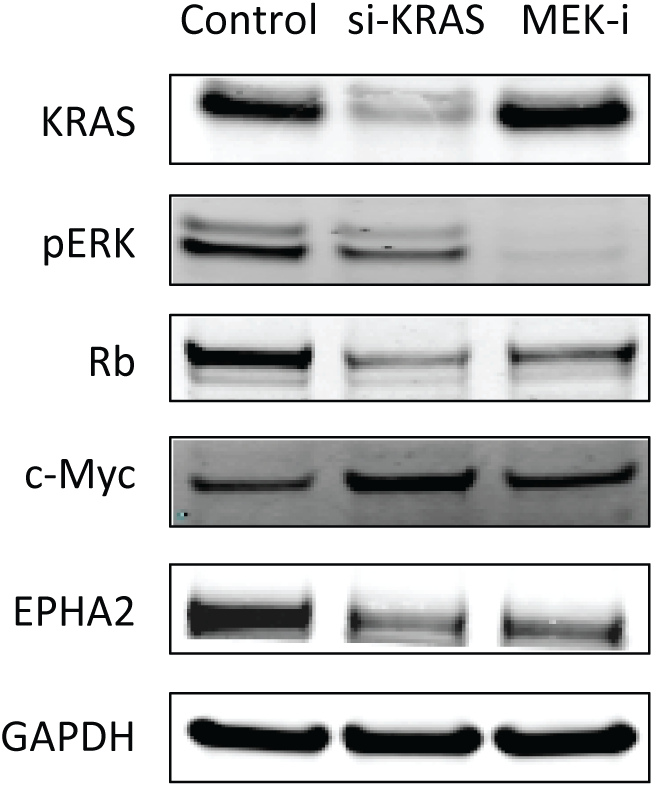
Western blots support SRM findings. Cells were treated with si-KRAS, MEK-i. or control siRNA/DMSO. Protein lysates were separated by SDS-PAGE, blotted, and probed by KRAS, pERK1/2, Rb, c-Myc, EPHA2, and GPDH specific antibodies.

**Figure S6.**
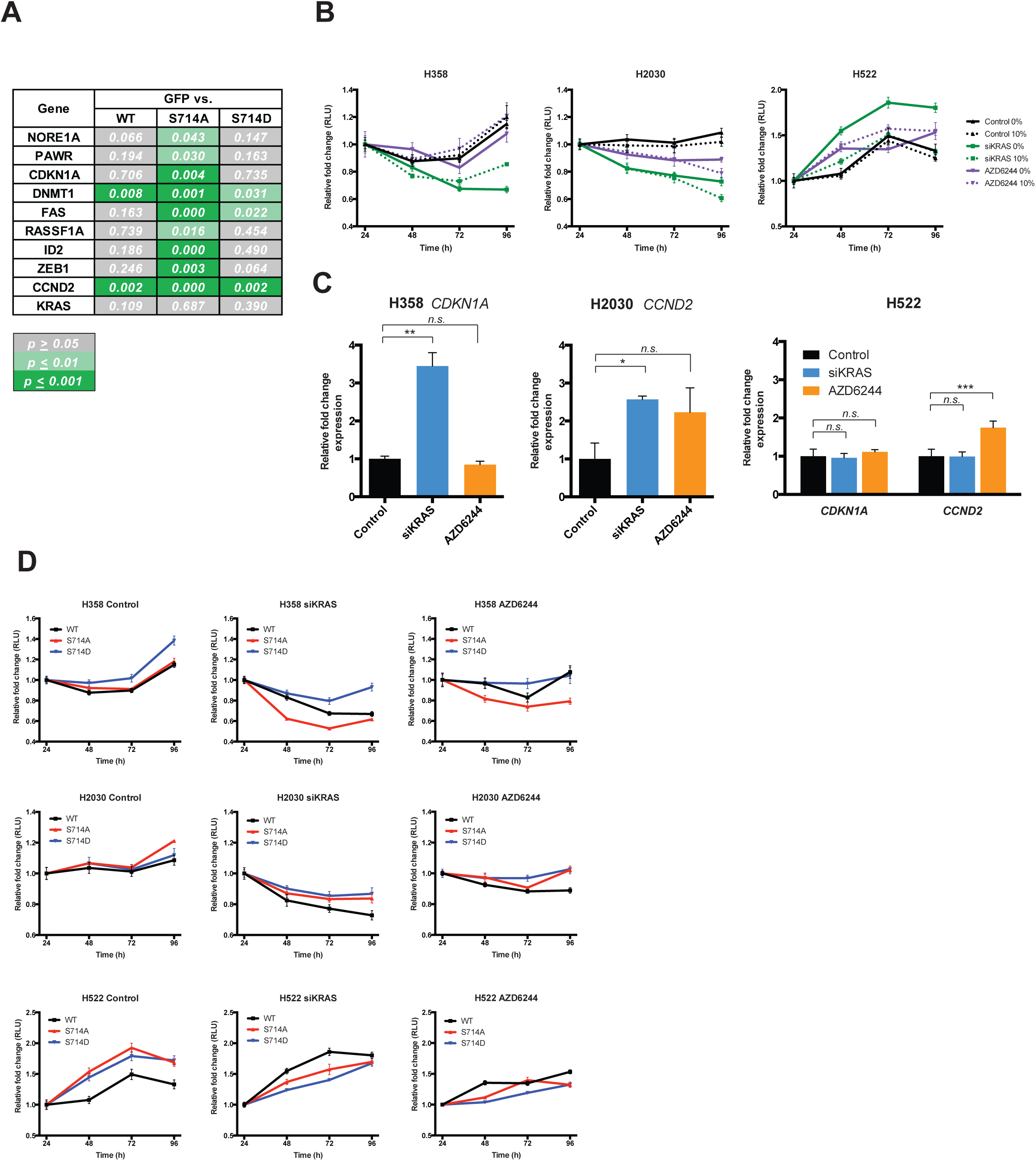
KRAS modulates phosphorylation of DNMT1 in a MEK-independent manner. MEK inhibition does not mimic KRAS knock­ down in the regulation of DNMT1 target genes. (**A**) Statistical significance of gene expression between H358 cells expressing GFP and DNMT1 WT, S714A or S714D mutants. Experiments were run in technical triplicates, and p-values from unpaired t-tests are shown. (**B**) Growth curves were generated for KRAS mutant and KRAS wild-type lines, treated with negative control siRNA+DMSO, siKRAS or S00nM AZD6244 in 10% or 0% serum. AZD6244 treatment only moderately affected proliferation in H2030 cells, and neither KRAS knockdown nor AZD6244 affected H522 cells. (**C**) AZD6244 treatment did not induce CDKN1A levels in H358 cells, consis­ tent with no change in cell proliferation. CCND2 levels were only moderately elevated in H2030 and H522 cells. Experiments were run in techni­ cal triplicates, and p-values from unpaired t-tests are shown. (**D**) Effects on proliferation by DNMT1 mutants were moderate under AZD6244 treatment compared to siKRAS. n.s. = *p*≥0.05.; * = *p*≤0.01; ** = *p*≤0.001

## References

1. Borras E, Jurado I, Hernan I, Gamundi MJ, Dias M, Marti I, Mane B, Arcusa A, Agundez JA, Blanca M, Carballo M: Clinical pharmacogenomic testing of KRAS, BRAF and EGFR mutations by high resolution melting analysis and ultra-deep pyrosequencing. BMC Cancer 2011, 11:406.

2. Imielinski M, Berger AH, Hammerman PS, Hernandez B, Pugh TJ, Hodis E, Cho J, Suh J, Capelletti M, Sivachenko A, et al: Mapping the hallmarks of lung adenocarcinoma with massively parallel sequencing. Cell 2012, 150:1107–1120.

3. Marks JL, Broderick S, Zhou Q, Chitale D, Li AR, Zakowski MF, Kris MG, Rusch VW, Azzoli CG, Seshan VE, et al: Prognostic and therapeutic implications of EGFR and KRAS mutations in resected lung adenocarcinoma. J Thorac Oncol 2008, 3:111–116.

4. De Roock W, Jonker DJ, Di Nicolantonio F, Sartore-Bianchi A, Tu D, Siena S, Lamba S, Arena S, Frattini M, Piessevaux H, et al: Association of KRAS p.G13D mutation with outcome in patients with chemotherapy-refractory metastatic colorectal cancer treated with cetuximab. JAMA 2010, 304:1812–1820.

5. Suda K, Tomizawa K, Mitsudomi T: Biological and clinical significance of KRAS mutations in lung cancer: an oncogenic driver that contrasts with EGFR mutation. Cancer Metastasis Rev 2010, 29:49–60.

6. McCormick F: KRAS as a Therapeutic Target. Clin Cancer Res 2015, 21:1797–1801.

7. Ribas A, Flaherty KT: BRAF targeted therapy changes the treatment paradigm in melanoma. Nat Rev Clin Oncol 2011, 8:426–433.

8. Caunt CJ, Sale MJ, Smith PD, Cook SJ: MEK1 and MEK2 inhibitors and cancer therapy: the long and winding road. Nat Rev Cancer 2015, 15:577–592.

9. Turke AB, Song Y, Costa C, Cook R, Arteaga CL, Asara JM, Engelman JA: MEK inhibition leads to PI3K/AKT activation by relieving a negative feedback on ERBB receptors. Cancer Res 2012, 72:3228–3237.

10. Hatzivassiliou G, Haling JR, Chen H, Song K, Price S, Heald R, Hewitt JF, Zak M, Peck A, Orr C, et al: Mechanism of MEK inhibition determines efficacy in mutant KRAS-versus BRAF-driven cancers. Nature 2013, 501:232–236.

11. Carriere A, Romeo Y, Acosta-Jaquez HA, Moreau J, Bonneil E, Thibault P, Fingar DC, Roux PP: ERK1/2 phosphorylate Raptor to promote Ras-dependent activation of mTOR complex 1 (mTORC1). J Biol Chem 2011, 286:567–577.

12. Lange V, Picotti P, Domon B, Aebersold R: Selected reaction monitoring for quantitative proteomics: a tutorial. Mol Syst Biol 2008, 4:222.

13. Wu G, Feng X, Stein L: A human functional protein interaction network and its application to cancer data analysis. Genome Biol 2010, 11:R53.

14. Hmitou I, Druillennec S, Valluet A, Peyssonnaux C, Eychene A: Differential regulation of B-raf isoforms by phosphorylation and autoinhibitory mechanisms. Mol Cell Biol 2007, 27:31–43.

15. Zimmermann S, Moelling K: Phosphorylation and regulation of Raf by Akt (protein kinase B). Science 1999, 286:1741–1744.

16. Dougherty MK, Muller J, Ritt DA, Zhou M, Zhou XZ, Copeland TD, Conrads TP, Veenstra TD, Lu KP, Morrison DK: Regulation of Raf-1 by direct feedback phosphorylation. Mol Cell 2005, 17:215–224.

17. Sharma P, Veeranna, Sharma M, Amin ND, Sihag RK, Grant P, Ahn N, Kulkarni AB, Pant HC: Phosphorylation of MEK1 by cdk5/p35 down-regulates the mitogen-activated protein kinase pathway. J Biol Chem 2002, 277:528–534.

18. Banks AS, McAllister FE, Camporez JP, Zushin PJ, Jurczak MJ, Laznik-Bogoslavski D, Shulman GI, Gygi SP, Spiegelman BM: An ERK/Cdk5 axis controls the diabetogenic actions of PPARgamma. Nature 2015, 517:391–395.

19. Joshi J, Fernandez-Marcos PJ, Galvez A, Amanchy R, Linares JF, Duran A, Pathrose P, Leitges M, Canamero M, Collado M, et al: Par-4 inhibits Akt and suppresses Ras-induced lung tumorigenesis. EMBO J 2008, 27:2181–2193.

20. Sunaga N, Shames DS, Girard L, Peyton M, Larsen JE, Imai H, Soh J, Sato M, Yanagitani N, Kaira K, et al: Knockdown of oncogenic KRAS in non-small cell lung cancers suppresses tumor growth and sensitizes tumor cells to targeted therapy. Mol Cancer Ther 2011, 10:336–346.

21. Valentino JD, Elliott VA, Zaytseva YY, Rychahou PG, Mustain WC, Wang C, Gao T, Evers BM: Novel small interfering RNA cotargeting strategy as treatment for colorectal cancer. Surgery 2012, 152:277–285.

22. Inoki K, Zhu T, Guan KL: TSC2 mediates cellular energy response to control cell growth and survival. Cell 2003, 115:577–590.

23. Inoki K, Ouyang H, Li Y, Guan KL: Signaling by target of rapamycin proteins in cell growth control. Microbiol Mol Biol Rev 2005, 69:79–100.

24. Sonntag AG, Dalle Pezze P, Shanley DP, Thedieck K: A modelling-experimental approach reveals insulin receptor substrate (IRS)-dependent regulation of adenosine monosphosphate-dependent kinase (AMPK) by insulin. FEBS J 2012, 279:3314–3328.

25. Chen CH, Shaikenov T, Peterson TR, Aimbetov R, Bissenbaev AK, Lee SW, Wu J, Lin HK, Sarbassov dos D: ER stress inhibits mTORC2 and Akt signaling through GSK-3beta-mediated phosphorylation of rictor. Sci Signal 2011, 4:ra10.

26. Meng J, Peng H, Dai B, Guo W, Wang L, Ji L, Minna JD, Chresta CM, Smith PD, Fang B, Roth JA: High level of AKT activity is associated with resistance to MEK inhibitor AZD6244 (ARRY-142886). Cancer Biol Ther 2009, 8:2073–2080.

27. Gao X, Chaturvedi D, Patel TB: Localization and retention of p90 ribosomal S6 kinase 1 in the nucleus: implications for its function. Mol Biol Cell 2012, 23:503–515.

28. Carriere A, Cargnello M, Julien LA, Gao H, Bonneil E, Thibault P, Roux PP: Oncogenic MAPK signaling stimulates mTORC1 activity by promoting RSK-mediated raptor phosphorylation. Curr Biol 2008, 18:1269–1277.

29. Assiddiq BF, Tan KY, Toy W, Chan SP, Chong PK, Lim YP: EGFR S1166 phosphorylation induced by a combination of EGF and gefitinib has a potentially negative impact on lung cancer cell growth. J Proteome Res 2012, 11:4110–4119.

30. Imami K, Sugiyama N, Imamura H, Wakabayashi M, Tomita M, Taniguchi M, Ueno T, Toi M, Ishihama Y: Temporal profiling of lapatinib-suppressed phosphorylation signals in EGFR/HER2 pathways. Mol Cell Proteomics 2012, 11:1741–1757.

31. Luo X, Prior M, He W, Hu X, Tang X, Shen W, Yadav S, Kiryu-Seo S, Miller R, Trapp BD, Yan R: Cleavage of neuregulin-1 by BACE1 or ADAM10 protein produces differential effects on myelination. J Biol Chem 2011, 286:23967–23974.

32. Villalonga P, Lopez-Alcala C, Bosch M, Chiloeches A, Rocamora N, Gil J, Marais R, Marshall CJ, Bachs O, Agell N: Calmodulin binds to K-Ras, but not to H- or N-Ras, and modulates its downstream signaling. Mol Cell Biol 2001, 21:7345–7354.

33. Moreto J, Vidal-Quadras M, Pol A, Santos E, Grewal T, Enrich C, Tebar F: Differential involvement of H- and K-Ras in Raf-1 activation determines the role of calmodulin in MAPK signaling. Cell Signal 2009, 21:1827–1836.

34. Alvarez-Moya B, Lopez-Alcala C, Drosten M, Bachs O, Agell N: K-Ras4B phosphorylation at Ser181 is inhibited by calmodulin and modulates K-Ras activity and function. Oncogene 2010, 29:5911–5922.

35. Bivona TG, Quatela SE, Bodemann BO, Ahearn IM, Soskis MJ, Mor A, Miura J, Wiener HH, Wright L, Saba SG, et al: PKC regulates a farnesyl-electrostatic switch on K-Ras that promotes its association with Bcl-XL on mitochondria and induces apoptosis. Mol Cell 2006, 21:481–493.

36. Wang MT, Holderfield M, Galeas J, Delrosario R, To MD, Balmain A, McCormick F: K-Ras Promotes Tumorigenicity through Suppression of Non-canonical Wnt Signaling. Cell 2015, 163:1237–1251.

37. Huelsenbeck J, May M, Schulz F, Schelle I, Ronkina N, Hohenegger M, Fritz G, Just I, Gerhard R, Genth H: Cytoprotective effect of the small GTPase RhoB expressed upon treatment of fibroblasts with the Ras-glucosylating Clostridium sordellii lethal toxin. FEBS Lett 2012, 586:3665–3673.

38. Jiang K, Sun J, Cheng J, Djeu JY, Wei S, Sebti S: Akt mediates Ras downregulation of RhoB, a suppressor of transformation, invasion, and metastasis. Mol Cell Biol 2004, 24:5565–5576.

39. Dhillon AS, Hagan S, Rath O, Kolch W: MAP kinase signalling pathways in cancer. Oncogene 2007, 26:3279–3290.

40. Berra E, Diaz-Meco MT, Moscat J: The activation of p38 and apoptosis by the inhibition of Erk is antagonized by the phosphoinositide 3-kinase/Akt pathway. J Biol Chem 1998, 273:10792–10797.

41. Shin YJ, Kim YB, Kim JH: Protein kinase CK2 phosphorylates and activates p21-activated kinase 1. Mol Biol Cell 2013, 24:2990–2999.

42. Son J, Lyssiotis CA, Ying H, Wang X, Hua S, Ligorio M, Perera RM, Ferrone CR, Mullarky E, Shyh-Chang N, et al: Glutamine supports pancreatic cancer growth through a KRAS-regulated metabolic pathway. Nature 2013, 496:101–105.

43. Shanware NP, Bray K, Eng CH, Wang F, Follettie M, Myers J, Fantin VR, Abraham RT: Glutamine deprivation stimulates mTOR-JNK-dependent chemokine secretion. Nat Commun 2014, 5:4900.

44. McCleland ML, Adler AS, Deming L, Cosino E, Lee L, Blackwood EM, Solon M, Tao J, Li L, Shames D, et al: Lactate dehydrogenase B is required for the growth of KRAS-dependent lung adenocarcinomas. Clin Cancer Res 2013, 19:773–784.

45. Viale A, Pettazzoni P, Lyssiotis CA, Ying H, Sanchez N, Marchesini M, Carugo A, Green T, Seth S, Giuliani V, et al: Oncogene ablation-resistant pancreatic cancer cells depend on mitochondrial function. Nature 2014, 514:628–632.

46. Hassler M, Singh S, Yue WW, Luczynski M, Lakbir R, Sanchez-Sanchez F, Bader T, Pearl LH, Mittnacht S: Crystal structure of the retinoblastoma protein N domain provides insight into tumor suppression, ligand interaction, and holoprotein architecture. Mol Cell 2007, 28:371–385.

47. Ren S, Rollins BJ: Cyclin C/cdk3 promotes Rb-dependent G0 exit. Cell 2004, 117:239–251.

48. Padmanabhan A, Li X, Bieberich CJ: Protein kinase A regulates MYC protein through transcriptional and post-translational mechanisms in a catalytic subunit isoform-specific manner. J Biol Chem 2013, 288:14158–14169.

49. Soucek L, Whitfield J, Martins CP, Finch AJ, Murphy DJ, Sodir NM, Karnezis AN, Swigart LB, Nasi S, Evan GI: Modelling Myc inhibition as a cancer therapy. Nature 2008, 455:679–683.

50. Maddika S, Ande SR, Wiechec E, Hansen LL, Wesselborg S, Los M: Akt-mediated phosphorylation of CDK2 regulates its dual role in cell cycle progression and apoptosis. J Cell Sci 2008, 121:979–988.

51. Wells NJ, Hickson ID: Human topoisomerase II alpha is phosphorylated in a cell-cycle phase-dependent manner by a proline-directed kinase. Eur J Biochem 1995, 231:491–497.

52. Eletr ZM, Wilkinson KD: An emerging model for BAP1’s role in regulating cell cycle progression. Cell Biochem Biophys 2011, 60:3–11.

53. Matheny SA, Chen C, Kortum RL, Razidlo GL, Lewis RE, White MA: Ras regulates assembly of mitogenic signalling complexes through the effector protein IMP. Nature 2004, 427:256–260.

54. Sakamaki J, Daitoku H, Ueno K, Hagiwara A, Yamagata K, Fukamizu A: Arginine methylation of BCL-2 antagonist of cell death (BAD) counteracts its phosphorylation and inactivation by Akt. Proc Natl Acad Sci U S A 2011, 108:6085–6090.

55. Lazzaretti D, Tournier I, Izaurralde E: The C-terminal domains of human TNRC6A, TNRC6B, and TNRC6C silence bound transcripts independently of Argonaute proteins. RNA 2009, 15:1059–1066.

56. Serra RW, Fang M, Park SM, Hutchinson L, Green MR: A KRAS-directed transcriptional silencing pathway that mediates the CpG island methylator phenotype. Elife 2014, 3:e02313.

57. Stewart ML, Tamayo P, Wilson AJ, Wang S, Chang YM, Kim JW, Khabele D, Shamji AF, Schreiber SL: KRAS Genomic Status Predicts the Sensitivity of Ovarian Cancer Cells to Decitabine. Cancer Res 2015, 75:2897–2906.

58. Song J, Rechkoblit O, Bestor TH, Patel DJ: Structure of DNMT1-DNA complex reveals a role for autoinhibition in maintenance DNA methylation. Science 2011, 331:1036–1040.

59. Kim JY, Welsh EA, Fang B, Bai Y, Kinose F, Eschrich SA, Koomen JM, Haura EB: Phosphoproteomics Reveals MAPK Inhibitors Enhance MET- and EGFR-Driven AKT Signaling in KRAS-Mutant Lung Cancer. Mol Cancer Res 2016, 14:1019–1029.

60. Schweppe DK, Rigas JR, Gerber SA: Quantitative phosphoproteomic profiling of human non-small cell lung cancer tumors. J Proteomics 2013, 91:286–296.

61. Gstaiger M, Aebersold R: Applying mass spectrometry-based proteomics to genetics, genomics and network biology. Nat Rev Genet 2009, 10:617–627.

62. Doll S, Burlingame AL: Mass spectrometry-based detection and assignment of protein posttranslational modifications. ACS Chem Biol 2015, 10:63–71.

63. Yuan TL, Amzallag A, Bagni R, Yi M, Afghani S, Burgan W, Fer N, Strathern LA, Powell K, Smith B, et al: Differential Effector Engagement by Oncogenic KRAS. Cell Rep 2018, 22:1889–1902.

64. Rueden CT, Schindelin J, Hiner MC, DeZonia BE, Walter AE, Arena ET, Eliceiri KW: ImageJ2: ImageJ for the next generation of scientific image data. BMC Bioinformatics 2017, 18:529.

65. Zheng L, Baumann U, Reymond JL: An efficient one-step site-directed and site-saturation mutagenesis protocol. Nucleic Acids Res 2004, 32:e115.

66. Ruperez P, Gago-Martinez A, Burlingame AL, Oses-Prieto JA: Quantitative phosphoproteomic analysis reveals a role for serine and threonine kinases in the cytoskeletal reorganization in early T cell receptor activation in human primary T cells. Mol Cell Proteomics 2012, 11:171–186.

67. MacLean B, Tomazela DM, Shulman N, Chambers M, Finney GL, Frewen B, Kern R, Tabb DL, Liebler DC, MacCoss MJ: Skyline: an open source document editor for creating and analyzing targeted proteomics experiments. Bioinformatics 2010, 26:966–968.

68. Sos ML, Levin RS, Gordan JD, Oses-Prieto JA, Webber JT, Salt M, Hann B, Burlingame AL, McCormick F, Bandyopadhyay S, Shokat KM: Oncogene mimicry as a mechanism of primary resistance to BRAF inhibitors. Cell Rep 2014, 8:1037–1048.

69. Chalkley RJ, Baker PR, Huang L, Hansen KC, Allen NP, Rexach M, Burlingame AL: Comprehensive analysis of a multidimensional liquid chromatography mass spectrometry dataset acquired on a quadrupole selecting, quadrupole collision cell, time-of-flight mass spectrometer: II. New developments in Protein Prospector allow for reliable and comprehensive automatic analysis of large datasets. Mol Cell Proteomics 2005, 4:1194–1204.

70. Vizcaino JA, Deutsch EW, Wang R, Csordas A, Reisinger F, Rios D, Dianes JA, Sun Z, Farrah T, Bandeira N, et al: ProteomeXchange provides globally coordinated proteomics data submission and dissemination. Nat Biotechnol 2014, 32:223–226.

71. Geiger T, Wehner A, Schaab C, Cox J, Mann M: Comparative proteomic analysis of eleven common cell lines reveals ubiquitous but varying expression of most proteins. Mol Cell Proteomics 2012, 11:M111 014050.

72. Croft D, Mundo AF, Haw R, Milacic M, Weiser J, Wu G, Caudy M, Garapati P, Gillespie M, Kamdar MR, et al: The Reactome pathway knowledgebase. Nucleic Acids Res 2014, 42:D472–477.

73. Altschul SF, Gish W, Miller W, Myers EW, Lipman DJ: Basic local alignment search tool. J Mol Biol 1990, 215:403–410.

74. Carr SA, Abbatiello SE, Ackermann BL, Borchers C, Domon B, Deutsch EW, Grant RP, Hoofnagle AN, Huttenhain R, Koomen JM, et al: Targeted peptide measurements in biology and medicine: best practices for mass spectrometry-based assay development using a fit-for-purpose approach. Mol Cell Proteomics 2014, 13:907–917.

75. Chang CY, Picotti P, Huttenhain R, Heinzelmann-Schwarz V, Jovanovic M, Aebersold R, Vitek O: Protein significance analysis in selected reaction monitoring (SRM) measurements. Mol Cell Proteomics 2012, 11:M111 014662.

76. Bantscheff M, Lemeer S, Savitski MM, Kuster B: Quantitative mass spectrometry in proteomics: critical review update from 2007 to the present. Anal Bioanal Chem 2012, 404:939–965.

77. Shannon P, Markiel A, Ozier O, Baliga NS, Wang JT, Ramage D, Amin N, Schwikowski B, Ideker T: Cytoscape: a software environment for integrated models of biomolecular interaction networks. Genome Res 2003, 13:2498–2504.

78. Morris JH, Kuchinsky A, Ferrin TE, Pico AR: enhancedGraphics: a Cytoscape app for enhanced node graphics. F1000Res 2014, 3:147.

