## Additional Files for "Targeted phosphoproteomics of the Ras signaling network reveal regulatory mechanisms mediated by oncogenic KRAS": Additional_File_3.pdf

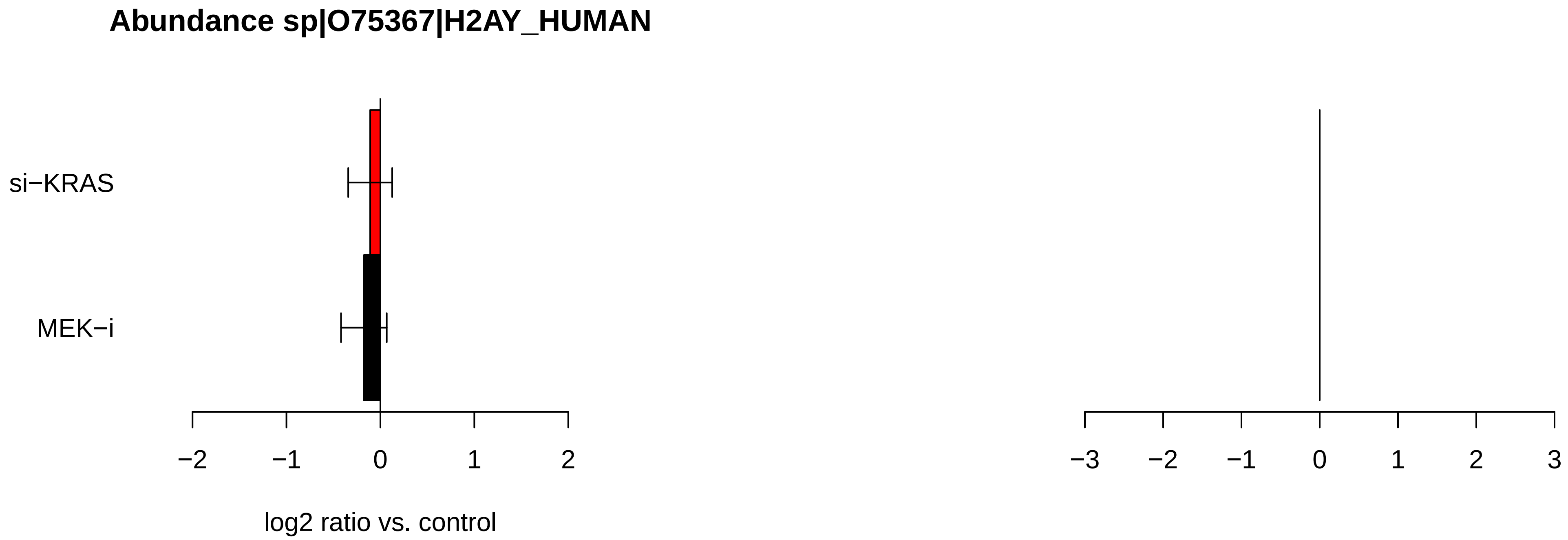
Abundance sp|O75367|H2AY_HUMAN
si−KRAS
MEK−i
−2
−1
0
1
2
log2 ratio vs. control
−3
−2
−1
0
1
2
3

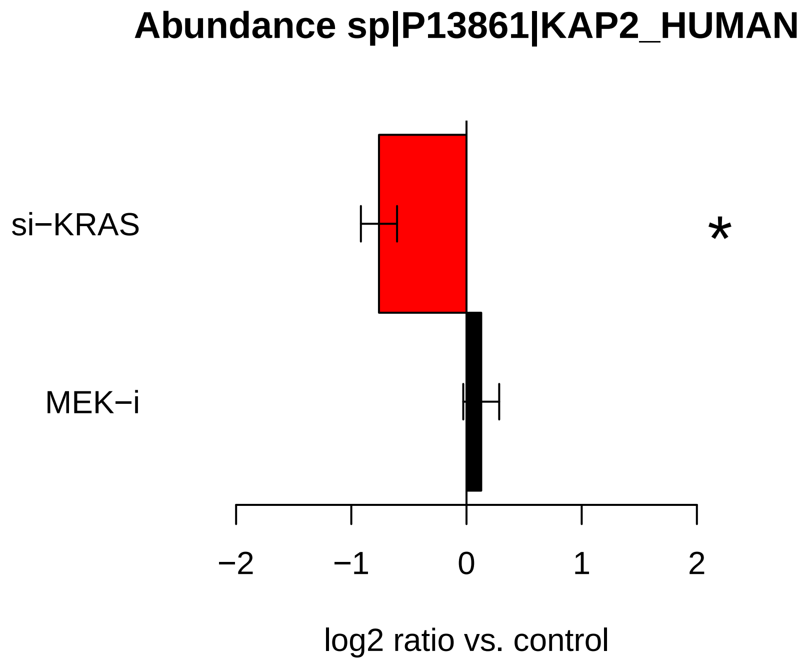
Abundance sp|P13861|KAP2_HUMAN
si−KRAS
MEK−i
*
−2
−1
0
1
2
log2 ratio vs. control

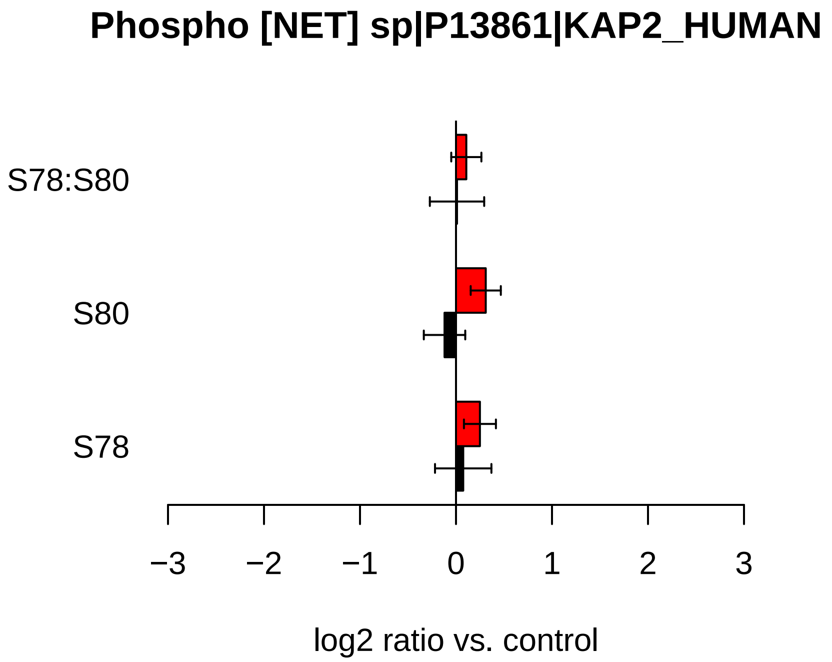
Phospho [NET] sp|P13861|KAP2_HUMAN
S78:S80
S80
S78
−3
−2
−1
0
1
2
3
log2 ratio vs. control

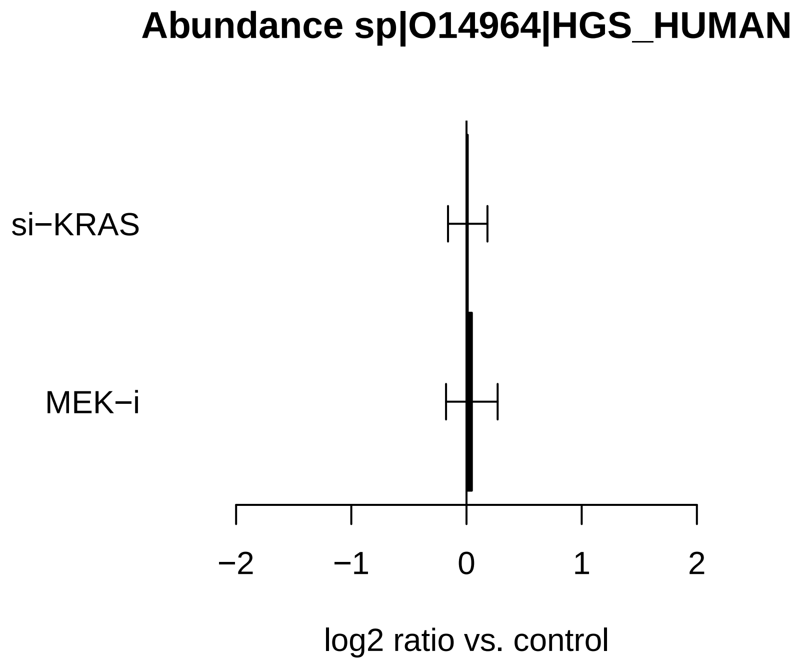
Abundance sp|O14964|HGS_HUMAN
si−KRAS
MEK−i
−2
−1
0
1
2
log2 ratio vs. control

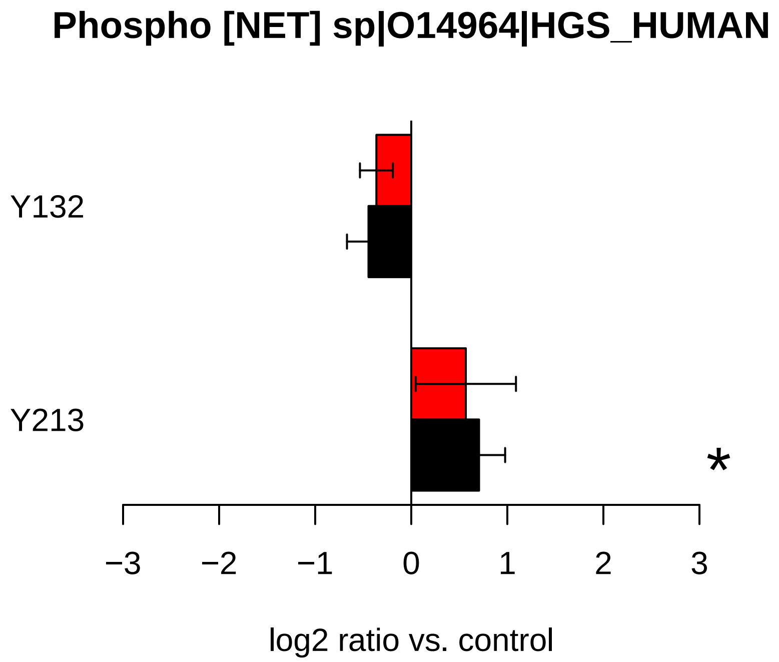
Phospho [NET] sp|O14964|HGS_HUMAN
Y132
Y213
*
−3
−2
−1
0
1
2
3
log2 ratio vs. control

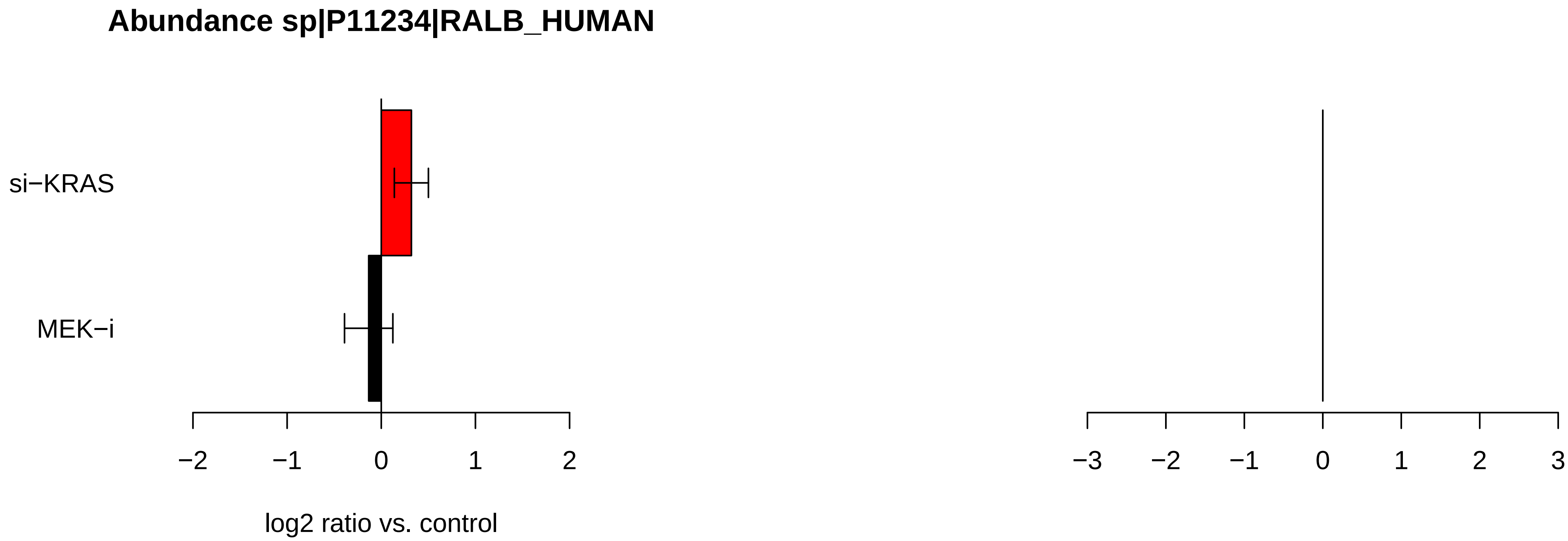

Abundance sp|P11234|RALB_HUMAN
si−KRAS
MEK−i
−2
−1
0
1
2
log2 ratio vs. control
−3
−2
−1
0
1
2
3

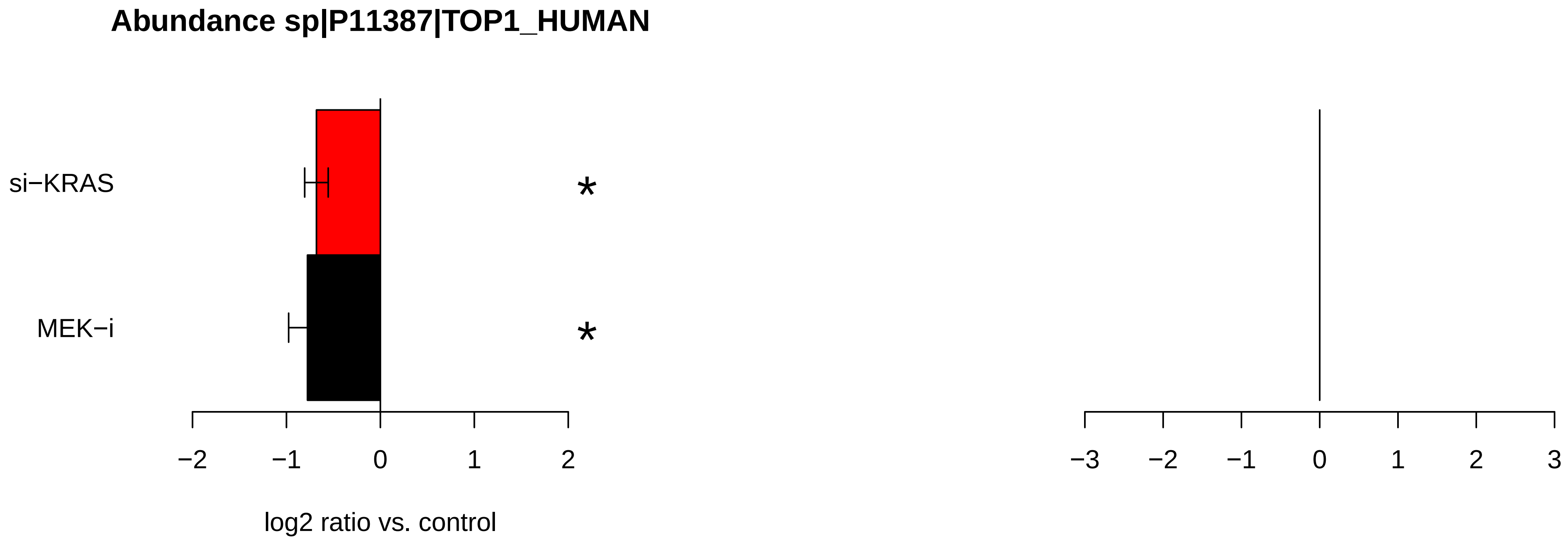
Abundance sp|P11387|TOP1_HUMAN
si−KRAS
MEK−i
*
*
−2
−1
0
1
2
log2 ratio vs. control
−3
−2
−1
0
1
2
3

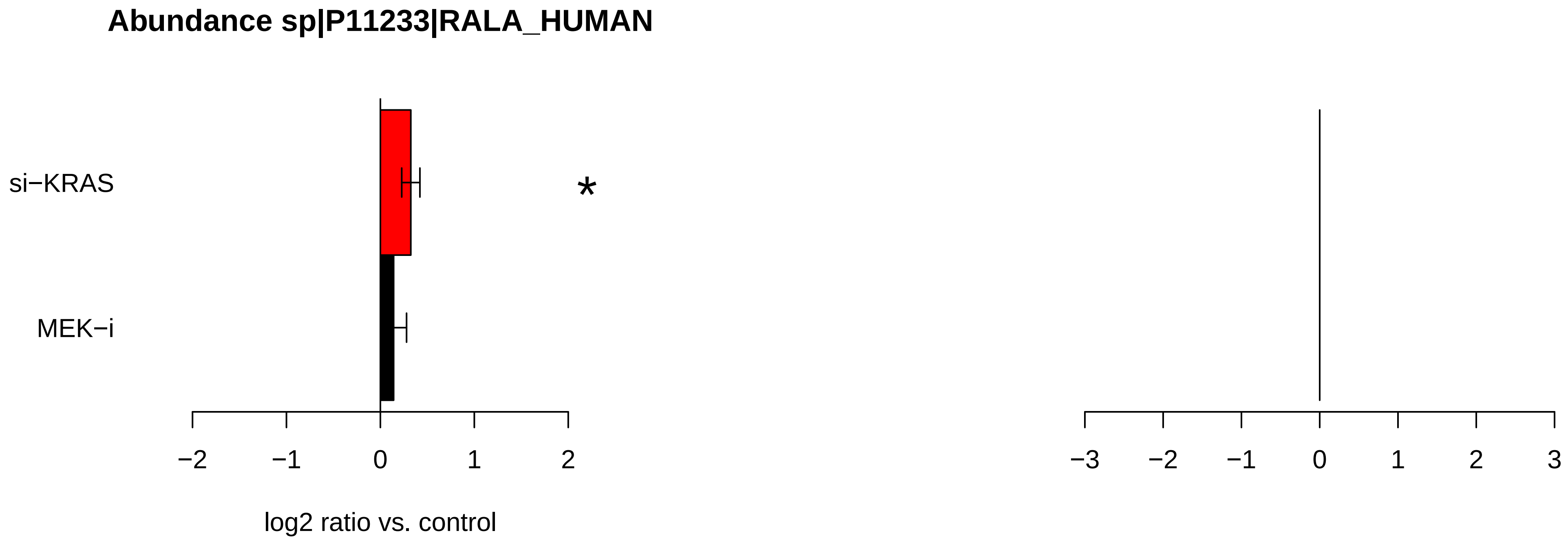

Abundance sp|P11233|RALA_HUMAN
si−KRAS
MEK−i
*
−2
−1
0
1
2
log2 ratio vs. control
−3
−2
−1
0
1
2
3

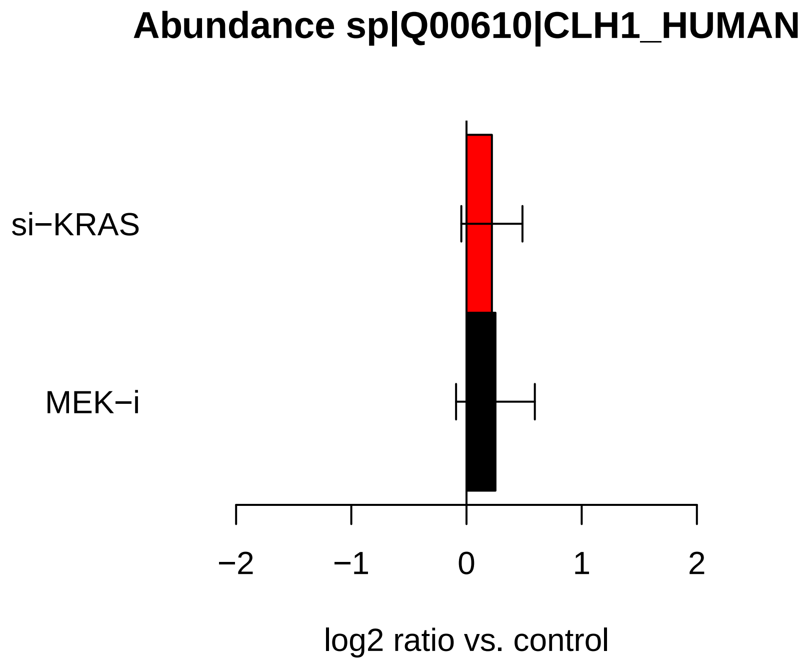

Abundance sp|Q00610|CLH1_HUMAN
si−KRAS
MEK−i
−2
−1
0
1
2
log2 ratio vs. control


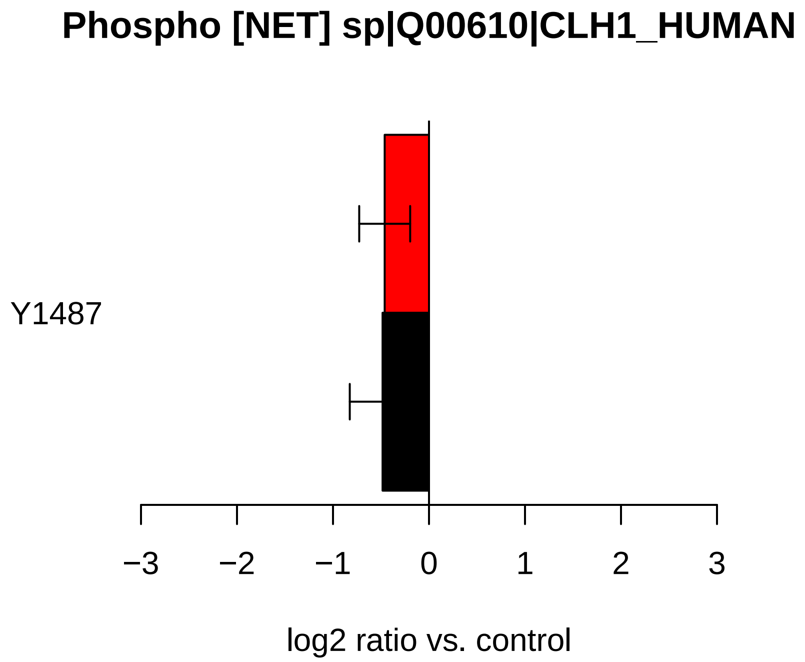

Phospho [NET] sp|Q00610|CLH1_HUMAN
Y1487
−3
−2
−1
0
1
2
3
log2 ratio vs. control

**Abundance sp|Q15118|PDK1_HUMAN**

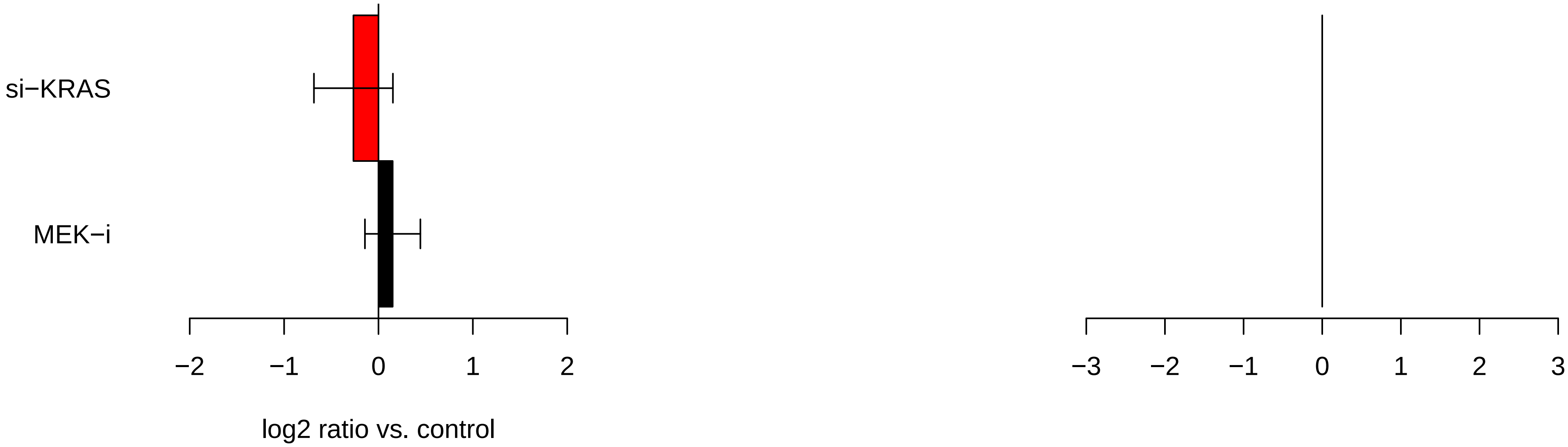

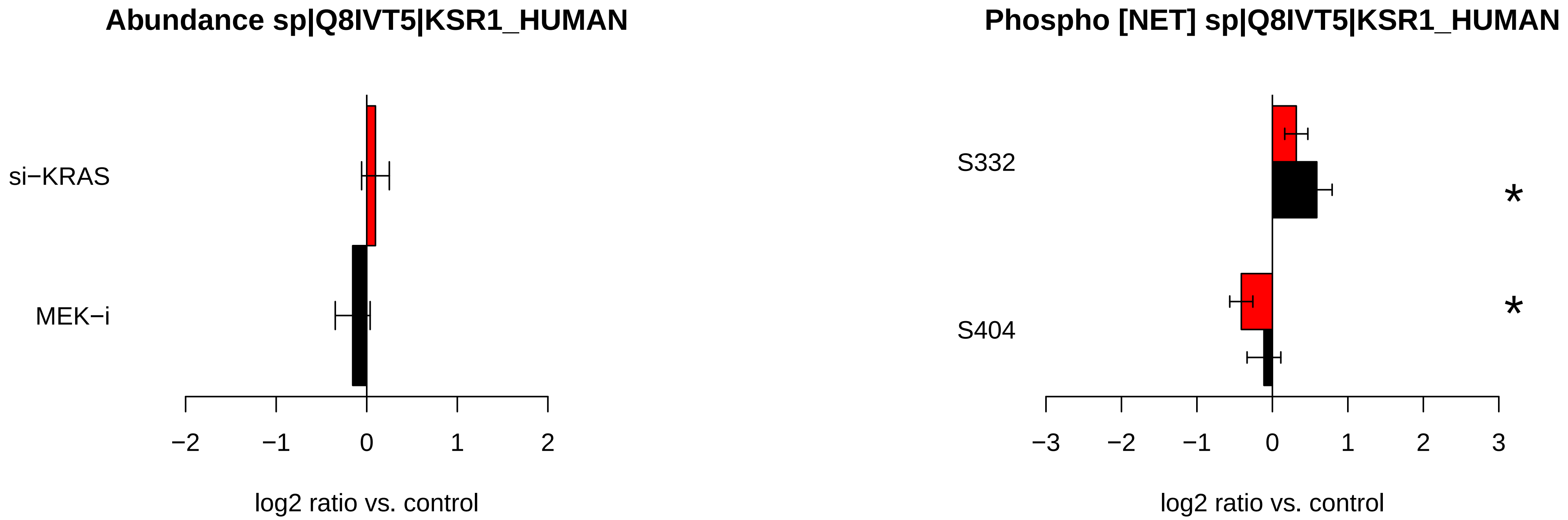

Abundance sp|Q8IVT5|KSR1_HUMAN
si−KRAS
MEK−i
−2
−1
0
1
2
log2 ratio vs. control
Phospho [NET] sp|Q8IVT5|KSR1_HUMAN
S332
S404
*
*
−3
−2
−1
0
1
2
3
log2 ratio vs. control

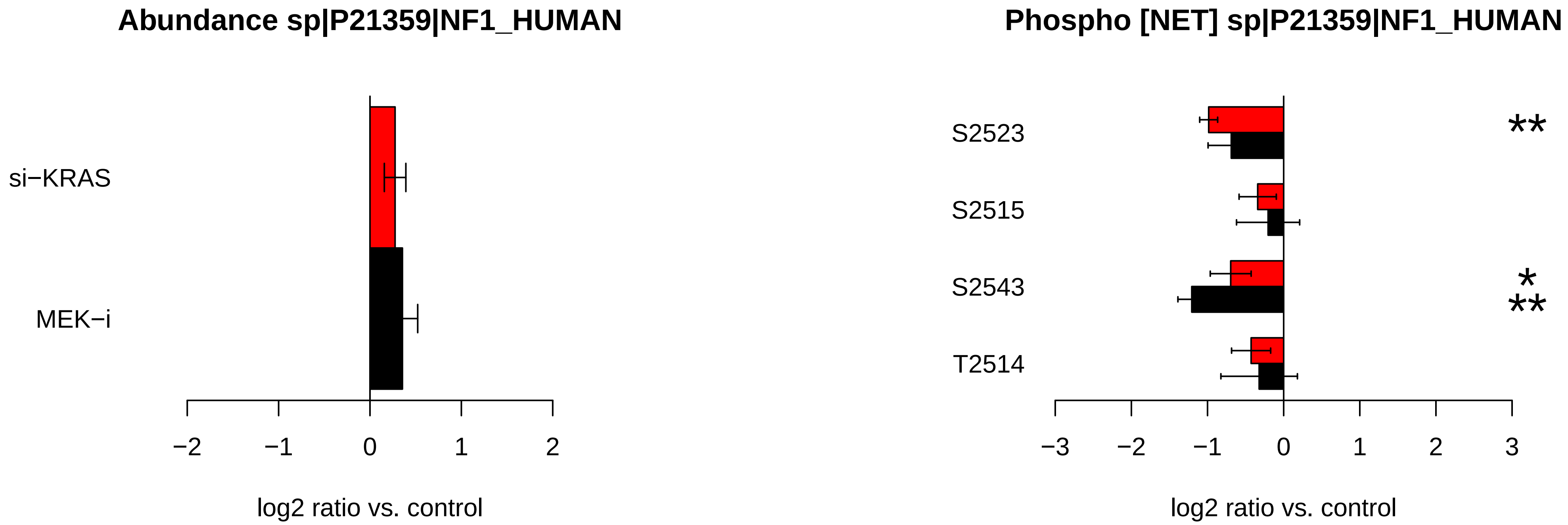

Abundance sp|P21359|NF1_HUMAN
si−KRAS
MEK−i
−2
−1
0
1
2
log2 ratio vs. control
Phospho [NET] sp|P21359|NF1_HUMAN
S2523
S2515
S2543
T2514
**
*
**
−3
−2
−1
0
1
2
3
log2 ratio vs. control

### Abundance sp|Q15149|PLEC_HUMAN

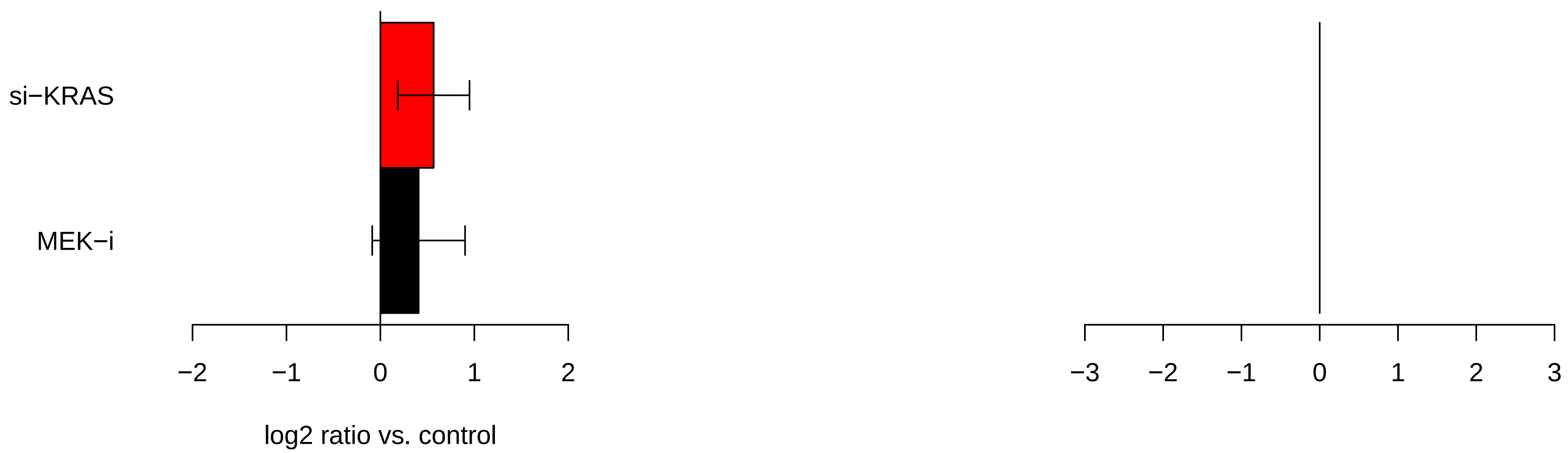

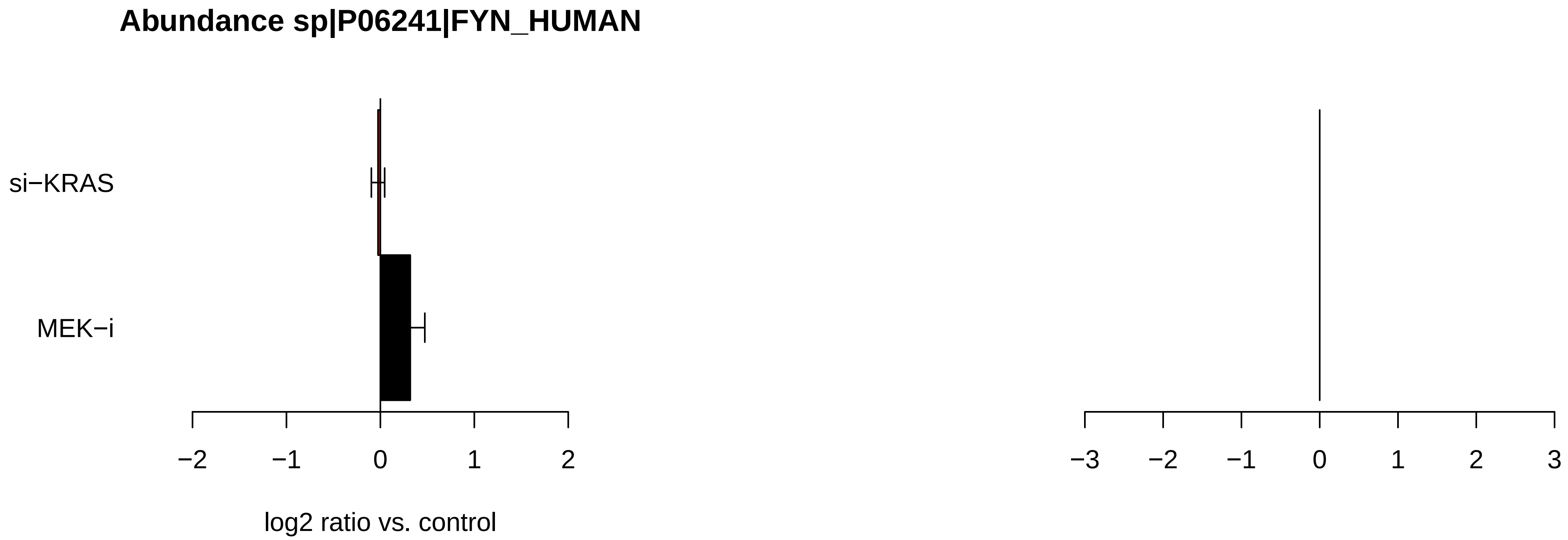
Abundance sp|P06241|FYN_HUMAN
si−KRAS
MEK−i
−2
−1
0
1
2
log2 ratio vs. control
−3
−2
−1
0
1
2
3

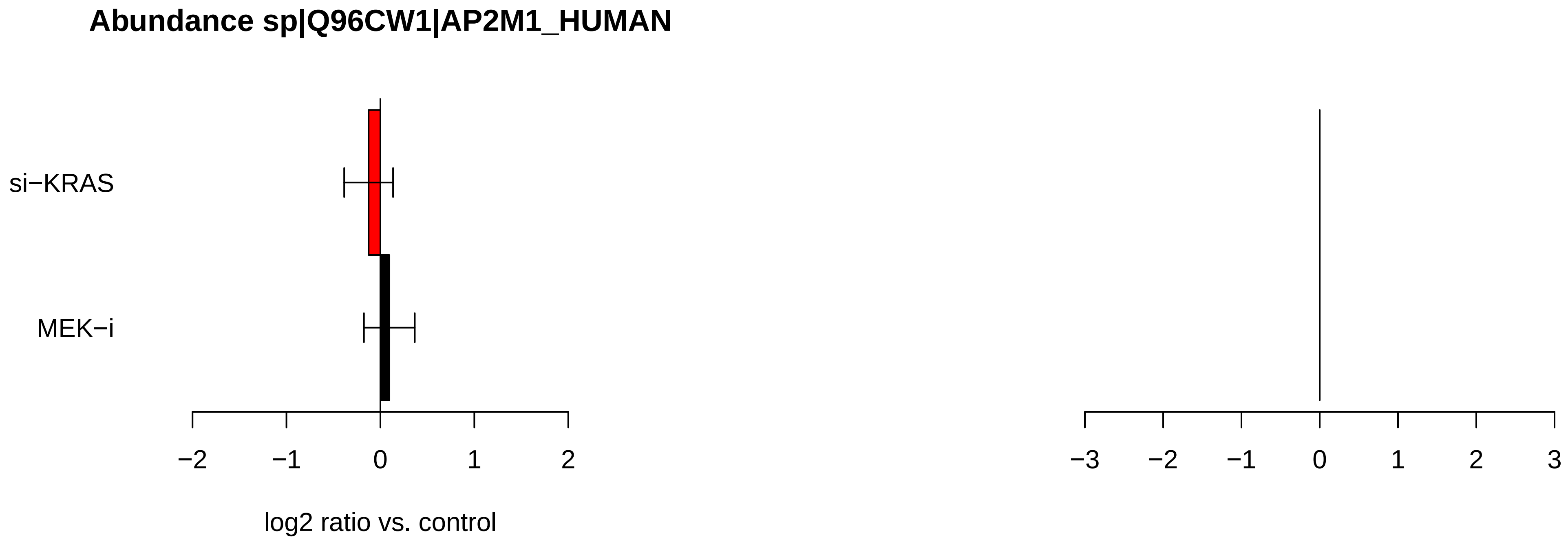

Abundance sp|Q96CW1|AP2M1_HUMAN
si−KRAS
MEK−i
−2
−1
0
1
2
log2 ratio vs. control
−3
−2
−1
0
1
2
3

### Abundance sp|P52789|HXK2_HUMAN

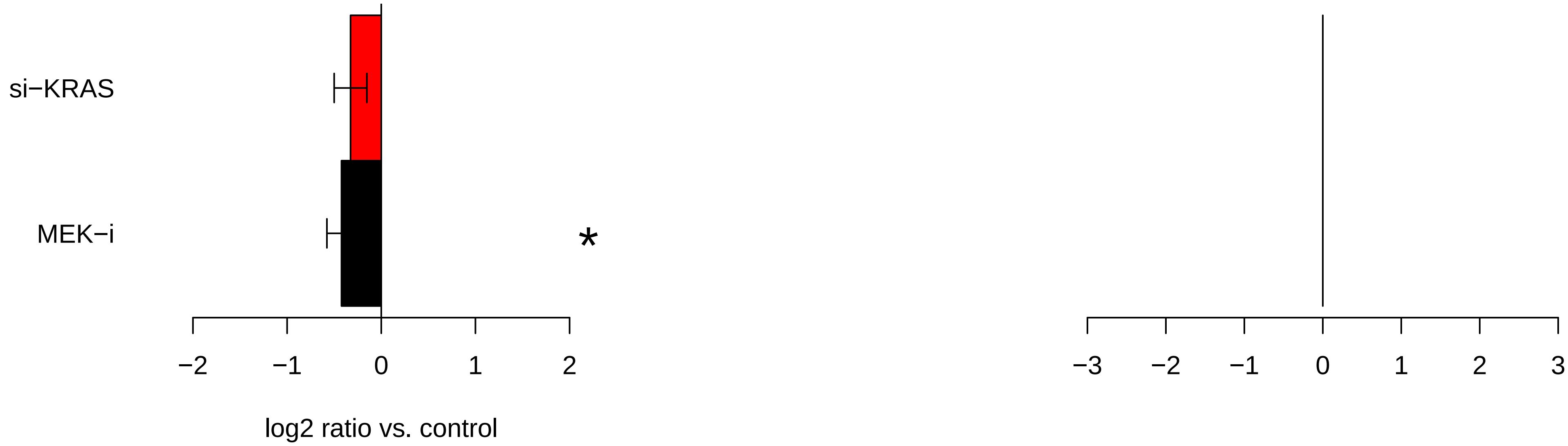

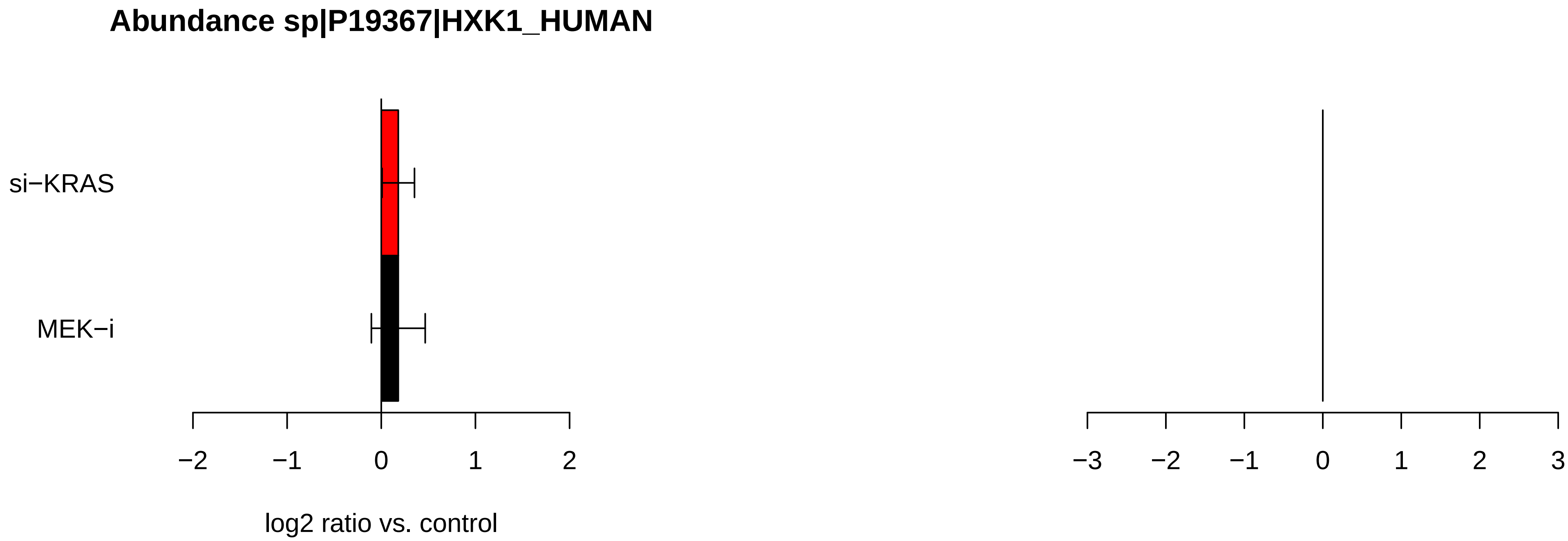
Abundance sp|P19367|HXK1_HUMAN
si−KRAS
MEK−i
−2
−1
0
1
2
log2 ratio vs. control
−3
−2
−1
0
1
2
3

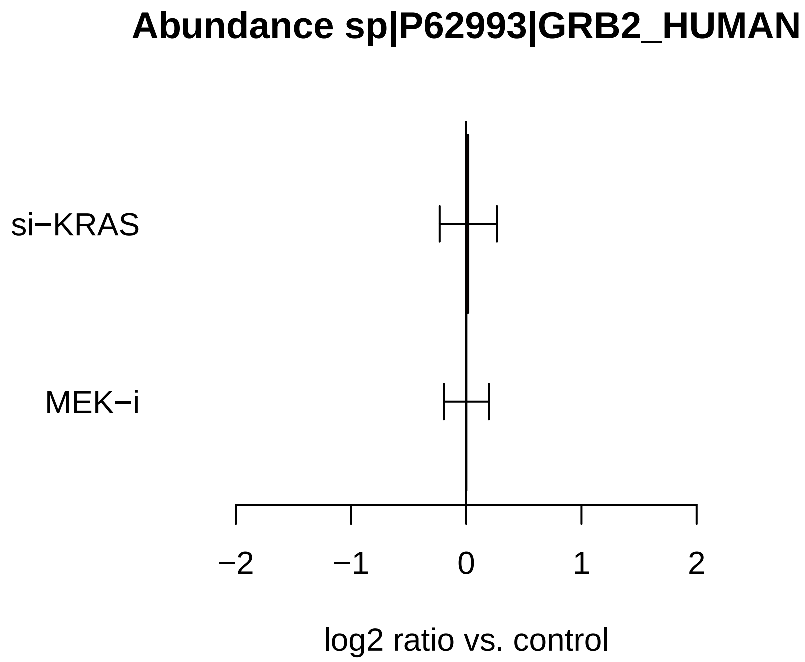
Abundance sp|P62993|GRB2_HUMAN
si−KRAS
MEK−i
−2
−1
0
1
2
log2 ratio vs. control

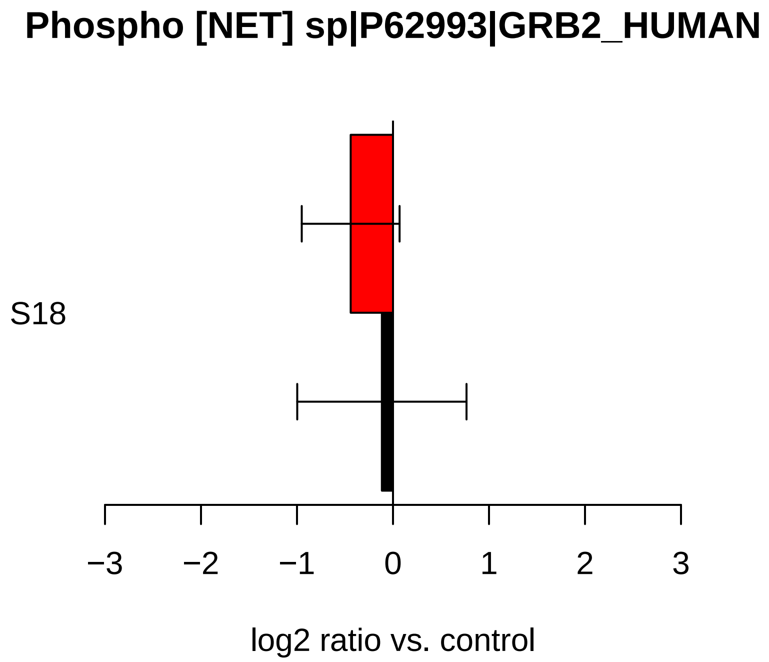
Phospho [NET] sp|P62993|GRB2_HUMAN
S18
−3
−2
−1
0
1
2
3
log2 ratio vs. control

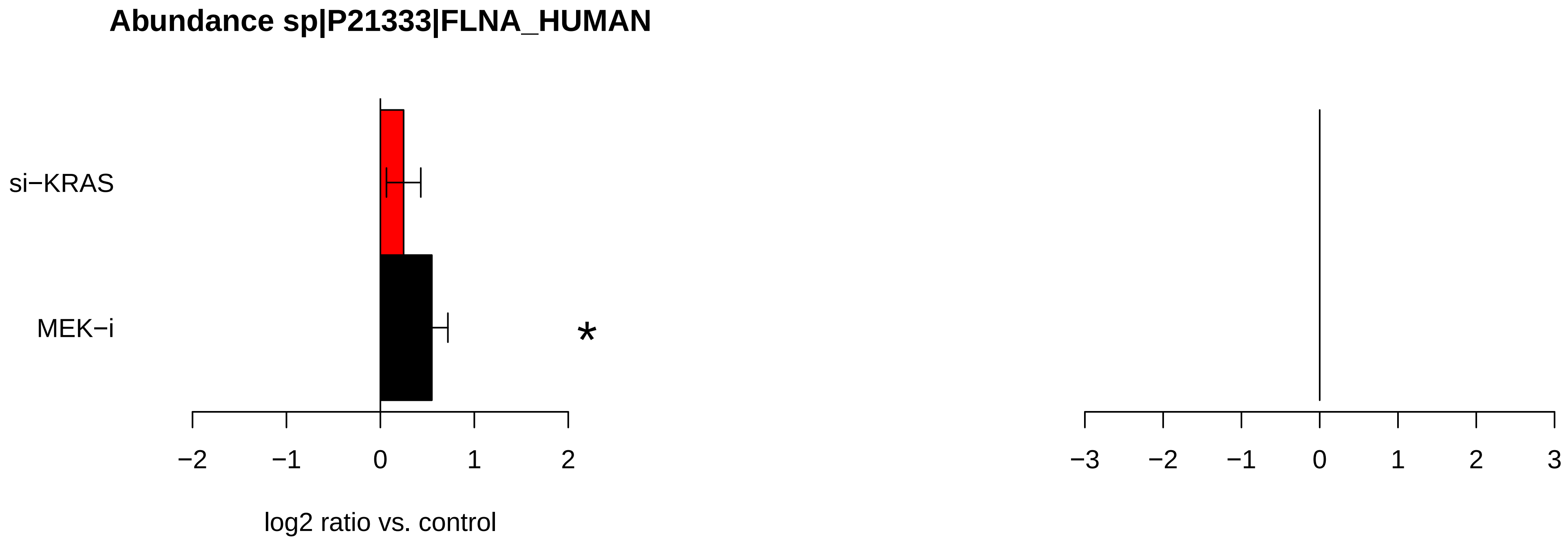
Abundance sp|P21333|FLNA_HUMAN
si−KRAS
MEK−i
*
−2
−1
0
1
2
log2 ratio vs. control
−3
−2
−1
0
1
2
3

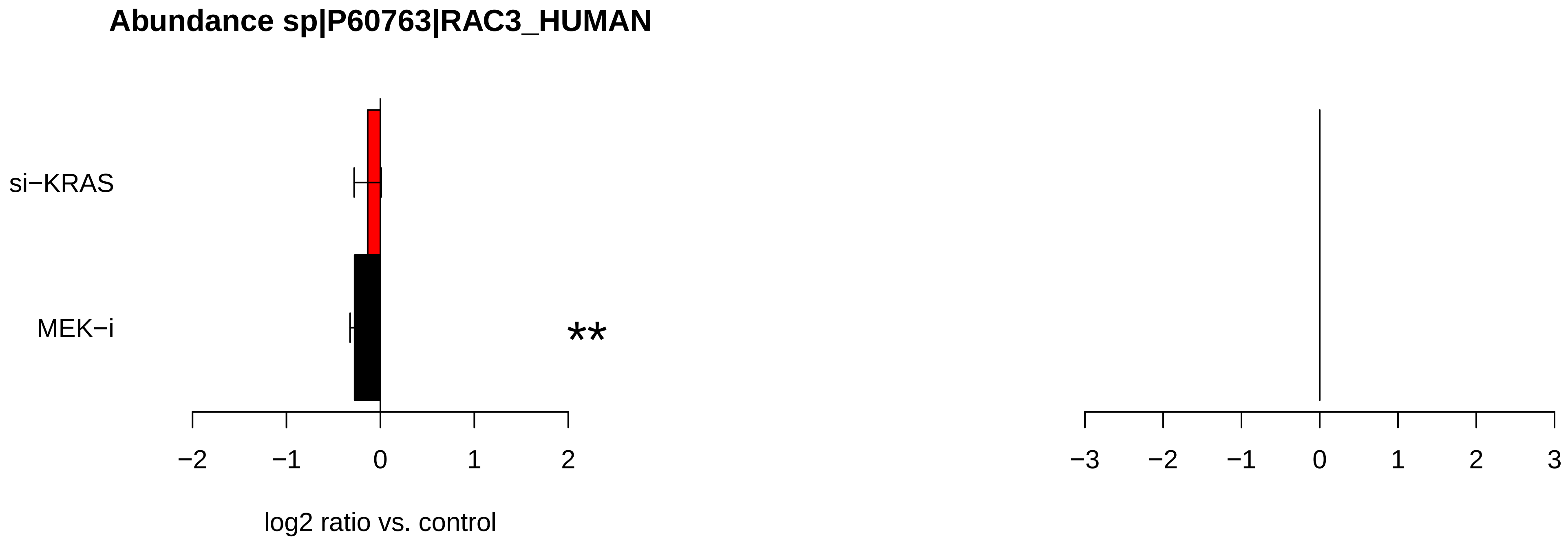

Abundance sp|P60763|RAC3_HUMAN
si−KRAS
MEK−i
**
−2
−1
0
1
2
log2 ratio vs. control
−3
−2
−1
0
1
2
3

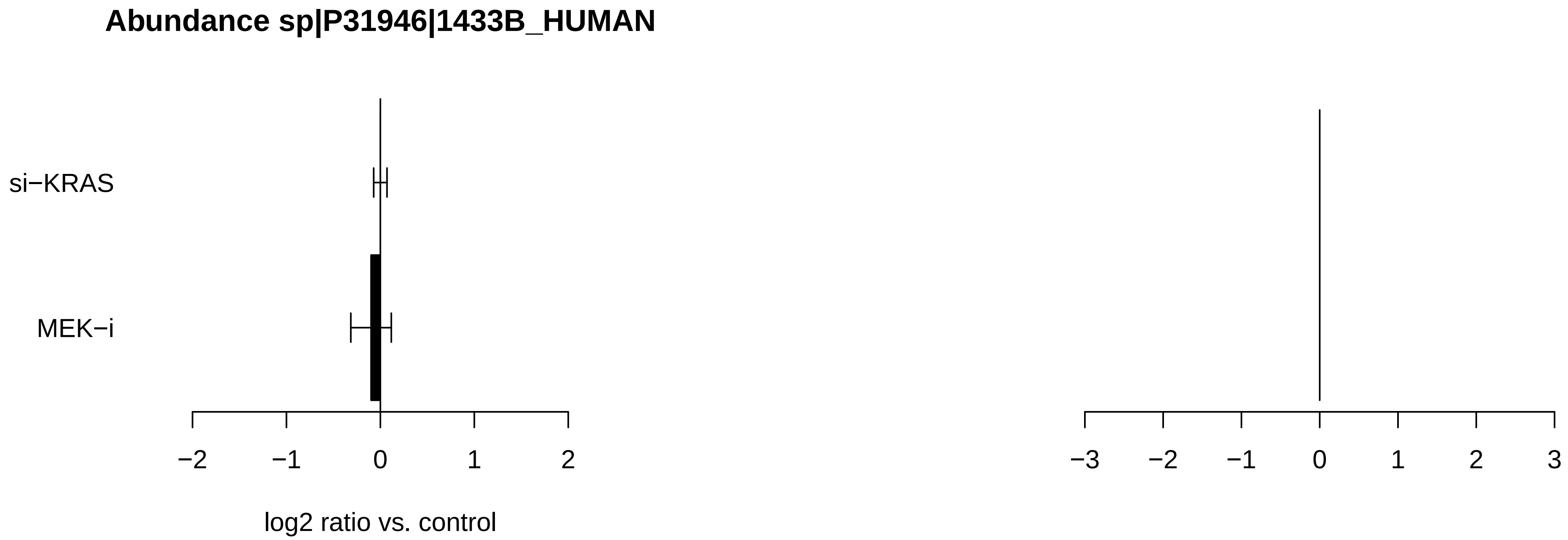
Abundance sp|P31946|1433B_HUMAN
si−KRAS
MEK−i
−2
−1
0
1
2
log2 ratio vs. control
−3
−2
−1
0
1
2
3

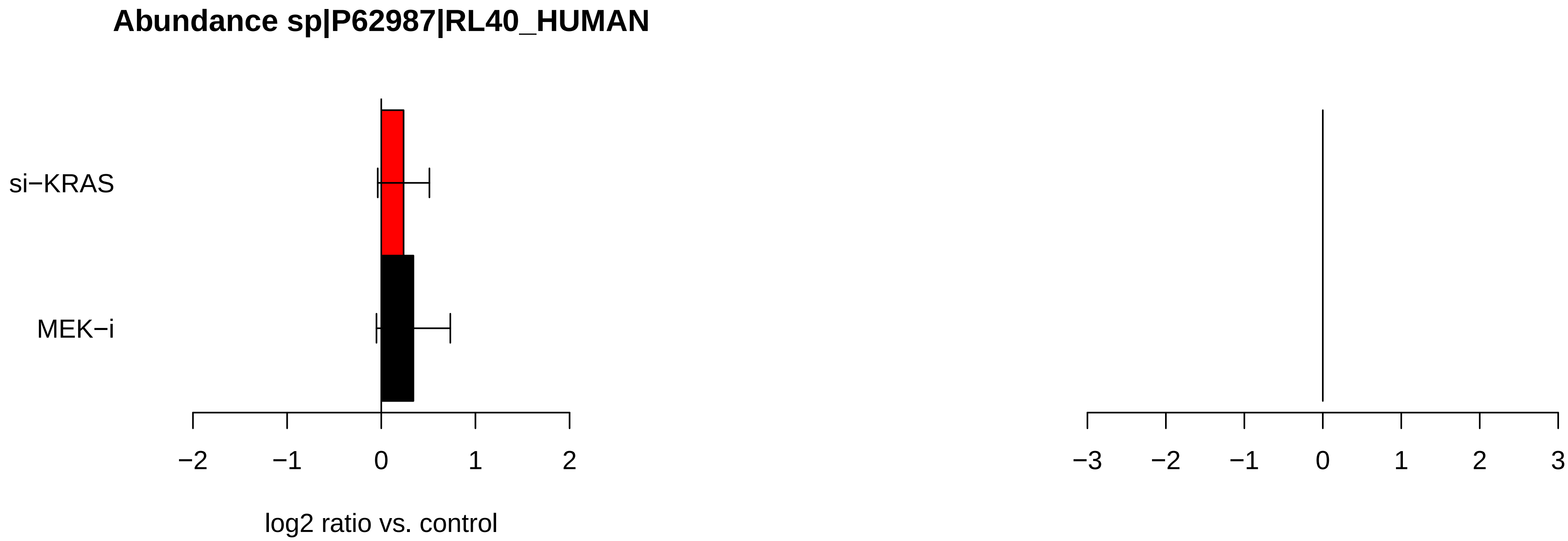

Abundance sp|P62987|RL40_HUMAN
si−KRAS
MEK−i
−2
−1
0
1
2
log2 ratio vs. control
−3
−2
−1
0
1
2
3

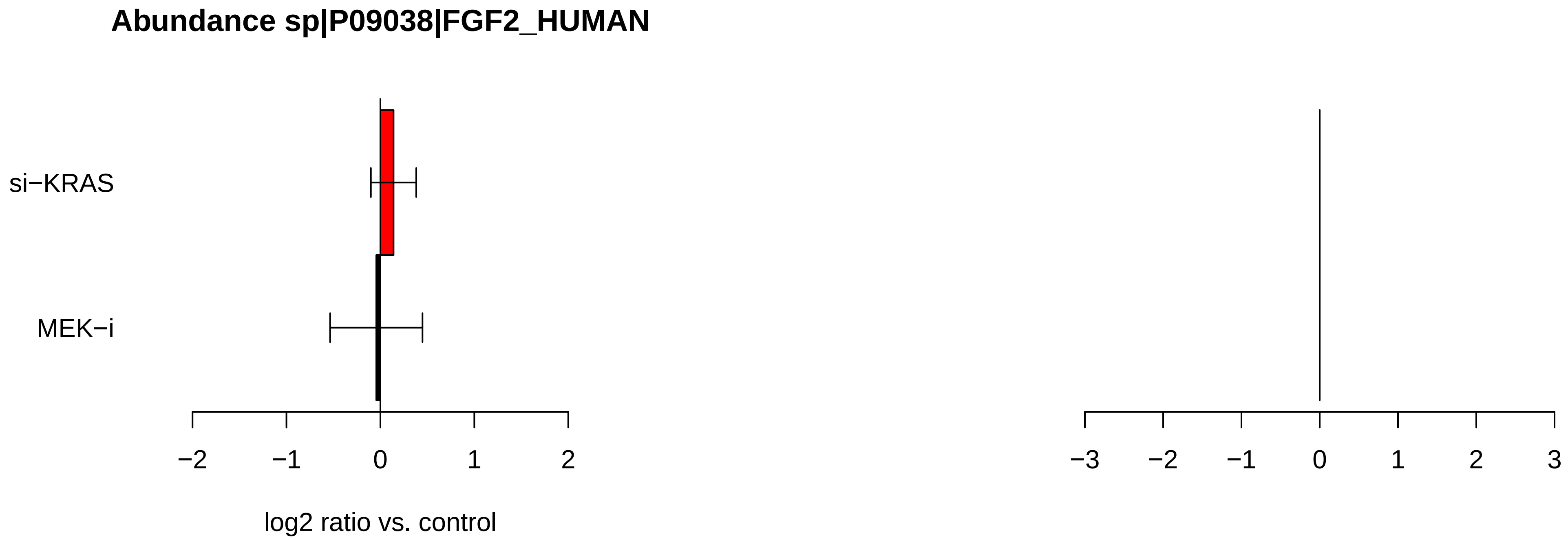
Abundance sp|P09038|FGF2_HUMAN
si−KRAS
MEK−i
−2
−1
0
1
2
log2 ratio vs. control
−3
−2
−1
0
1
2
3

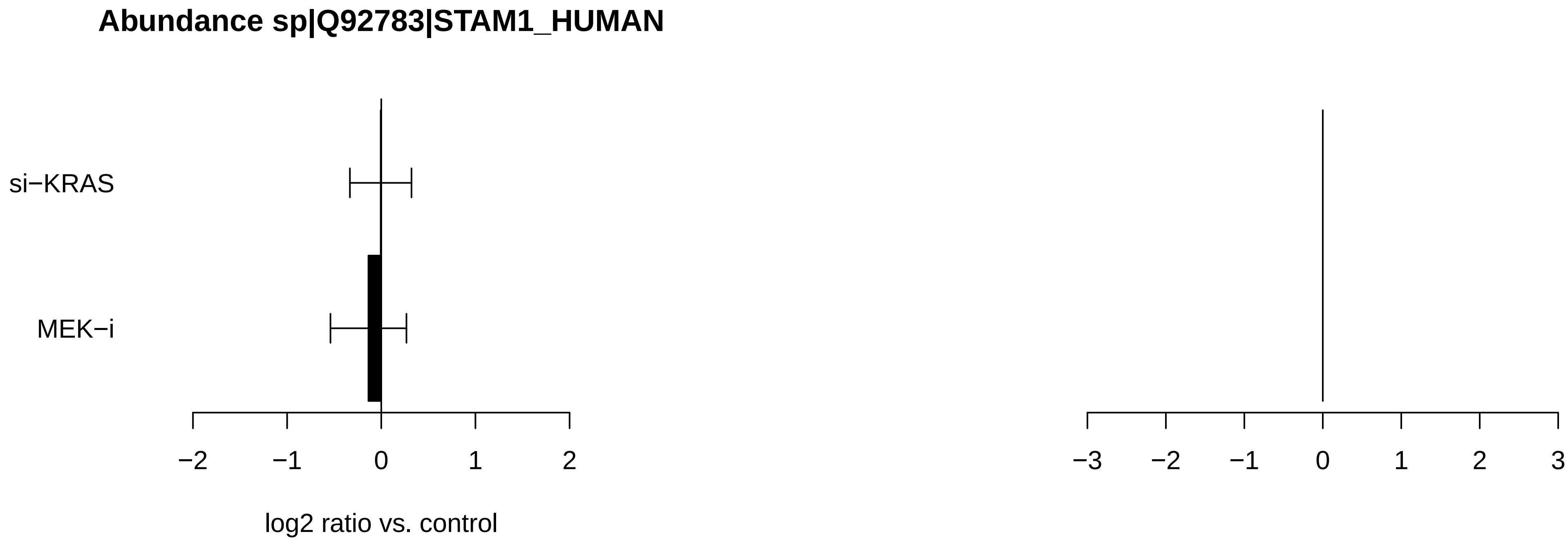

Abundance sp|Q92783|STAM1_HUMAN
si−KRAS
MEK−i
−2
−1
0
1
2
log2 ratio vs. control
−3
−2
−1
0
1
2
3

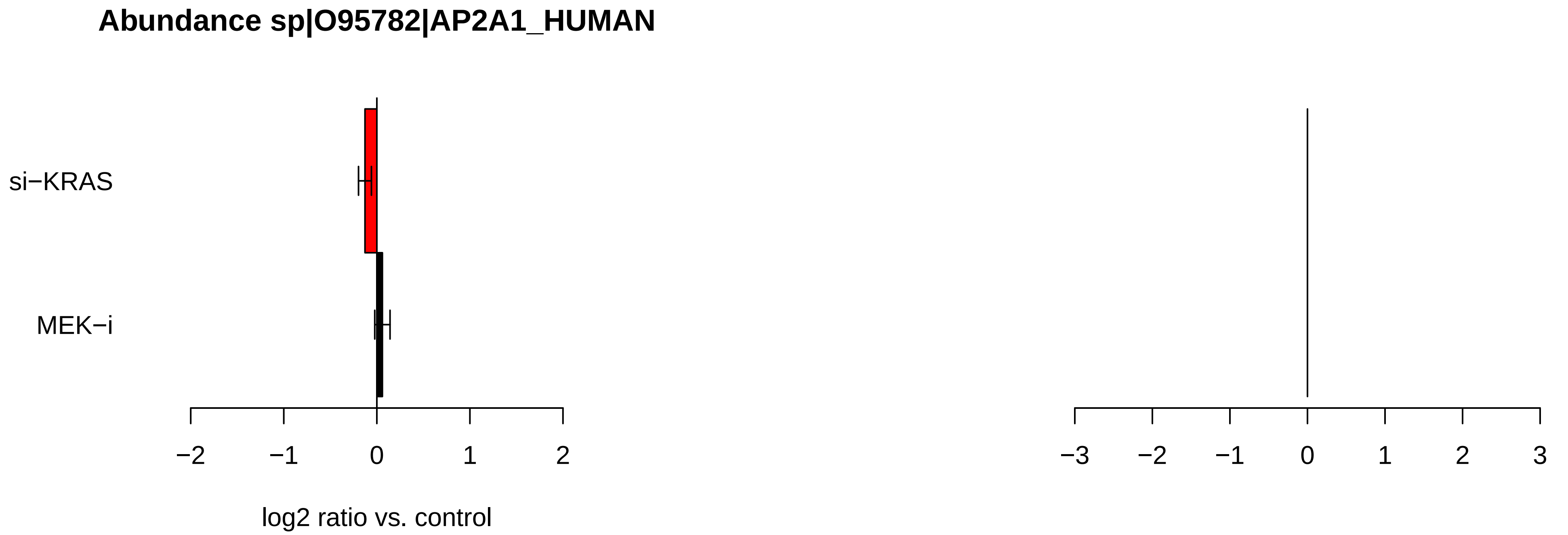
Abundance sp|O95782|AP2A1_HUMAN
si−KRAS
MEK−i
−2
−1
0
1
2
log2 ratio vs. control
−3
−2
−1
0
1
2
3

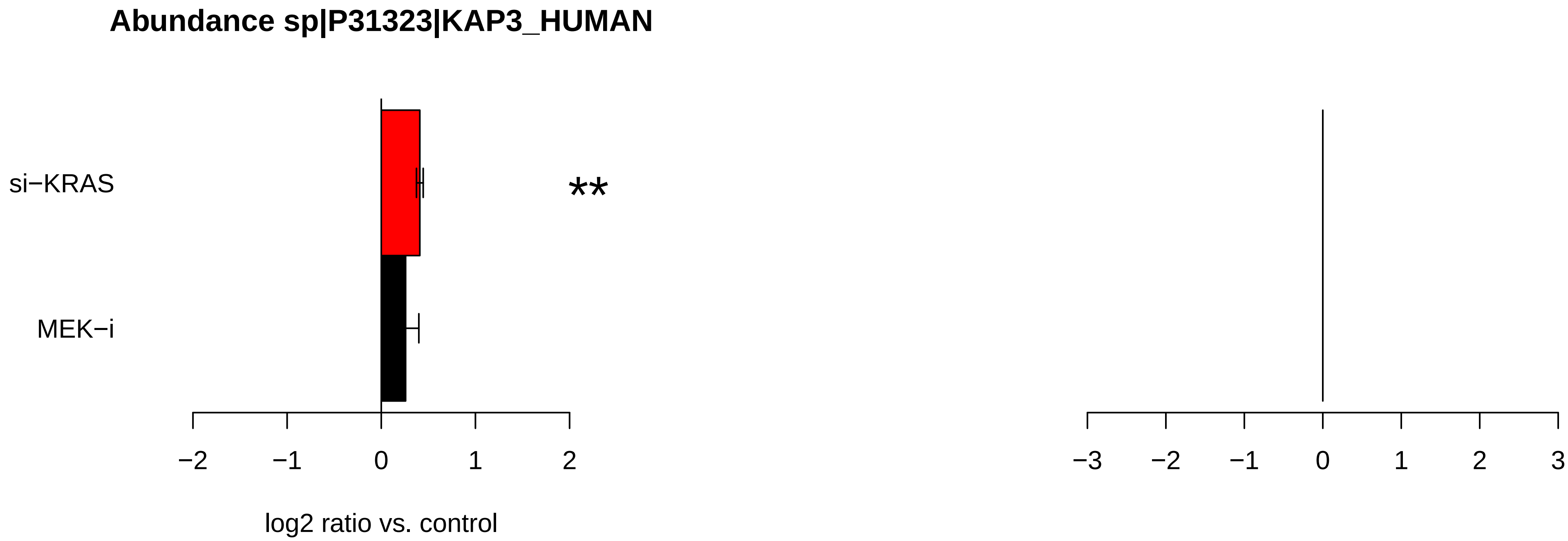
Abundance sp|P31323|KAP3_HUMAN
si−KRAS
MEK−i
**
−2
−1
0
1
2
log2 ratio vs. control
−3
−2
−1
0
1
2
3

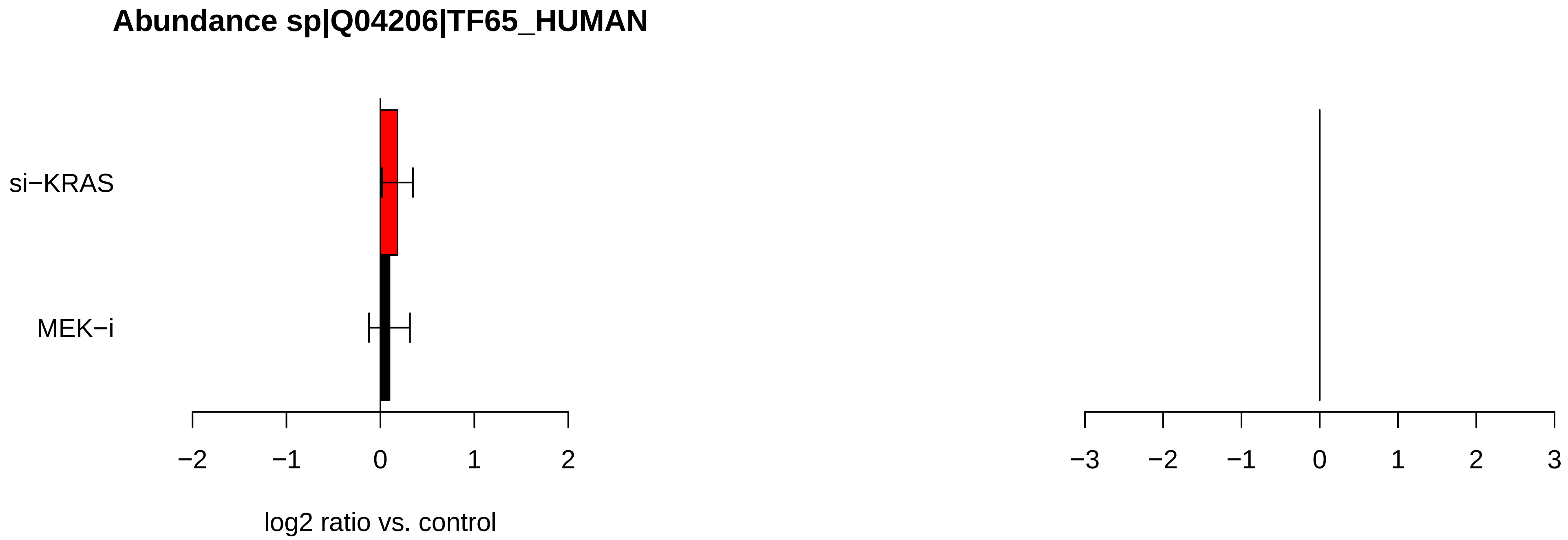
Abundance sp|Q04206|TF65_HUMAN
si−KRAS
MEK−i
−2
−1
0
1
2
log2 ratio vs. control
−3
−2
−1
0
1
2
3

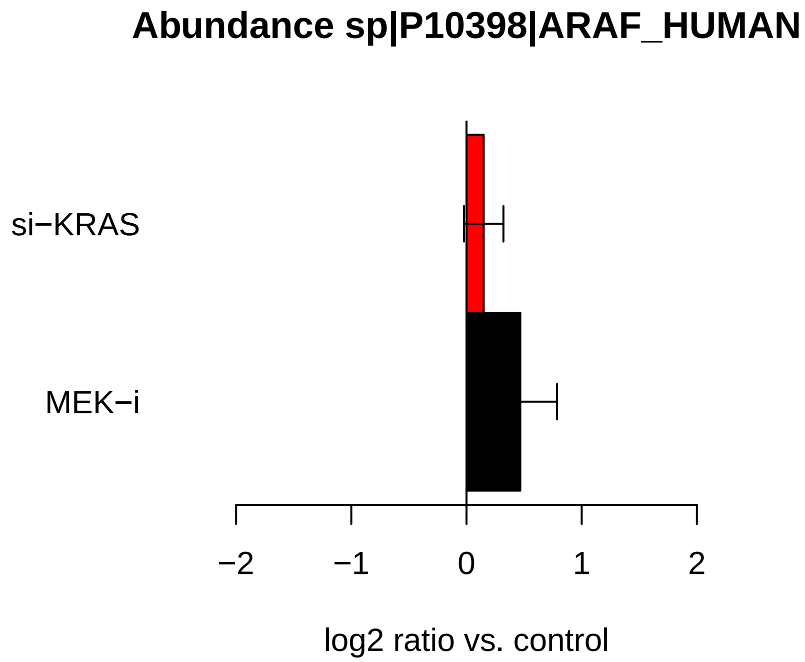
Abundance sp|P10398|ARAF_HUMAN
si−KRAS
MEK−i
−2
−1
0
1
2
log2 ratio vs. control

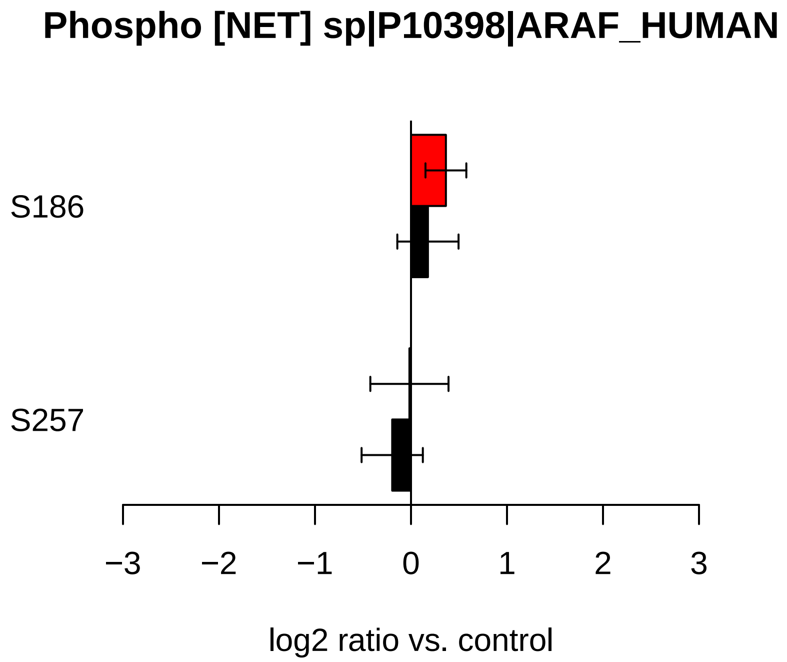
Phospho [NET] sp|P10398|ARAF_HUMAN
S186
S257
−3
−2
−1
0
1
2
3
log2 ratio vs. control

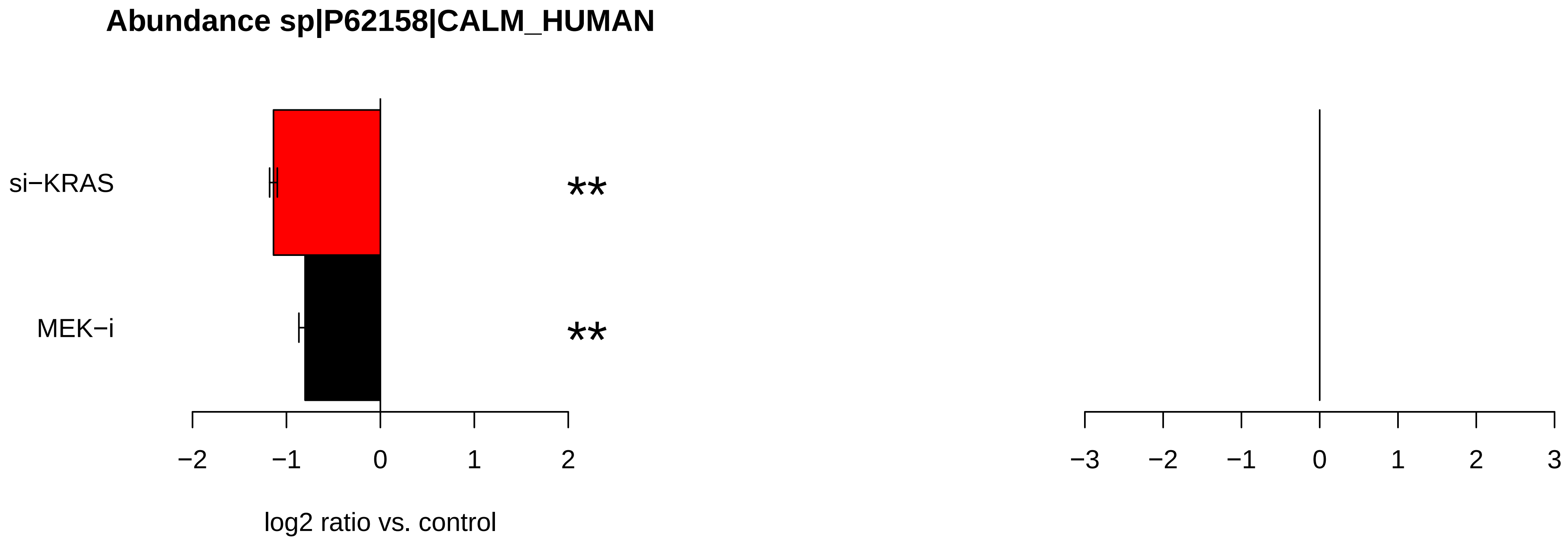
Abundance sp|P62158|CALM_HUMAN
si−KRAS
MEK−i
**
**
−2
−1
0
1
2
−3
−2
−1
0
1
2
3
log2 ratio vs. control

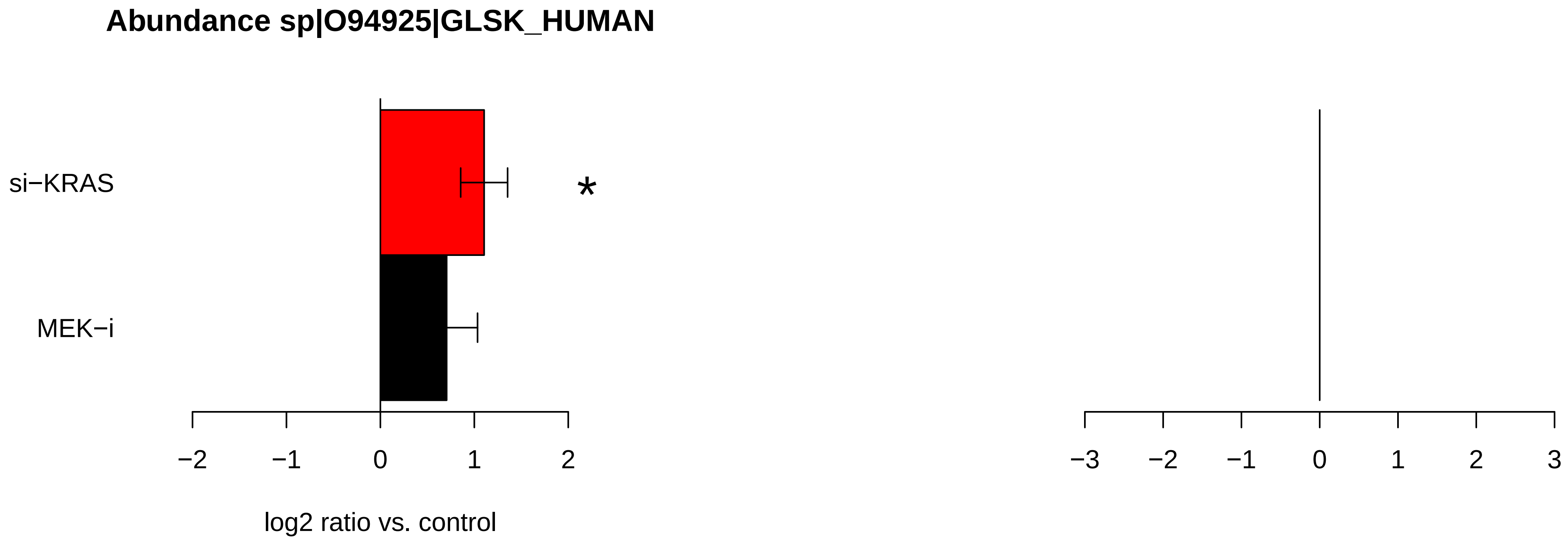
Abundance sp|O94925|GLSK_HUMAN
si−KRAS
MEK−i
*
−2
−1
0
1
2
log2 ratio vs. control
−3
−2
−1
0
1
2
3

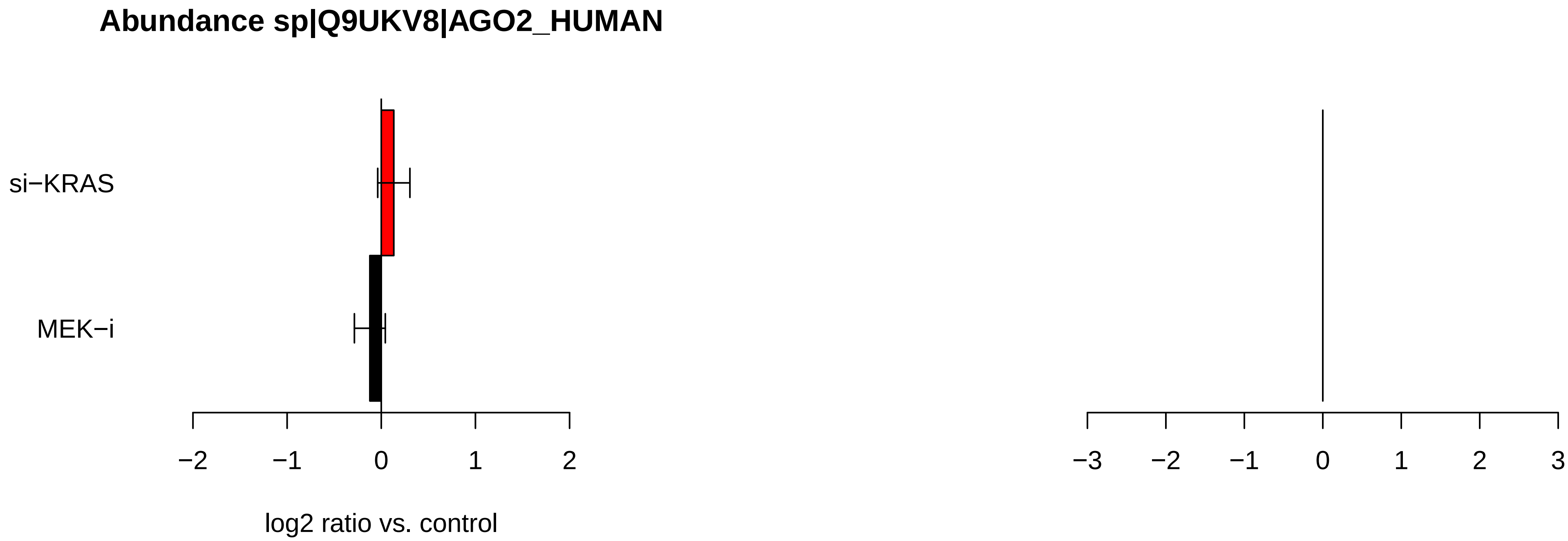
Abundance sp|Q9UKV8|AGO2_HUMAN
si−KRAS
MEK−i
−2
−1
0
1
2
log2 ratio vs. control
−3
−2
−1
0
1
2
3

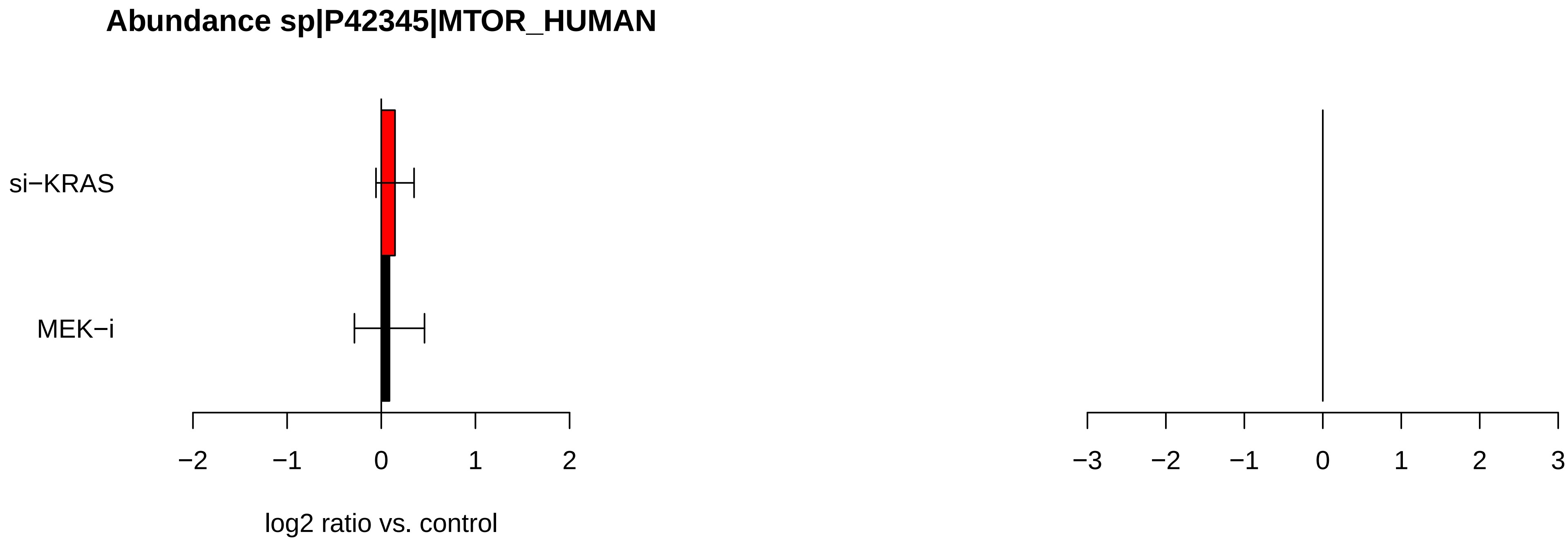
Abundance sp|P42345|MTOR_HUMAN
si−KRAS
MEK−i
−2
−1
0
1
2
log2 ratio vs. control
−3
−2
−1
0
1
2
3

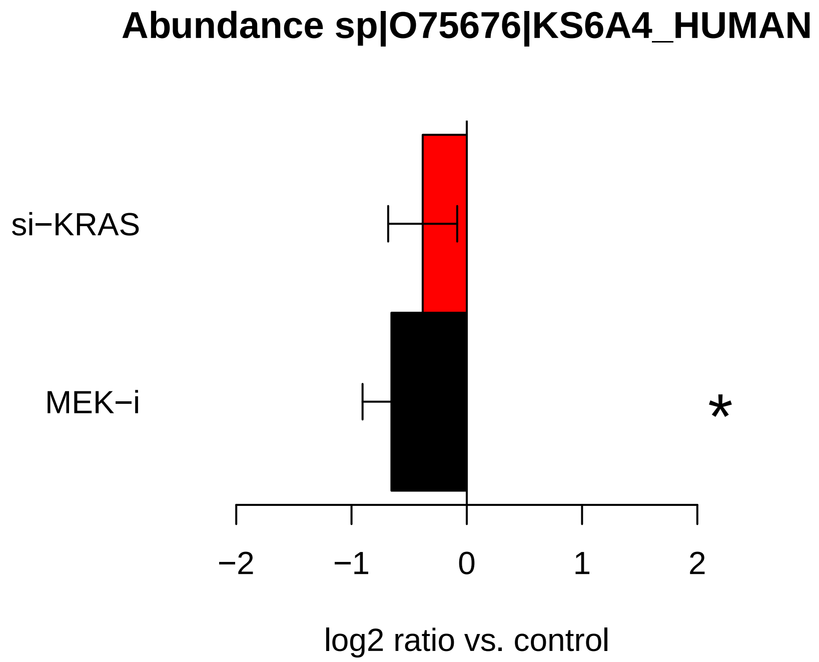
Abundance sp|O75676|KS6A4_HUMAN
si−KRAS
MEK−i
*
−2
−1
0
1
2
log2 ratio vs. control

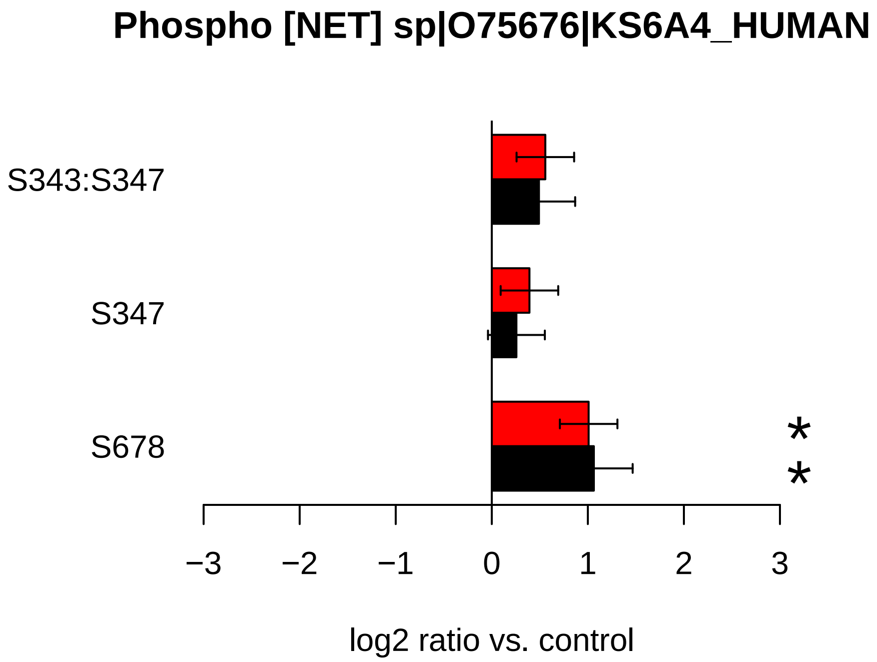
Phospho [NET] sp|O75676|KS6A4_HUMAN
S343:S347
S347
S678
*
*
−3
−2
−1
0
1
2
3
log2 ratio vs. control

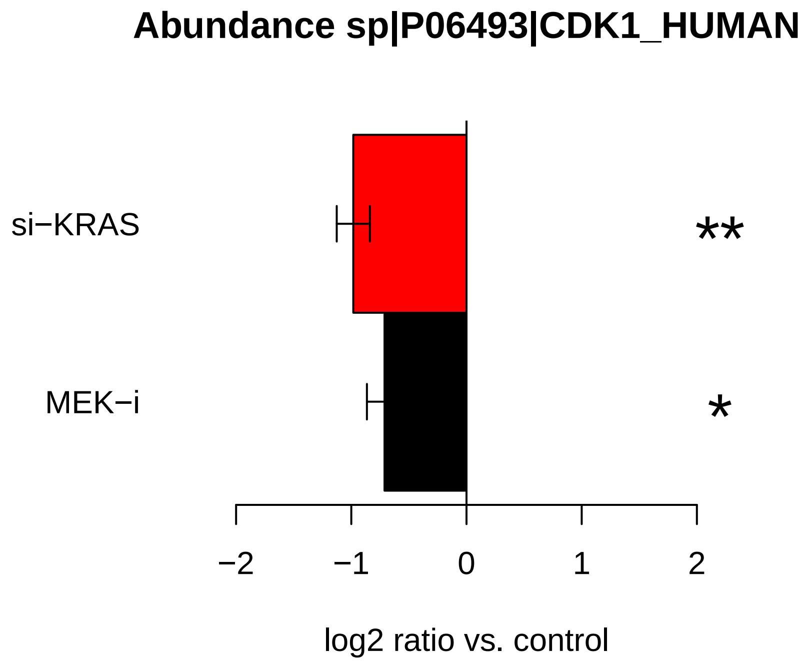

Abundance sp|P06493|CDK1_HUMAN
si−KRAS
MEK−i
**
*
−2
−1
0
1
2
log2 ratio vs. control


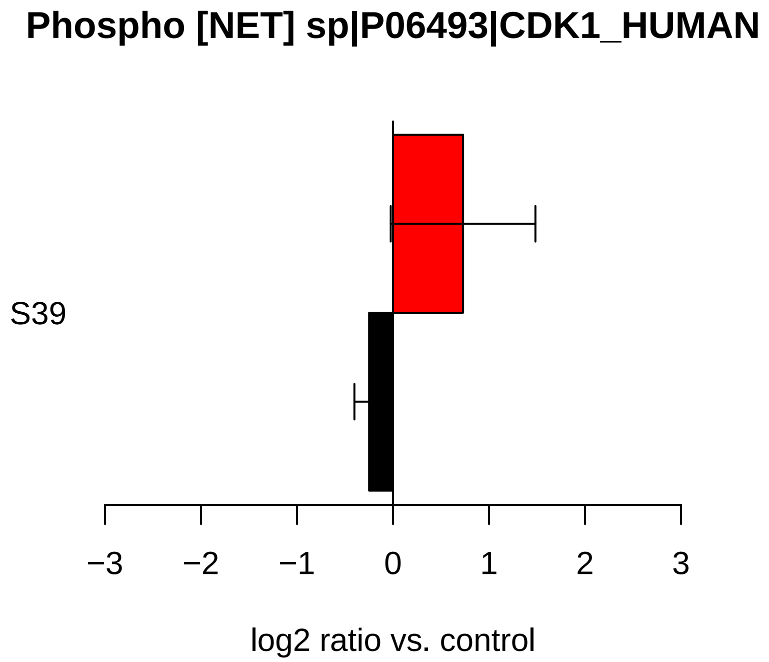

Phospho [NET] sp|P06493|CDK1_HUMAN
S39
−3
−2
−1
0
1
2
3
log2 ratio vs. control

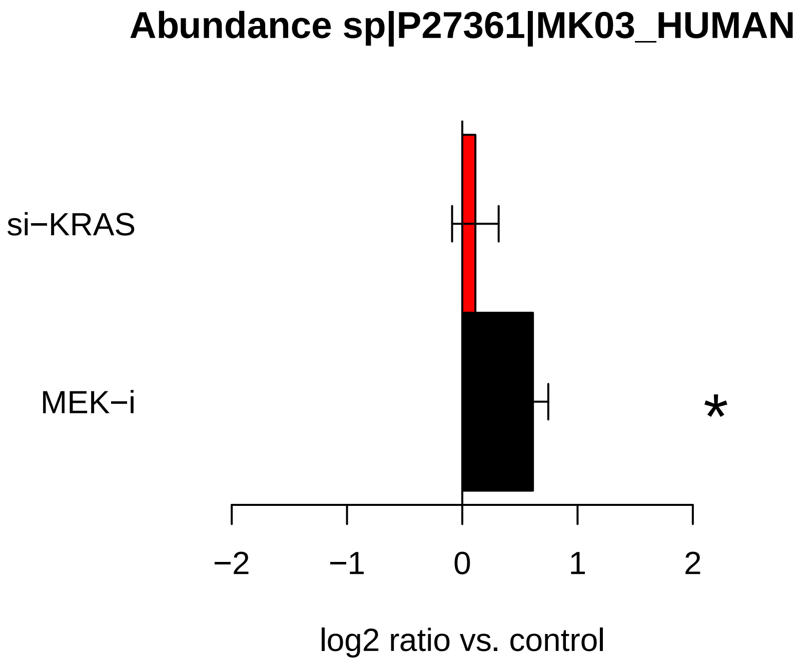
Abundance sp|P27361|MK03_HUMAN
si−KRAS
MEK−i
*
−2
−1
0
1
2
log2 ratio vs. control

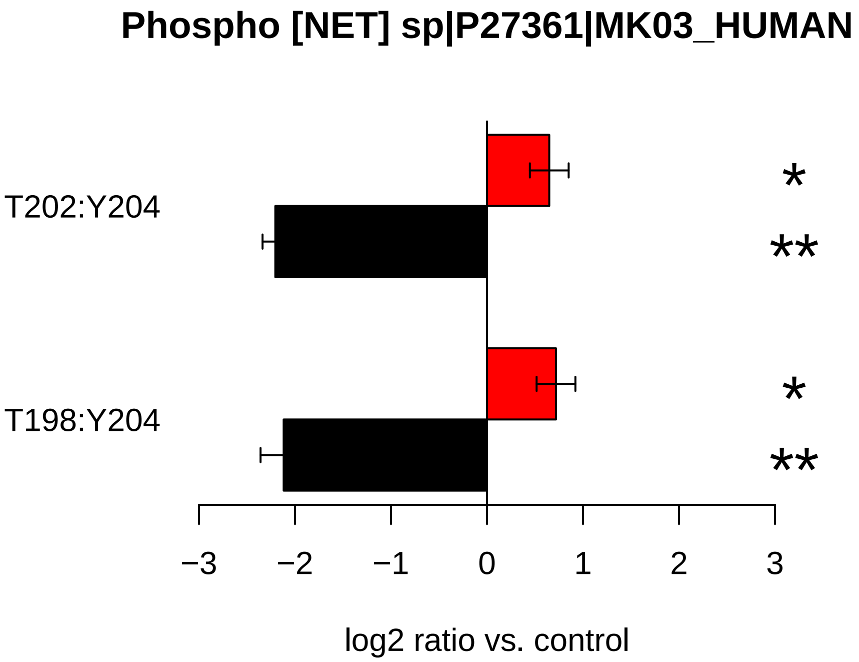
Phospho [NET] sp|P27361|MK03_HUMAN
T202:Y204
T198:Y204
*
**
*
**
−3
−2
−1
0
1
2
3
log2 ratio vs. control

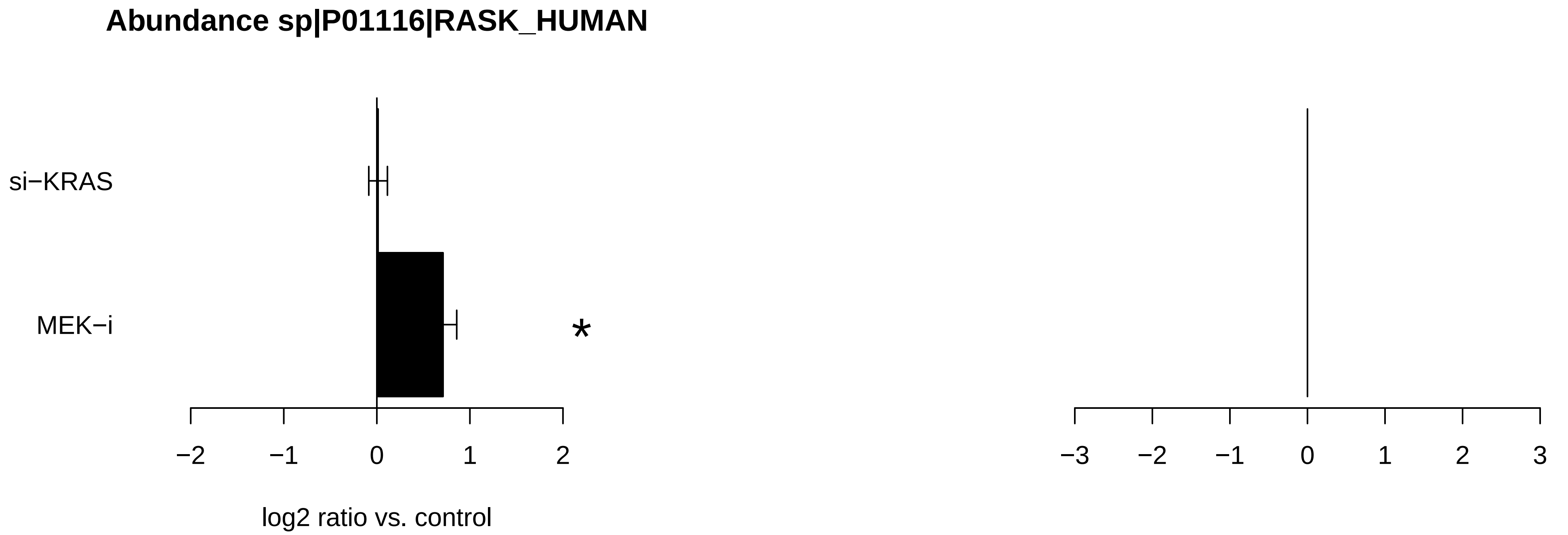

Abundance sp|P01116|RASK_HUMAN
si−KRAS
MEK−i
*
−2
−1
0
1
2
log2 ratio vs. control
−3
−2
−1
0
1
2
3

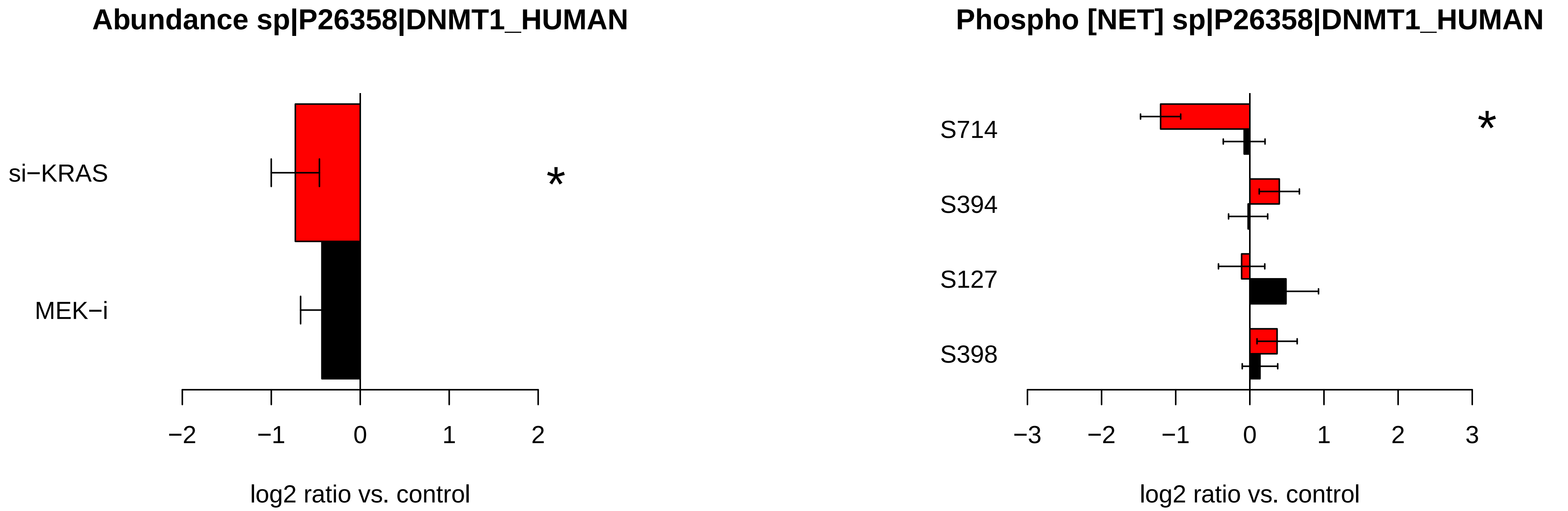

Abundance sp|P26358|DNMT1_HUMAN
si−KRAS
MEK−i
*
−2
−1
0
1
2
log2 ratio vs. control
Phospho [NET] sp|P26358|DNMT1_HUMAN
S714
S394
S127
S398
*
−3
−2
−1
0
1
2
3
log2 ratio vs. control

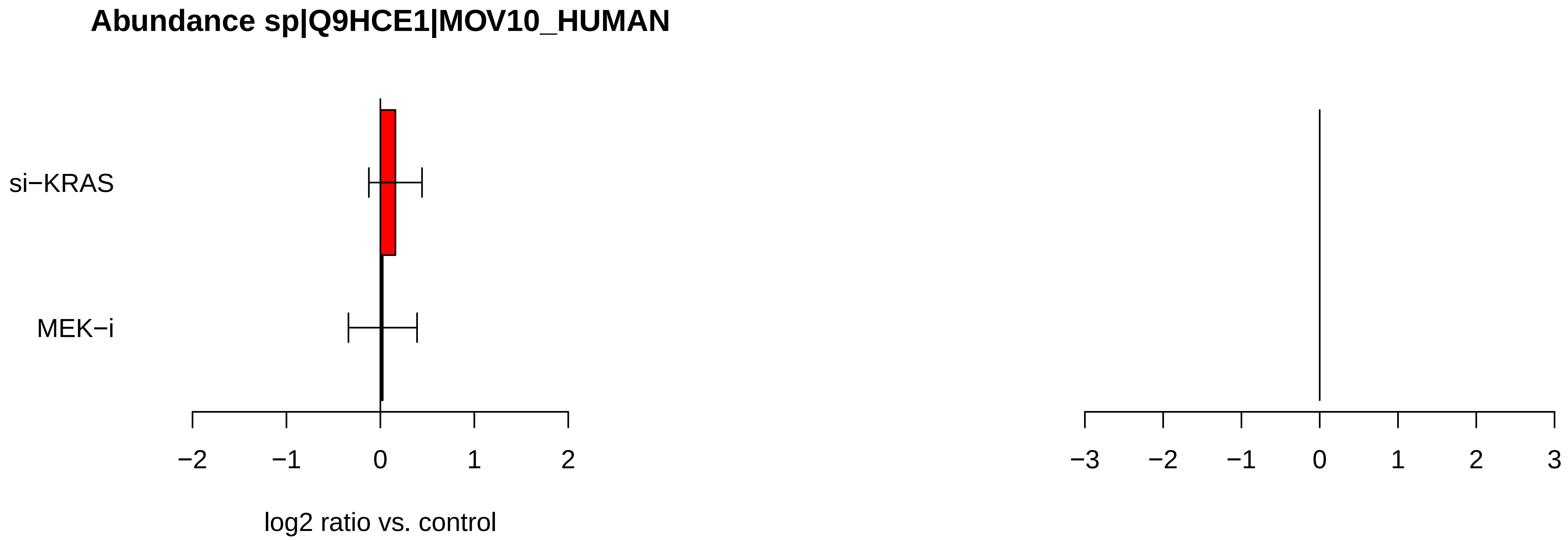
Abundance sp|Q9HCE1|MOV10_HUMAN
si−KRAS
MEK−i
−2
−1
0
1
2
log2 ratio vs. control
−3
−2
−1
0
1
2
3

**Abundance sp|O00459|P85B_HUMAN**

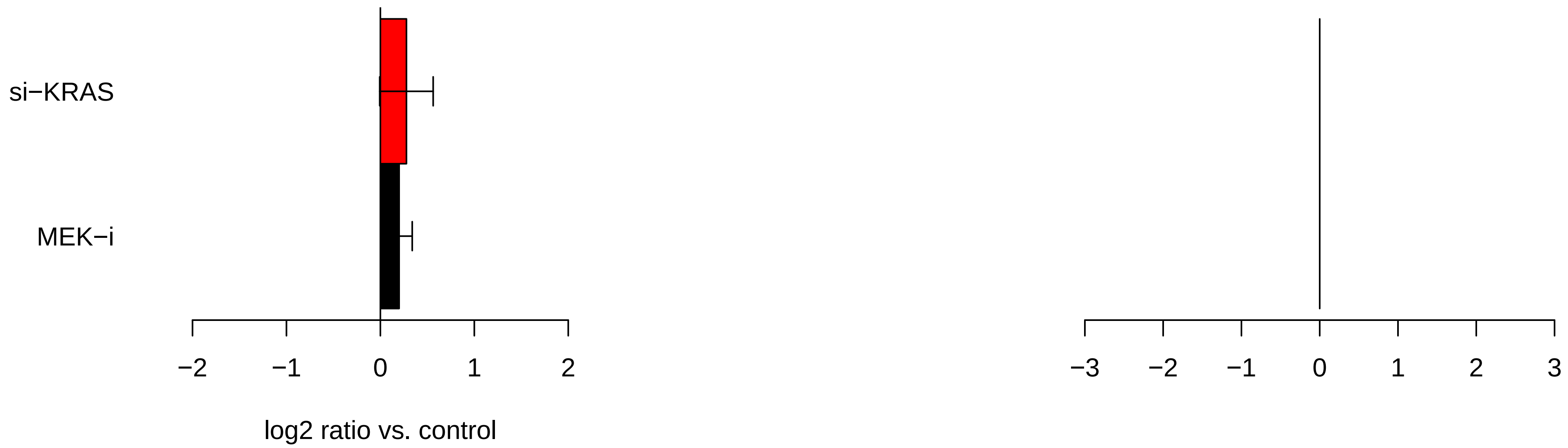

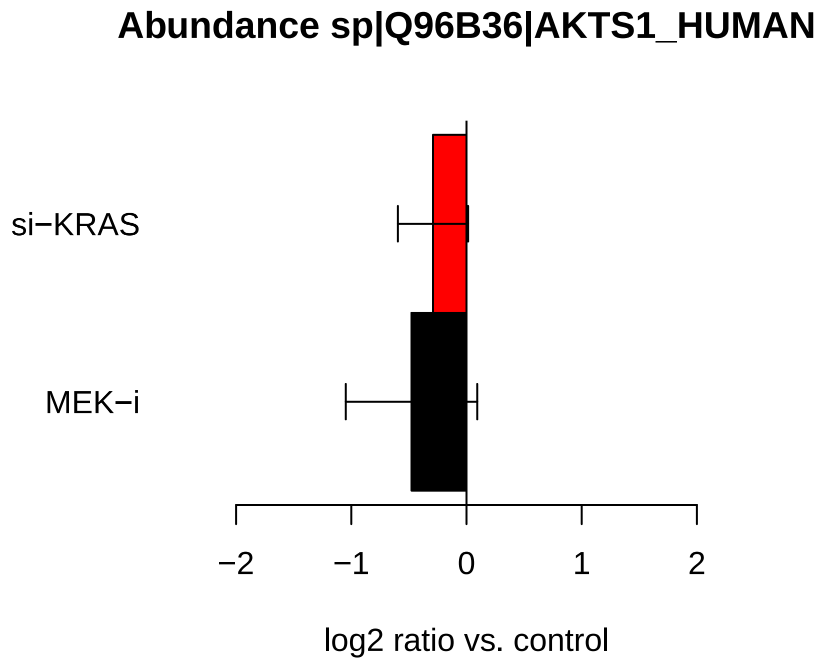

Abundance sp|Q96B36|AKTS1_HUMAN
si−KRAS
MEK−i
−2
−1
0
1
2
log2 ratio vs. control


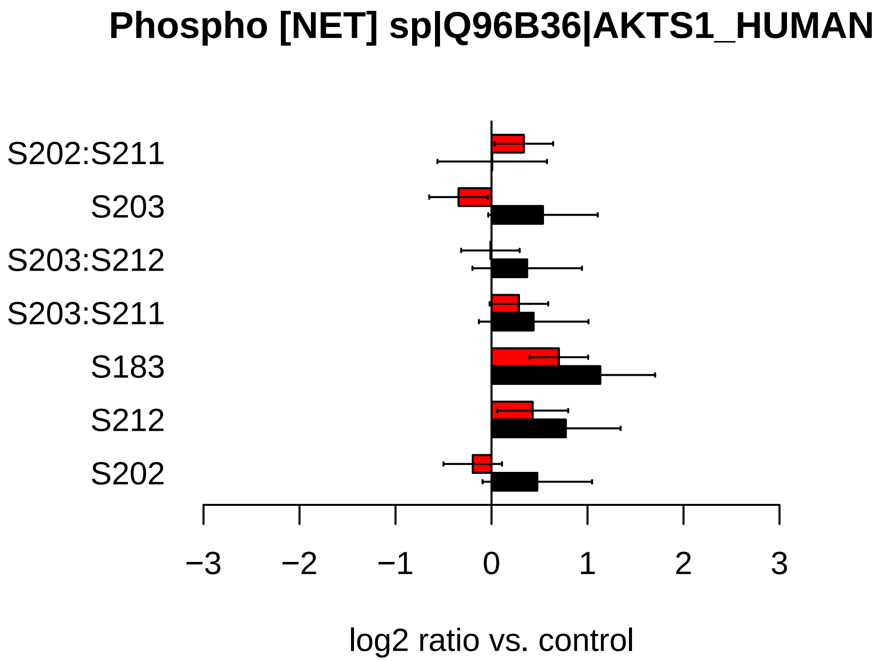

Phospho [NET] sp|Q96B36|AKTS1_HUMAN
S202:S211
S203
S203:S212
S203:S211
S183
S212
S202
−3
−2
−1
0
1
2
3
log2 ratio vs. control

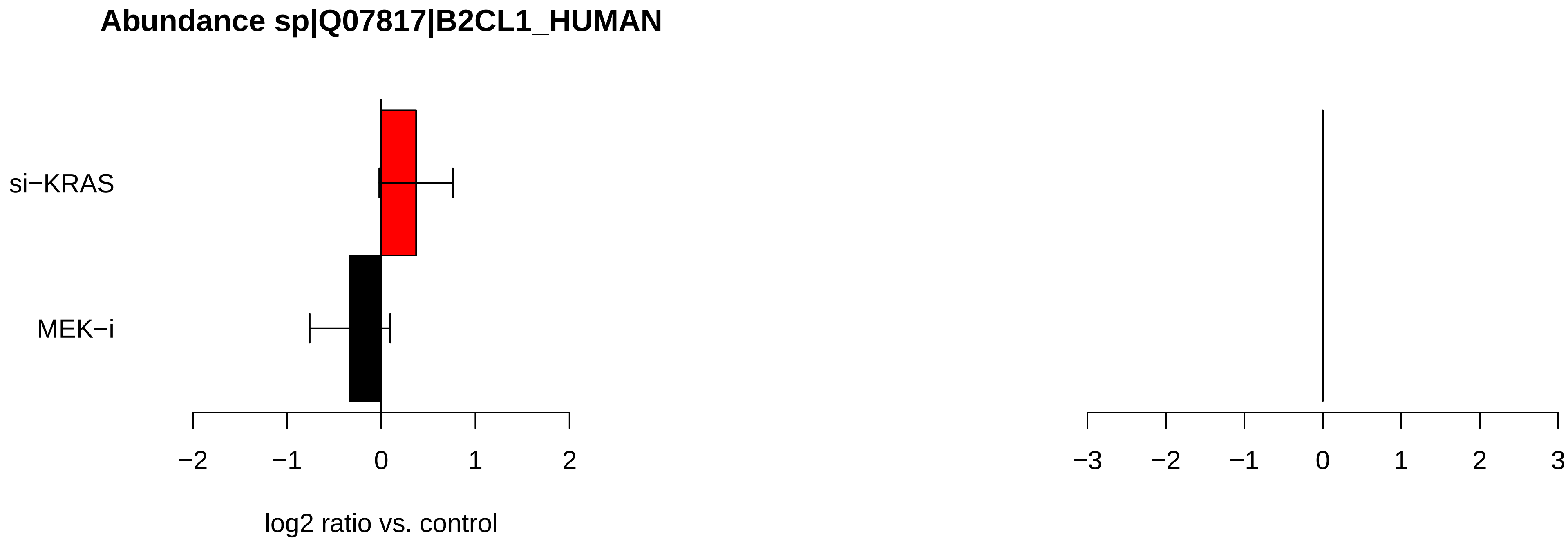
Abundance sp|Q07817|B2CL1_HUMAN
si−KRAS
MEK−i
−2
−1
0
1
2
log2 ratio vs. control
−3
−2
−1
0
1
2
3

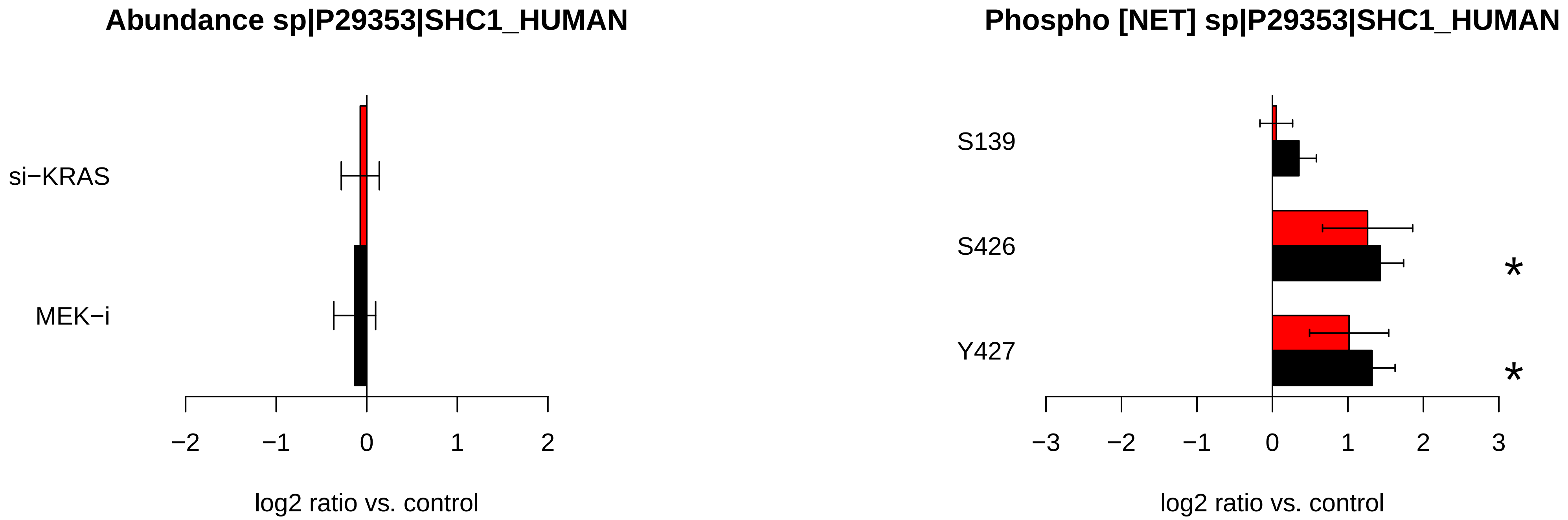
Abundance sp|P29353|SHC1_HUMAN
si−KRAS
MEK−i
−2
−1
0
1
2
log2 ratio vs. control
Phospho [NET] sp|P29353|SHC1_HUMAN
S139
S426
Y427
*
*
−3
−2
−1
0
1
2
3
log2 ratio vs. control

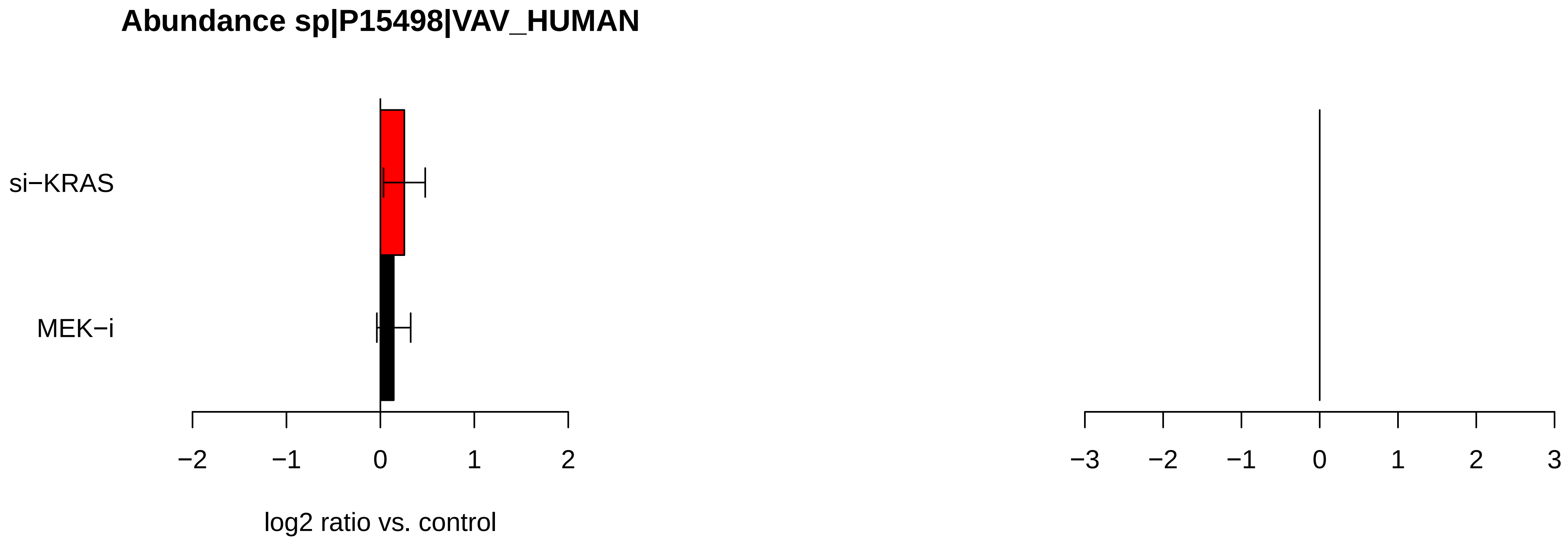

Abundance sp|P15498|VAV_HUMAN
si−KRAS
MEK−i
−2
−1
0
1
2
log2 ratio vs. control
−3
−2
−1
0
1
2
3

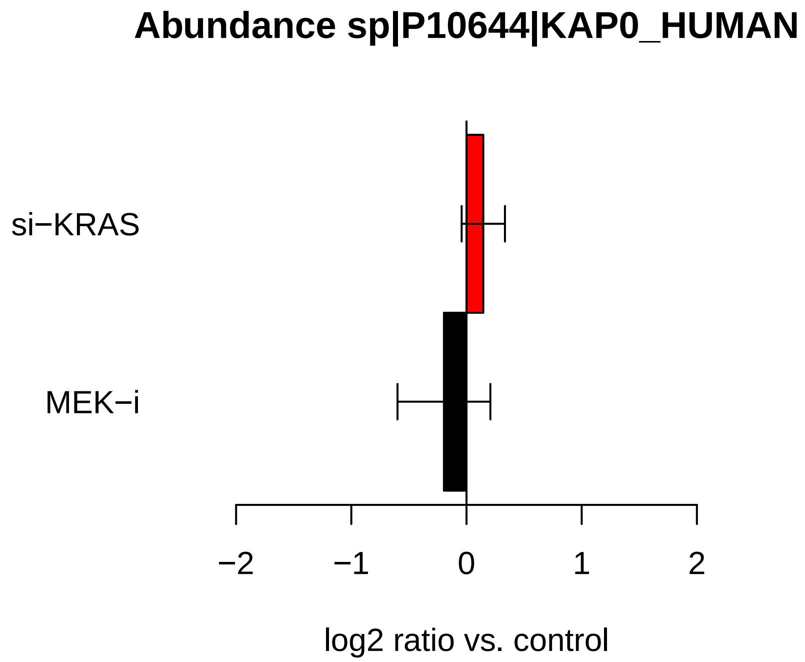
Abundance sp|P10644|KAP0_HUMAN
si−KRAS
MEK−i
−2
−1
0
1
2
log2 ratio vs. control
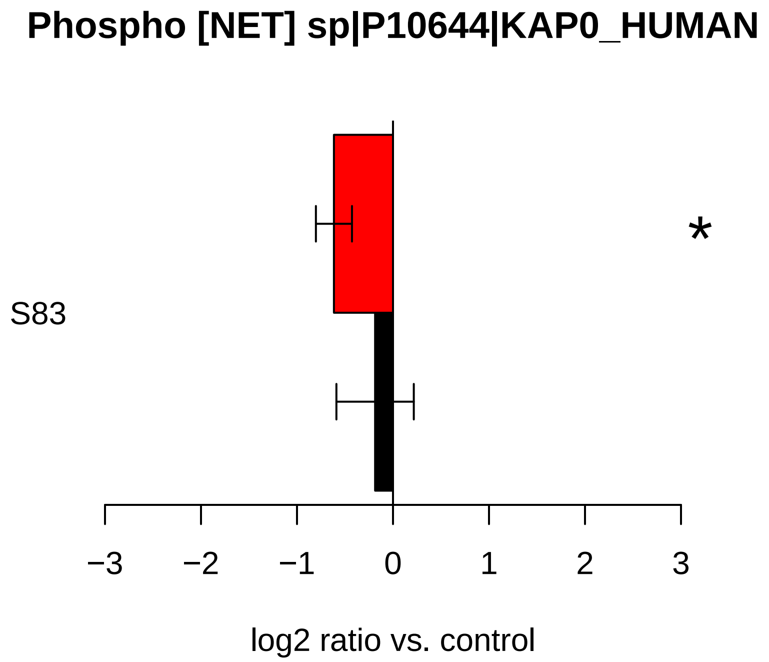
Phospho [NET] sp|P10644|KAP0_HUMAN
S83
*
−3
−2
−1
0
1
2
3
log2 ratio vs. control

### Abundance sp|P19174|PLCG1_HUMAN

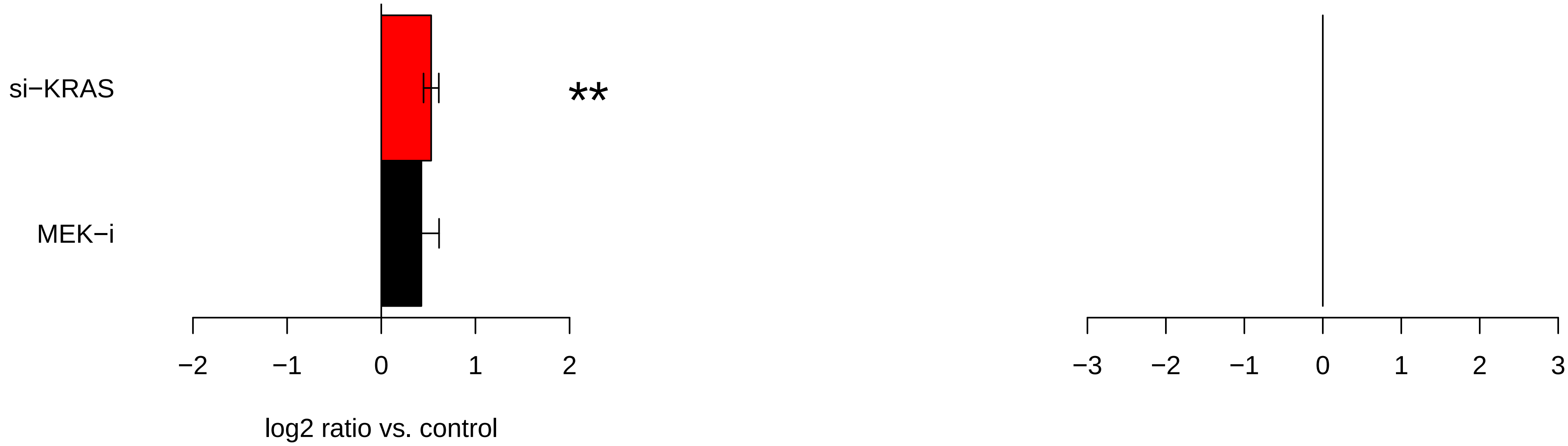

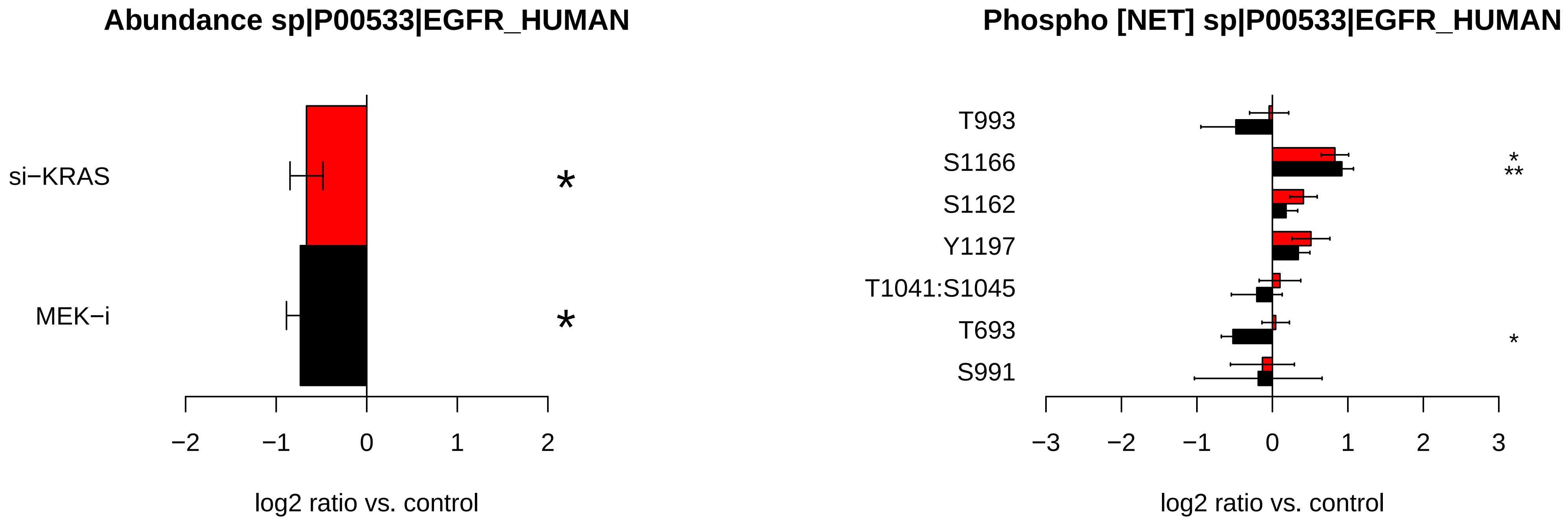
Abundance sp|P00533|EGFR_HUMAN
si−KRAS
MEK−i
*
*
−2
−1
0
1
2
log2 ratio vs. control
Phospho [NET] sp|P00533|EGFR_HUMAN
T993
S1166
S1162
Y1197
T1041:S1045
T693
S991
*
**
*
−3
−2
−1
0
1
2
3
log2 ratio vs. control

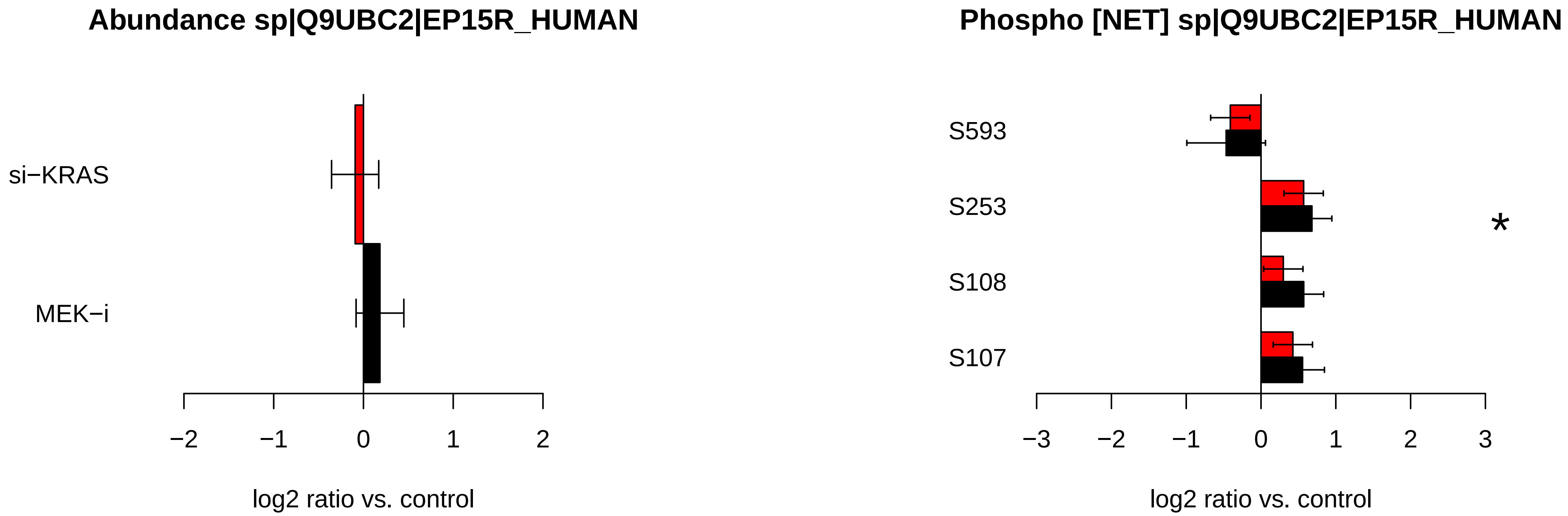
Abundance sp|Q9UBC2|EP15R_HUMAN
si−KRAS
MEK−i
−2
−1
0
1
2
log2 ratio vs. control
Phospho [NET] sp|Q9UBC2|EP15R_HUMAN
S593
S253
S108
S107
*
−3
−2
−1
0
1
2
3
log2 ratio vs. control

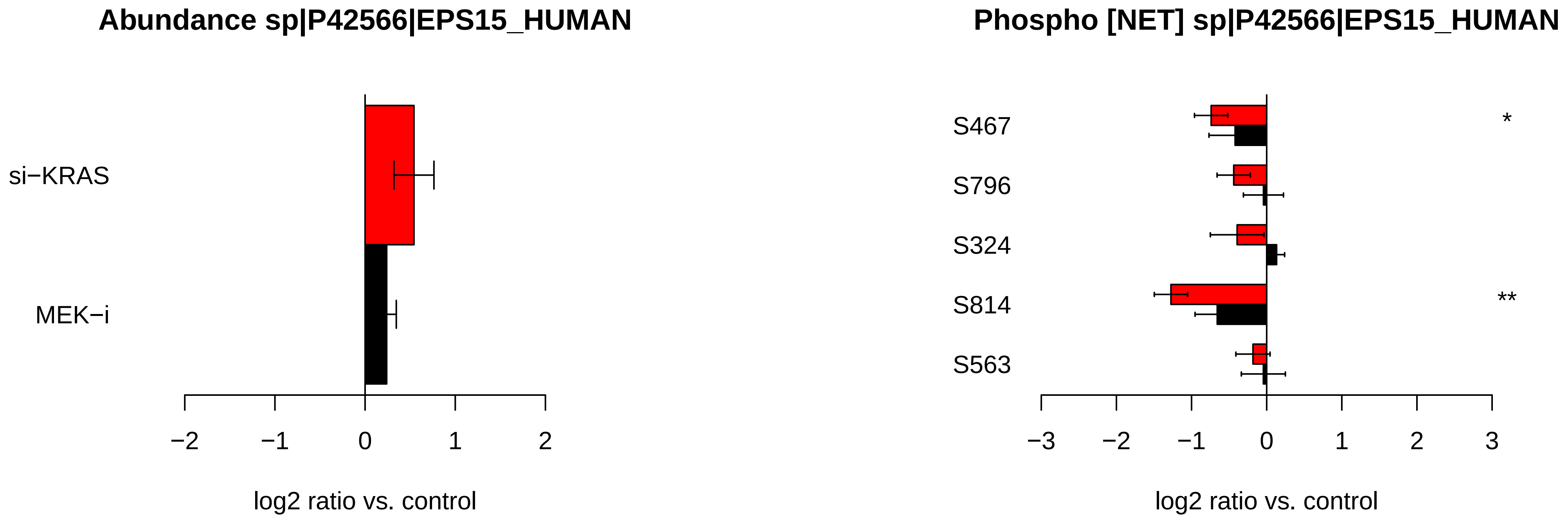

Abundance sp|P42566|EPS15_HUMAN
si−KRAS
MEK−i
−2
−1
0
1
2
log2 ratio vs. control
Phospho [NET] sp|P42566|EPS15_HUMAN
S467
S796
S324
S814
S563
*
**
−3
−2
−1
0
1
2
3
log2 ratio vs. control

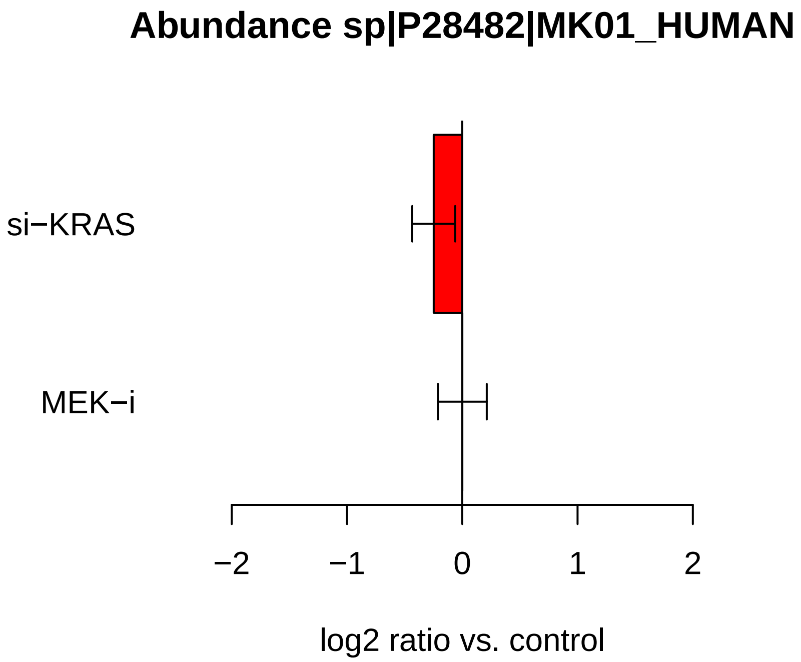

Abundance sp|P28482|MK01_HUMAN
si−KRAS
MEK−i
−2
−1
0
1
2
log2 ratio vs. control


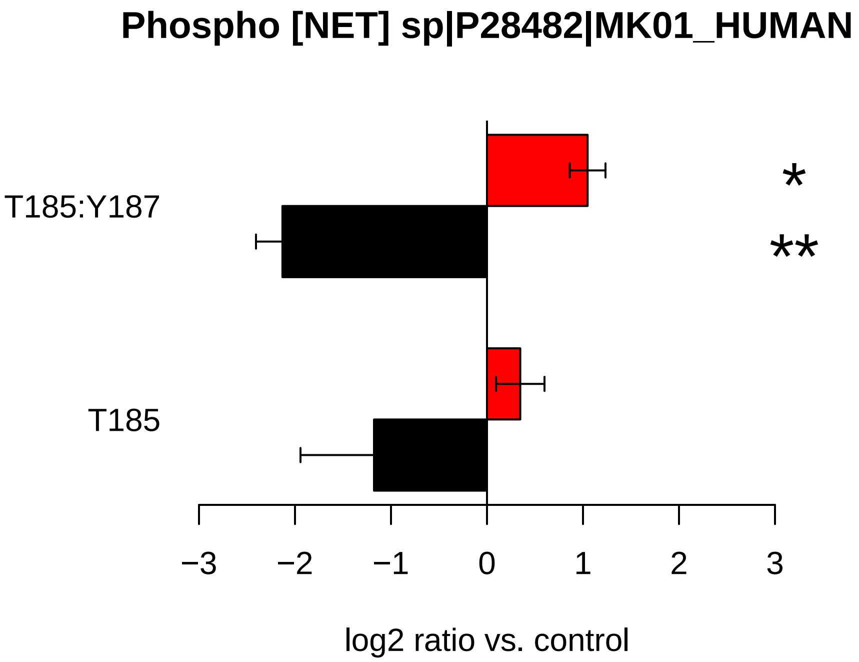

Phospho [NET] sp|P28482|MK01_HUMAN
T185:Y187
T185
*
**
−3
−2
−1
0
1
2
3
log2 ratio vs. control

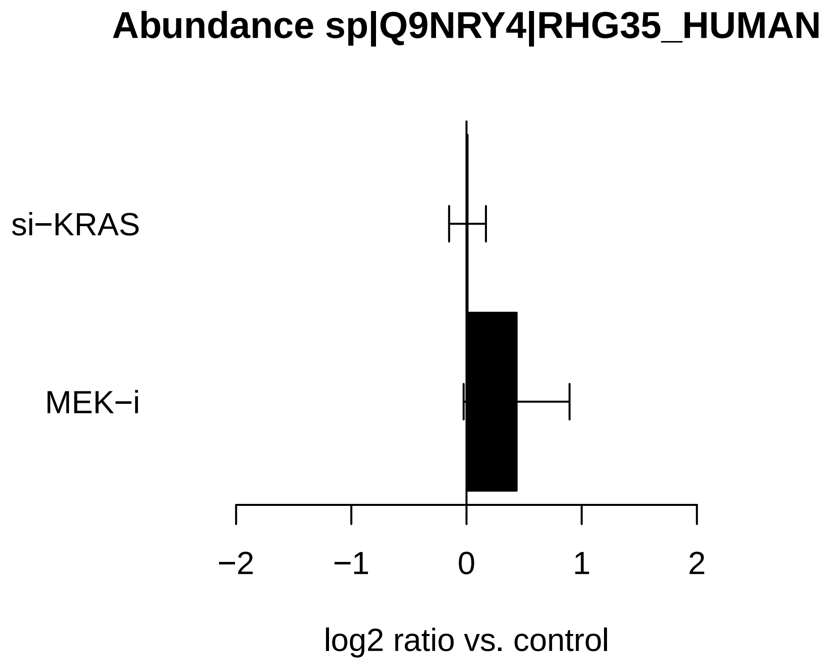

Abundance sp|Q9NRY4|RHG35_HUMAN
si−KRAS
MEK−i
−2
−1
0
1
2
log2 ratio vs. control


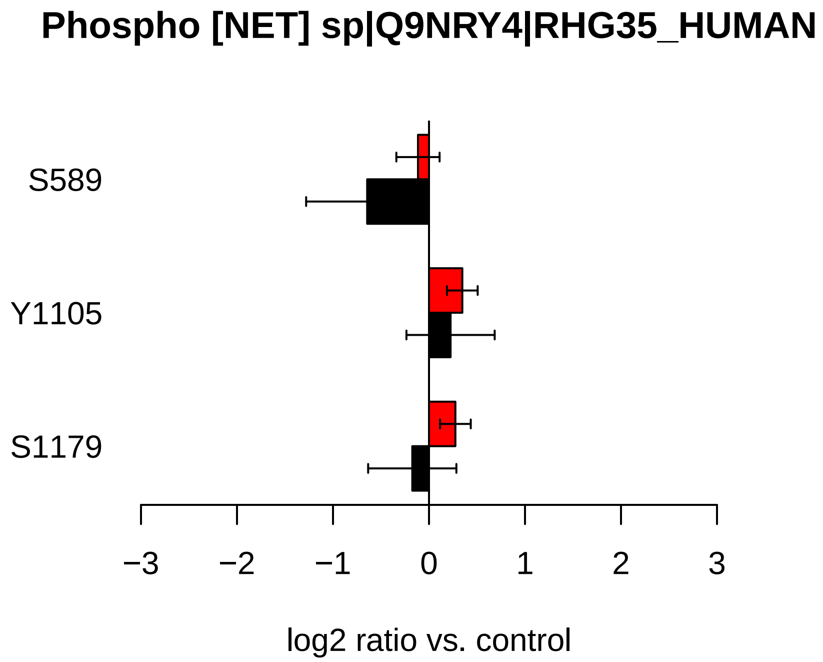

Phospho [NET] sp|Q9NRY4|RHG35_HUMAN
S589
Y1105
S1179
−3
−2
−1
0
1
2
3
log2 ratio vs. control

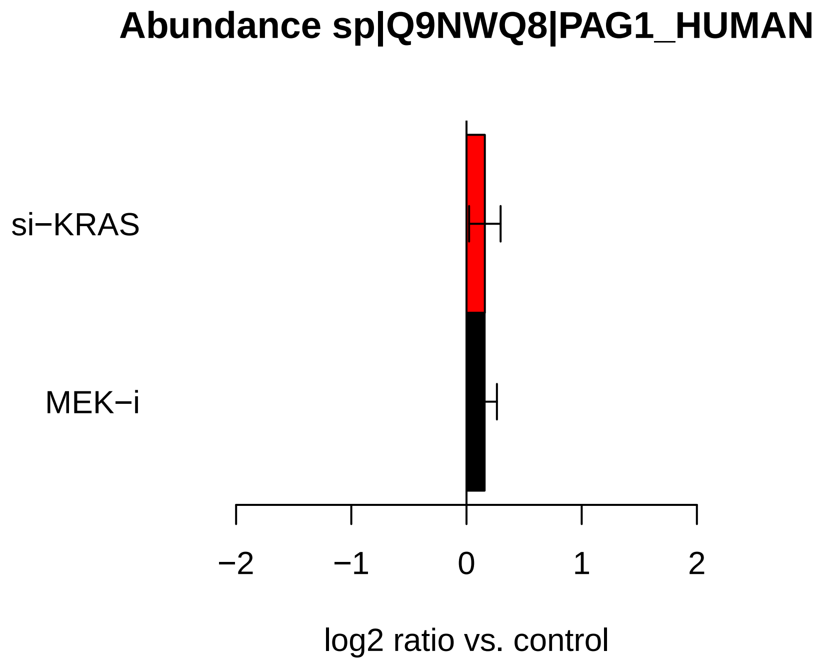

Abundance sp|Q9NWQ8|PAG1_HUMAN
si−KRAS
MEK−i
−2
−1
0
1
2
log2 ratio vs. control


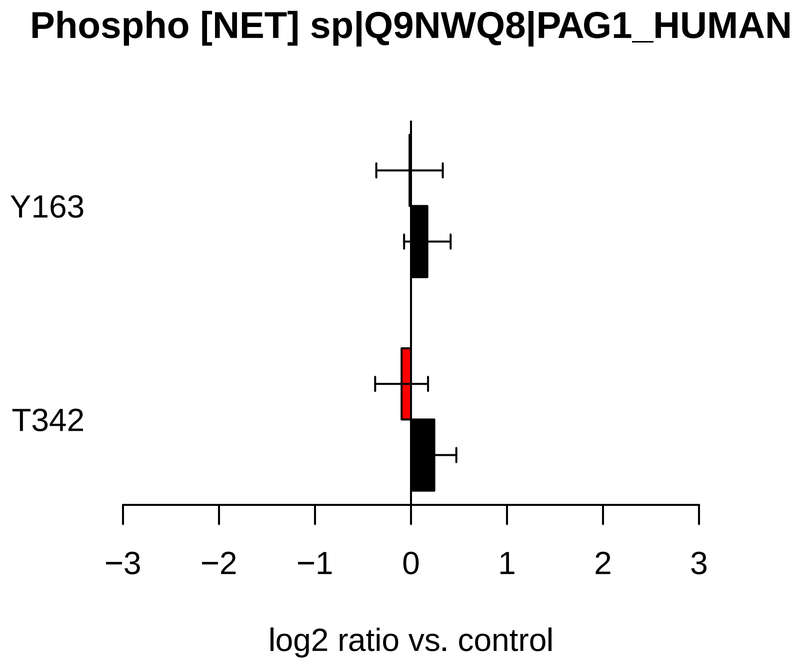

Phospho [NET] sp|Q9NWQ8|PAG1_HUMAN
Y163
T342
−3
−2
−1
0
1
2
3
log2 ratio vs. control

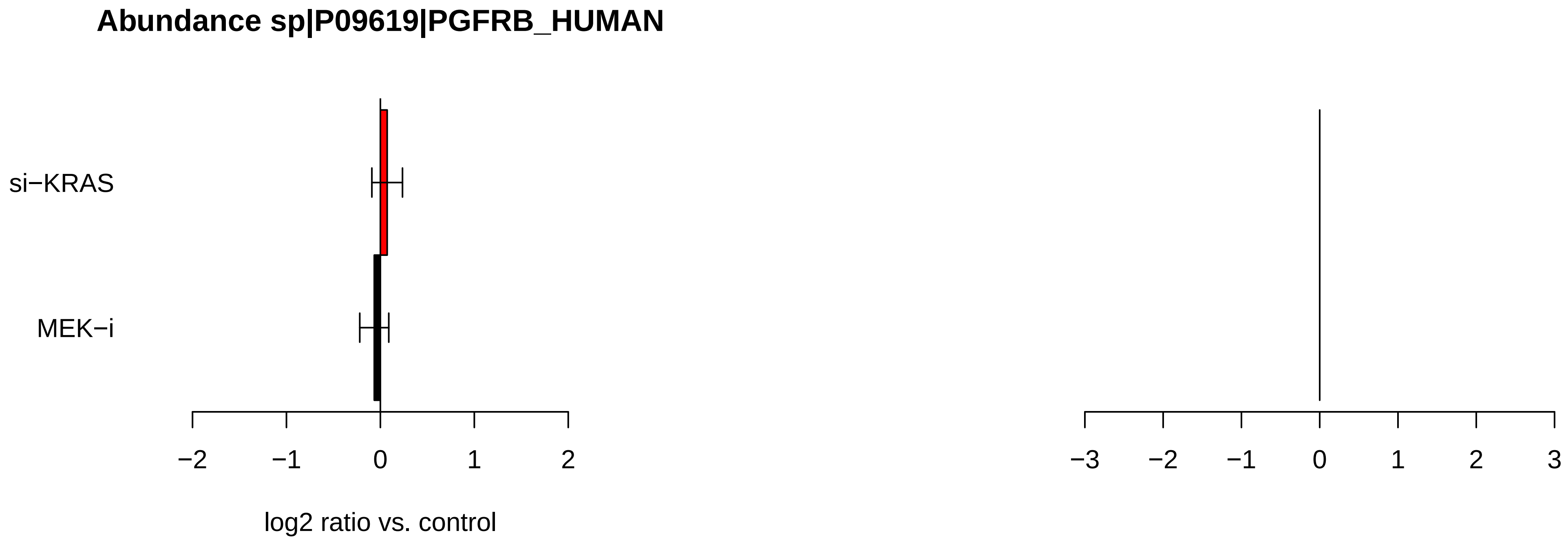

Abundance sp|P09619|PGFRB_HUMAN
si−KRAS
MEK−i
−2
−1
0
1
2
log2 ratio vs. control
−3
−2
−1
0
1
2
3

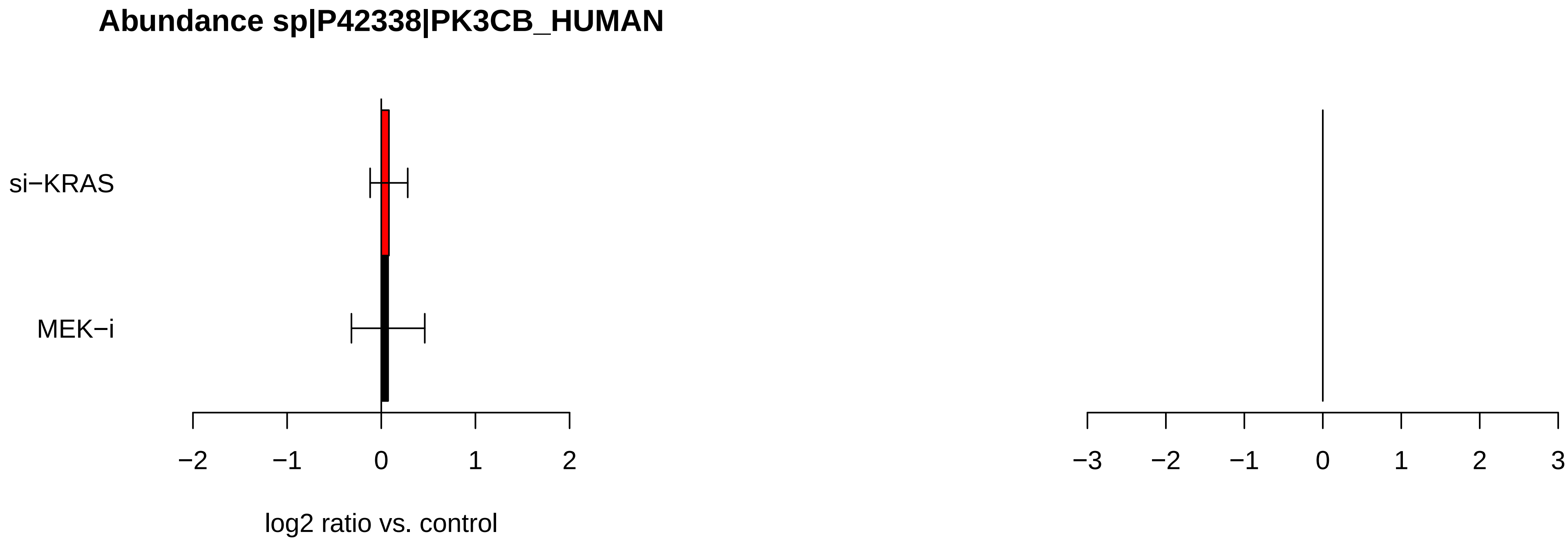

Abundance sp|P42338|PK3CB_HUMAN
si−KRAS
MEK−i
−2
−1
0
1
2
log2 ratio vs. control
−3
−2
−1
0
1
2
3

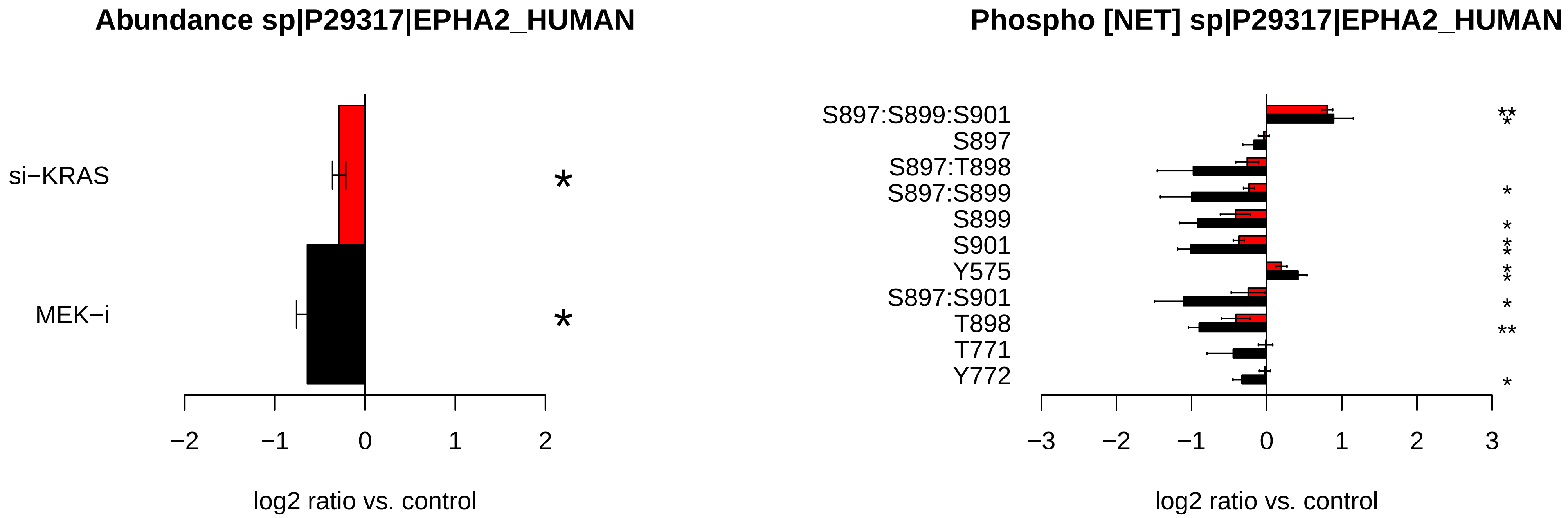

Abundance sp|P29317|EPHA2_HUMAN
si−KRAS
MEK−i
*
*
−2
−1
0
1
2
log2 ratio vs. control
Phospho [NET] sp|P29317|EPHA2_HUMAN
S897:S899:S901
S897
S897:T898
S897:S899
S899
S901
Y575
S897:S901
T898
T771
Y772
−3
−2
−1
0
1
2
3
log2 ratio vs. control

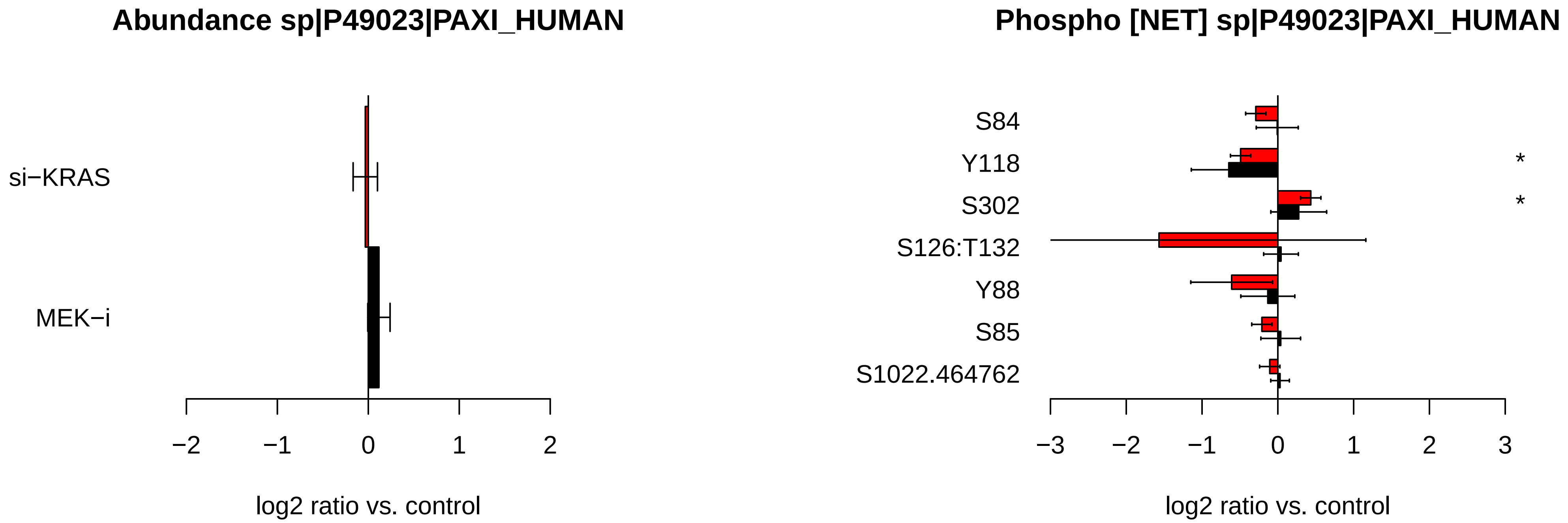
Abundance sp|P49023|PAXI_HUMAN
si−KRAS
MEK−i
−2
−1
0
1
2
log2 ratio vs. control
Phospho [NET] sp|P49023|PAXI_HUMAN
S84
Y118
S302
S126:T132
Y88
S85
S1022.464762
*
*
−3
−2
−1
0
1
2
3
log2 ratio vs. control

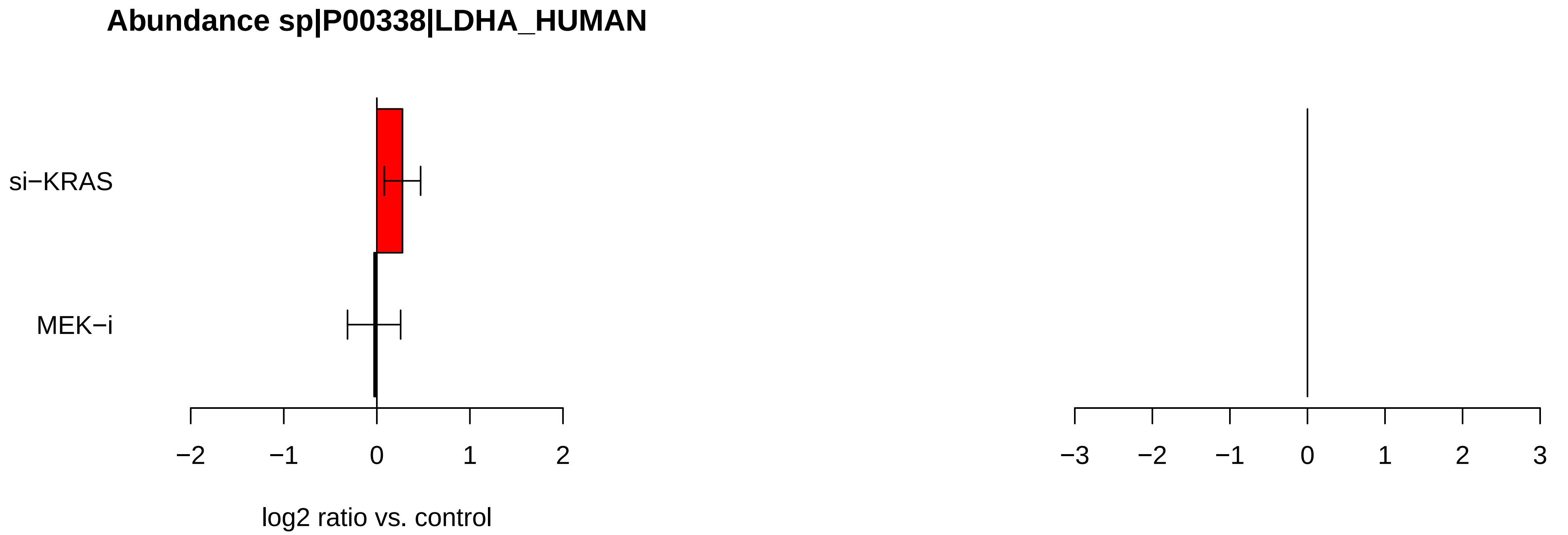
Abundance sp|P00338|LDHA_HUMAN
si−KRAS
MEK−i
−2
−1
0
1
2
log2 ratio vs. control
−3
−2
−1
0
1
2
3

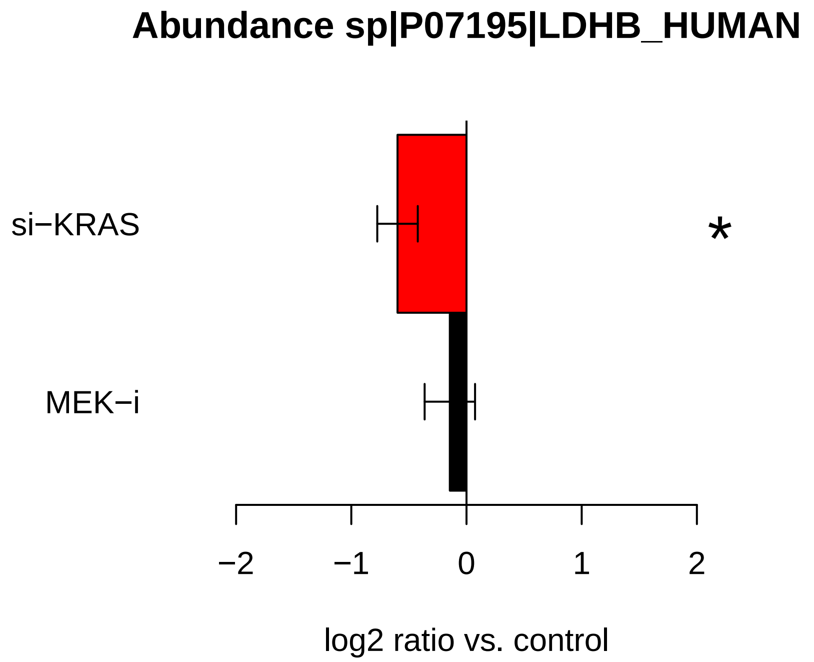
Abundance sp|P07195|LDHB_HUMAN
si−KRAS
MEK−i
*
−2
−1
0
1
2
log2 ratio vs. control

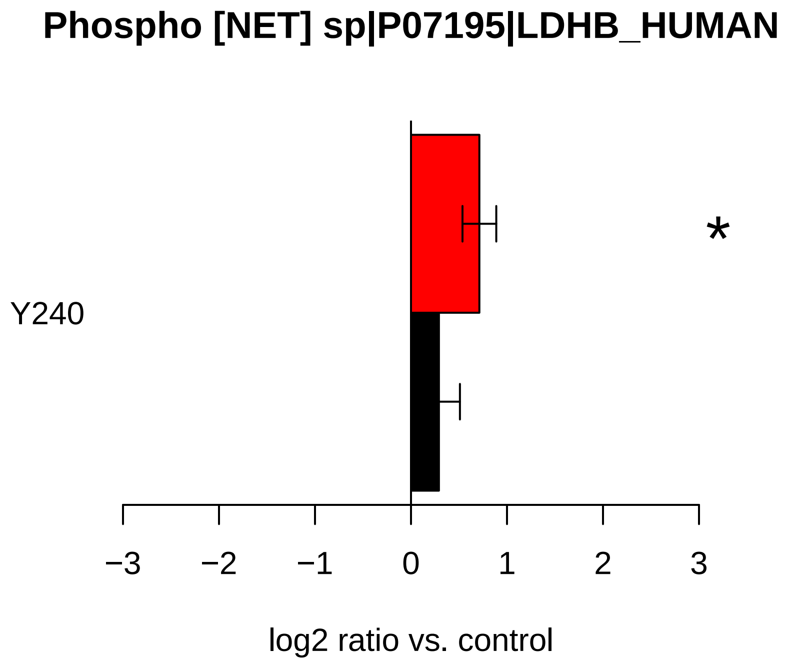
Phospho [NET] sp|P07195|LDHB_HUMAN
Y240
*
−3
−2
−1
0
1
2
3
log2 ratio vs. control

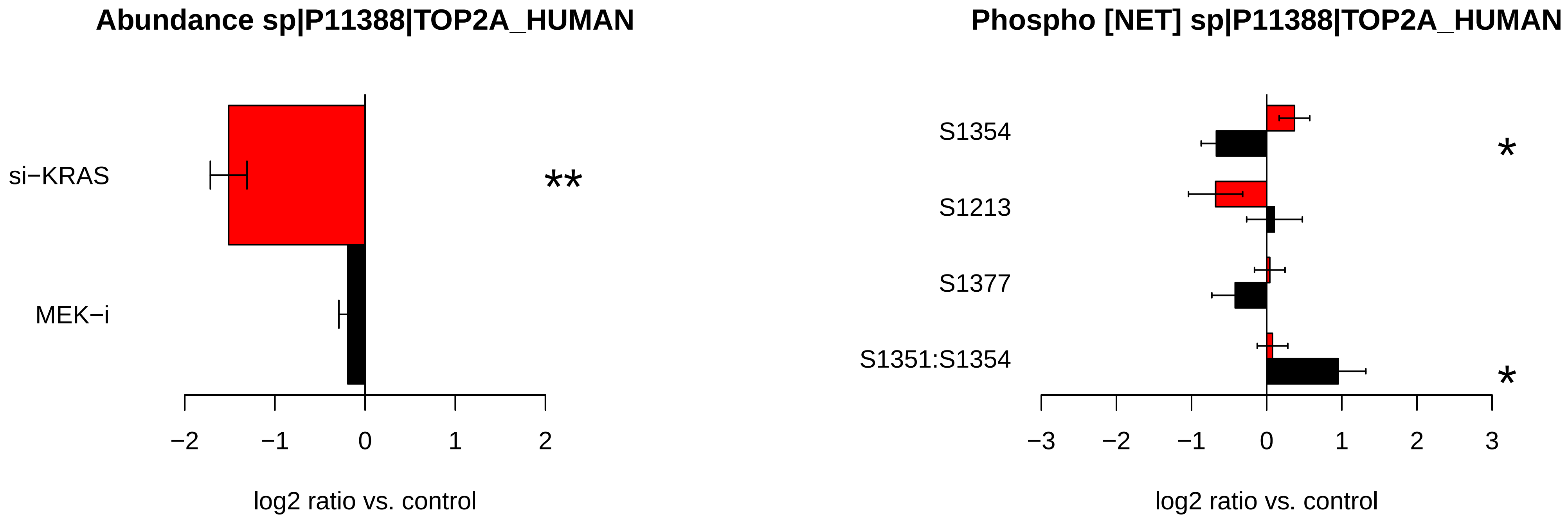
Abundance sp|P11388|TOP2A_HUMAN
si−KRAS
MEK−i
**
−2
−1
0
1
2
log2 ratio vs. control
Phospho [NET] sp|P11388|TOP2A_HUMAN
S1354
S1213
S1377
S1351:S1354
*
*
−3
−2
−1
0
1
2
3
log2 ratio vs. control

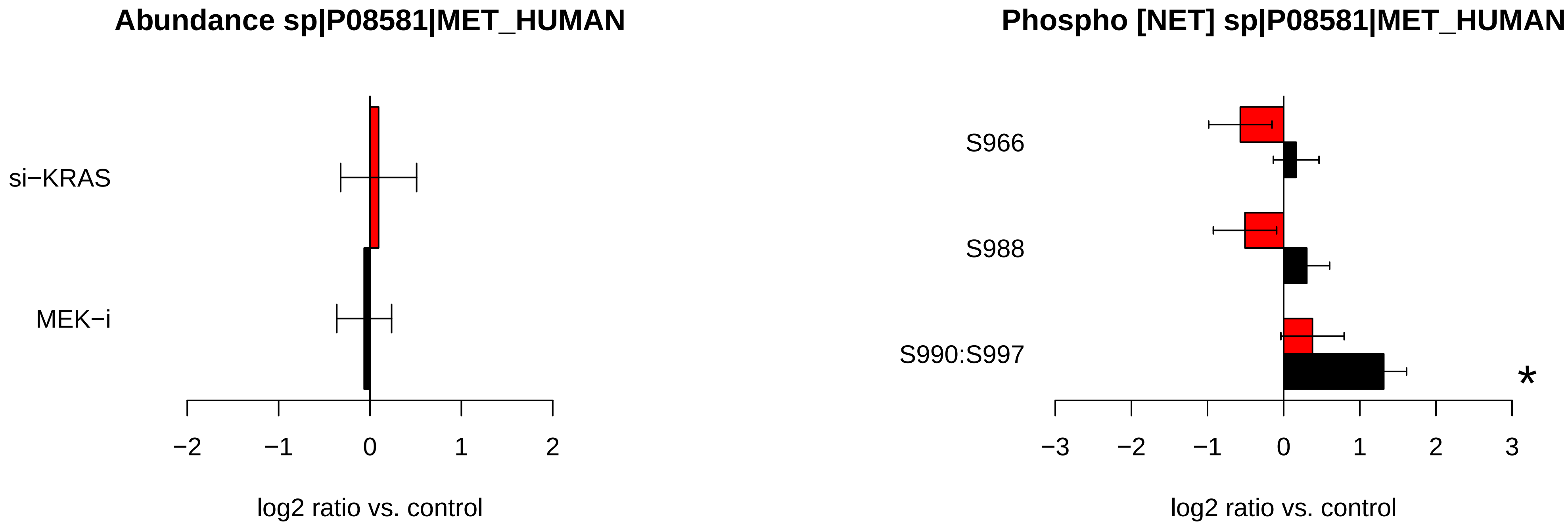
Abundance sp|P08581|MET_HUMAN
si−KRAS
MEK−i
−2
−1
0
1
2
log2 ratio vs. control
Phospho [NET] sp|P08581|MET_HUMAN
S966
S988
S990:S997
*
−3
−2
−1
0
1
2
3
log2 ratio vs. control

### Abundance sp|P46734|MP2K3_HUMAN

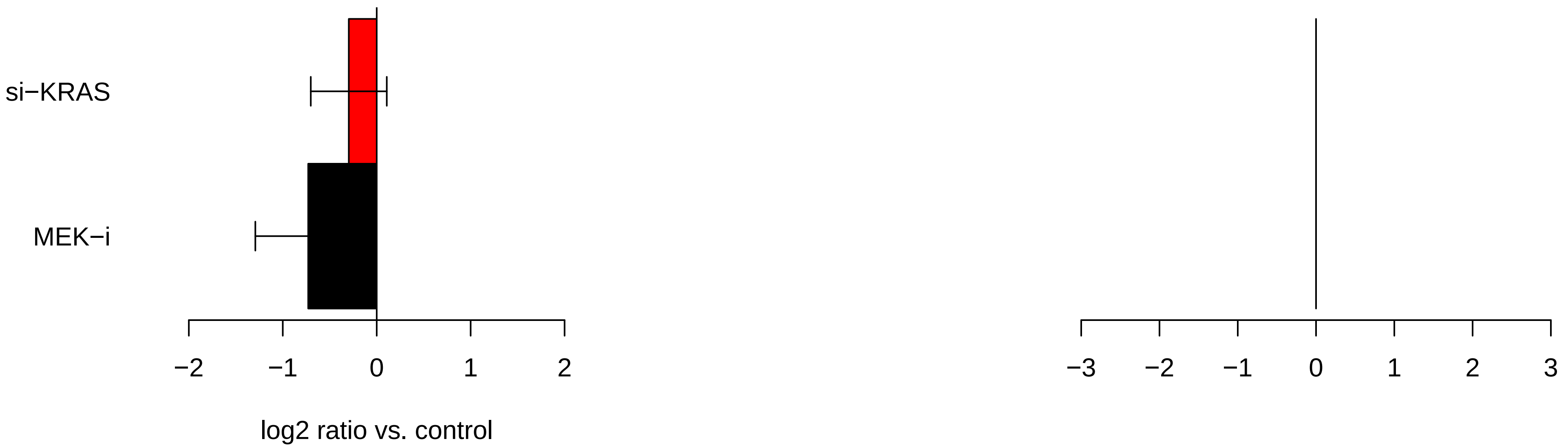

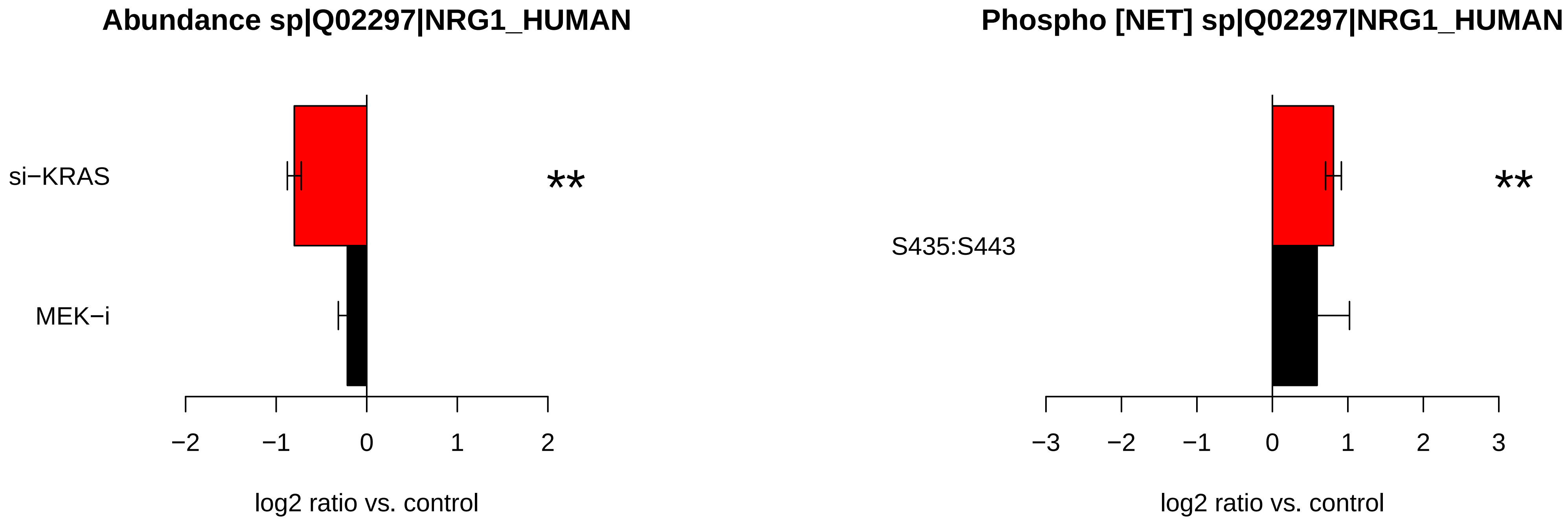

Abundance sp|Q02297|NRG1_HUMAN
si−KRAS
MEK−i
**
−2
−1
0
1
2
log2 ratio vs. control
Phospho [NET] sp|Q02297|NRG1_HUMAN
S435:S443
**
−3
−2
−1
0
1
2
3
log2 ratio vs. control

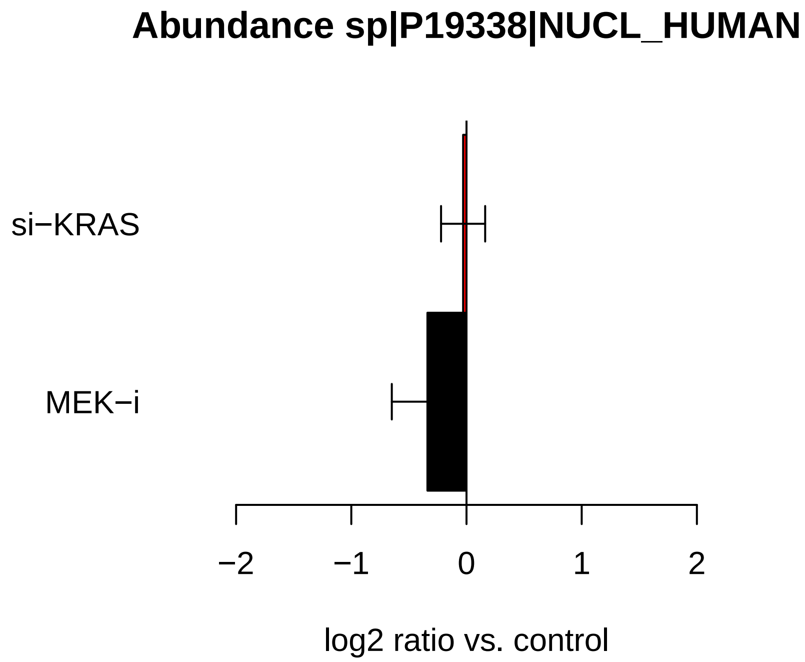

Abundance sp|P19338|NUCL_HUMAN
si−KRAS
MEK−i
−2
−1
0
1
2
log2 ratio vs. control


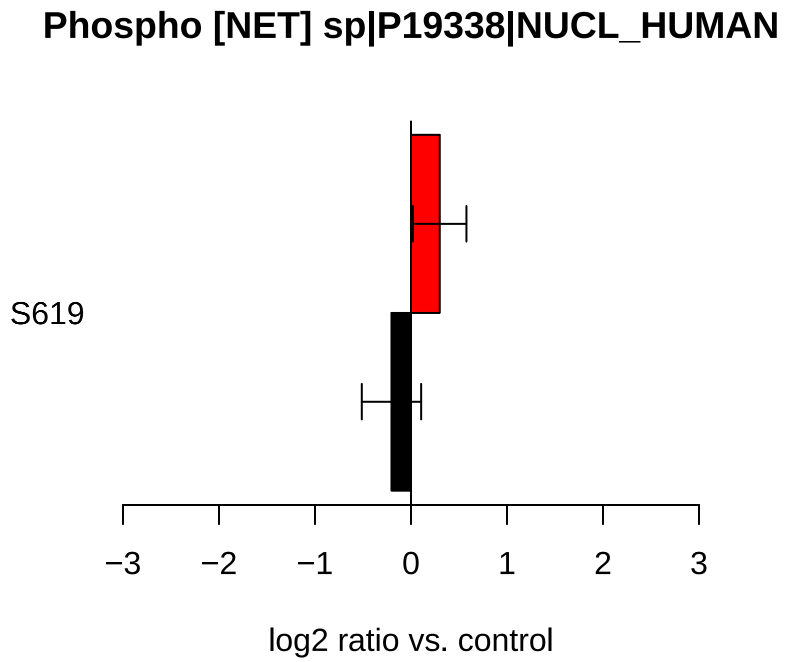

Phospho [NET] sp|P19338|NUCL_HUMAN
S619
−3
−2
−1
0
1
2
3
log2 ratio vs. control

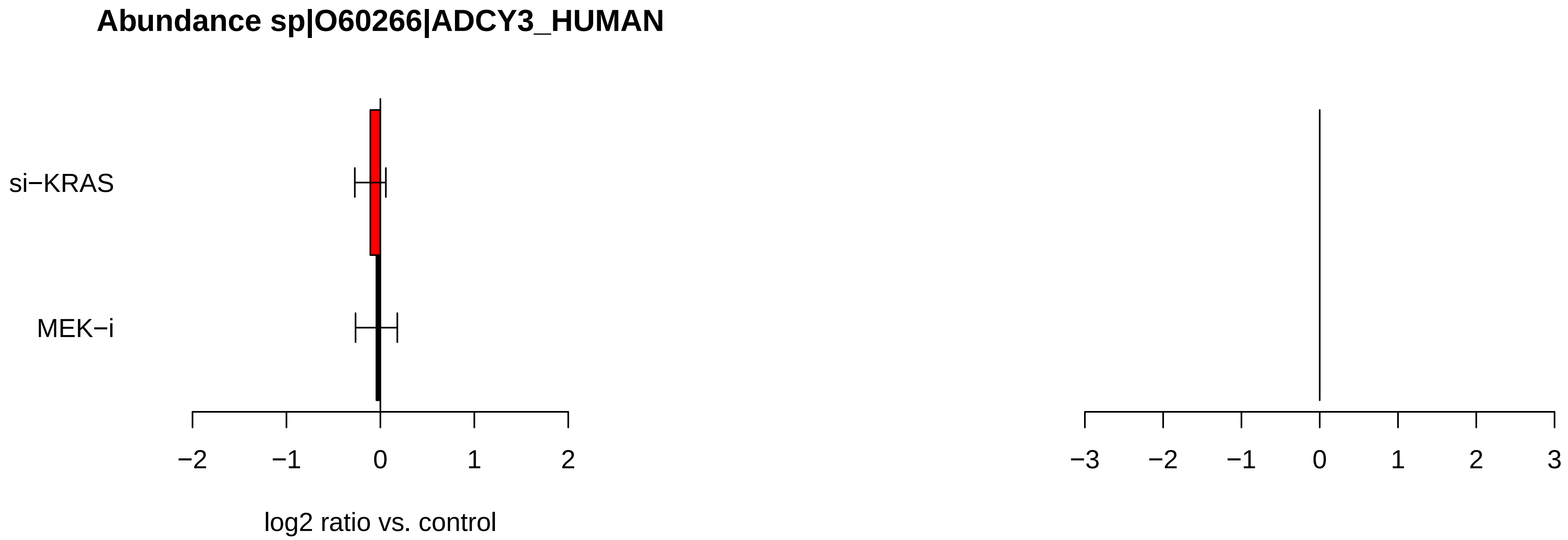

Abundance sp|O60266|ADCY3_HUMAN
si−KRAS
MEK−i
−2
−1
0
1
2
log2 ratio vs. control
−3
−2
−1
0
1
2
3

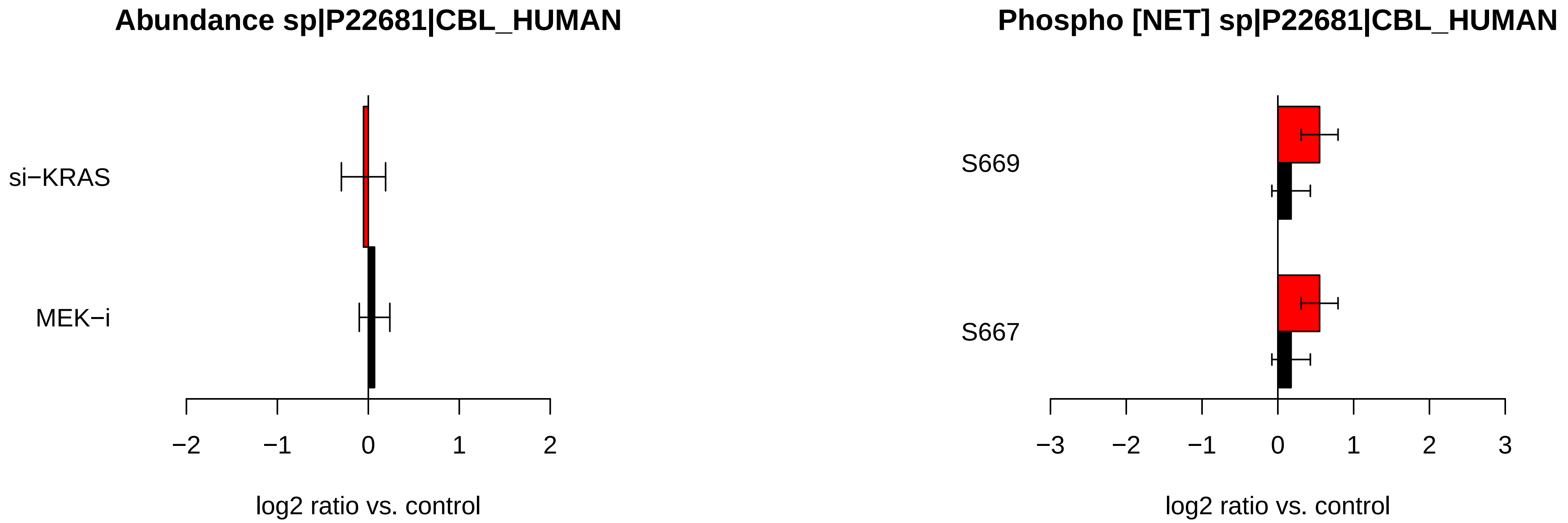

Abundance sp|P22681|CBL_HUMAN
si−KRAS
MEK−i
−2
−1
0
1
2
log2 ratio vs. control
Phospho [NET] sp|P22681|CBL_HUMAN
S669
S667
−3
−2
−1
0
1
2
3
log2 ratio vs. control

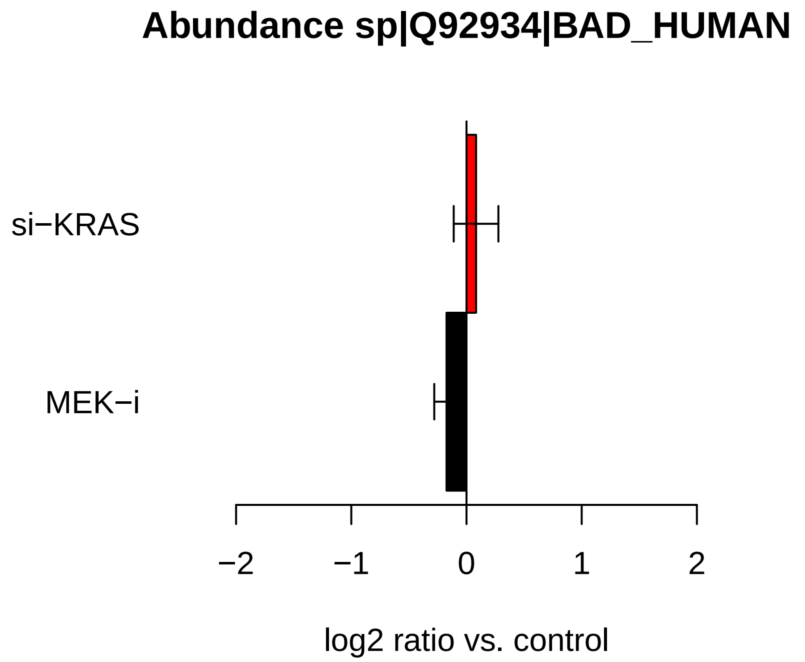

Abundance sp|Q92934|BAD_HUMAN
si−KRAS
MEK−i
−2
−1
0
1
2
log2 ratio vs. control


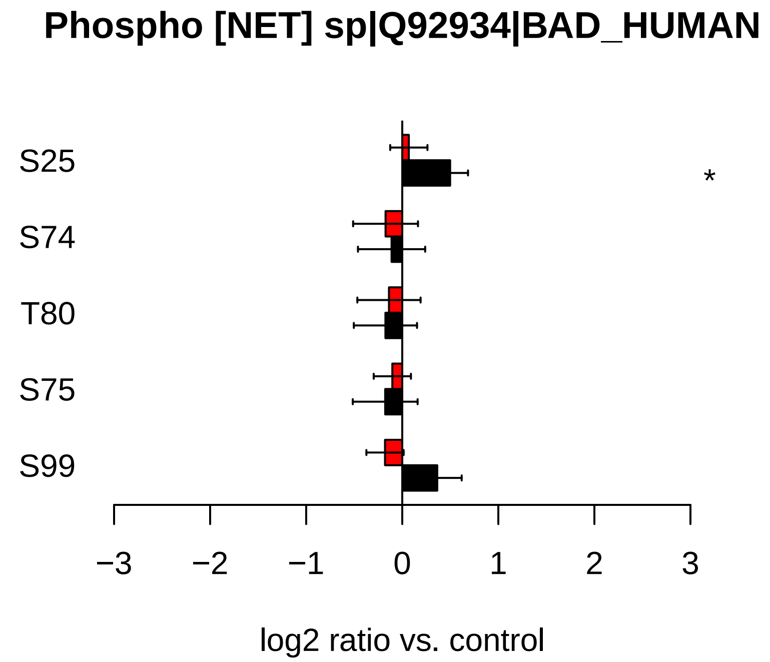

Phospho [NET] sp|Q92934|BAD_HUMAN
S25
S74
T80
S75
S99
*
−3
−2
−1
0
1
2
3
log2 ratio vs. control

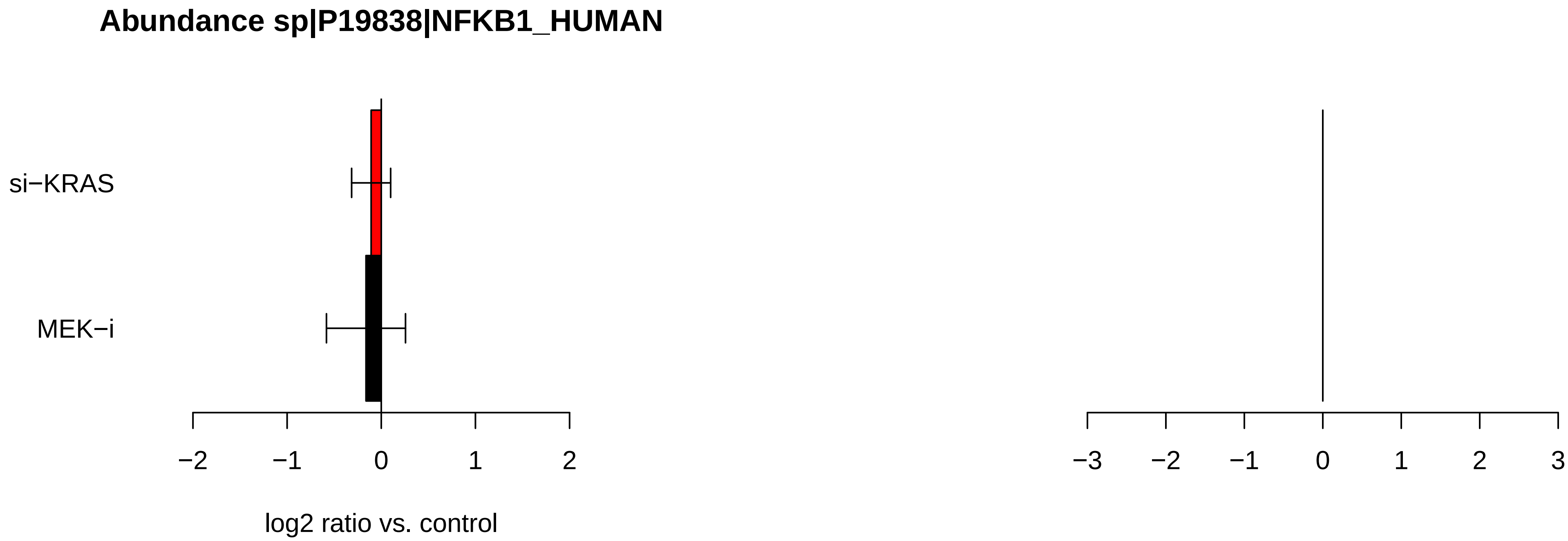

Abundance sp|P19838|NFKB1_HUMAN
si−KRAS
MEK−i
−2
−1
0
1
2
log2 ratio vs. control
−3
−2
−1
0
1
2
3

### Abundance sp|P63000|RAC1_HUMAN

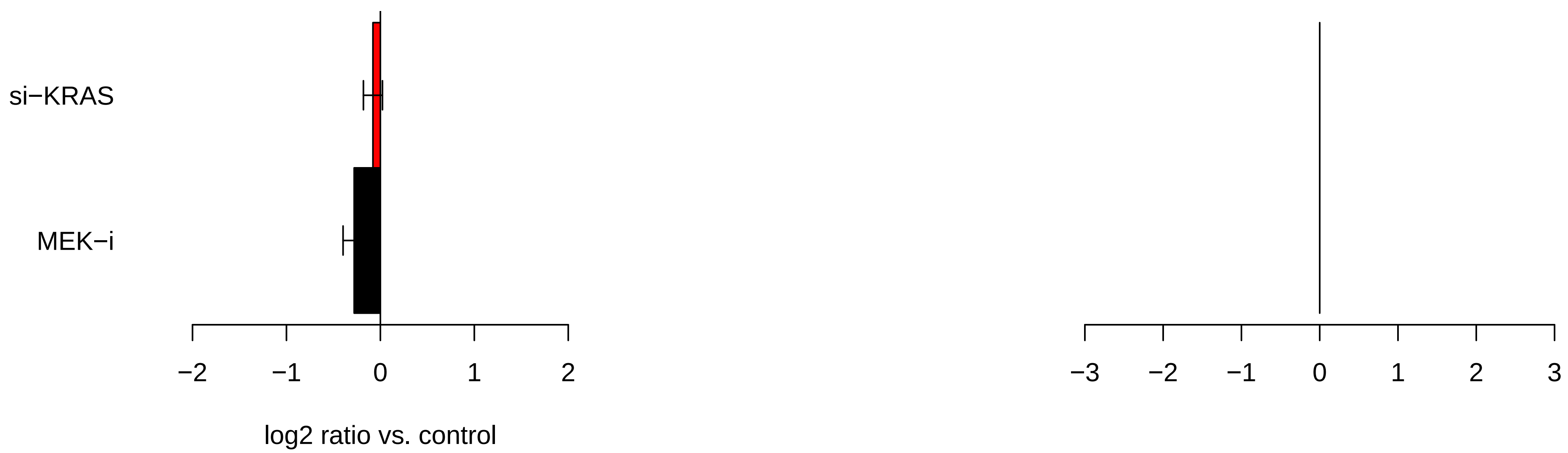

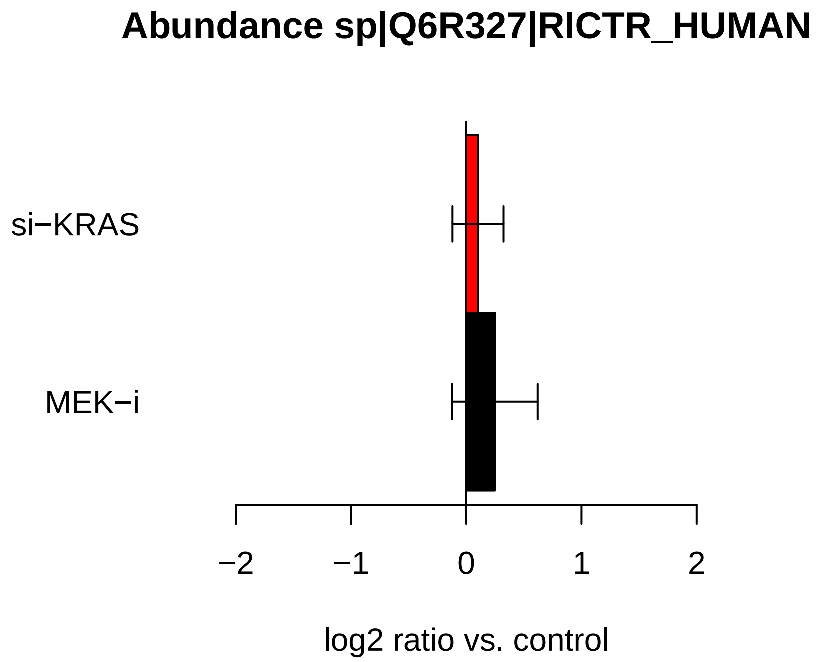

Abundance sp|Q6R327|RICTR_HUMAN
si−KRAS
MEK−i
−2
−1
0
1
2
log2 ratio vs. control


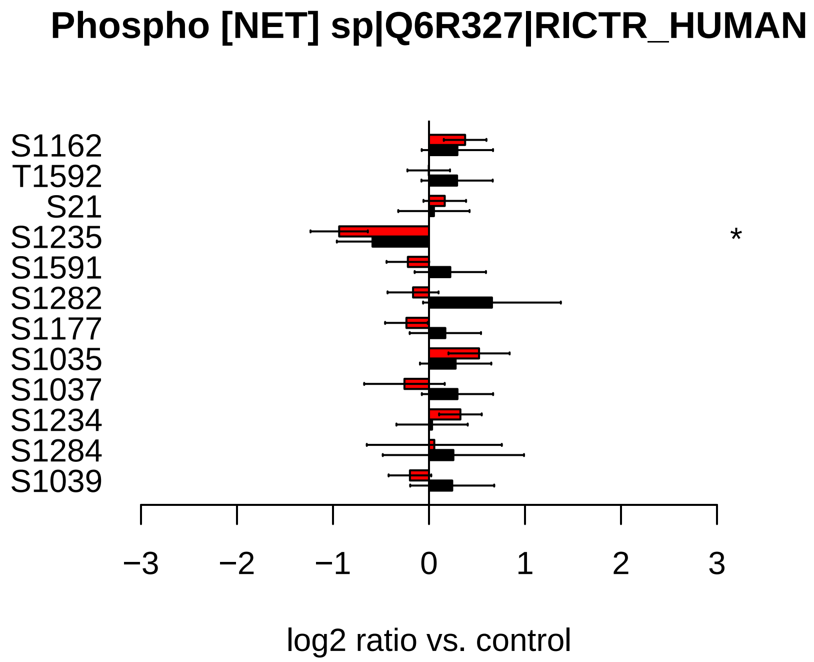

Phospho [NET] sp|Q6R327|RICTR_HUMAN
S1162
T1592
S21
S1235
S1591
S1282
S1177
S1035
S1037
S1234
S1284
S1039
*
−3
−2
−1
0
1
2
3
log2 ratio vs. control

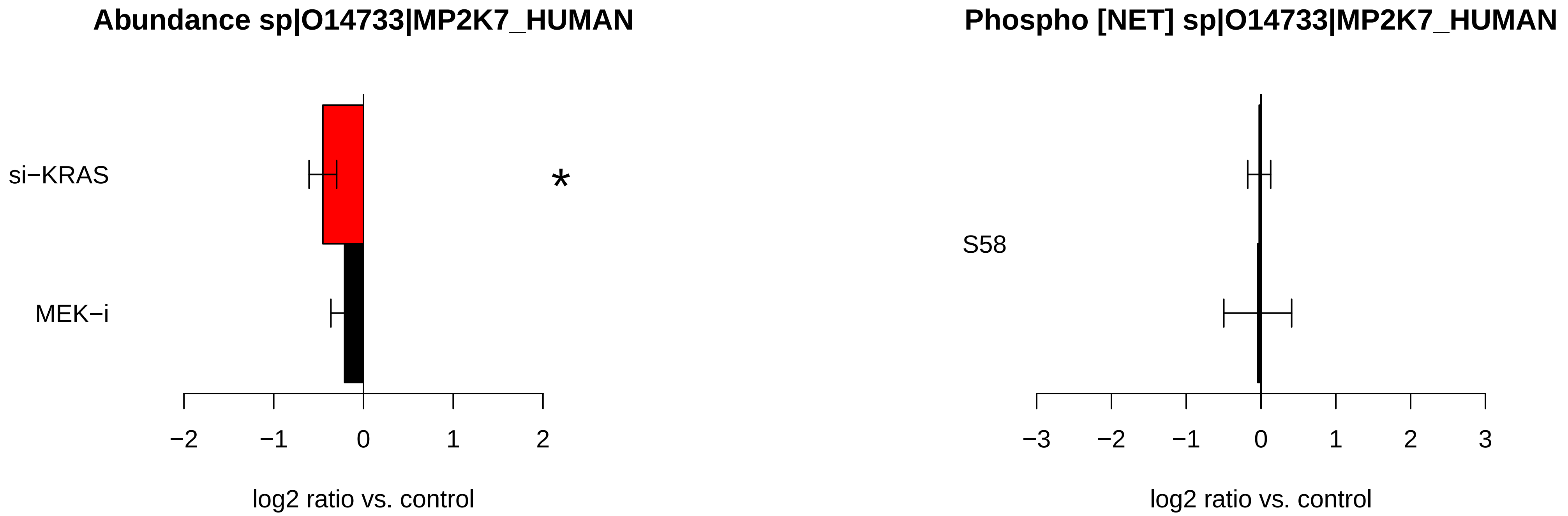

Abundance sp|O14733|MP2K7_HUMAN
si−KRAS
MEK−i
*
−2
−1
0
1
2
log2 ratio vs. control
Phospho [NET] sp|O14733|MP2K7_HUMAN
S58
−3
−2
−1
0
1
2
3
log2 ratio vs. control

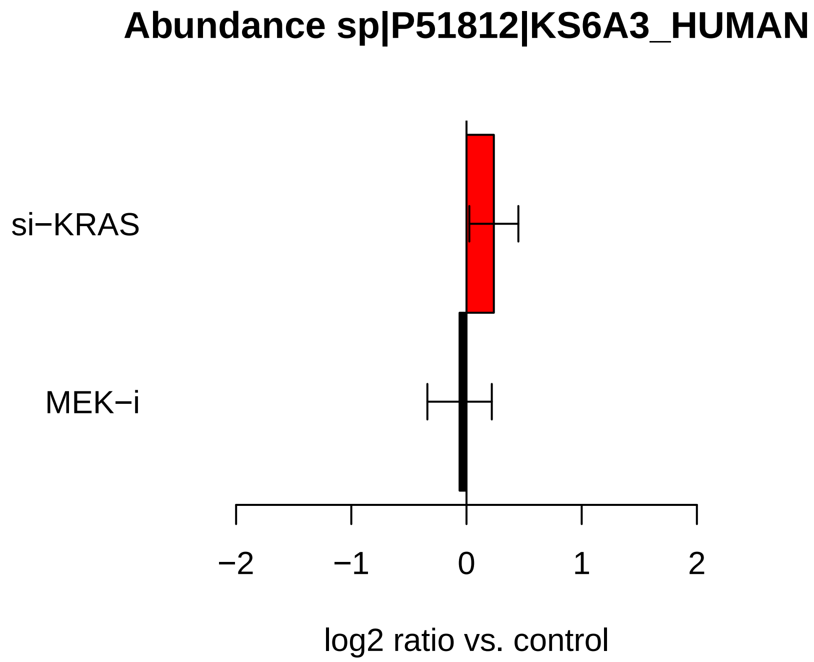
Abundance sp|P51812|KS6A3_HUMAN
si−KRAS
MEK−i
−2
−1
0
1
2
log2 ratio vs. control

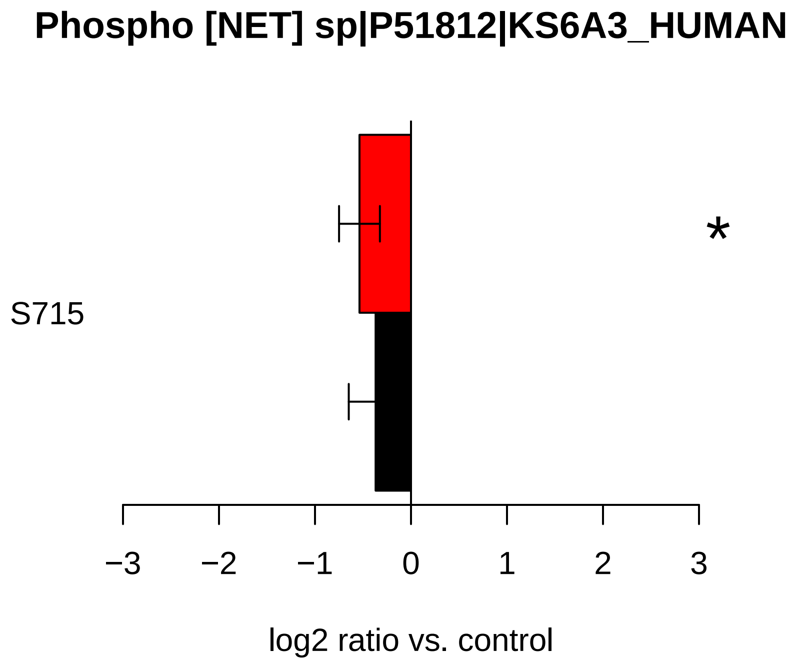
Phospho [NET] sp|P51812|KS6A3_HUMAN
S715
*
−3
−2
−1
0
1
2
3
log2 ratio vs. control

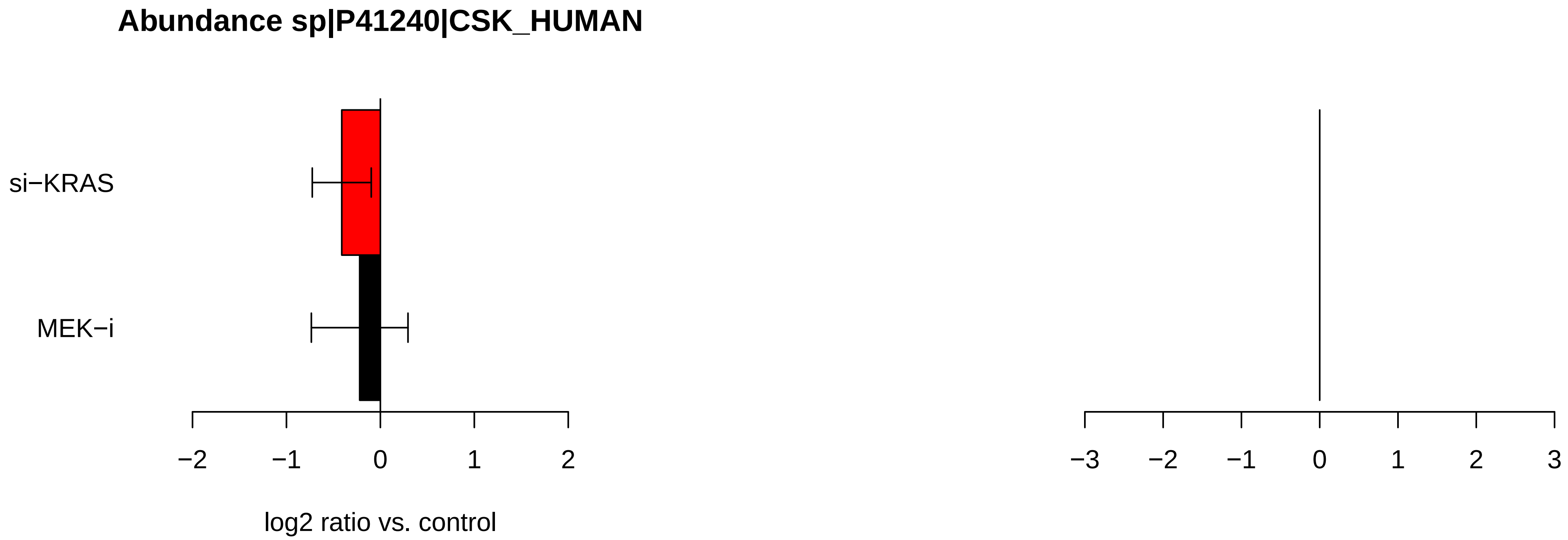

Abundance sp|P41240|CSK_HUMAN
si−KRAS
MEK−i
−2
−1
0
1
2
−3
−2
−1
0
1
2
3
log2 ratio vs. control

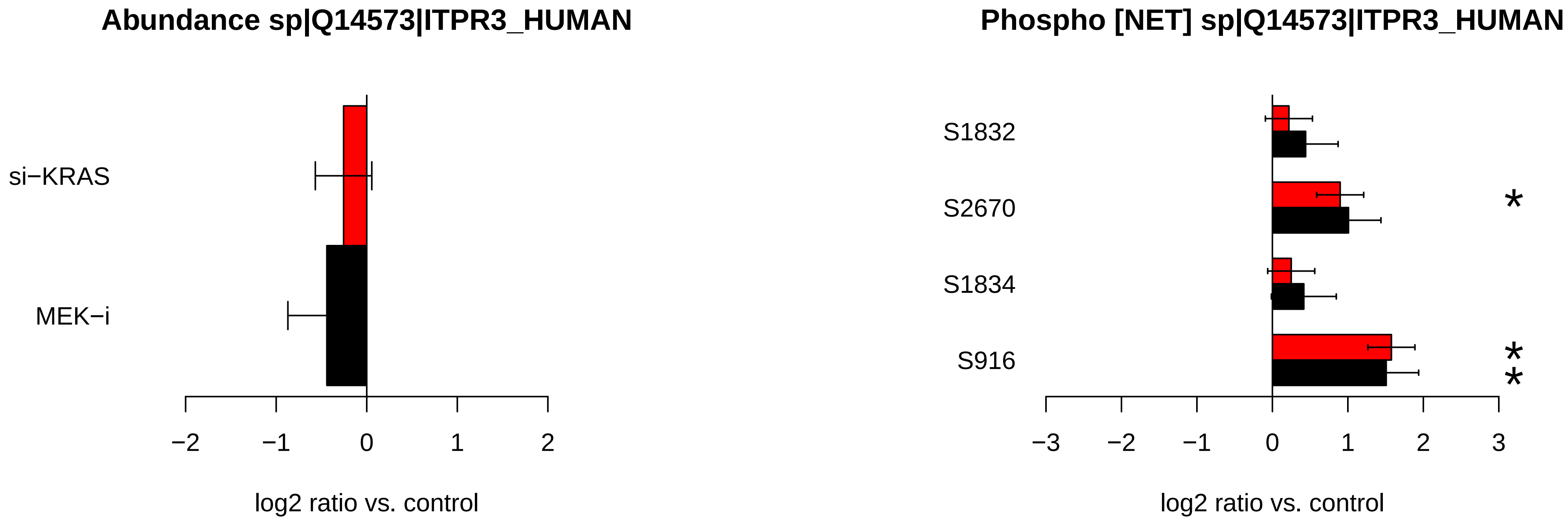
Abundance sp|Q14573|ITPR3_HUMAN
si−KRAS
MEK−i
−2
−1
0
1
2
log2 ratio vs. control
Phospho [NET] sp|Q14573|ITPR3_HUMAN
S1832
S2670
S1834
S916
*
*
*
−3
−2
−1
0
1
2
3
log2 ratio vs. control

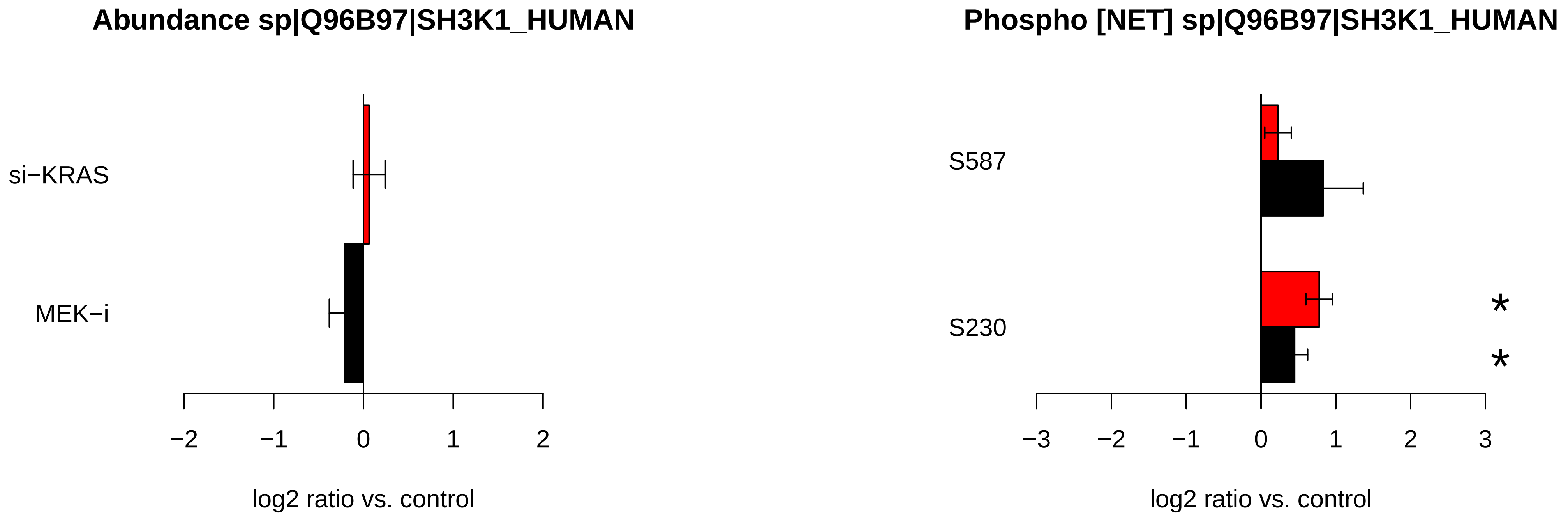
Abundance sp|Q96B97|SH3K1_HUMAN
si−KRAS
MEK−i
−2
−1
0
1
2
log2 ratio vs. control
Phospho [NET] sp|Q96B97|SH3K1_HUMAN
S587
S230
*
*
−3
−2
−1
0
1
2
3
log2 ratio vs. control

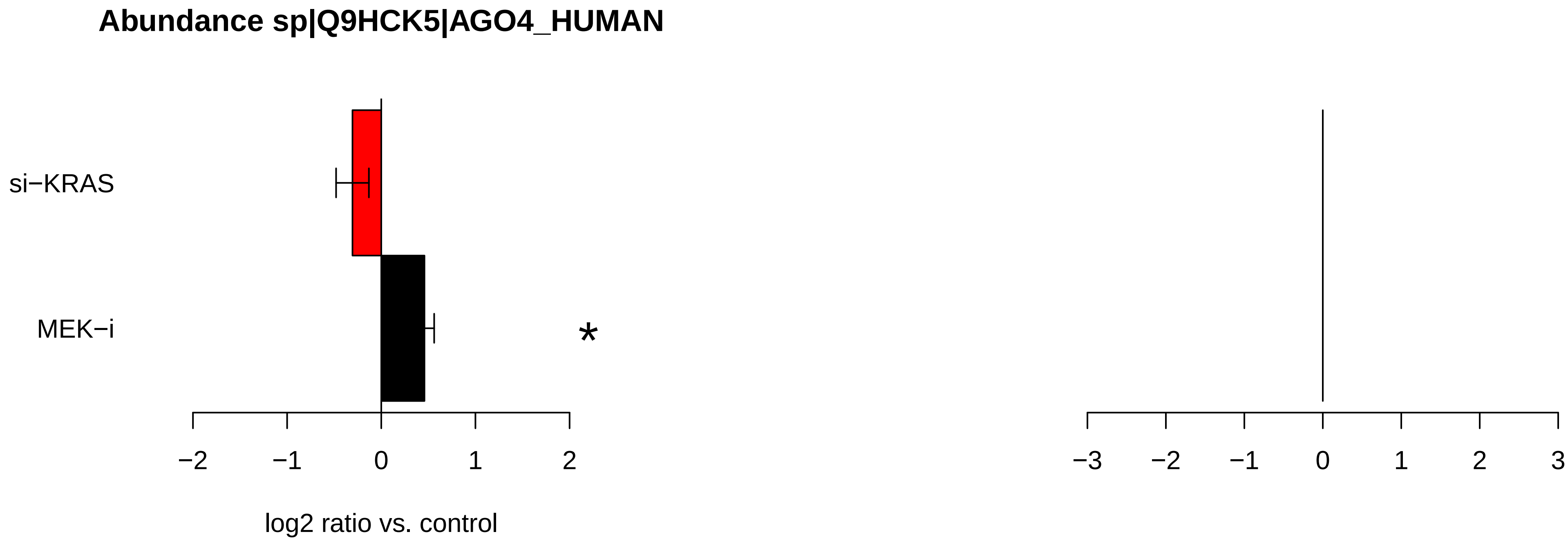

Abundance sp|Q9HCK5|AGO4_HUMAN
si−KRAS
MEK−i
*
−2
−1
0
1
2
log2 ratio vs. control
−3
−2
−1
0
1
2
3

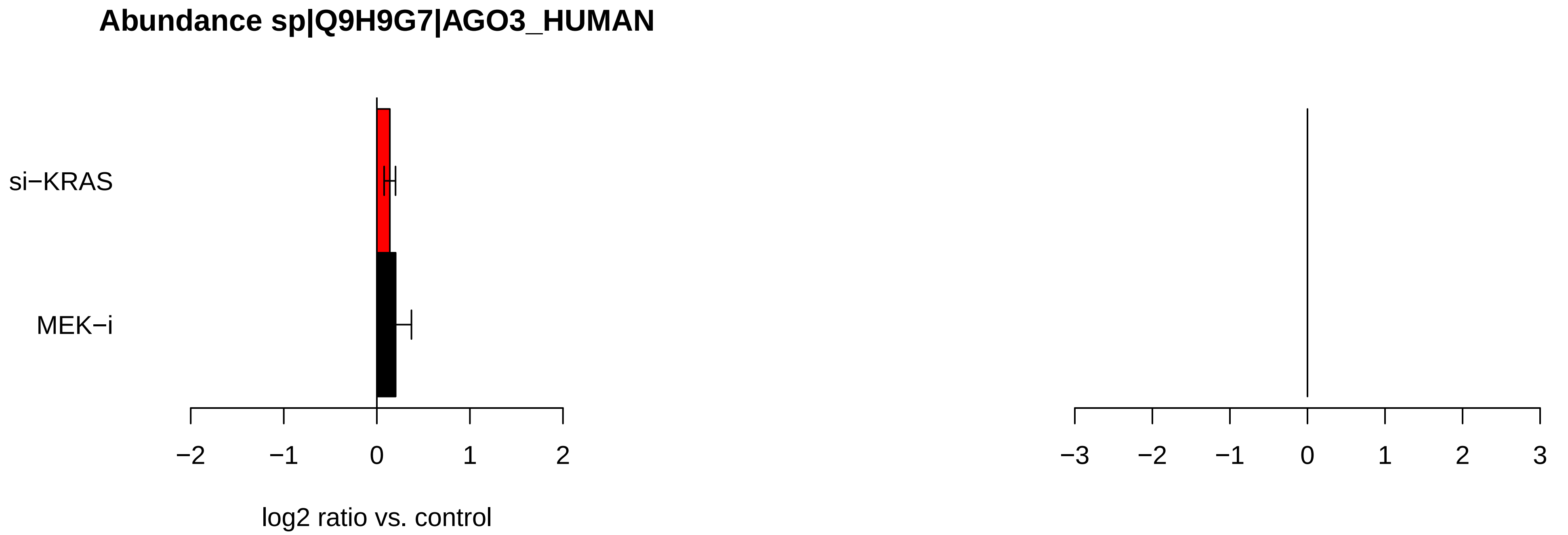
Abundance sp|Q9H9G7|AGO3_HUMAN
si−KRAS
MEK−i
−2
−1
0
1
2
log2 ratio vs. control
−3
−2
−1
0
1
2
3

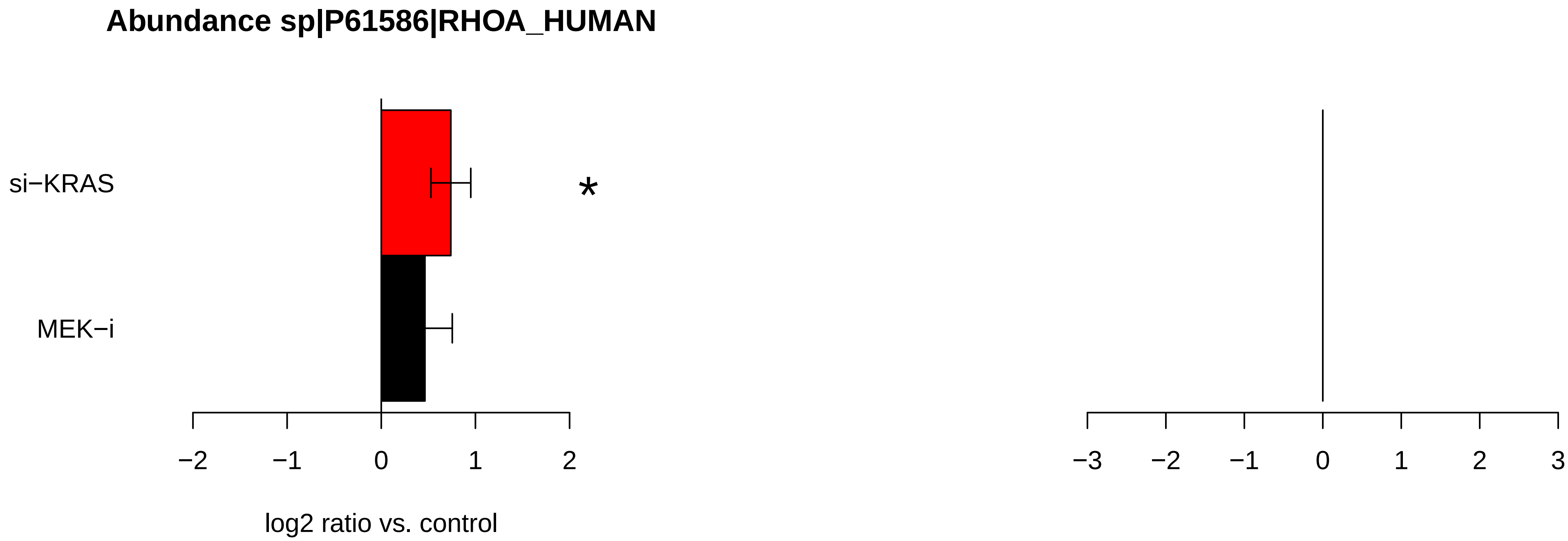
Abundance sp|P61586|RHOA_HUMAN
si−KRAS
MEK−i
*
−2
−1
0
1
2
−3
−2
−1
0
1
2
3
log2 ratio vs. control

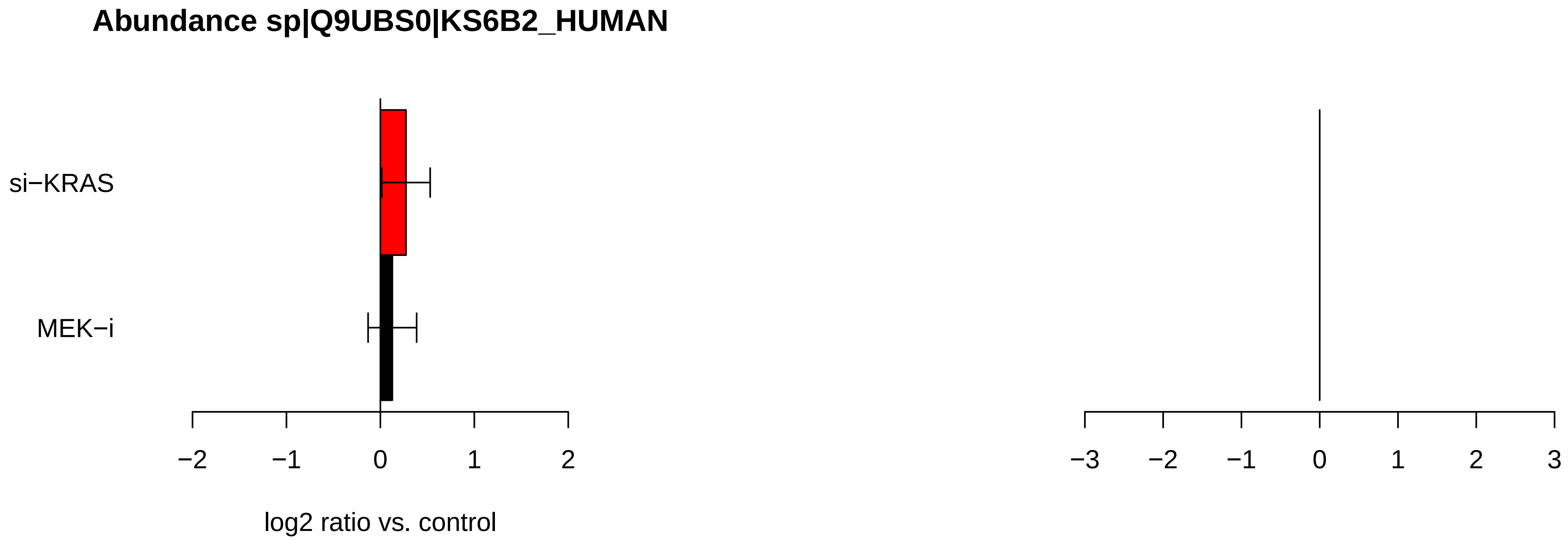

Abundance sp|Q9UBS0|KS6B2_HUMAN
si−KRAS
MEK−i
−2
−1
0
1
2
log2 ratio vs. control
−3
−2
−1
0
1
2
3

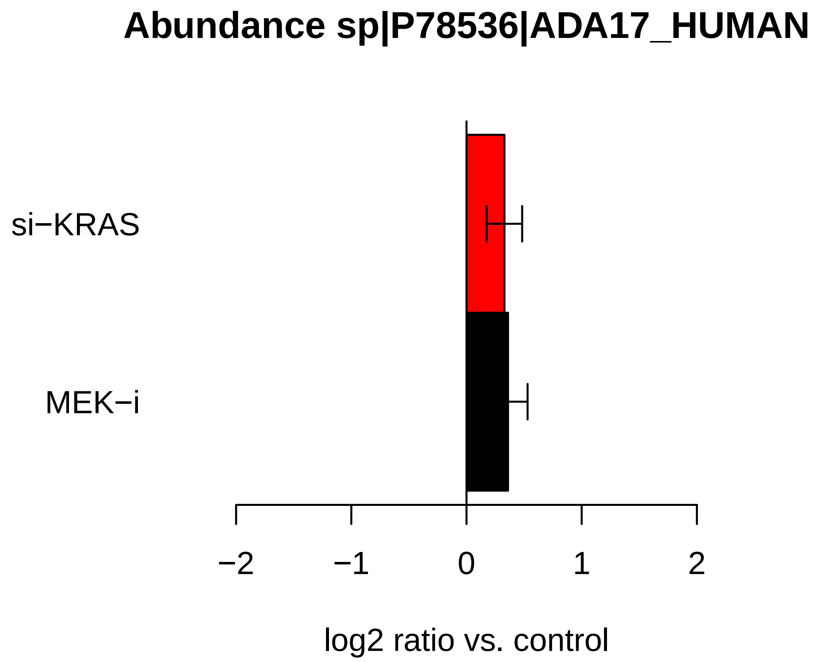

Abundance sp|P78536|ADA17_HUMAN
si−KRAS
MEK−i
−2
−1
0
1
2
log2 ratio vs. control


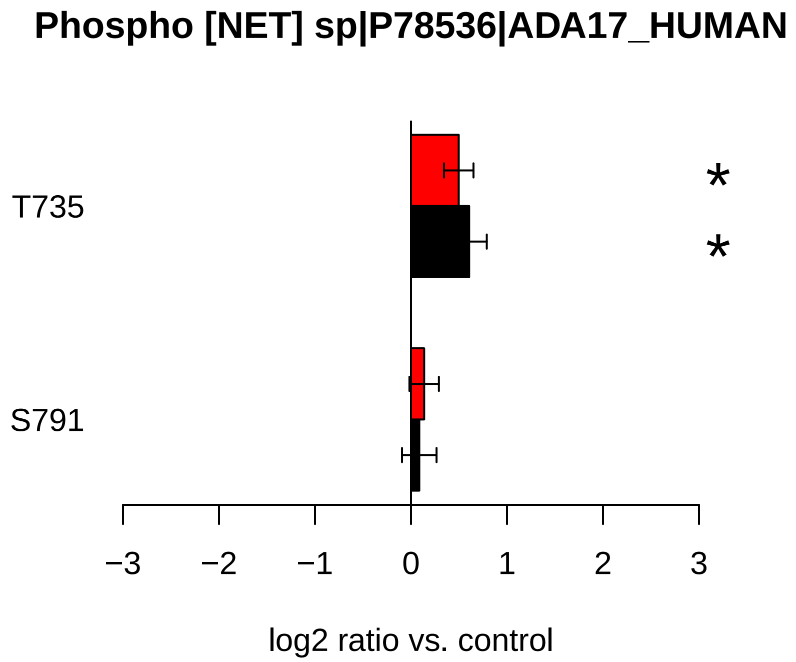

Phospho [NET] sp|P78536|ADA17_HUMAN
T735
S791
*
*
−3
−2
−1
0
1
2
3
log2 ratio vs. control

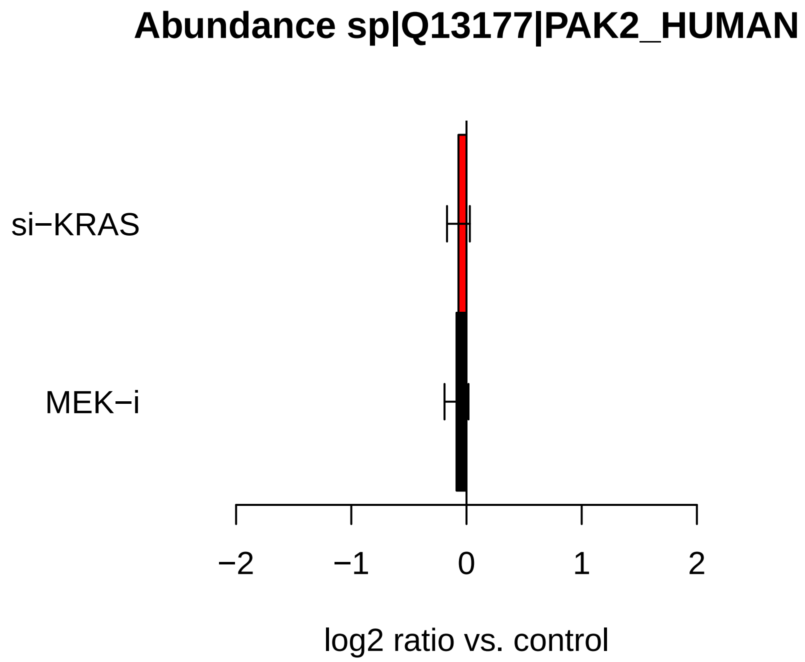
Abundance sp|Q13177|PAK2_HUMAN
si−KRAS
MEK−i
−2
−1
0
1
2
log2 ratio vs. control

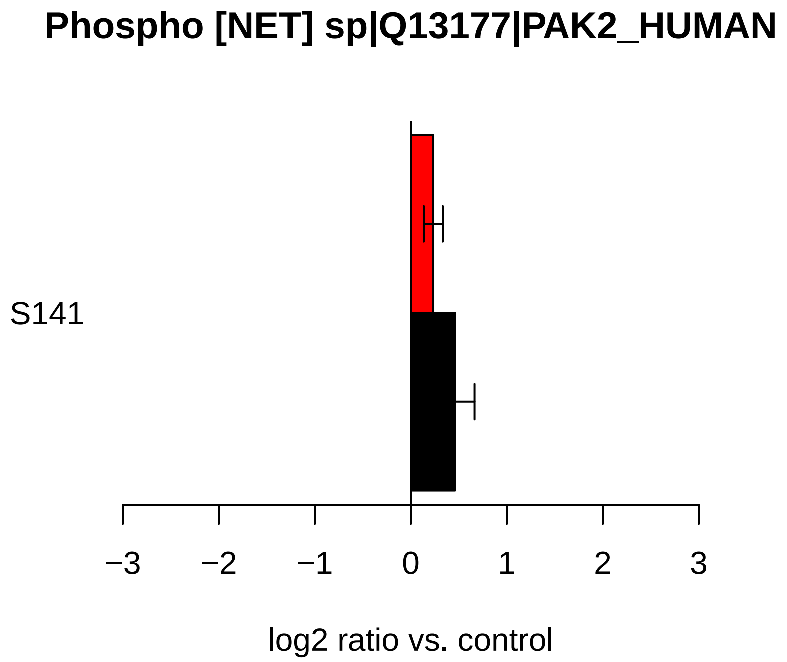
Phospho [NET] sp|Q13177|PAK2_HUMAN
S141
−3
−2
−1
0
1
2
3
log2 ratio vs. control

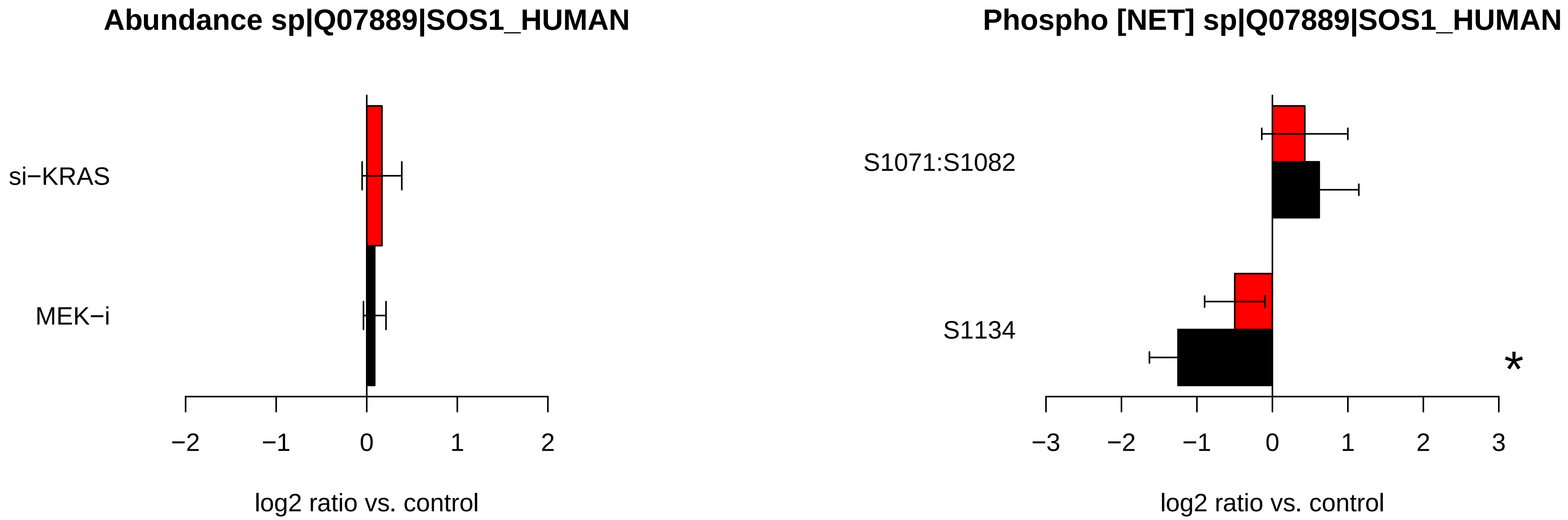

Abundance sp|Q07889|SOS1_HUMAN
si−KRAS
MEK−i
−2
−1
0
1
2
log2 ratio vs. control
Phospho [NET] sp|Q07889|SOS1_HUMAN
S1071:S1082
S1134
*
−3
−2
−1
0
1
2
3
log2 ratio vs. control

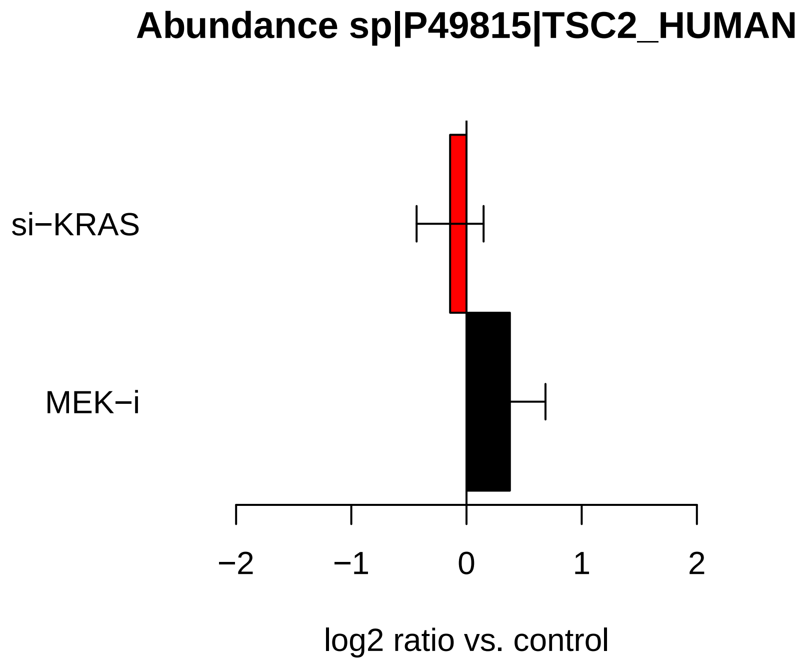
Abundance sp|P49815|TSC2_HUMAN
si−KRAS
MEK−i
−2
−1
0
1
2
log2 ratio vs. control

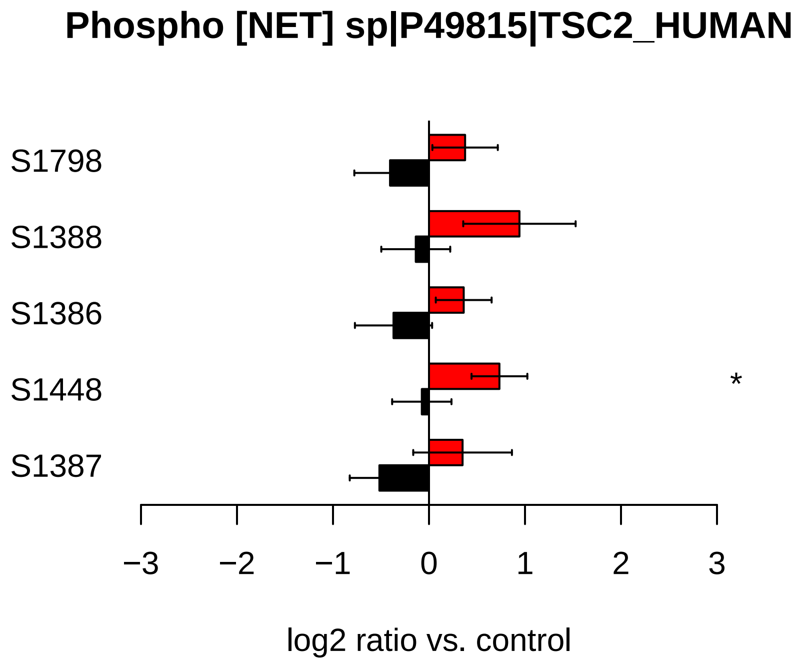
Phospho [NET] sp|P49815|TSC2_HUMAN
S1798
S1388
S1386
S1448
S1387
*
−3
−2
−1
0
1
2
3
log2 ratio vs. control

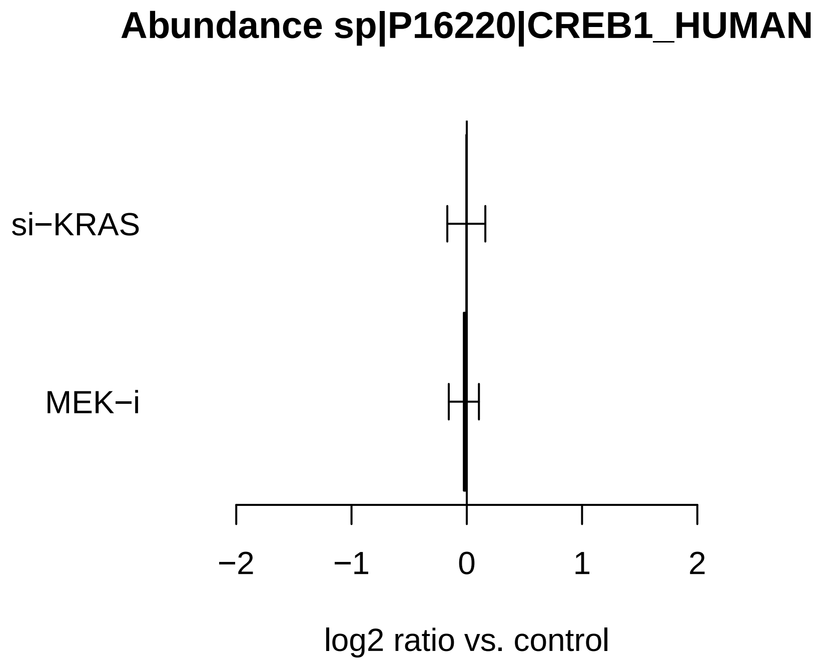
Abundance sp|P16220|CREB1_HUMAN
si−KRAS
MEK−i
−2
−1
0
1
2
log2 ratio vs. control

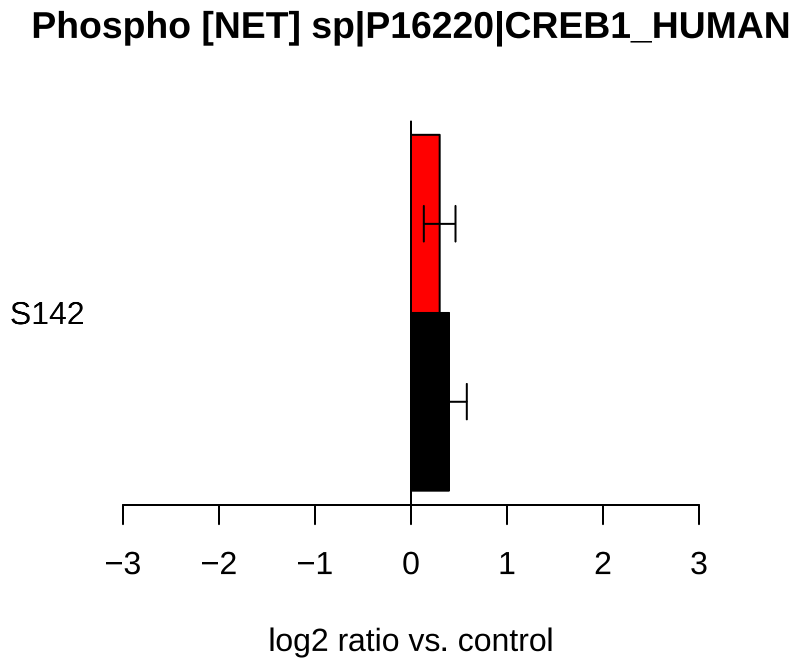
Phospho [NET] sp|P16220|CREB1_HUMAN
S142
−3
−2
−1
0
1
2
3
log2 ratio vs. control

### Abundance sp|P17612|KAPCA_HUMAN

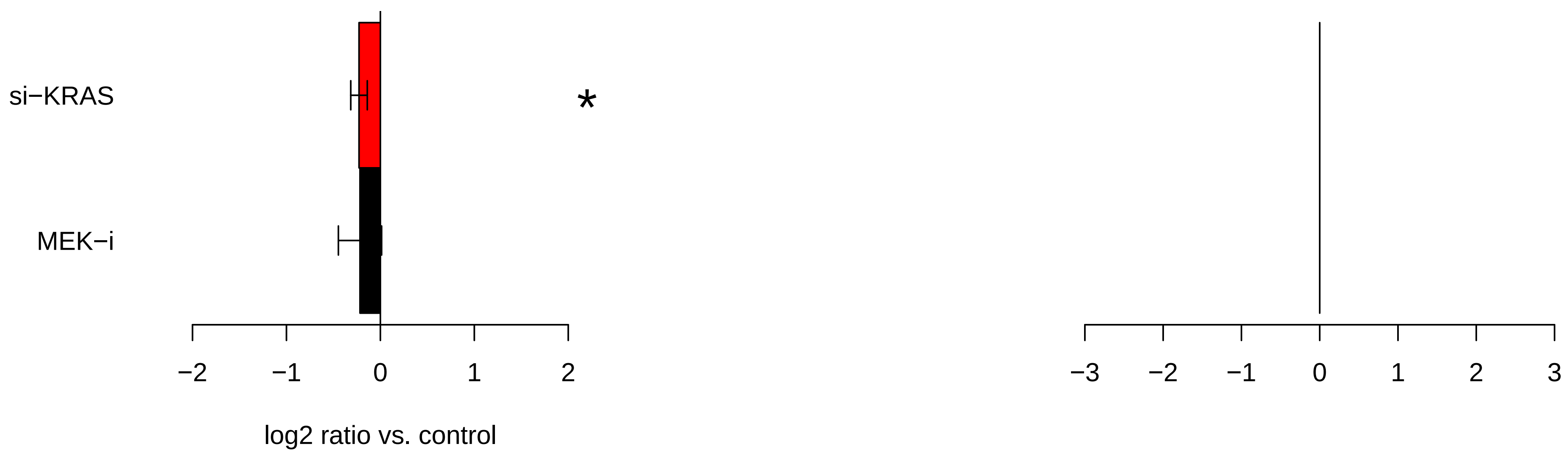

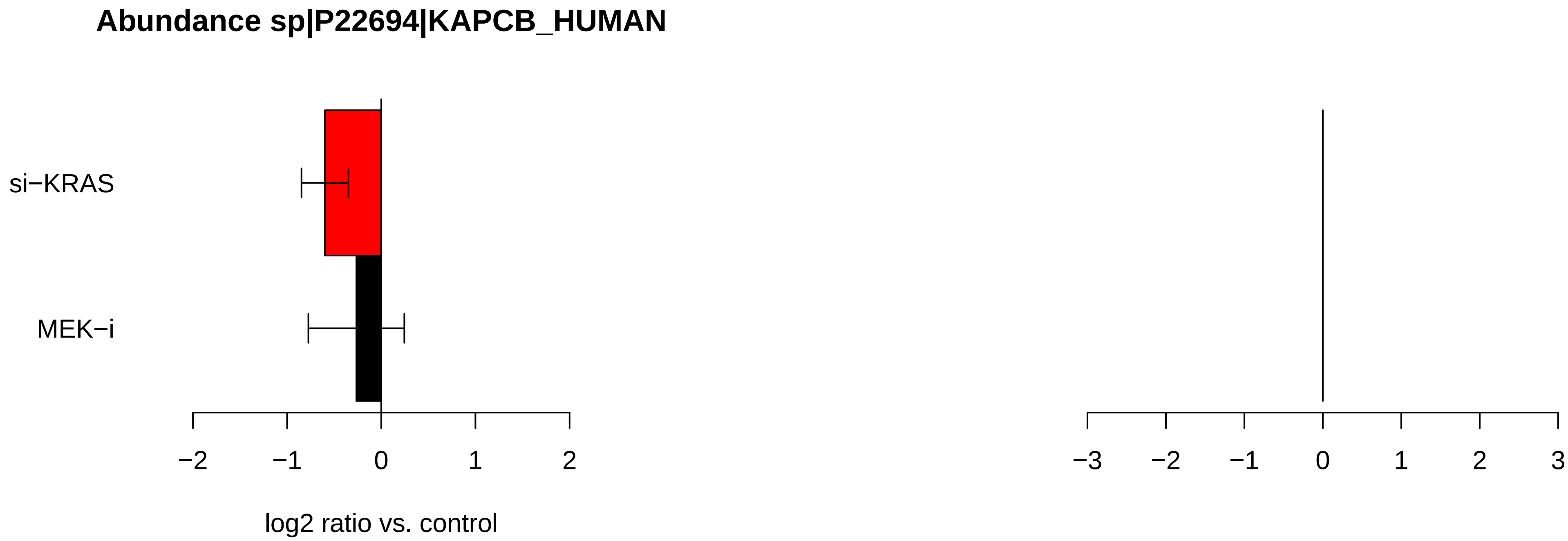

Abundance sp|P22694|KAPCB_HUMAN
si−KRAS
MEK−i
−2
−1
0
1
2
log2 ratio vs. control
−3
−2
−1
0
1
2
3

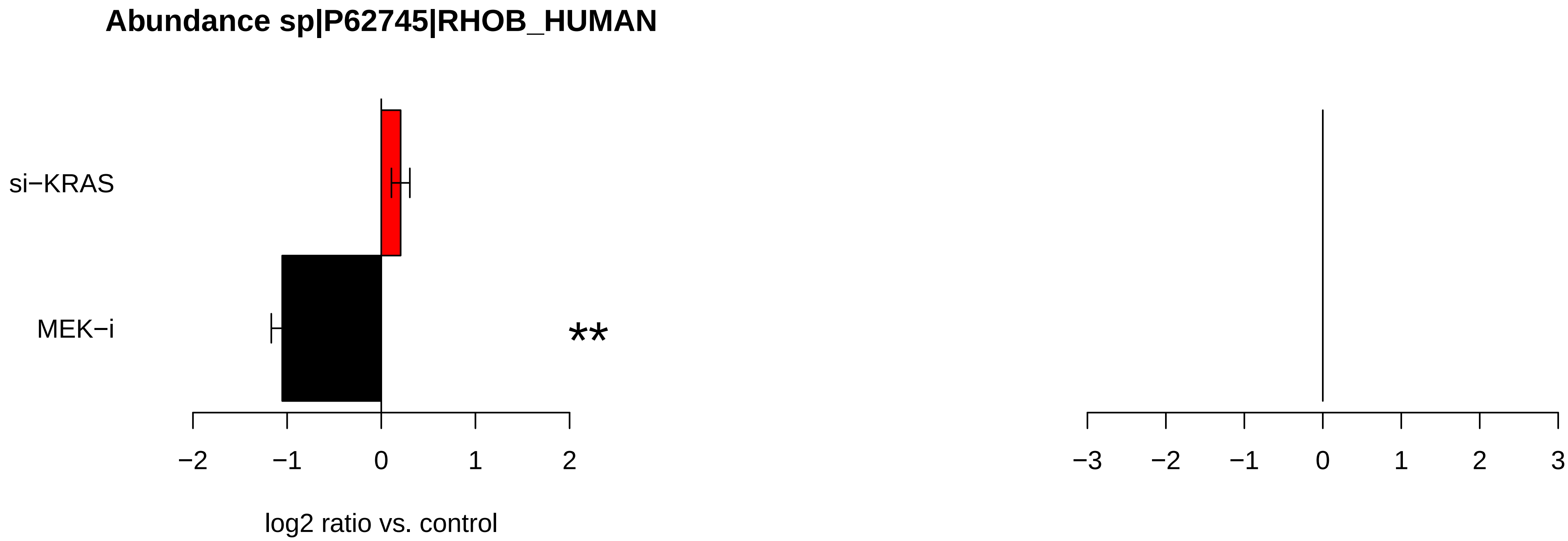
Abundance sp|P62745|RHOB_HUMAN
si−KRAS
MEK−i
**
−2
−1
0
1
2
log2 ratio vs. control
−3
−2
−1
0
1
2
3

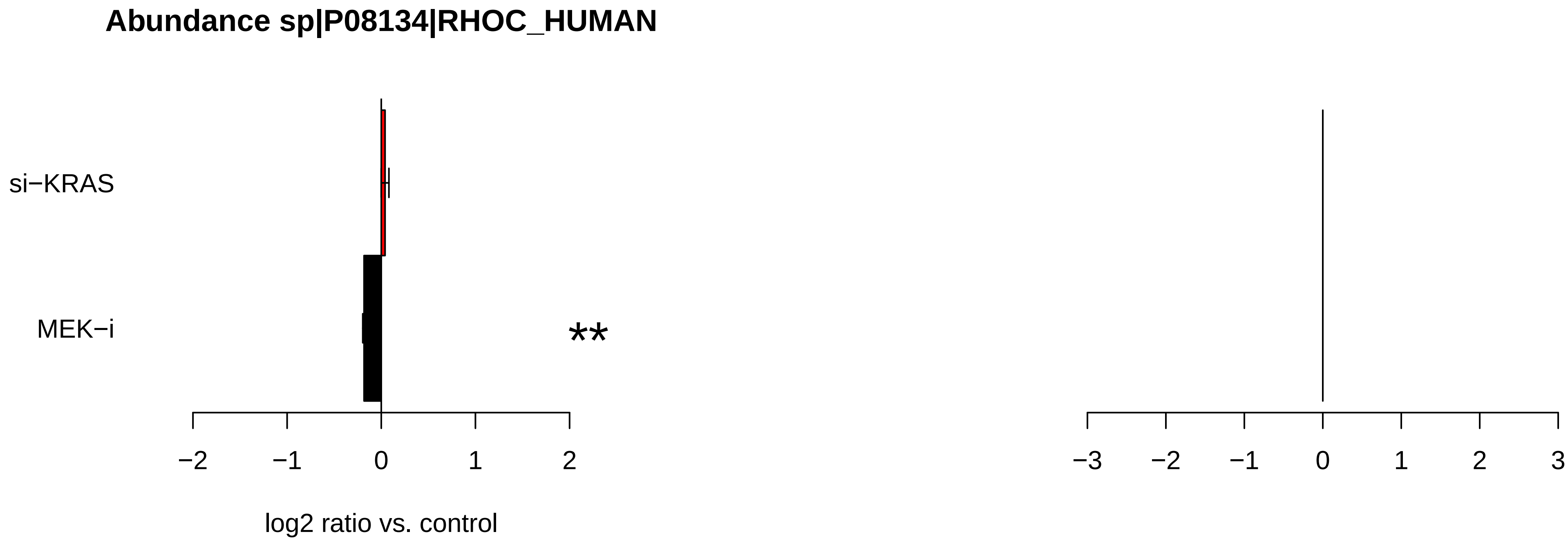
Abundance sp|P08134|RHOC_HUMAN
si−KRAS
MEK−i
**
−2
−1
0
1
2
log2 ratio vs. control
−3
−2
−1
0
1
2
3

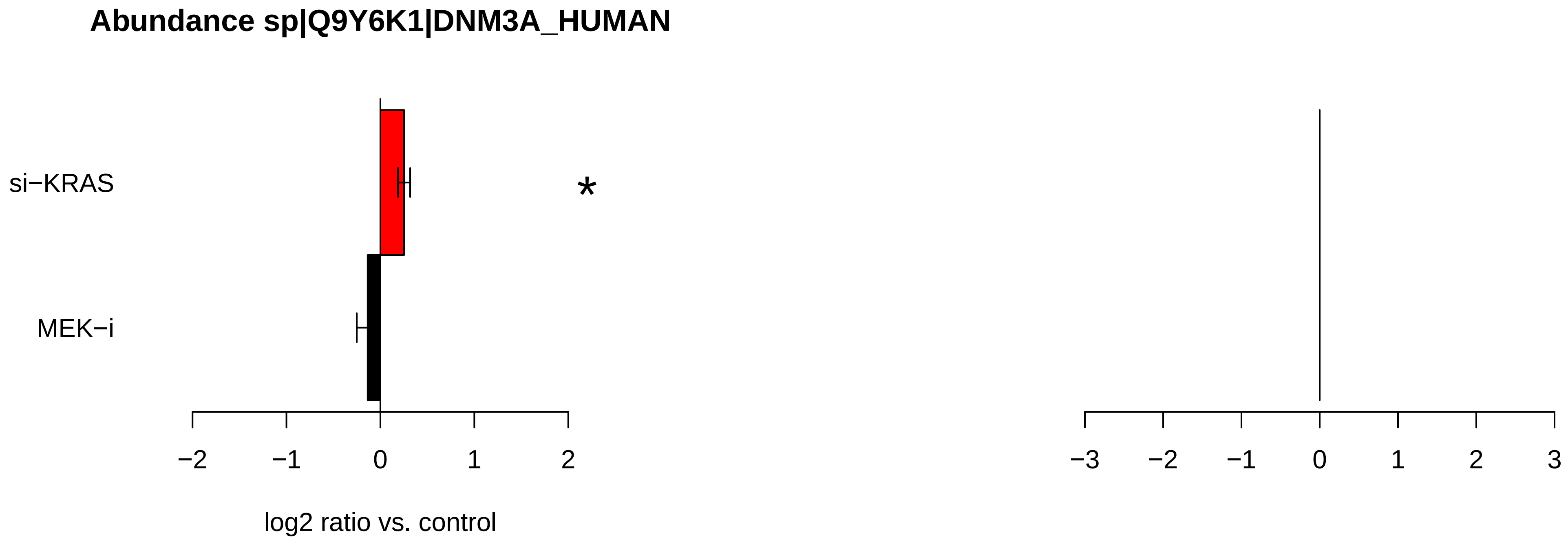
Abundance sp|Q9Y6K1|DNM3A_HUMAN
si−KRAS
MEK−i
*
−2
−1
0
1
2
log2 ratio vs. control
−3
−2
−1
0
1
2
3

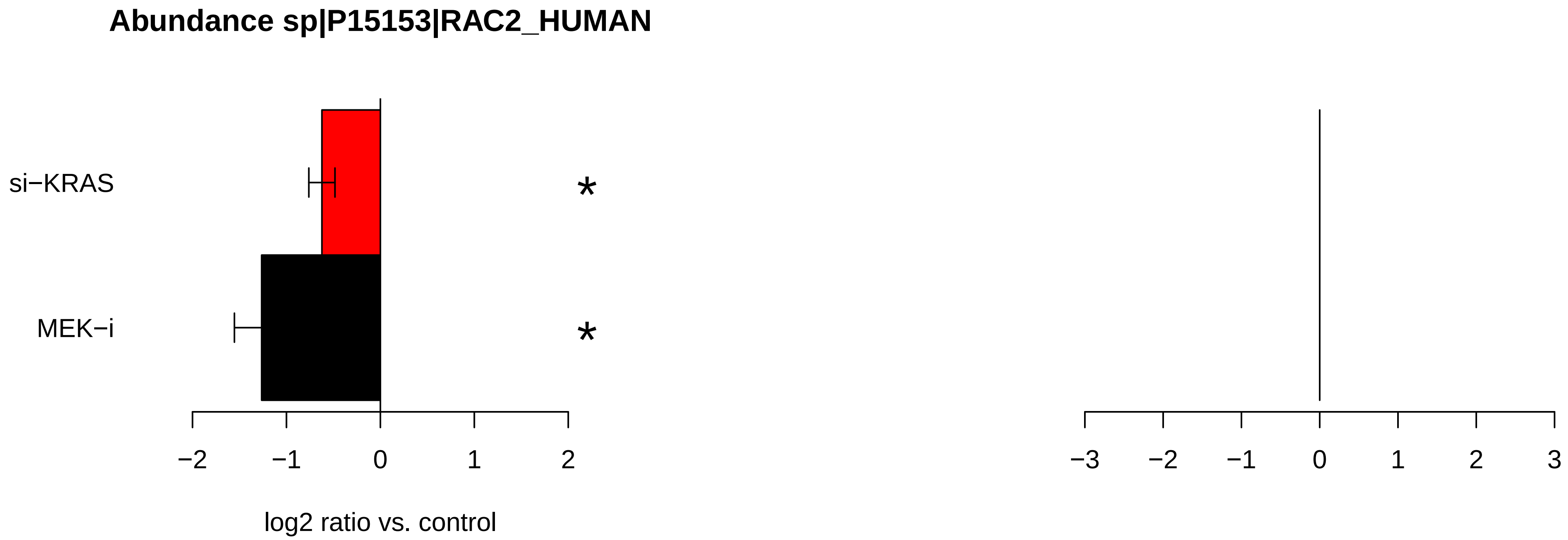
Abundance sp|P15153|RAC2_HUMAN
si−KRAS
MEK−i
*
*
−2
−1
0
1
2
log2 ratio vs. control
−3
−2
−1
0
1
2
3

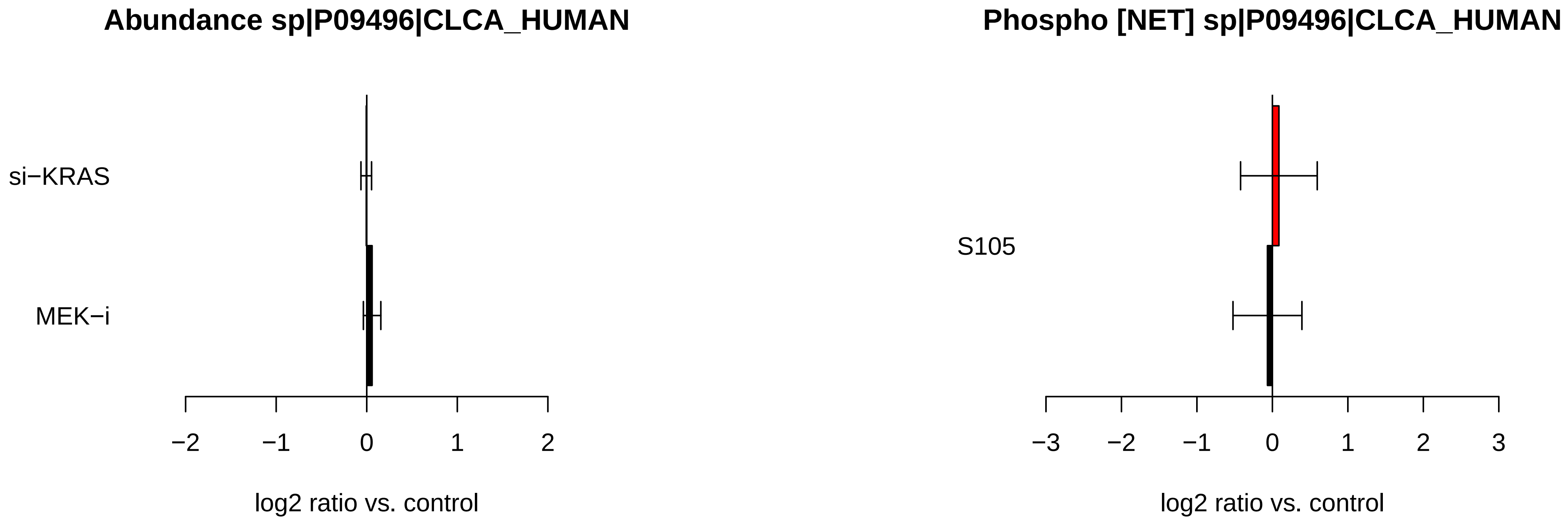

Abundance sp|P09496|CLCA_HUMAN
si−KRAS
MEK−i
−2
−1
0
1
2
log2 ratio vs. control
Phospho [NET] sp|P09496|CLCA_HUMAN
S105
−3
−2
−1
0
1
2
3
log2 ratio vs. control

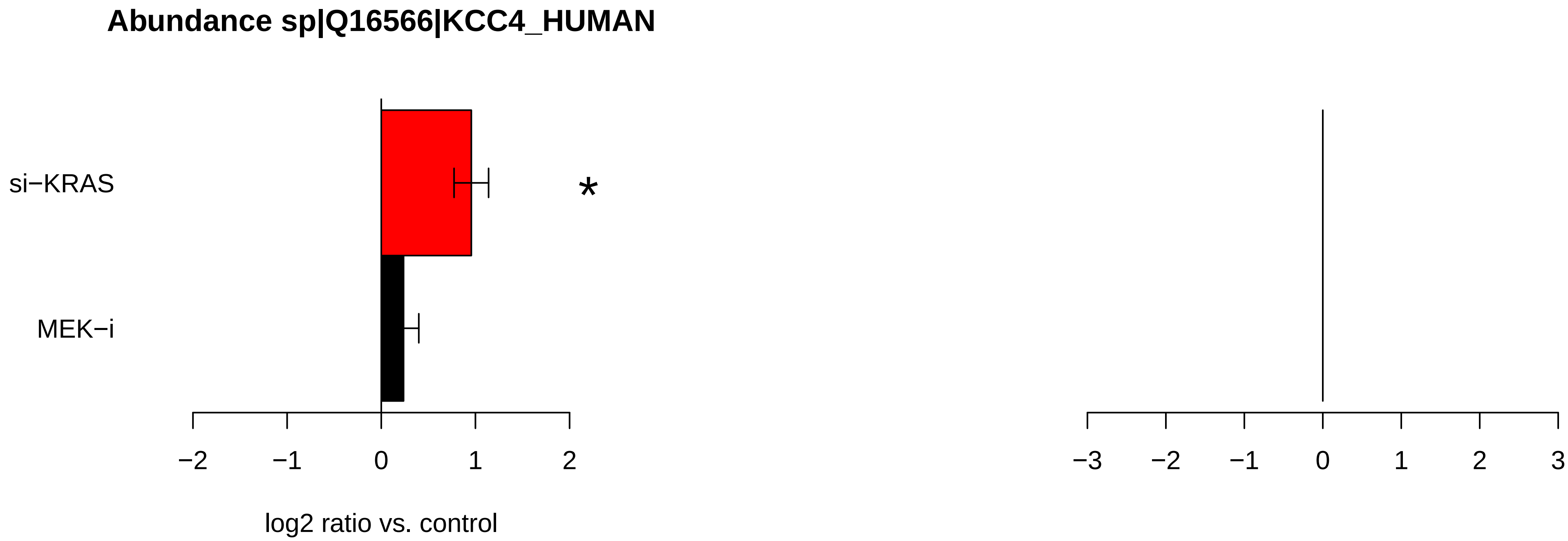
Abundance sp|Q16566|KCC4_HUMAN
si−KRAS
MEK−i
*
−2
−1
0
1
2
log2 ratio vs. control
−3
−2
−1
0
1
2
3

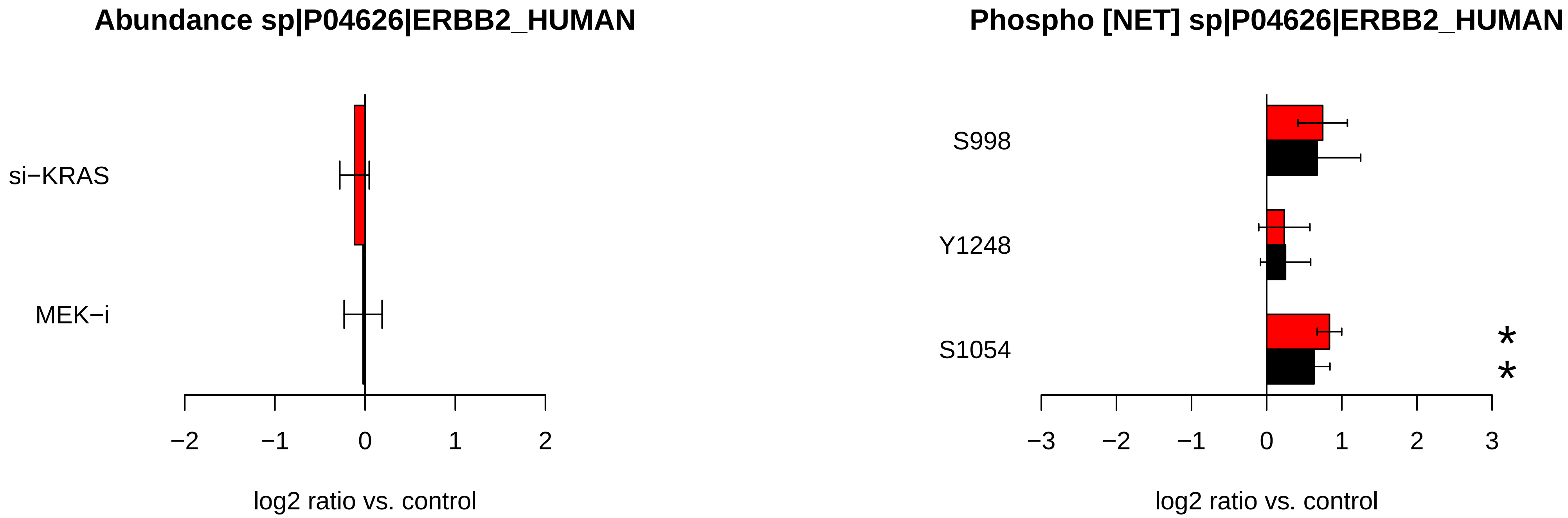
Abundance sp|P04626|ERBB2_HUMAN
si−KRAS
MEK−i
−2
−1
0
1
2
log2 ratio vs. control
Phospho [NET] sp|P04626|ERBB2_HUMAN
S998
Y1248
S1054
*
*
−3
−2
−1
0
1
2
3
log2 ratio vs. control

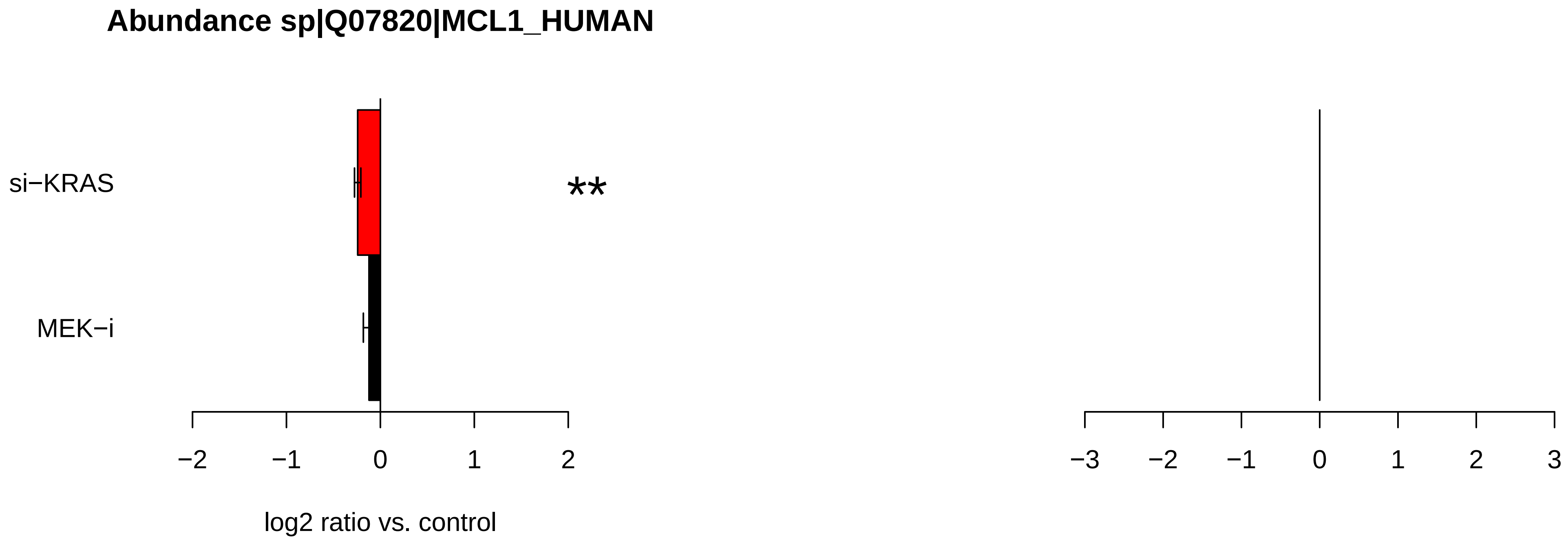

Abundance sp|Q07820|MCL1_HUMAN
si−KRAS
MEK−i
**
−2
−1
0
1
2
log2 ratio vs. control
−3
−2
−1
0
1
2
3

### Abundance sp|Q02880|TOP2B_HUMAN

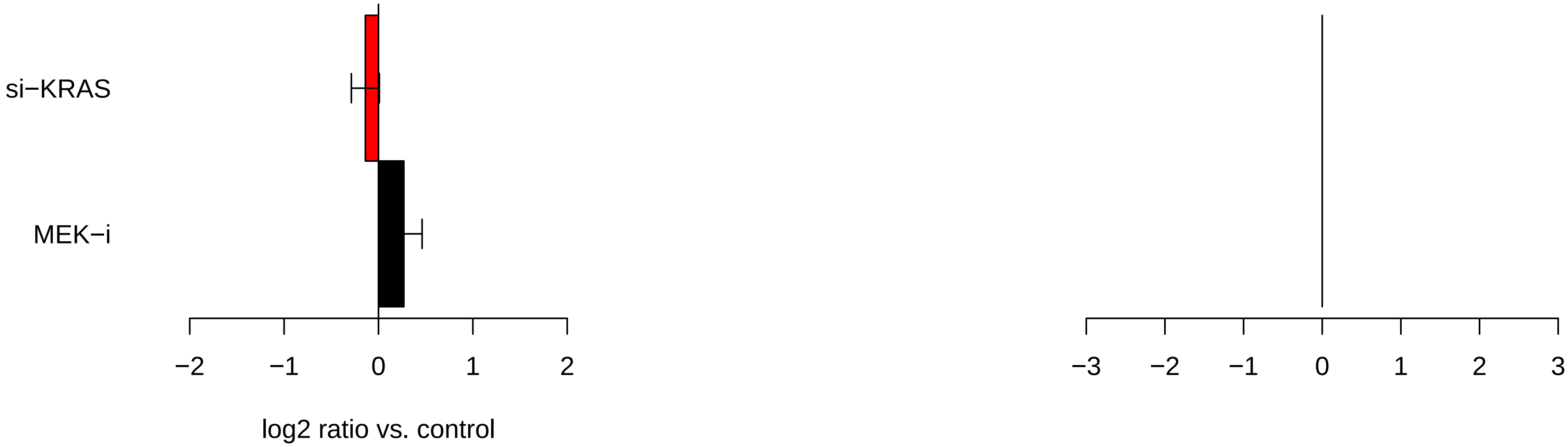

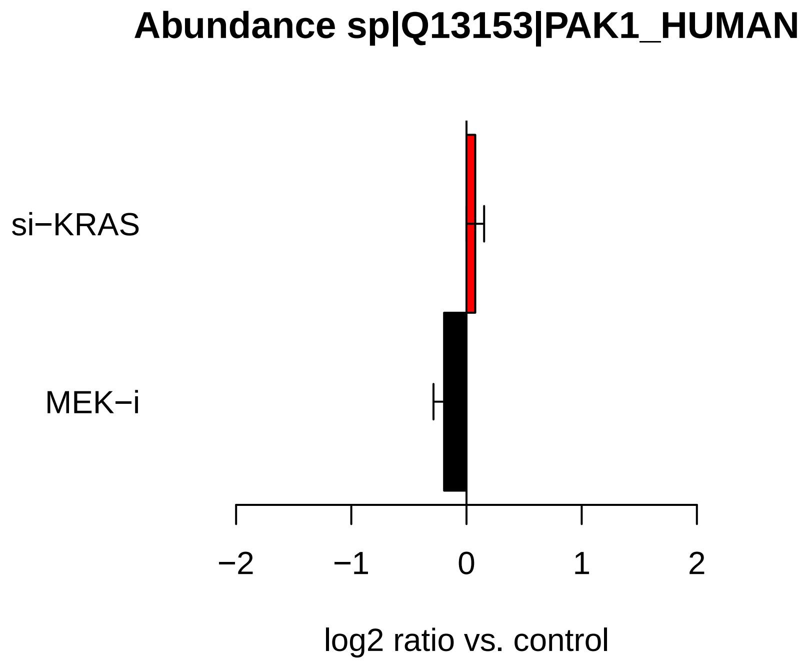

Abundance sp|Q13153|PAK1_HUMAN
si−KRAS
MEK−i
−2
−1
0
1
2
log2 ratio vs. control


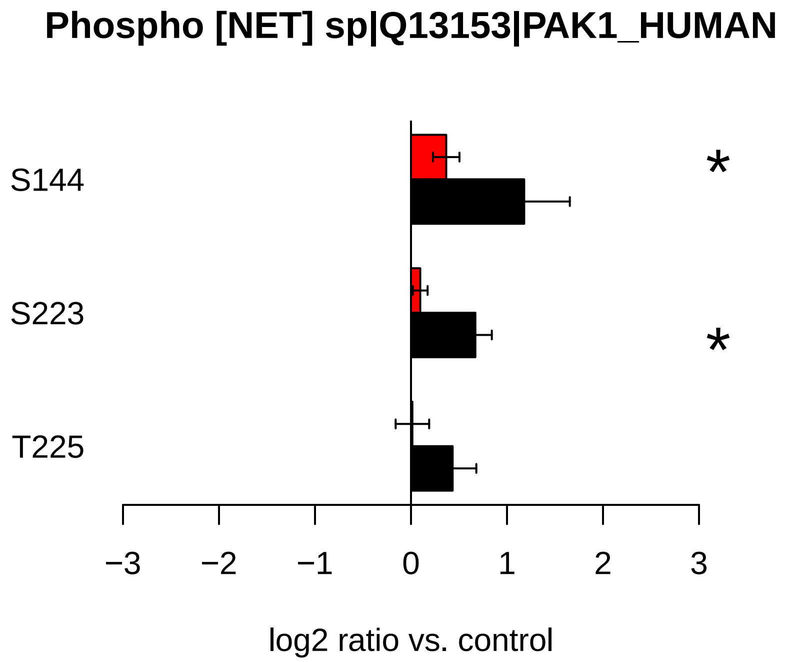

Phospho [NET] sp|Q13153|PAK1_HUMAN
S144
S223
T225
*
*
−3
−2
−1
0
1
2
3
log2 ratio vs. control

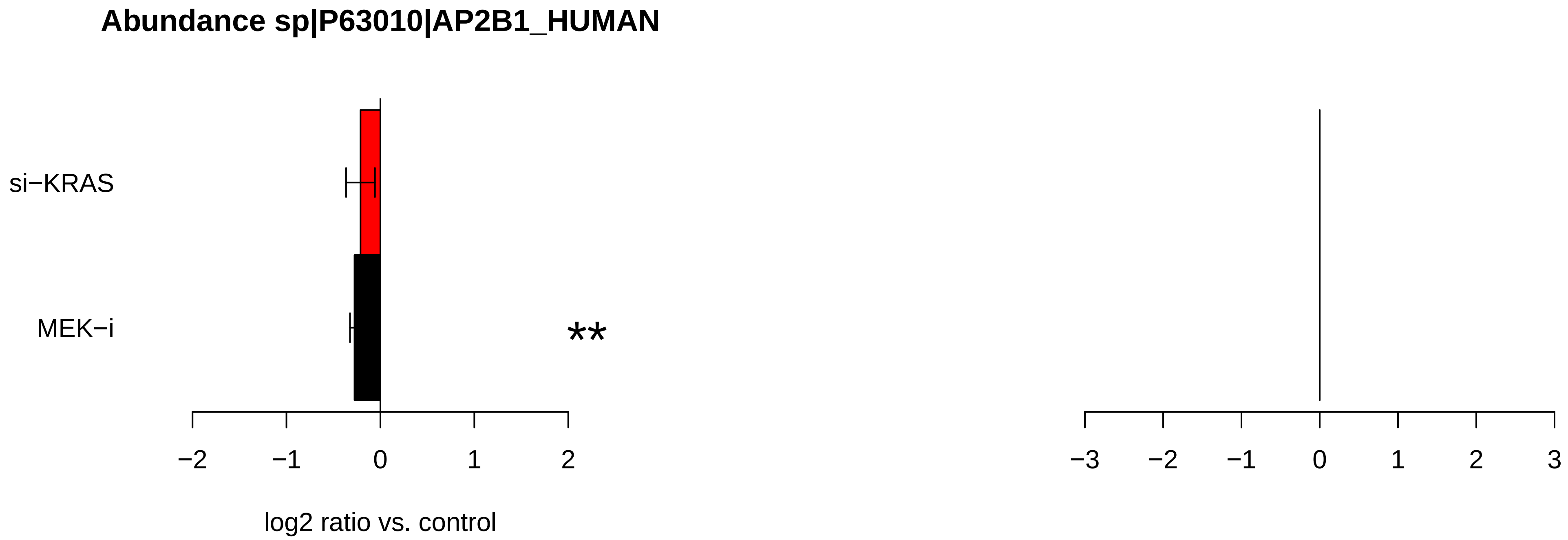
Abundance sp|P63010|AP2B1_HUMAN
si−KRAS
MEK−i
**
−2
−1
0
1
2
log2 ratio vs. control
−3
−2
−1
0
1
2
3

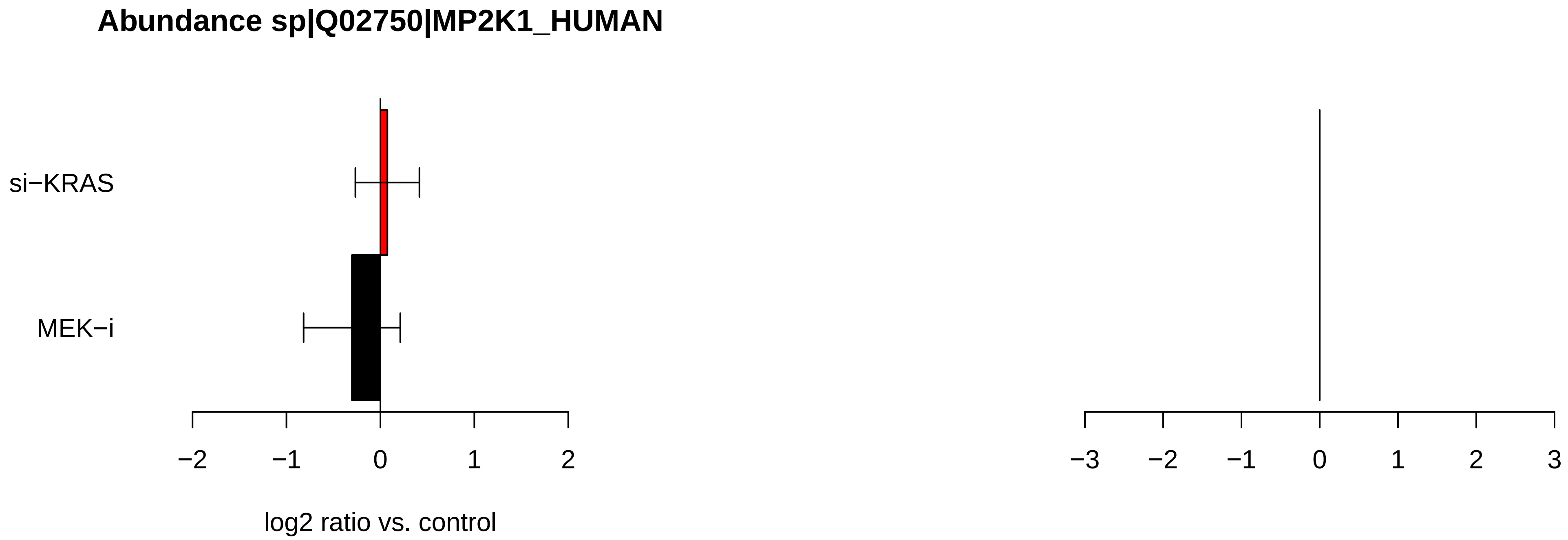

Abundance sp|Q02750|MP2K1_HUMAN
si−KRAS
MEK−i
−2
−1
0
1
2
log2 ratio vs. control
−3
−2
−1
0
1
2
3

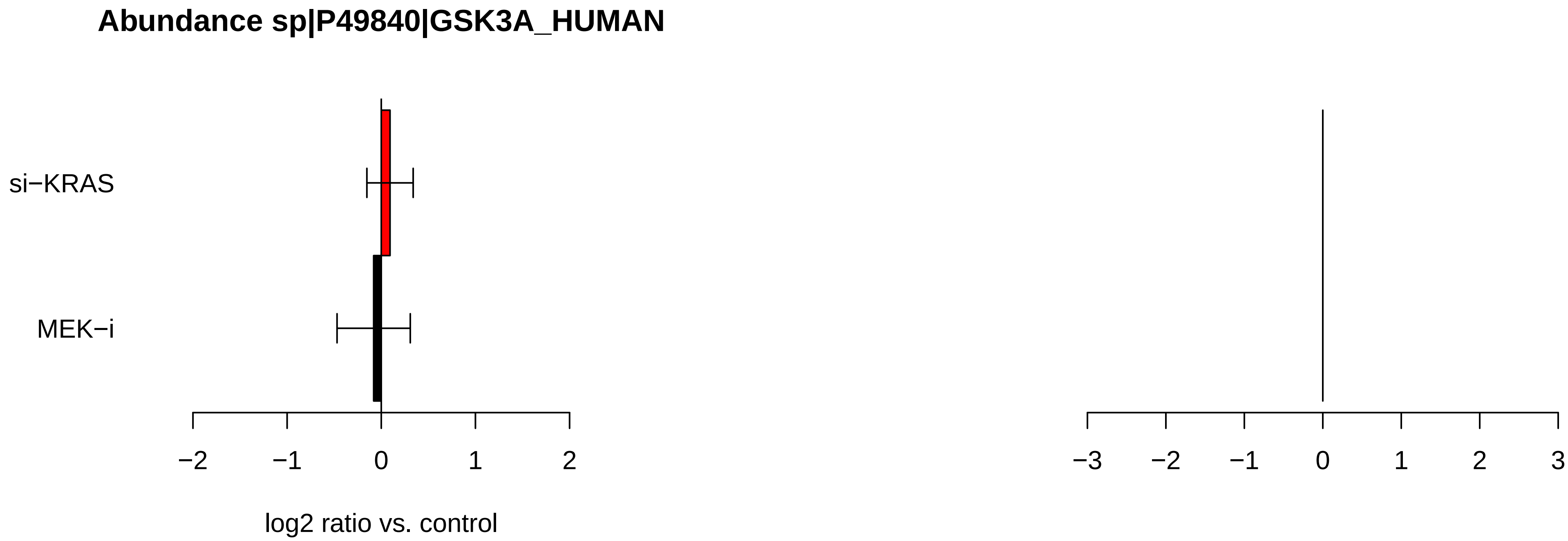
Abundance sp|P49840|GSK3A_HUMAN
si−KRAS
MEK−i
−2
−1
0
1
2
log2 ratio vs. control
−3
−2
−1
0
1
2
3

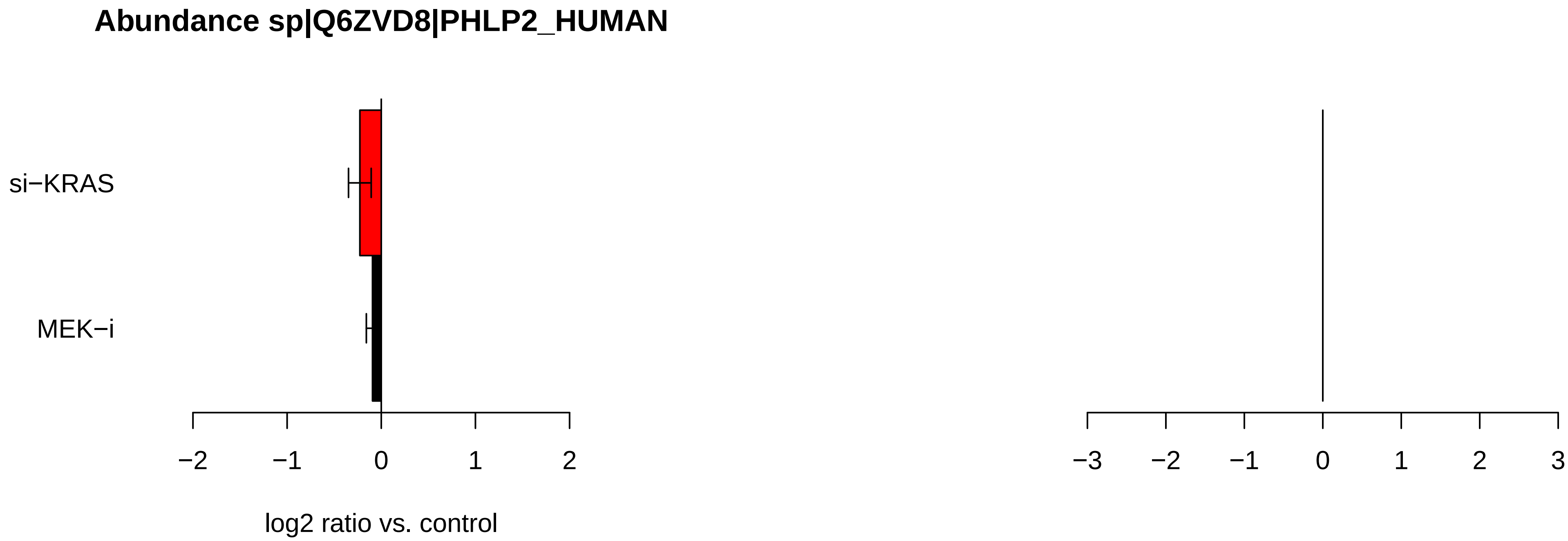

Abundance sp|Q6ZVD8|PHLP2_HUMAN
si−KRAS
MEK−i
−2
−1
0
1
2
log2 ratio vs. control
−3
−2
−1
0
1
2
3

### Abundance sp|P49841|GSK3B_HUMAN

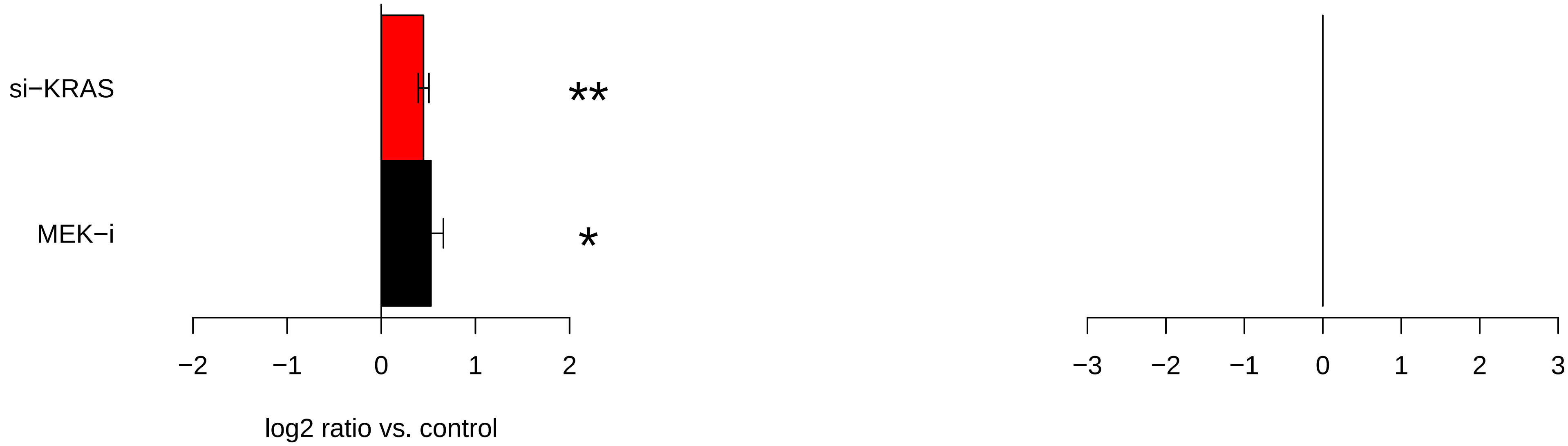

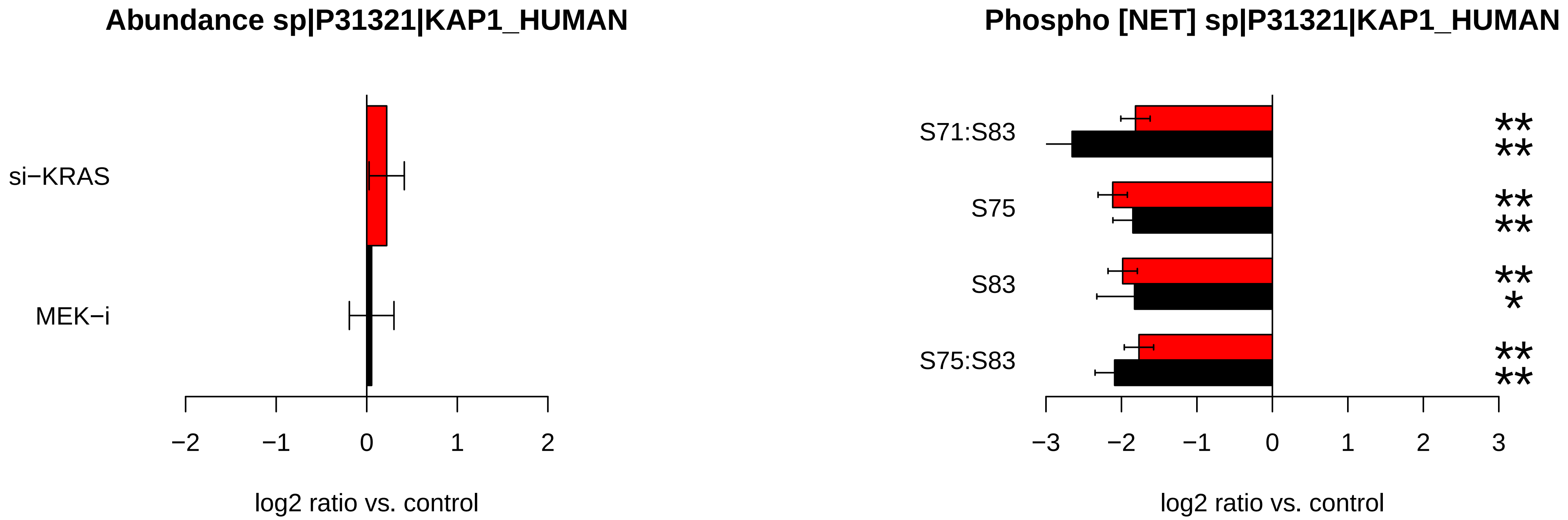
Abundance sp|P31321|KAP1_HUMAN
si−KRAS
MEK−i
−2
−1
0
1
2
log2 ratio vs. control
Phospho [NET] sp|P31321|KAP1_HUMAN
S71:S83
S75
S83
S75:S83
**
**
**
**
**
*
**
**
−3
−2
−1
0
1
2
3
log2 ratio vs. control

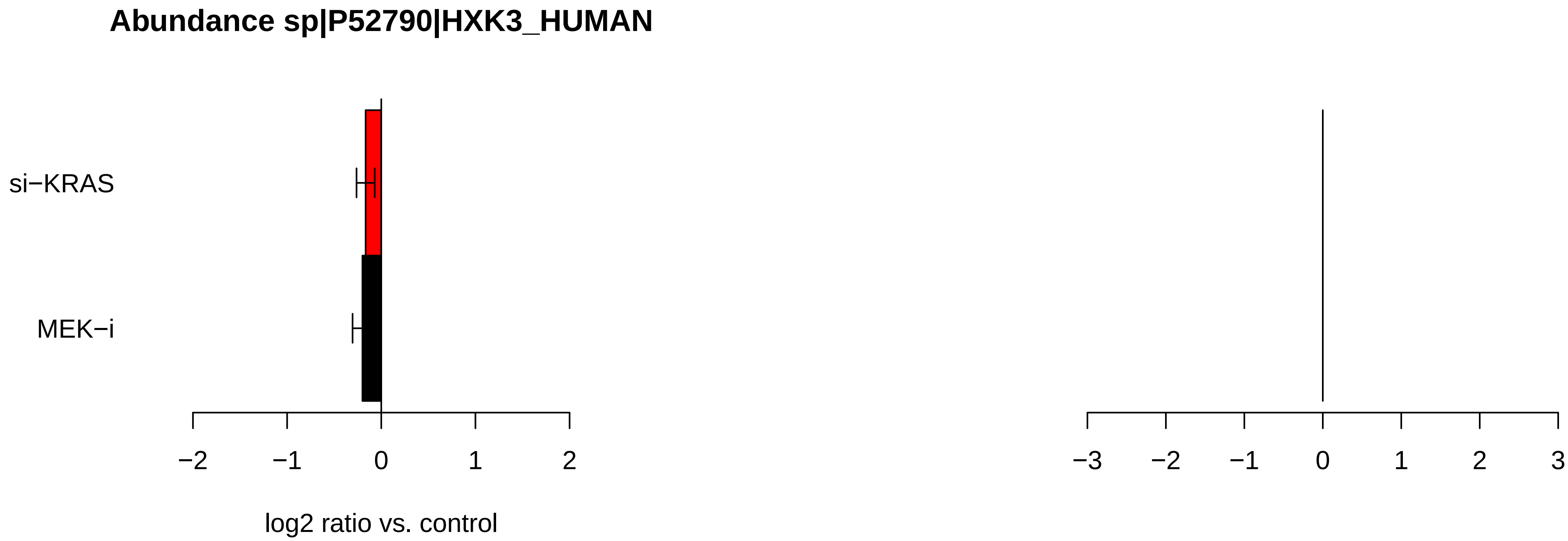

Abundance sp|P52790|HXK3_HUMAN
si−KRAS
MEK−i
−2
−1
0
1
2
log2 ratio vs. control
−3
−2
−1
0
1
2
3

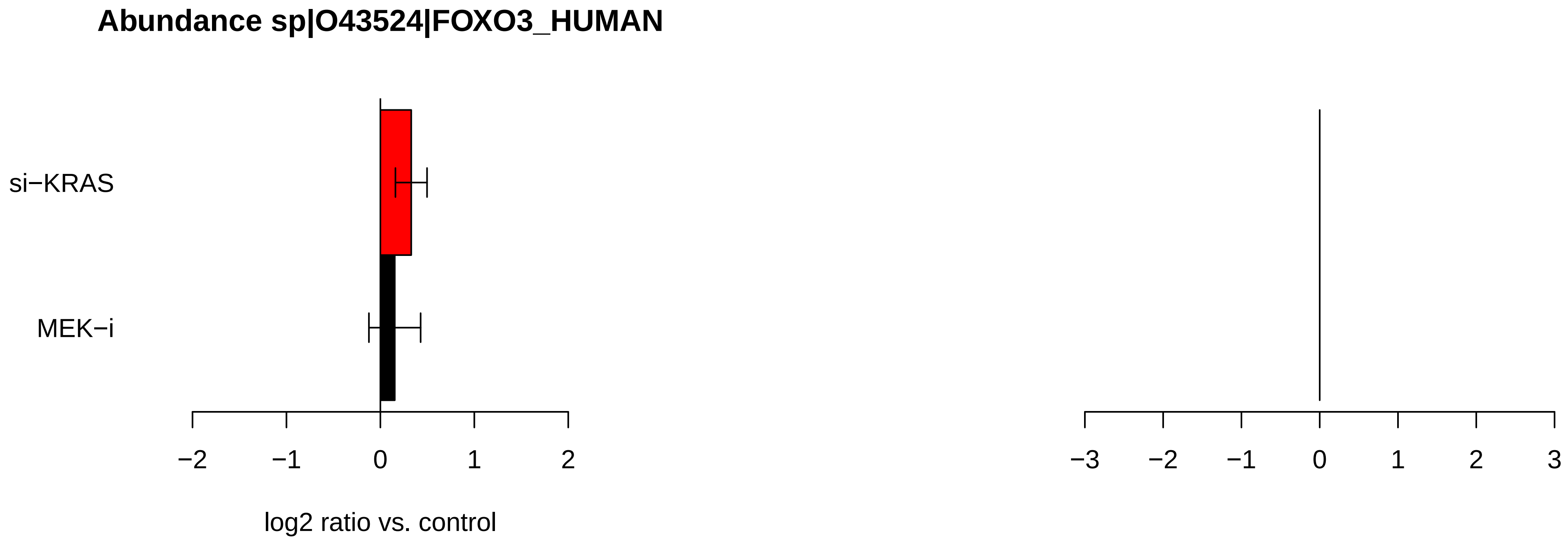
Abundance sp|O43524|FOXO3_HUMAN
si−KRAS
MEK−i
−2
−1
0
1
2
log2 ratio vs. control
−3
−2
−1
0
1
2
3

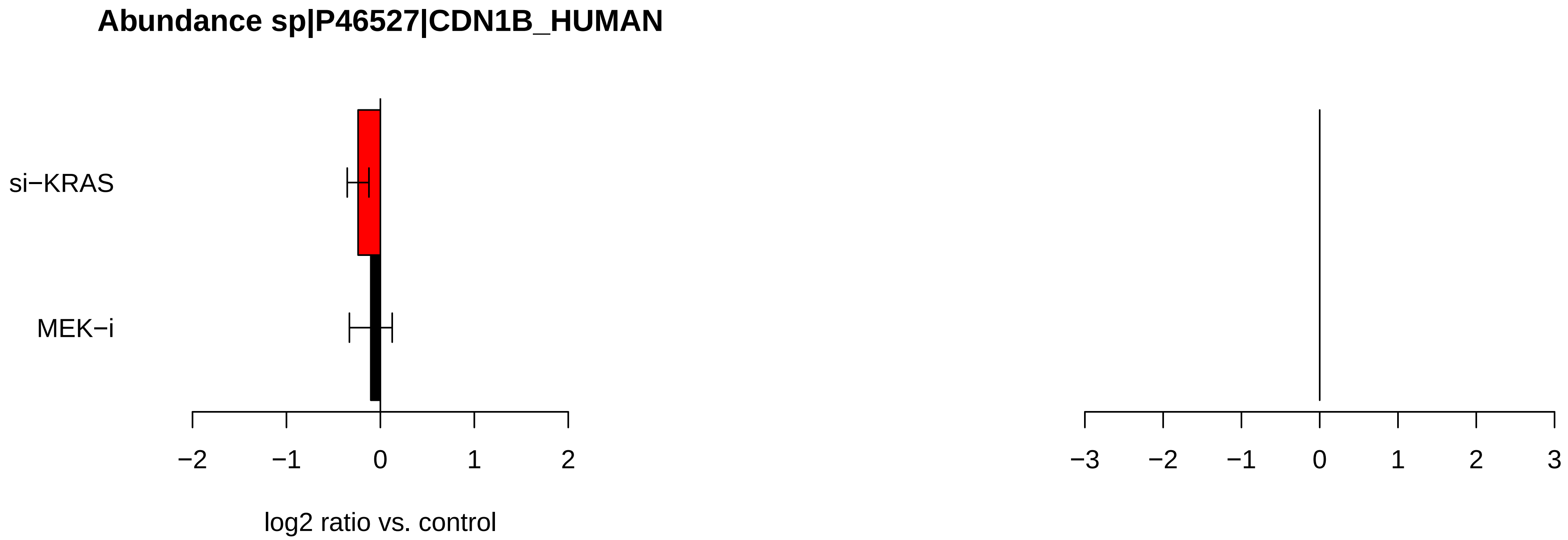

Abundance sp|P46527|CDN1B_HUMAN
si−KRAS
MEK−i
−2
−1
0
1
2
log2 ratio vs. control
−3
−2
−1
0
1
2
3

### Abundance sp|Q14155|ARHG7_HUMAN

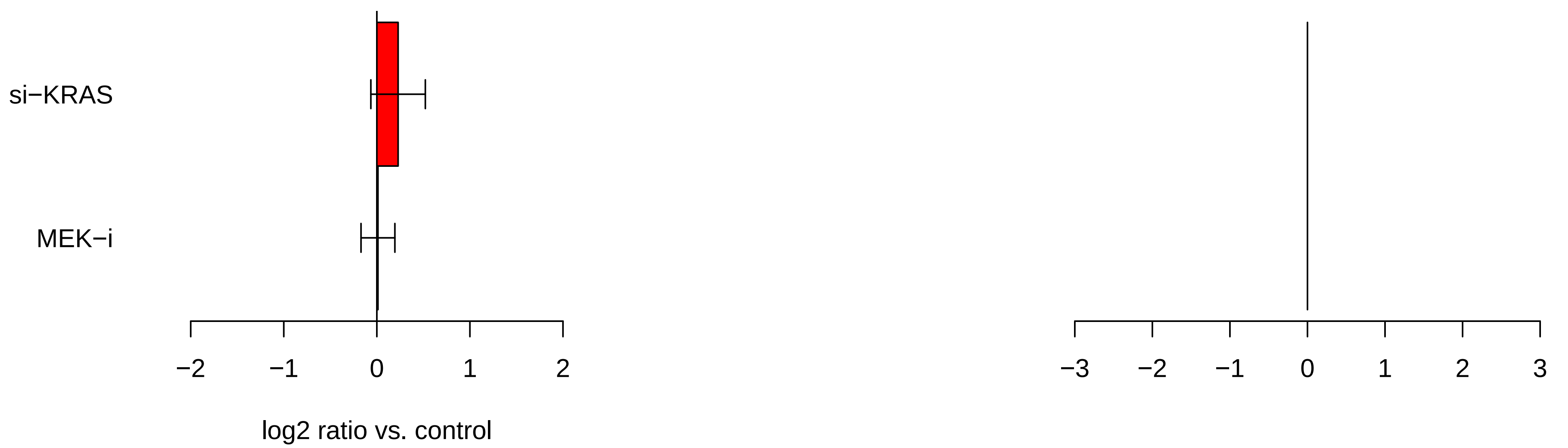

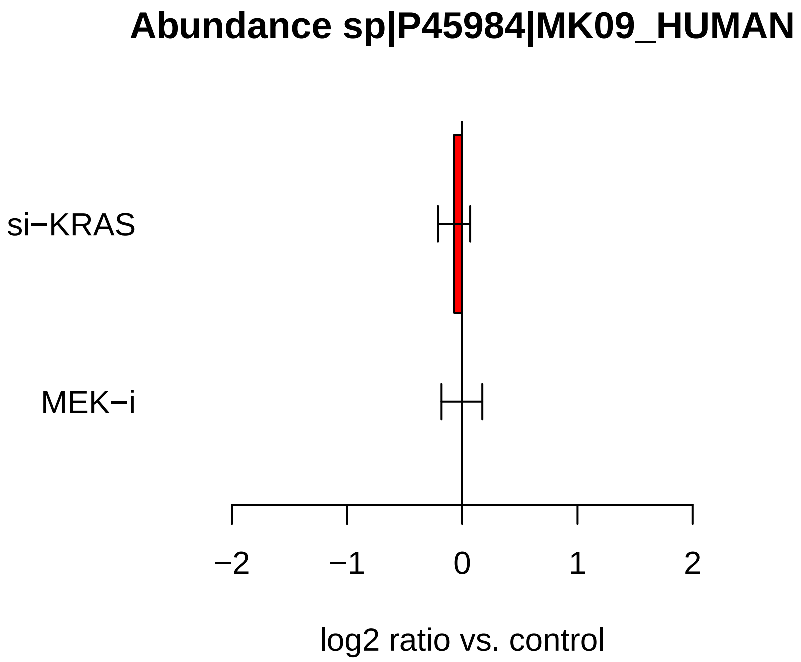

Abundance sp|P45984|MK09_HUMAN
si−KRAS
MEK−i
−2
−1
0
1
2
log2 ratio vs. control

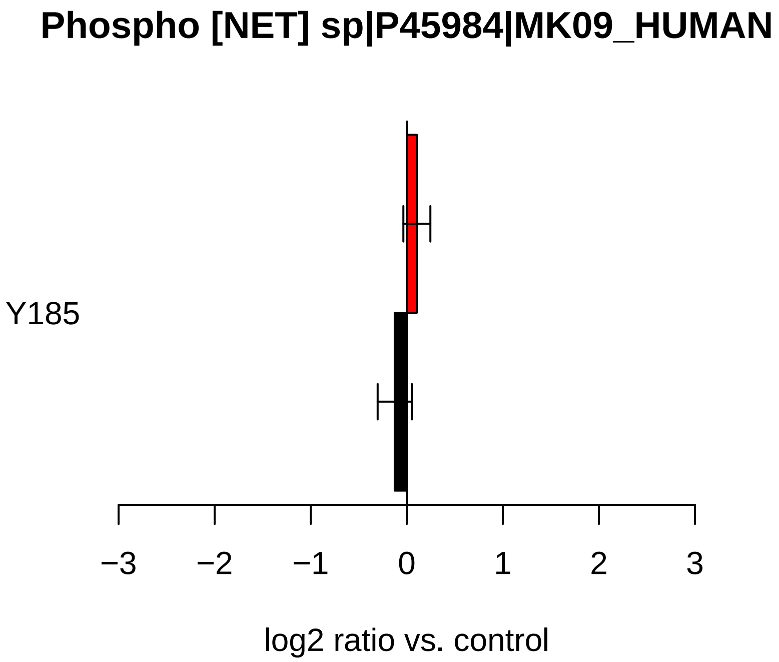

Phospho [NET] sp|P45984|MK09_HUMAN
Y185
−3
−2
−1
0
1
2
3
log2 ratio vs. control

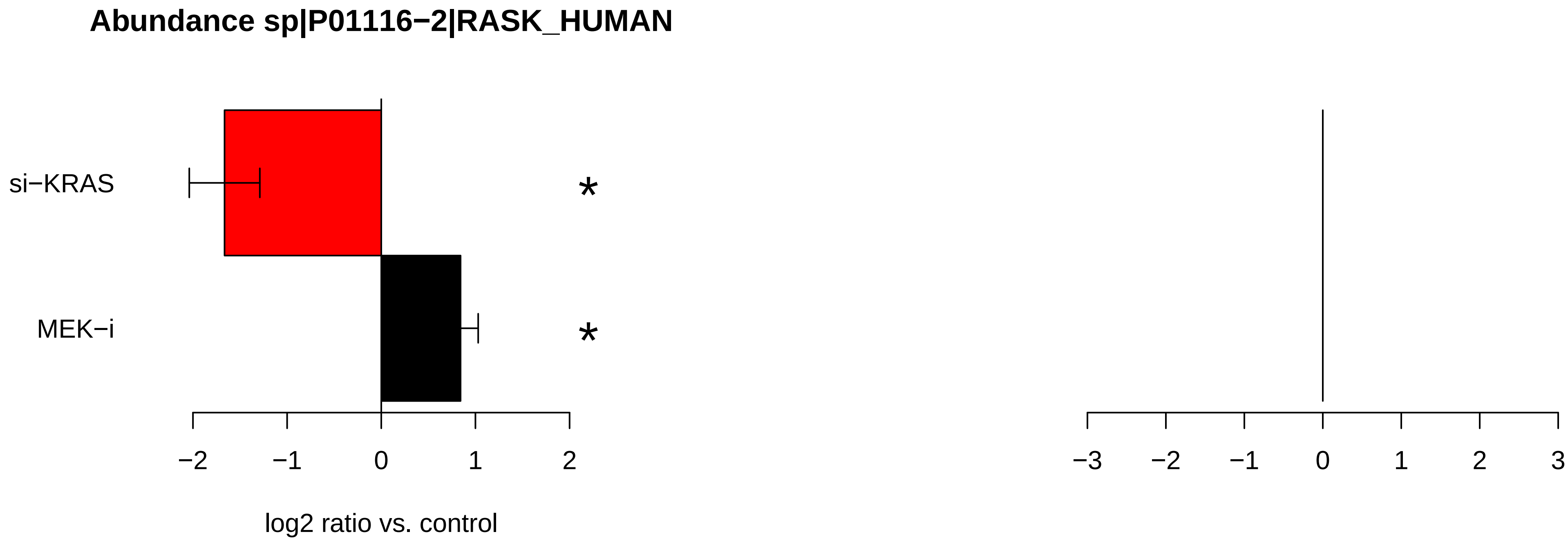
Abundance sp|P01116−2|RASK_HUMAN
si−KRAS
MEK−i
*
*
−2
−1
0
1
2
log2 ratio vs. control
−3
−2
−1
0
1
2
3

### Abundance sp|Q9UL18|AGO1_HUMAN

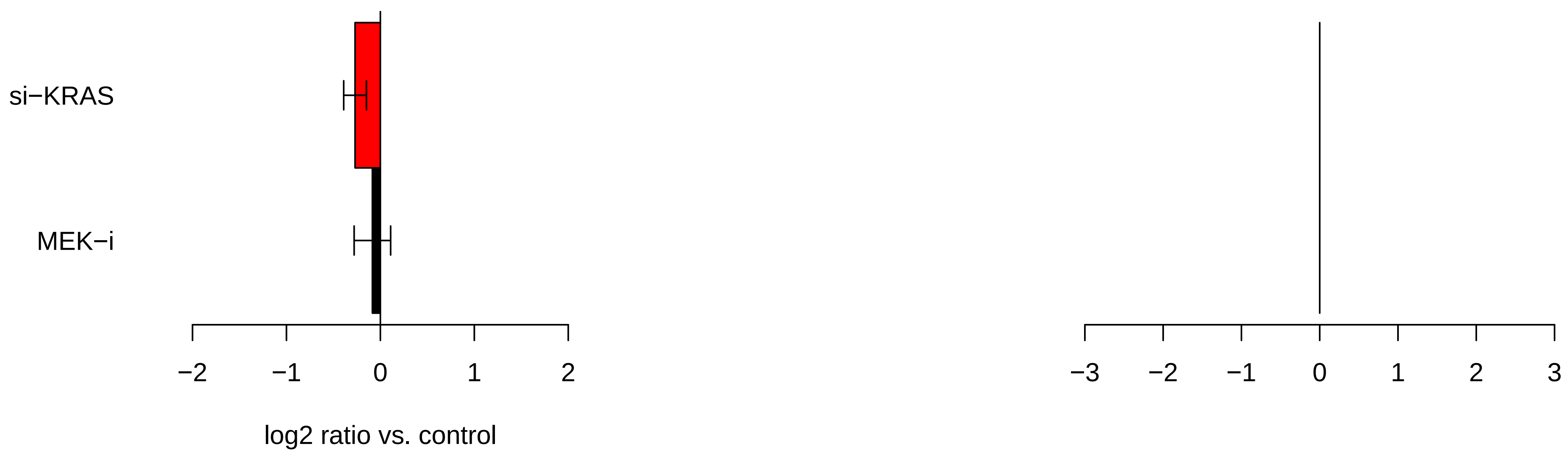

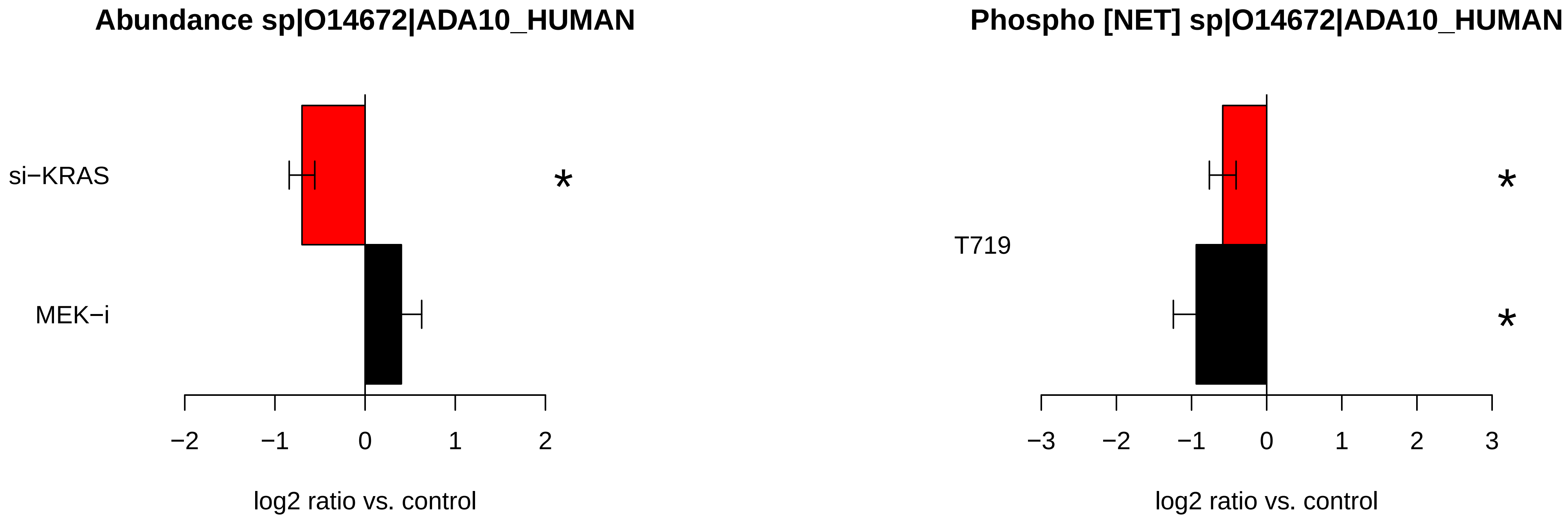

Abundance sp|O14672|ADA10_HUMAN
si−KRAS
MEK−i
*
−2
−1
0
1
2
log2 ratio vs. control
Phospho [NET] sp|O14672|ADA10_HUMAN
T719
*
*
−3
−2
−1
0
1
2
3
log2 ratio vs. control

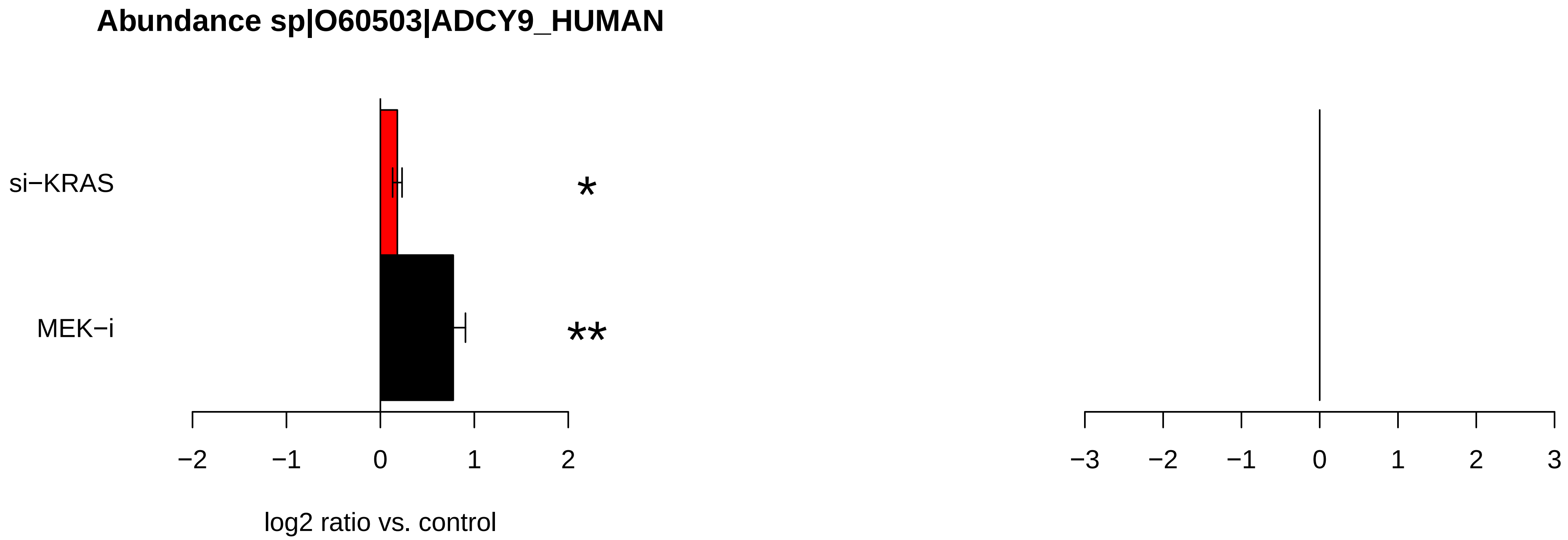
Abundance sp|O60503|ADCY9_HUMAN
si−KRAS
MEK−i
*
**
−2
−1
0
1
2
−3
−2
−1
0
1
2
3
log2 ratio vs. control

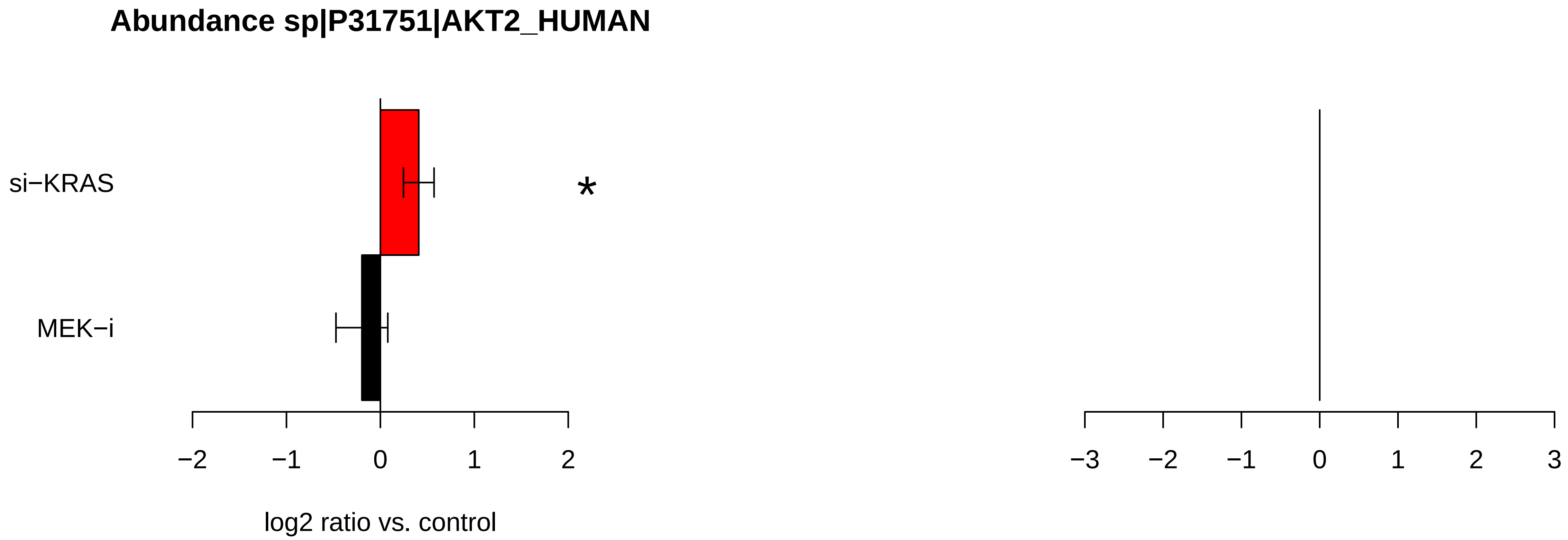
Abundance sp|P31751|AKT2_HUMAN
si−KRAS
MEK−i
*
−2
−1
0
1
2
log2 ratio vs. control
−3
−2
−1
0
1
2
3

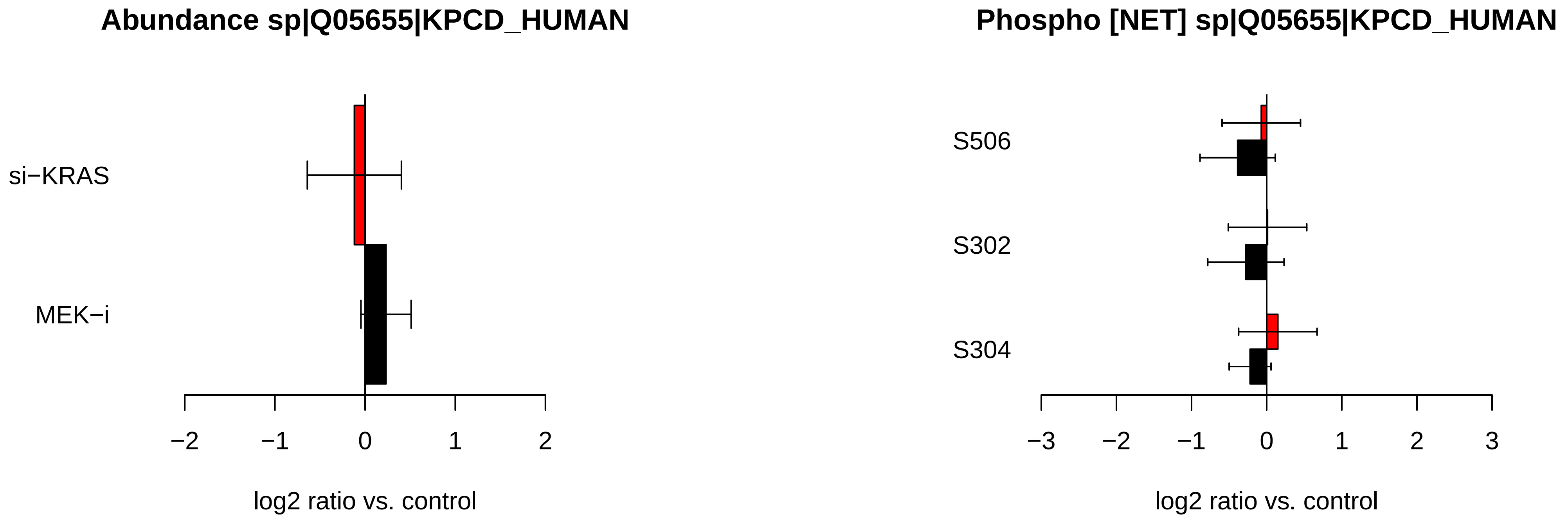
Abundance sp|Q05655|KPCD_HUMAN
si−KRAS
MEK−i
−2
−1
0
1
2
log2 ratio vs. control
Phospho [NET] sp|Q05655|KPCD_HUMAN
S506
S302
S304
−3
−2
−1
0
1
2
3
log2 ratio vs. control

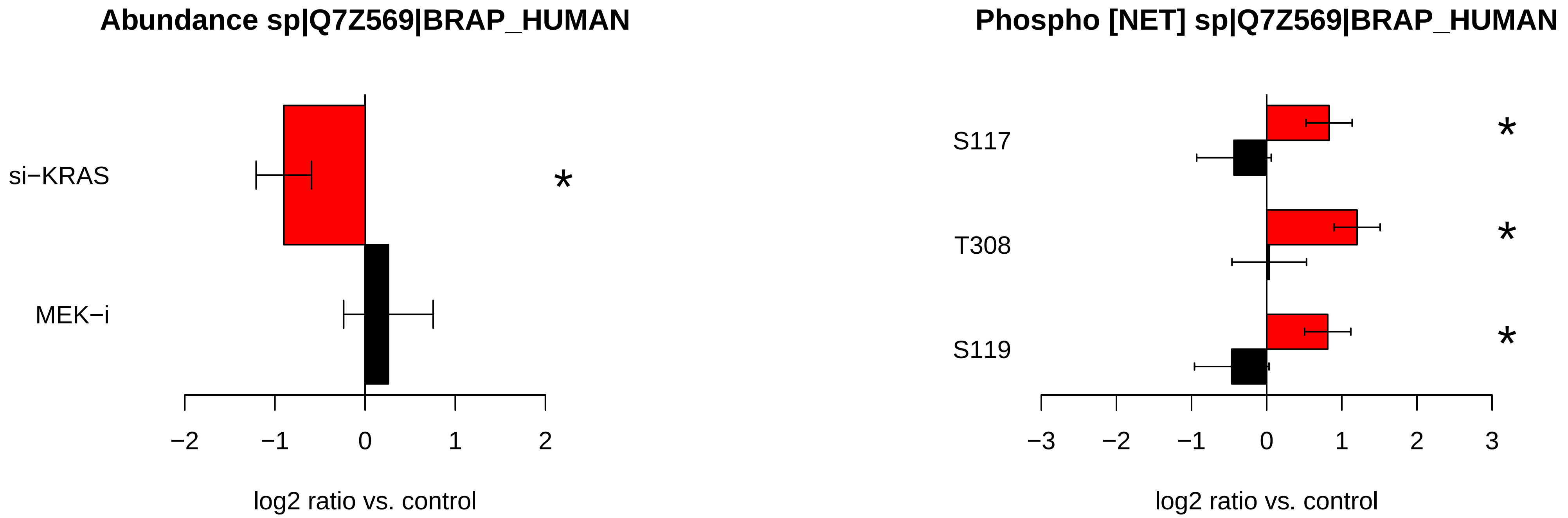
Abundance sp|Q7Z569|BRAP_HUMAN
si−KRAS
MEK−i
*
−2
−1
0
1
2
log2 ratio vs. control
Phospho [NET] sp|Q7Z569|BRAP_HUMAN
S117
T308
S119
*
*
*
−3
−2
−1
0
1
2
3
log2 ratio vs. control

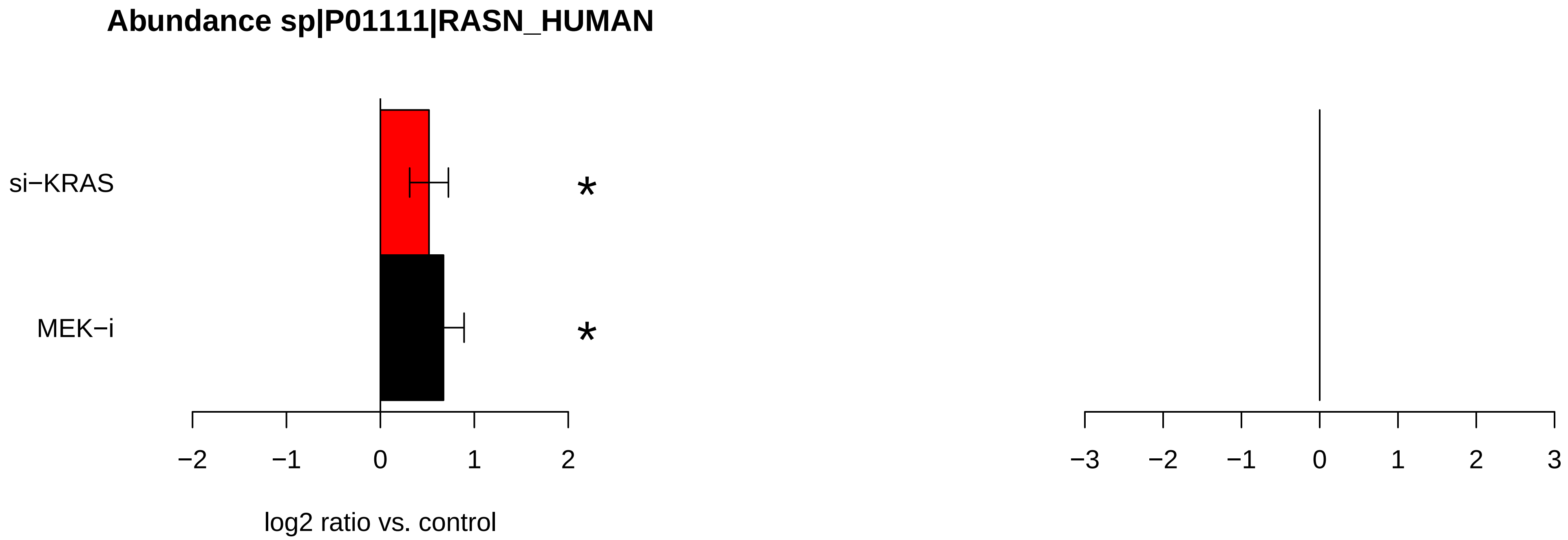
Abundance sp|P01111|RASN_HUMAN
si−KRAS
MEK−i
*
*
−2
−1
0
1
2
log2 ratio vs. control
−3
−2
−1
0
1
2
3

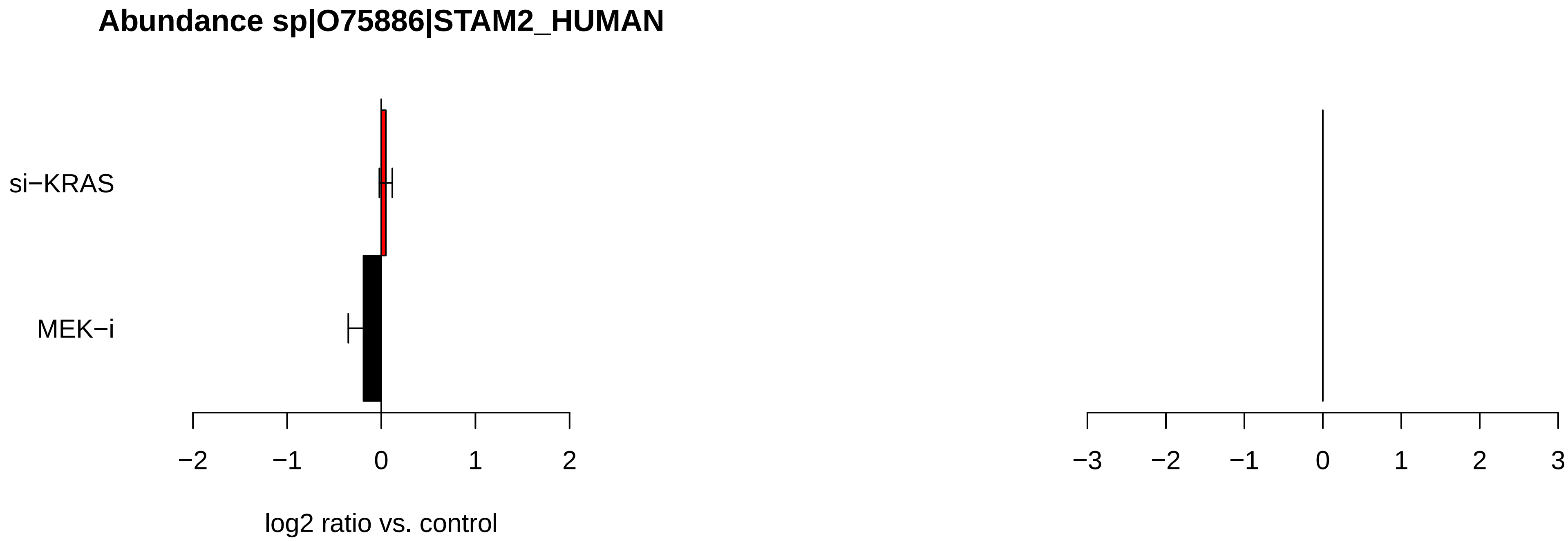

Abundance sp|O75886|STAM2_HUMAN
si−KRAS
MEK−i
−2
−1
0
1
2
log2 ratio vs. control
−3
−2
−1
0
1
2
3

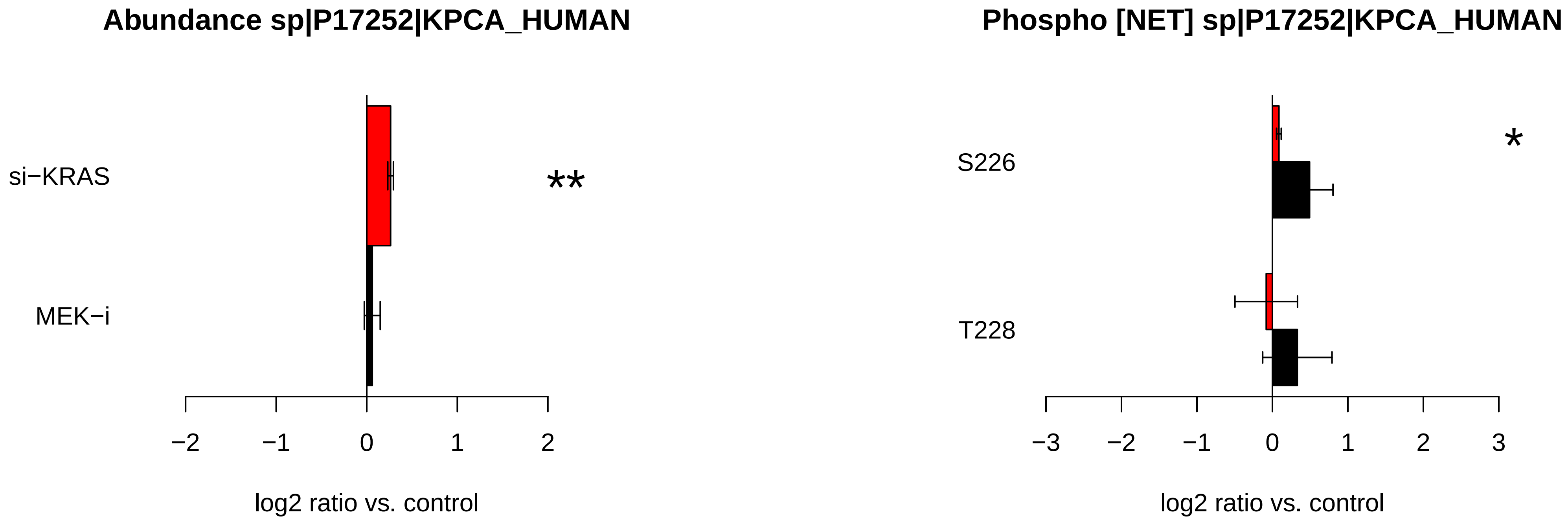

Abundance sp|P17252|KPCA_HUMAN
si−KRAS
MEK−i
**
−2
−1
0
1
2
log2 ratio vs. control
Phospho [NET] sp|P17252|KPCA_HUMAN
S226
T228
*
−3
−2
−1
0
1
2
3
log2 ratio vs. control

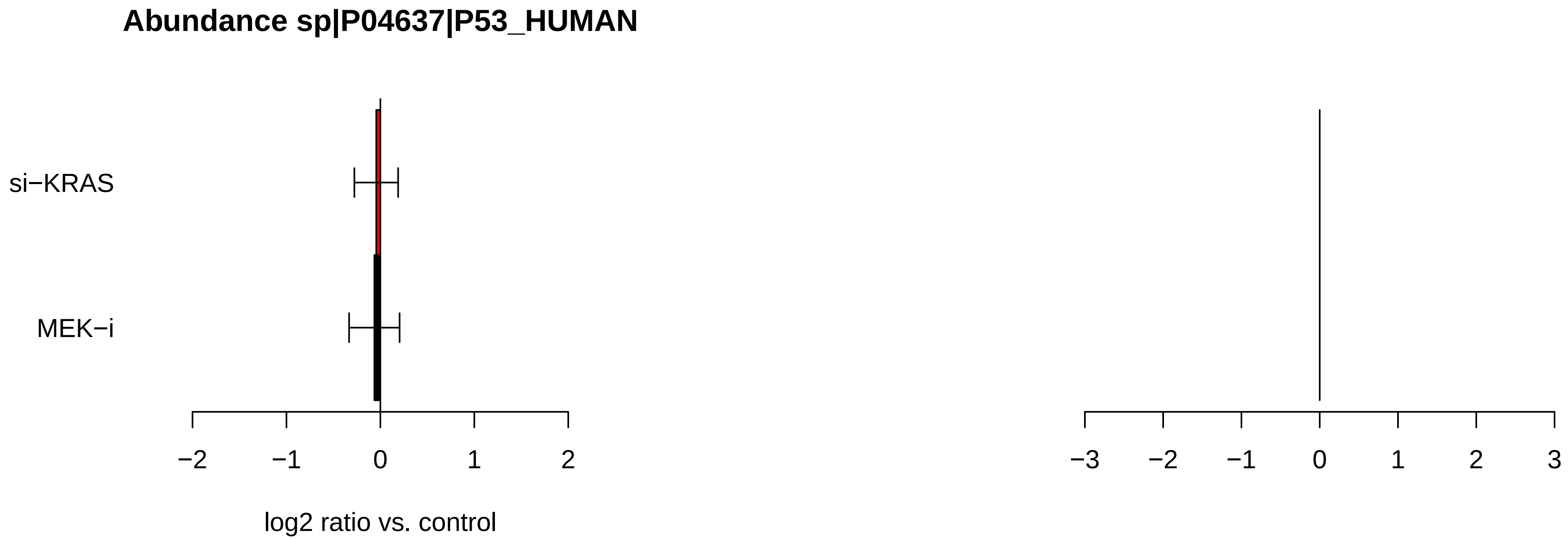
Abundance sp|P04637|P53_HUMAN
si−KRAS
MEK−i
−2
−1
0
1
2
log2 ratio vs. control
−3
−2
−1
0
1
2
3

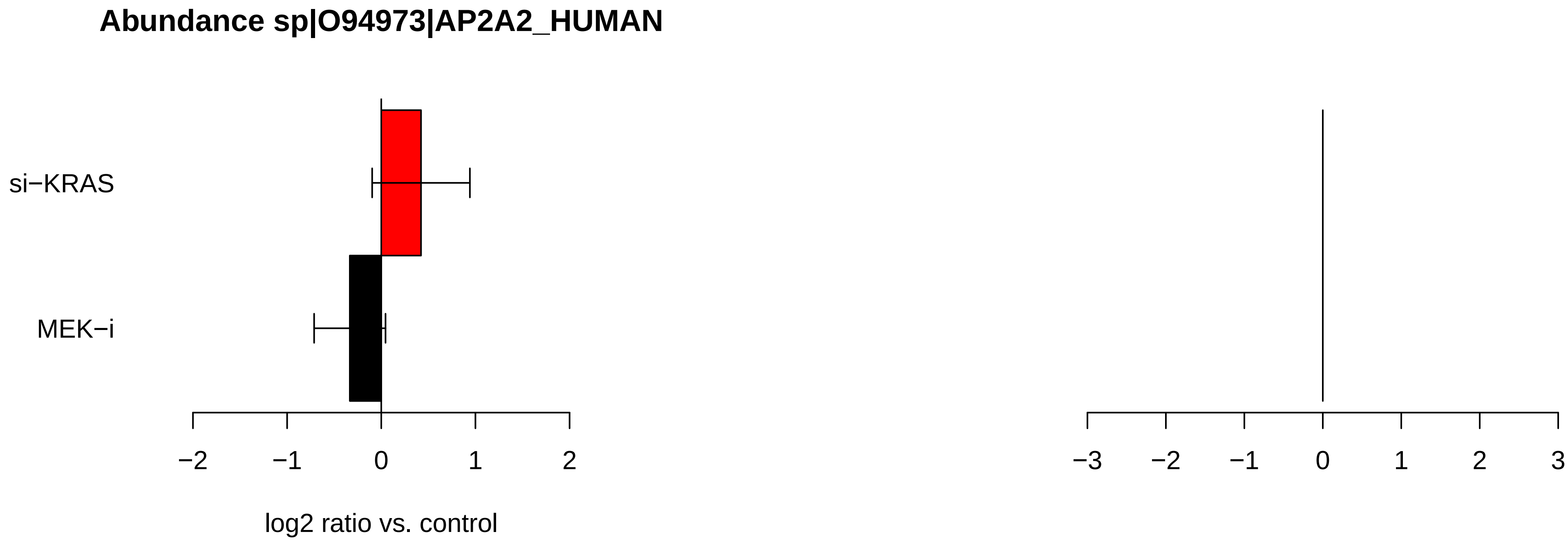
Abundance sp|O94973|AP2A2_HUMAN
si−KRAS
MEK−i
−2
−1
0
1
2
log2 ratio vs. control
−3
−2
−1
0
1
2
3

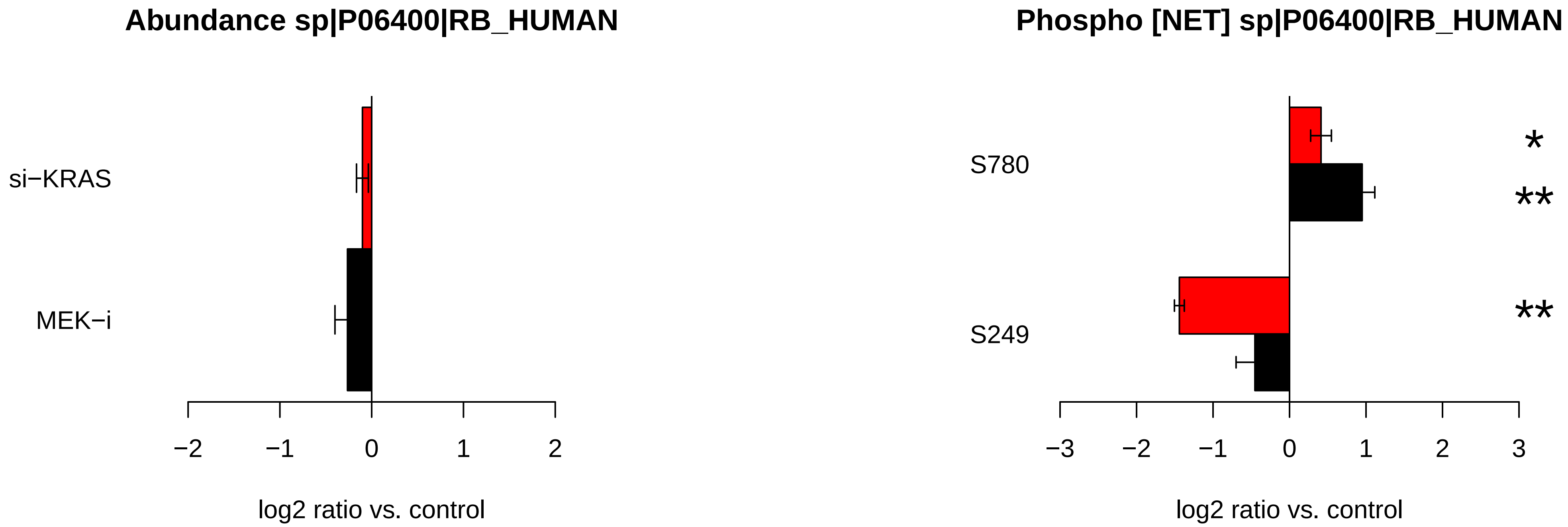

Abundance sp|P06400|RB_HUMAN
si−KRAS
MEK−i
−2
−1
0
1
2
log2 ratio vs. control
Phospho [NET] sp|P06400|RB_HUMAN
S780
S249
*
**
**
−3
−2
−1
0
1
2
3
log2 ratio vs. control

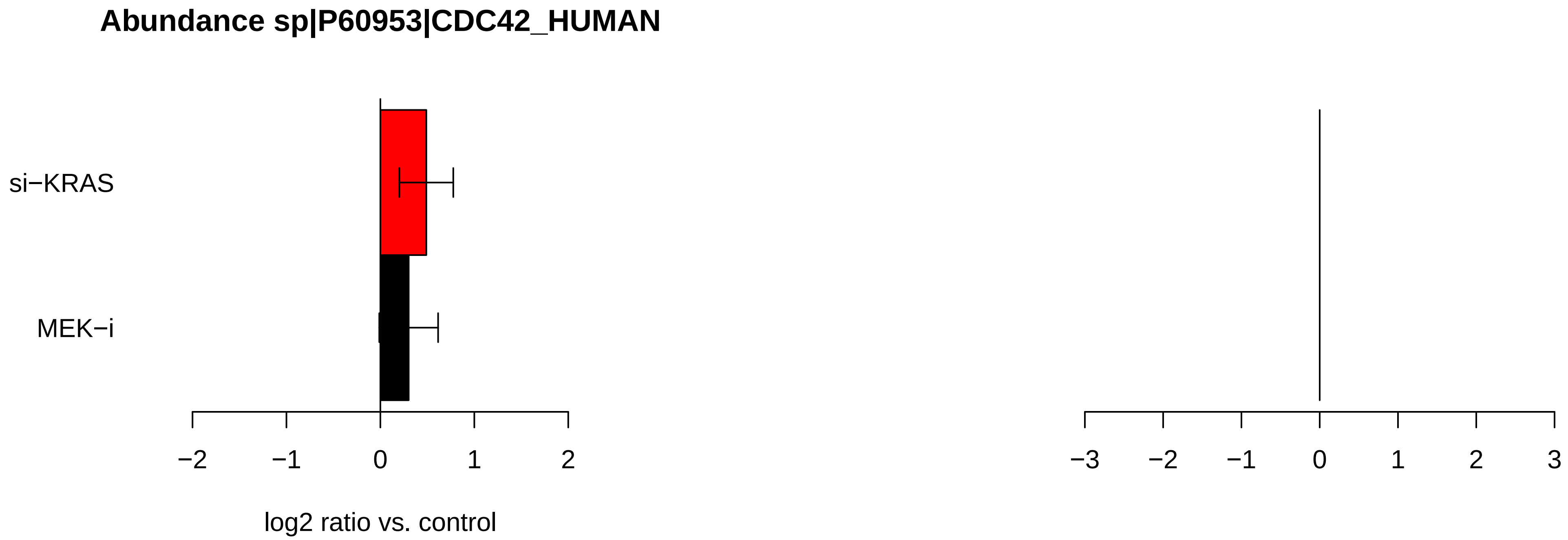
Abundance sp|P60953|CDC42_HUMAN
si−KRAS
MEK−i
−2
−1
0
1
2
log2 ratio vs. control
−3
−2
−1
0
1
2
3

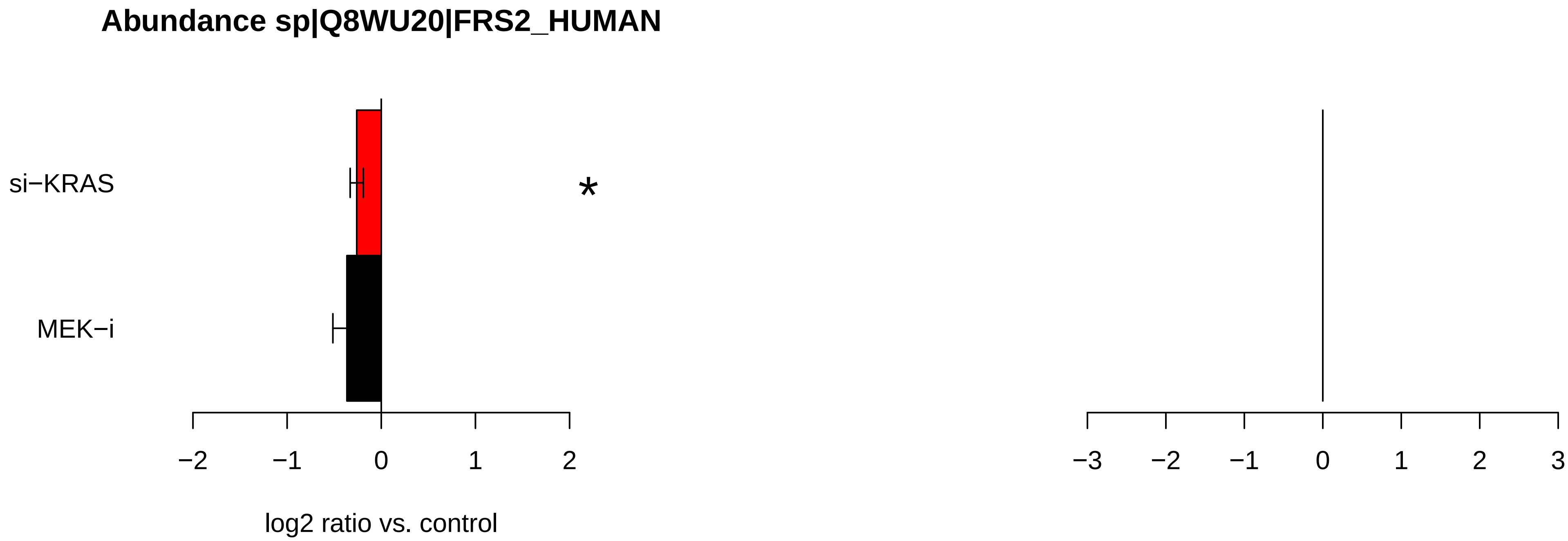

Abundance sp|Q8WU20|FRS2_HUMAN
si−KRAS
MEK−i
*
−2
−1
0
1
2
log2 ratio vs. control
−3
−2
−1
0
1
2
3

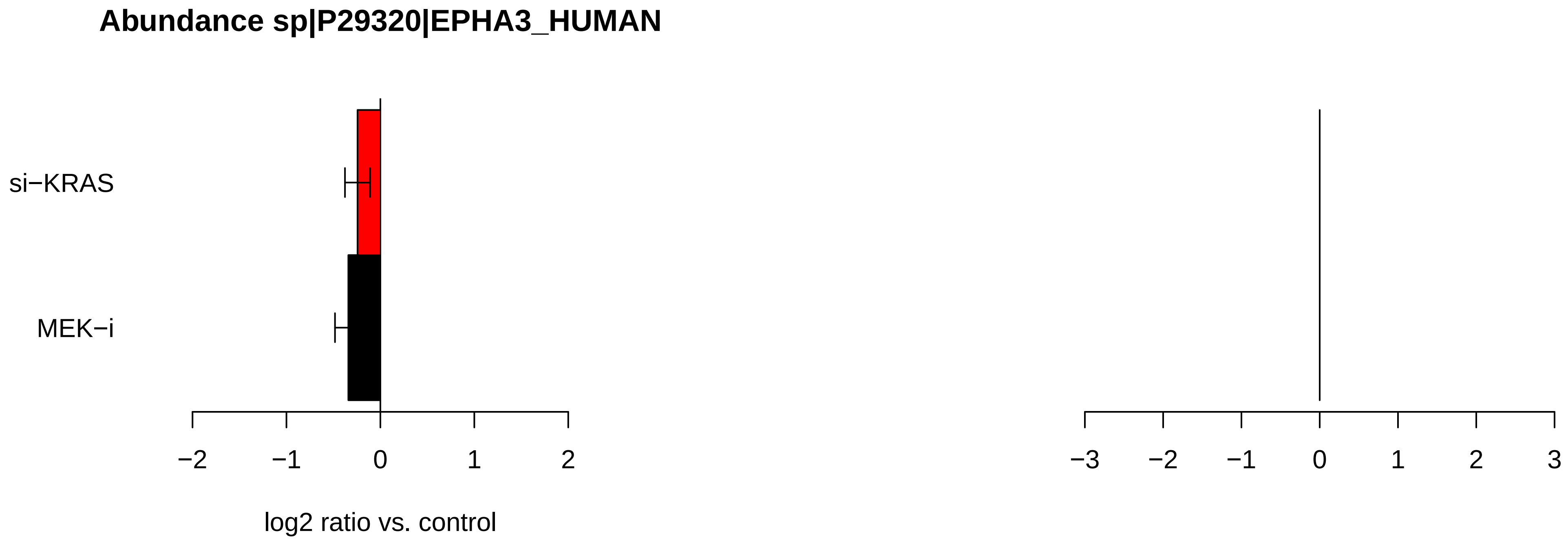
Abundance sp|P29320|EPHA3_HUMAN
si−KRAS
MEK−i
−2
−1
0
1
2
log2 ratio vs. control
−3
−2
−1
0
1
2
3

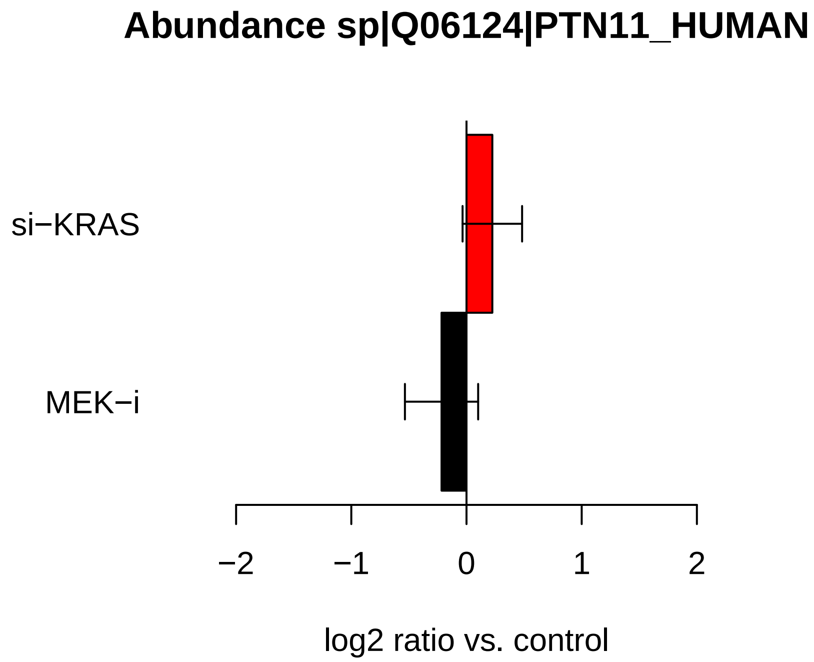
Abundance sp|Q06124|PTN11_HUMAN
si−KRAS
MEK−i
−2
−1
0
1
2
log2 ratio vs. control

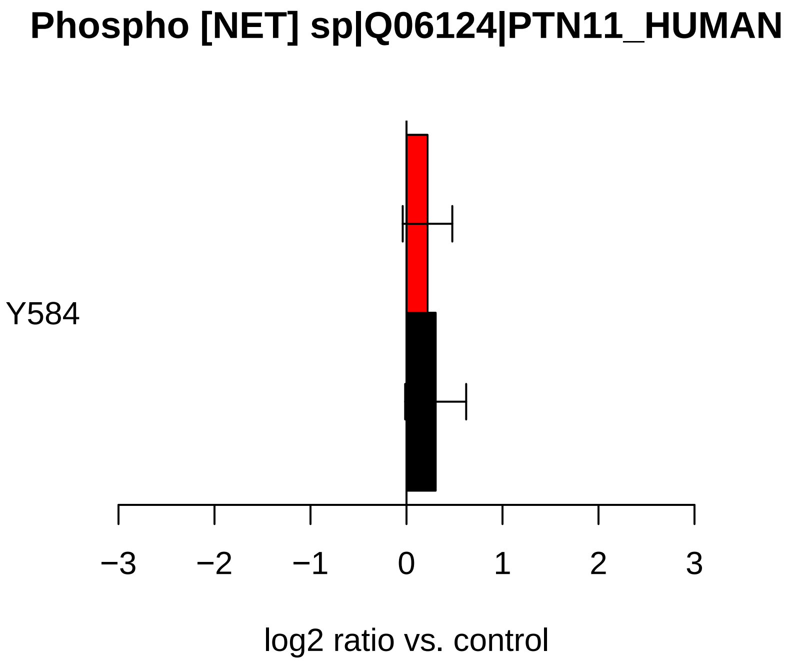
Phospho [NET] sp|Q06124|PTN11_HUMAN
Y584
−3
−2
−1
0
1
2
3
log2 ratio vs. control

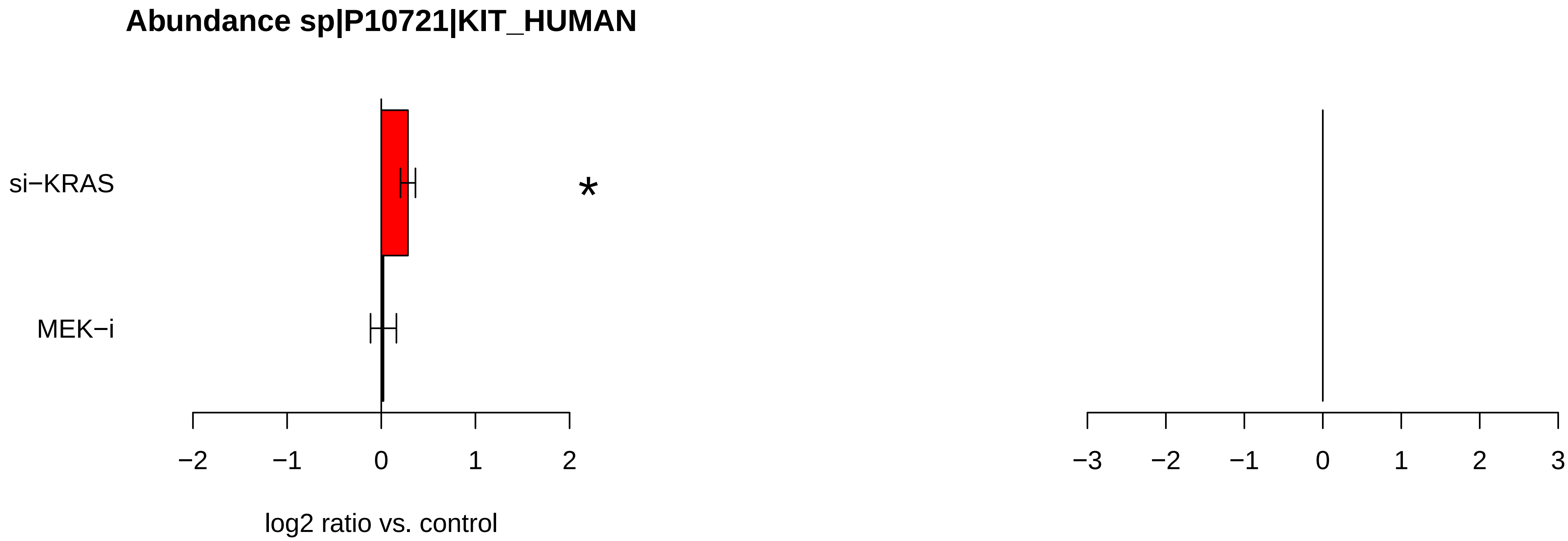

Abundance sp|P10721|KIT_HUMAN
si−KRAS
MEK−i
*
−2
−1
0
1
2
log2 ratio vs. control
−3
−2
−1
0
1
2
3

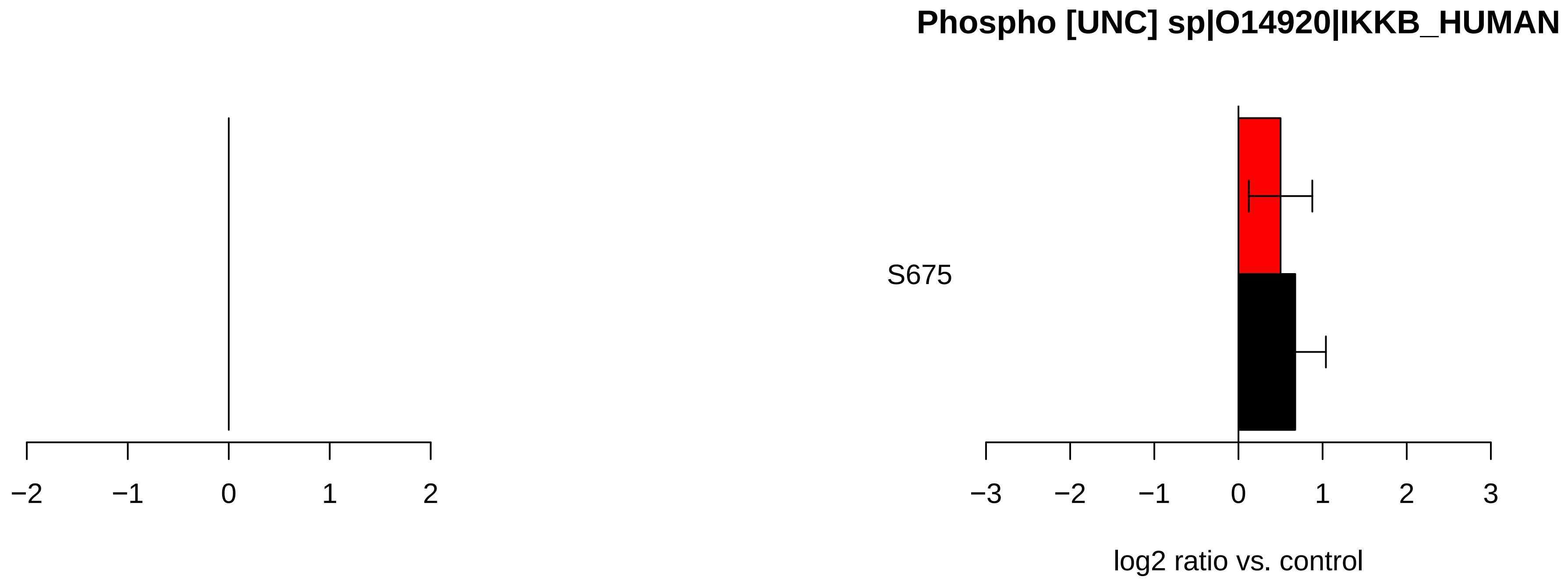

Phospho [UNC] sp|O14920|IKKB_HUMAN
S675
−2
−1
0
1
2
−3
−2
−1
0
1
2
3
log2 ratio vs. control

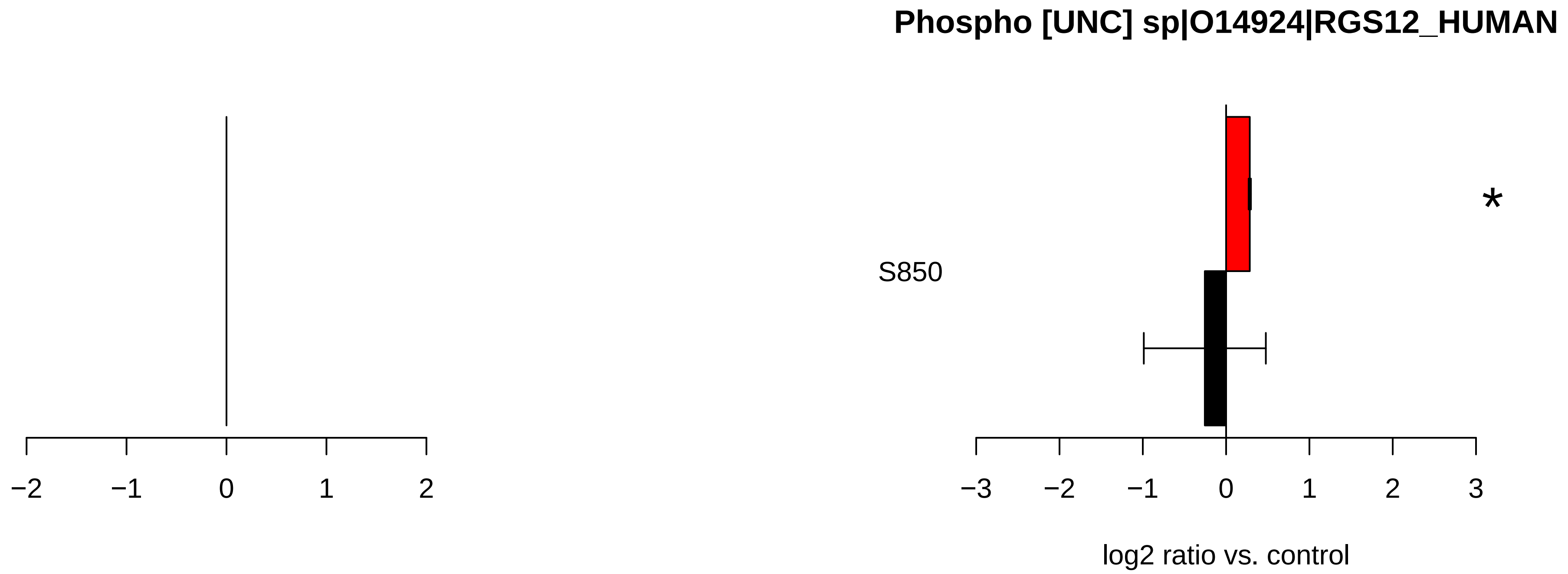
Phospho [UNC] sp|O14924|RGS12_HUMAN
S850
*
−2
−1
0
1
2
−3
−2
−1
0
1
2
3
log2 ratio vs. control

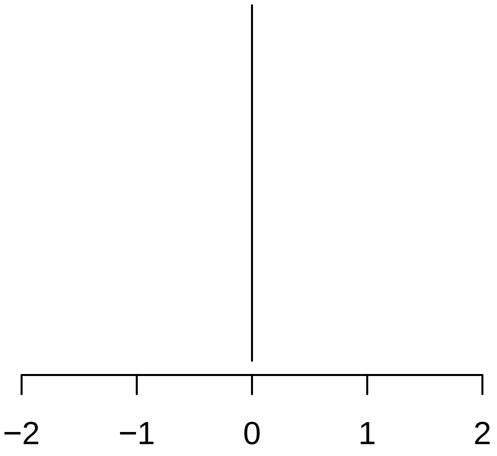

−2
−1
0
1
2


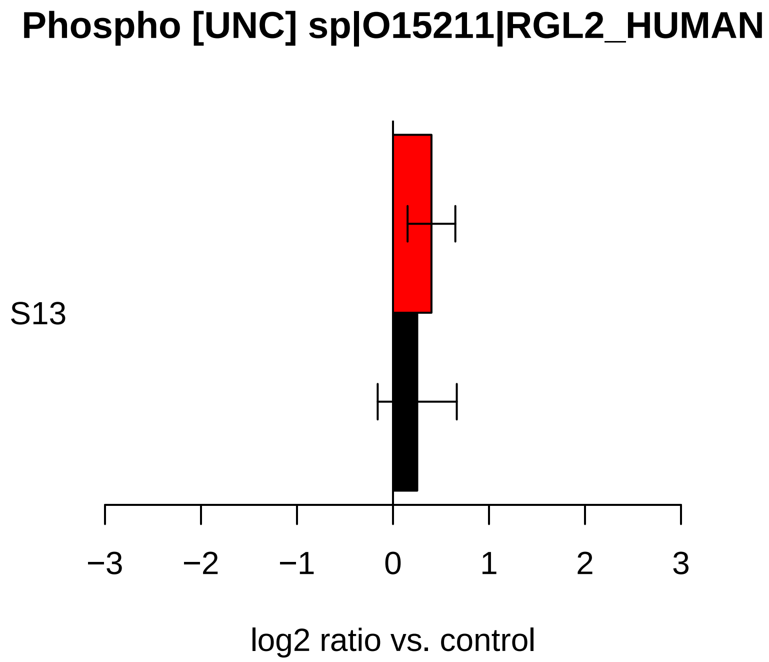

Phospho [UNC] sp|O15211|RGL2_HUMAN
S13
−3
−2
−1
0
1
2
3
log2 ratio vs. control

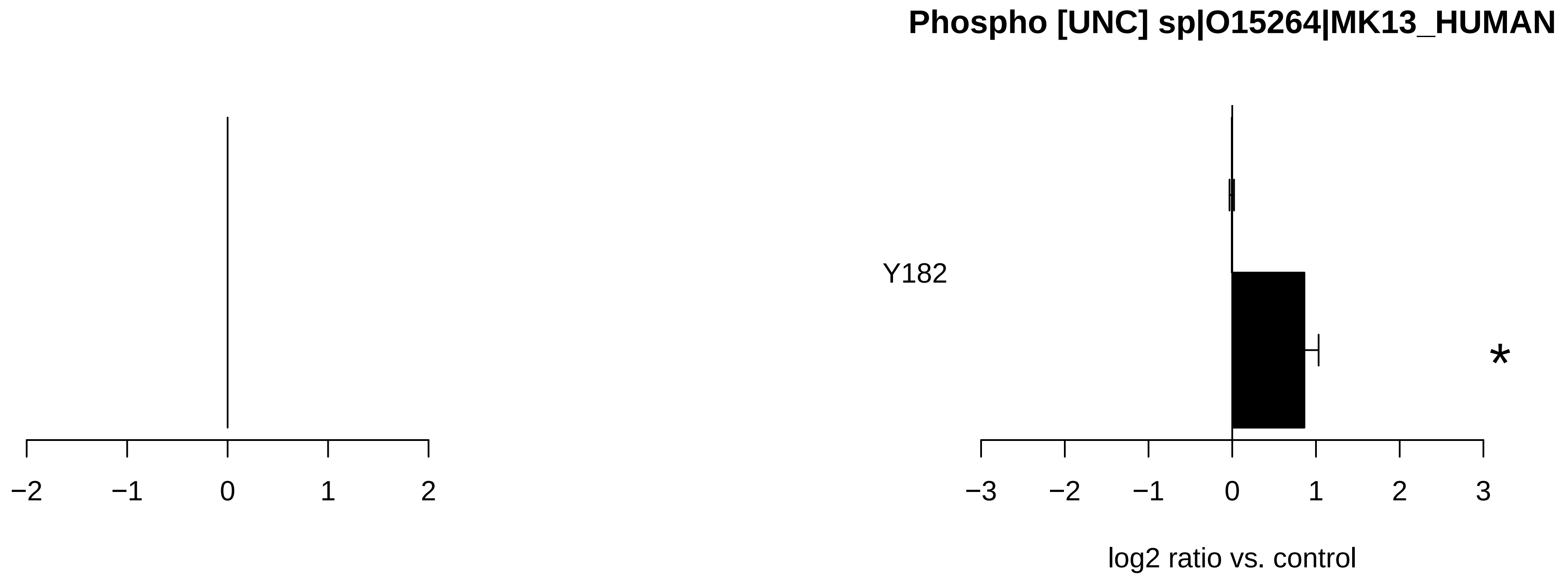
Phospho [UNC] sp|O15264|MK13_HUMAN
Y182
*
−2
−1
0
1
2
−3
−2
−1
0
1
2
3
log2 ratio vs. control

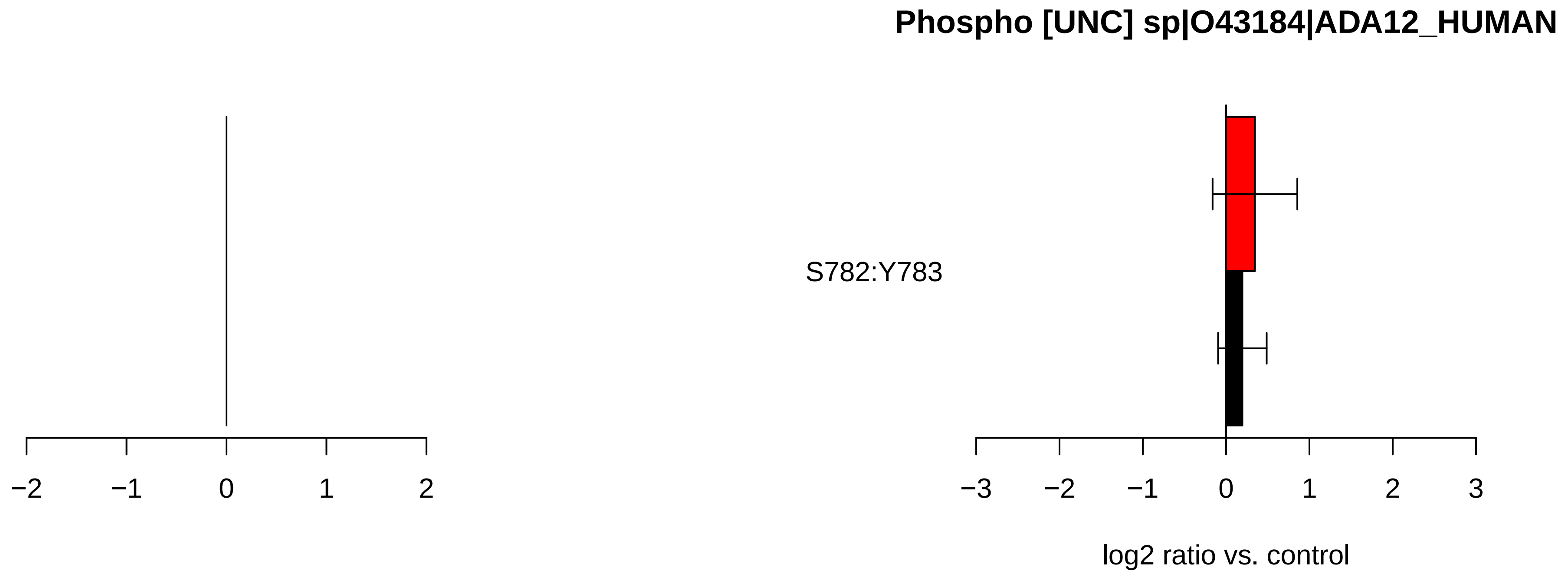

Phospho [UNC] sp|O43184|ADA12_HUMAN
S782:Y783
−2
−1
0
1
2
−3
−2
−1
0
1
2
3
log2 ratio vs. control

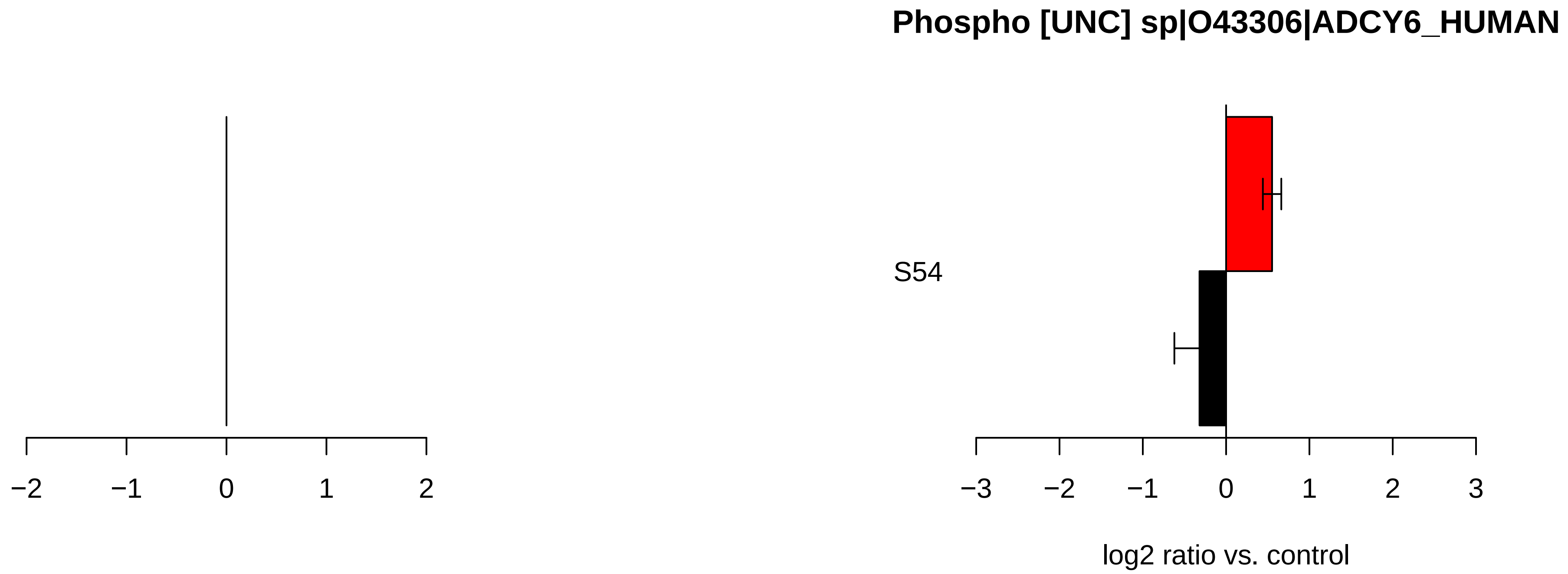

Phospho [UNC] sp|O43306|ADCY6_HUMAN
S54
−2
−1
0
1
2
−3
−2
−1
0
1
2
3
log2 ratio vs. control

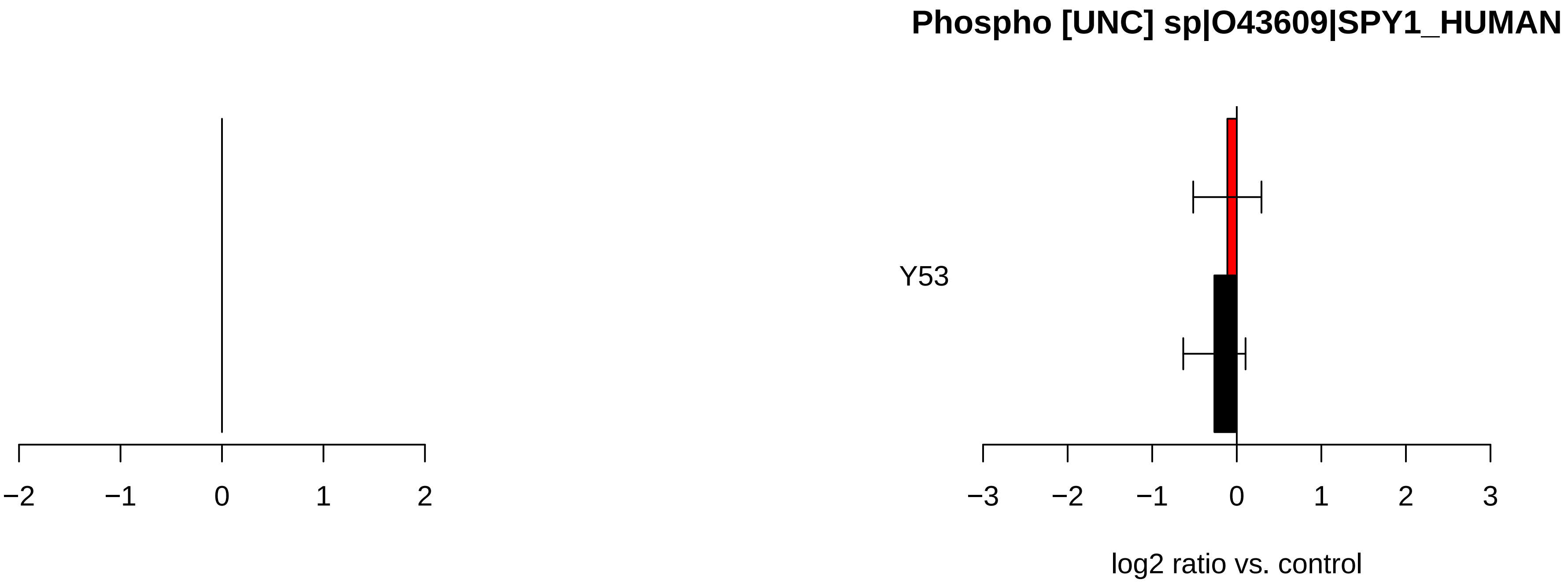
Phospho [UNC] sp|O43609|SPY1_HUMAN
Y53
−2
−1
0
1
2
−3
−2
−1
0
1
2
3
log2 ratio vs. control

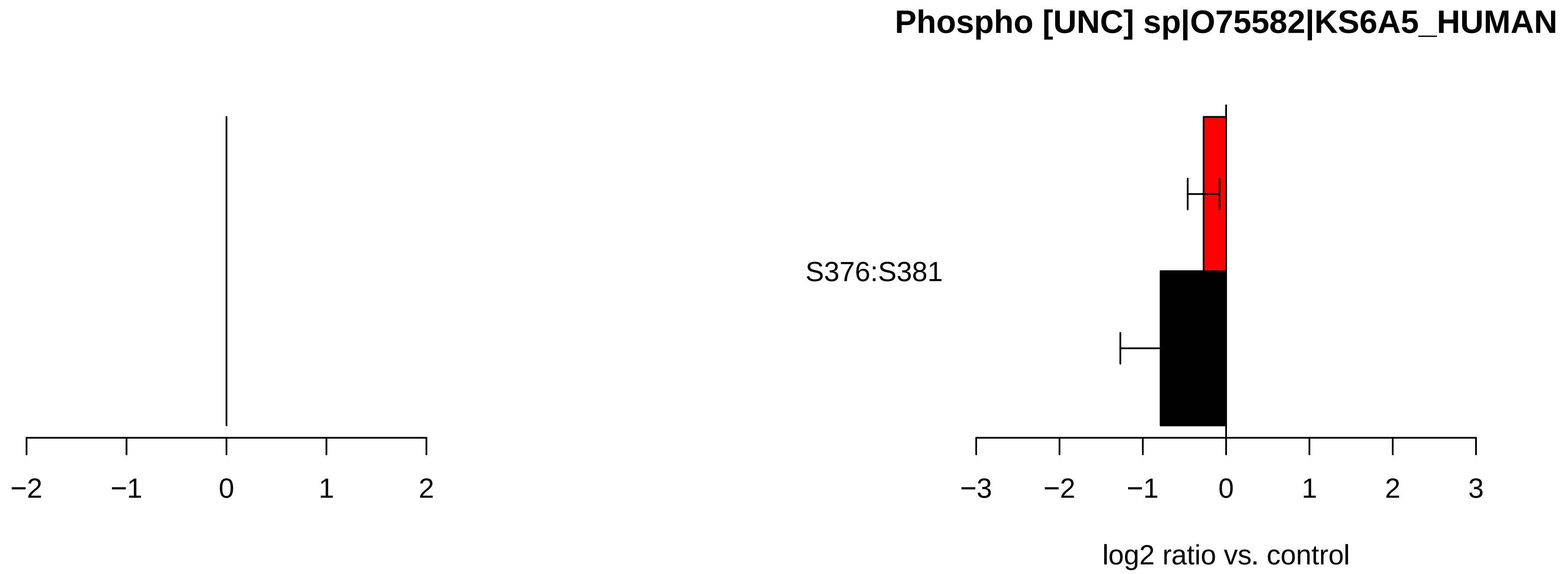

Phospho [UNC] sp|O75582|KS6A5_HUMAN
S376:S381
−2
−1
0
1
2
−3
−2
−1
0
1
2
3
log2 ratio vs. control

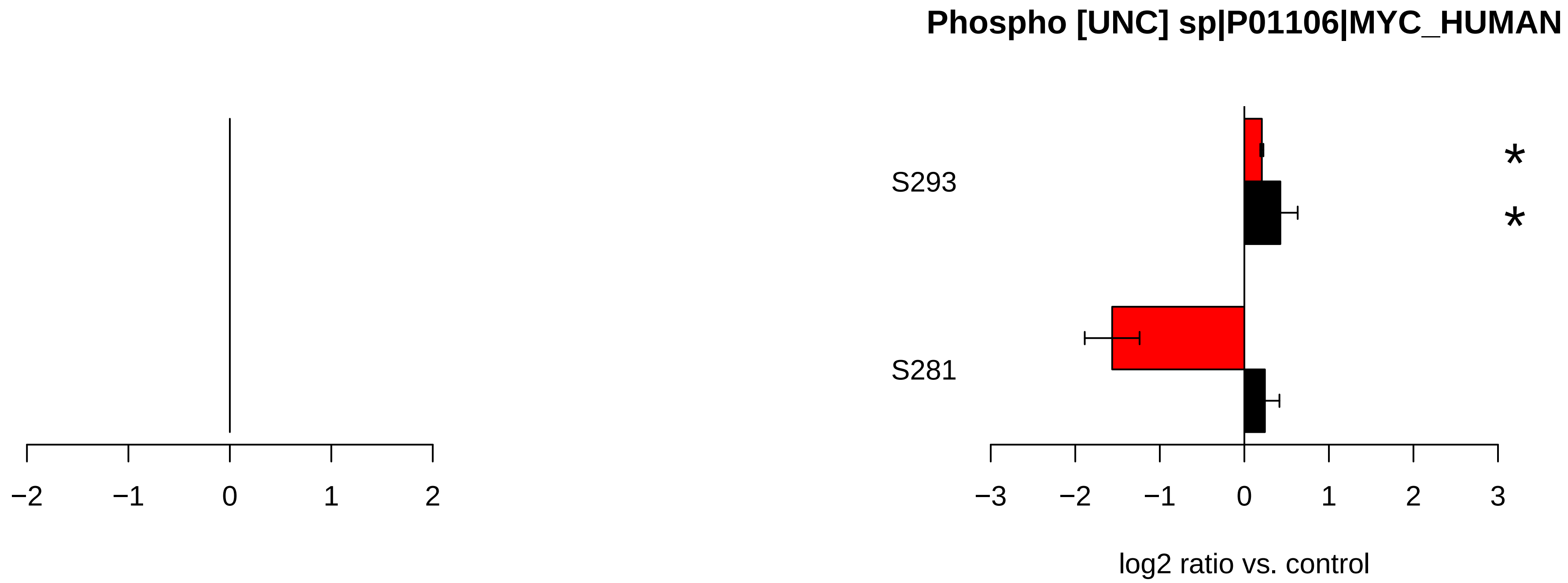

Phospho [UNC] sp|P01106|MYC_HUMAN
S293
S281
*
*
−2
−1
0
1
2
−3
−2
−1
0
1
2
3
log2 ratio vs. control

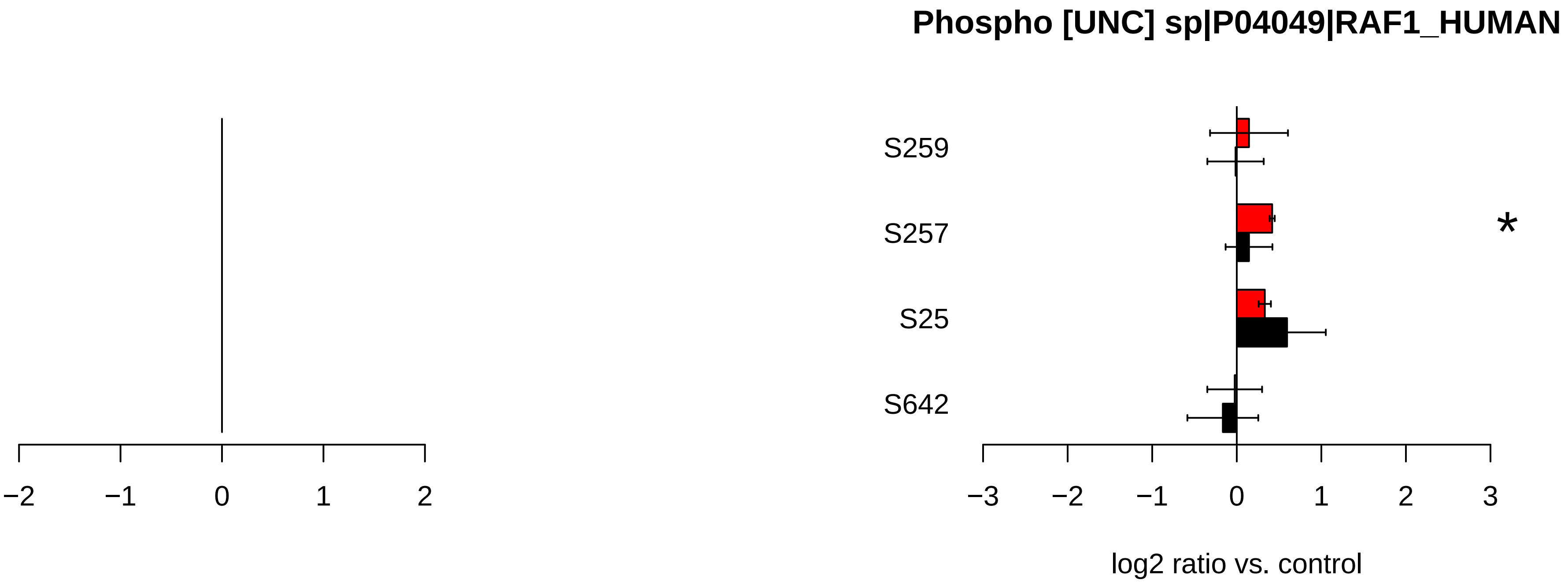

Phospho [UNC] sp|P04049|RAF1_HUMAN
S259
S257
S25
S642
*
−2
−1
0
1
2
−3
−2
−1
0
1
2
3
log2 ratio vs. control

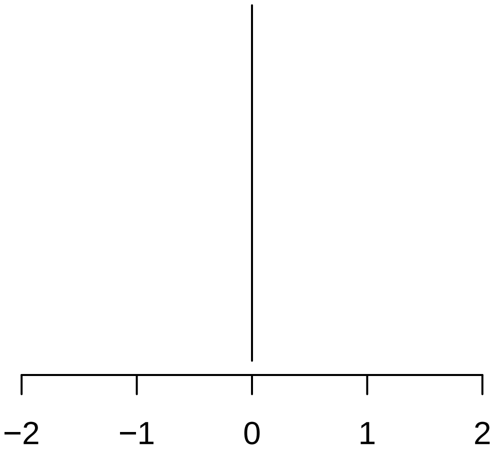


### Phospho [UNC] sp|P15056|BRAF_HUMAN

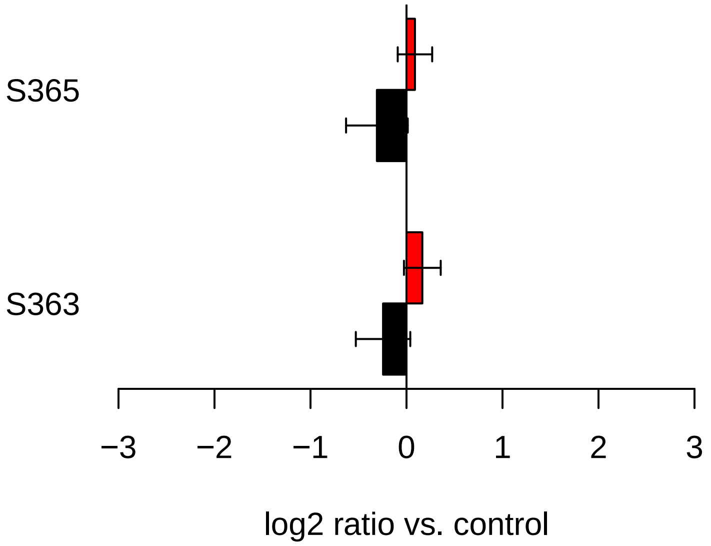

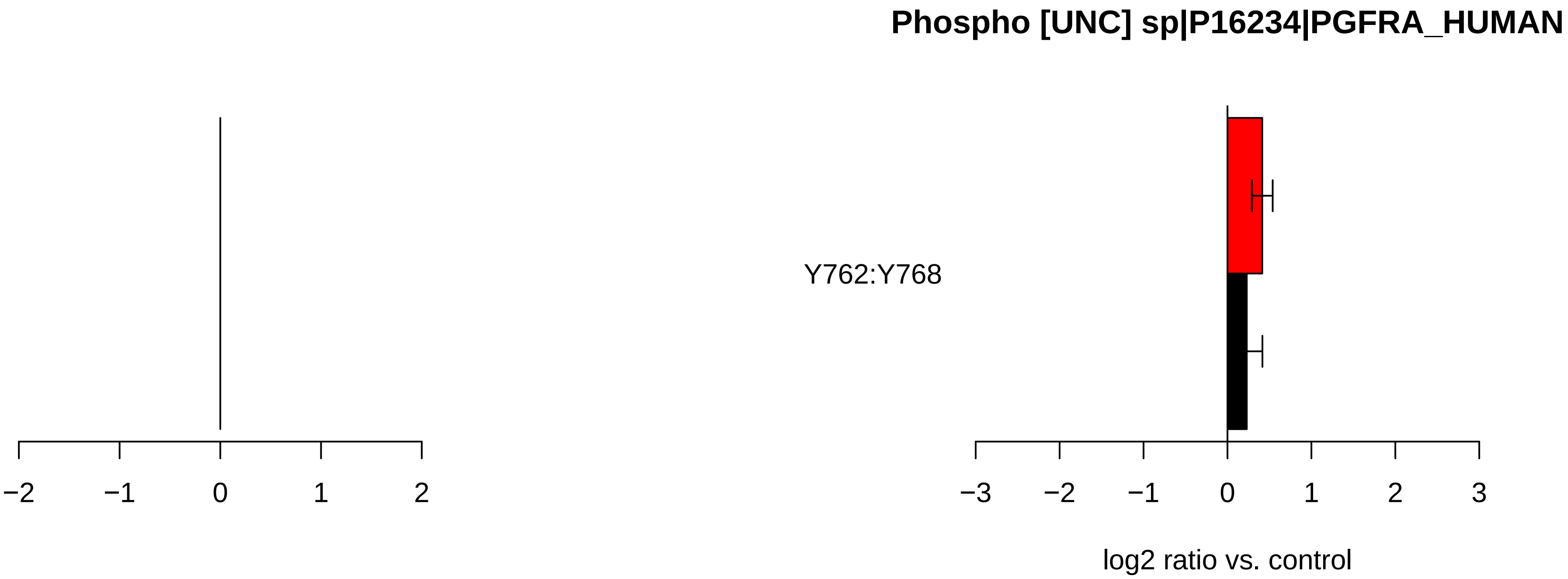
Phospho [UNC] sp|P16234|PGFRA_HUMAN
Y762:Y768
−2
−1
0
1
2
−3
−2
−1
0
1
2
3
log2 ratio vs. control

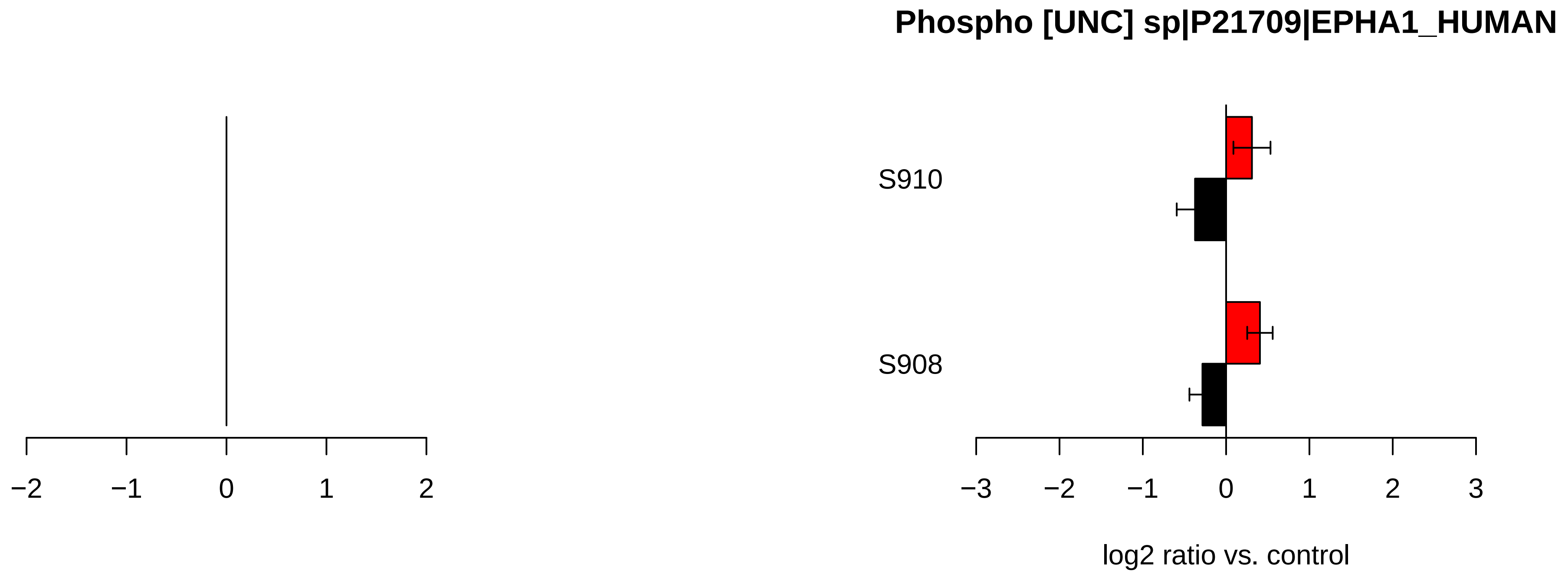

Phospho [UNC] sp|P21709|EPHA1_HUMAN
S910
S908
−2
−1
0
1
2
−3
−2
−1
0
1
2
3
log2 ratio vs. control

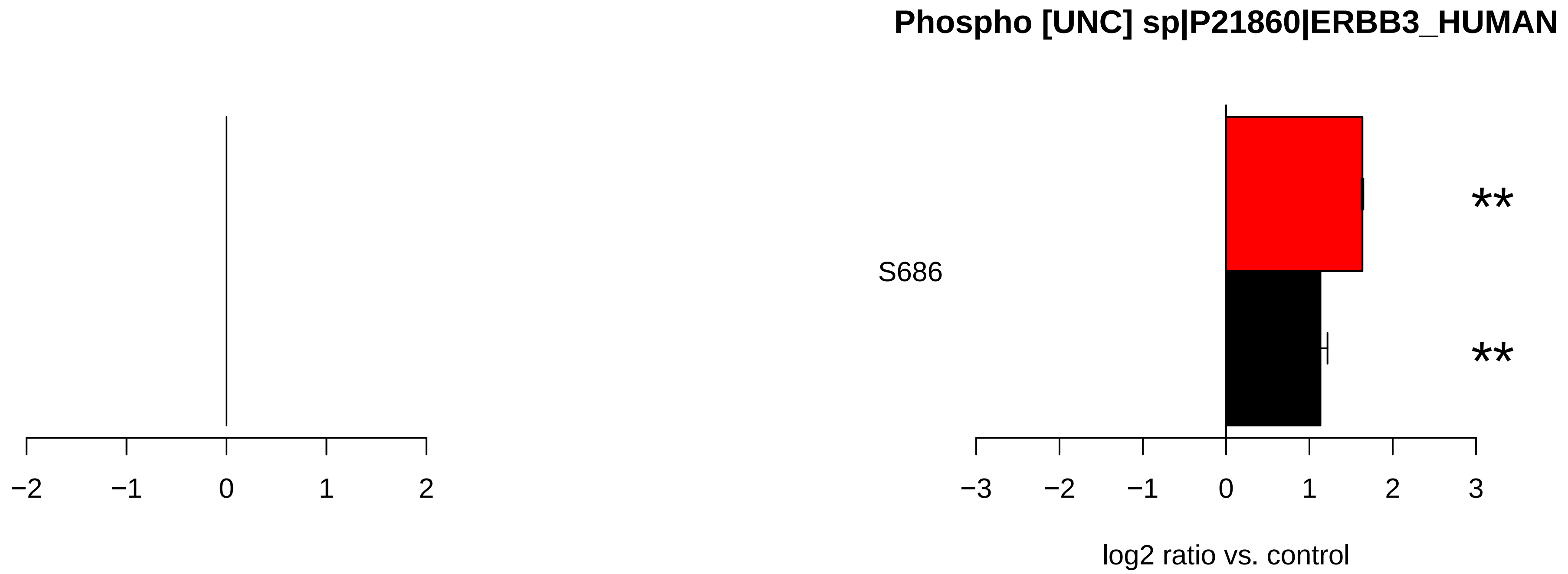
Phospho [UNC] sp|P21860|ERBB3_HUMAN
S686
**
**
−2
−1
0
1
2
−3
−2
−1
0
1
2
3
log2 ratio vs. control

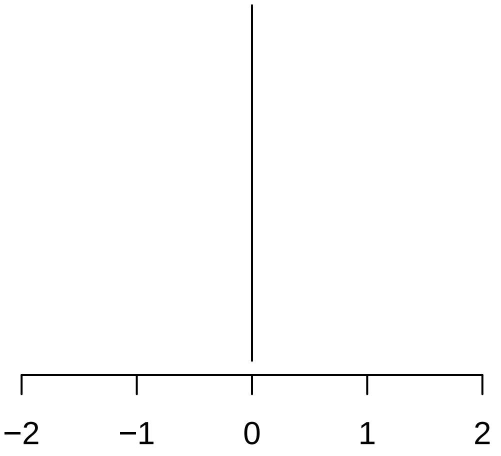

−2
−1
0
1
2


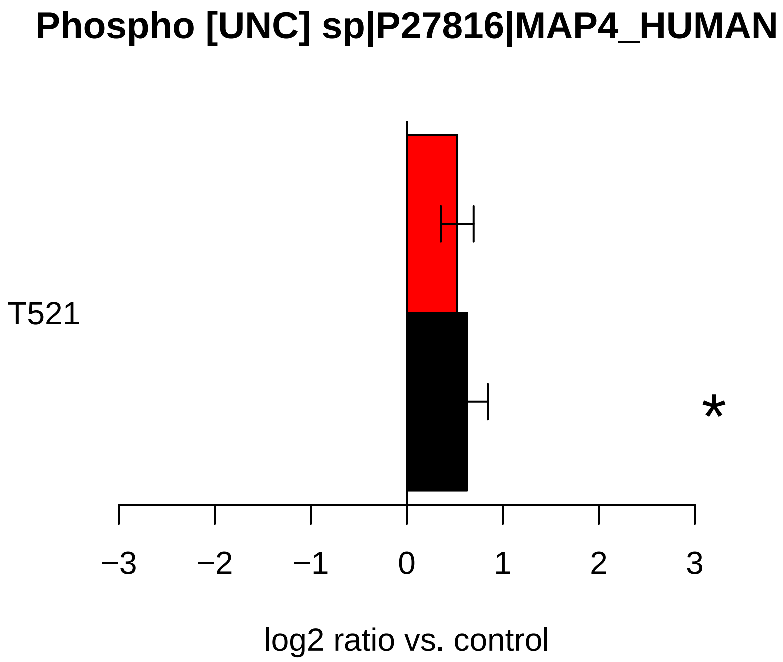

Phospho [UNC] sp|P27816|MAP4_HUMAN
T521
*
−3
−2
−1
0
1
2
3
log2 ratio vs. control

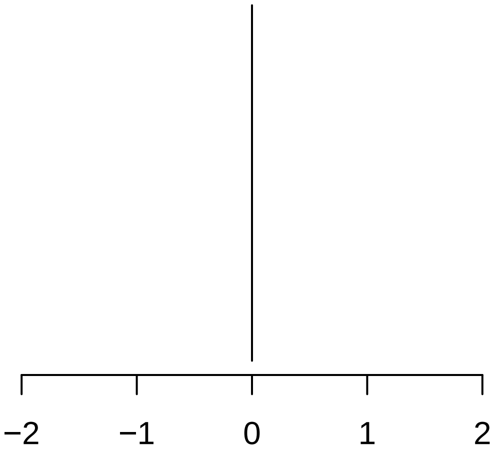

−2
−1
0
1
2


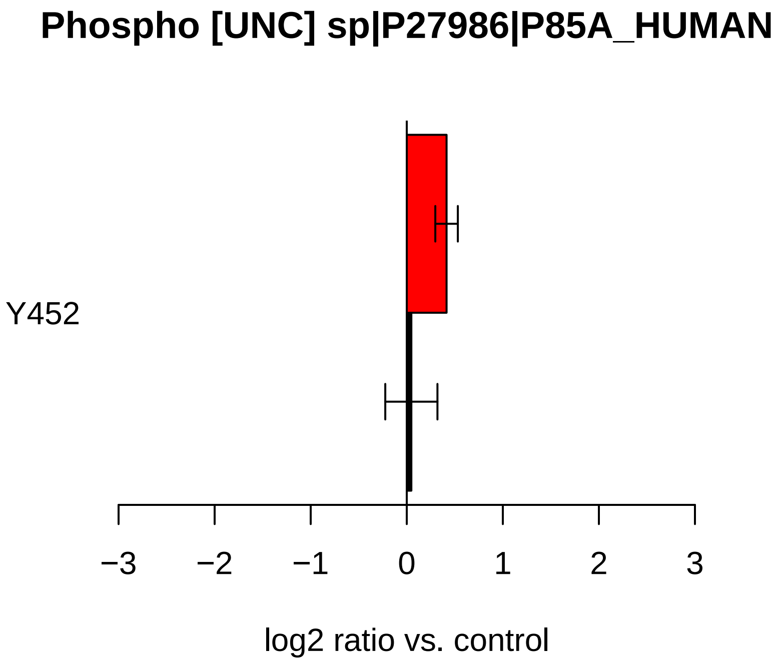

Phospho [UNC] sp|P27986|P85A_HUMAN
Y452
−3
−2
−1
0
1
2
3
log2 ratio vs. control

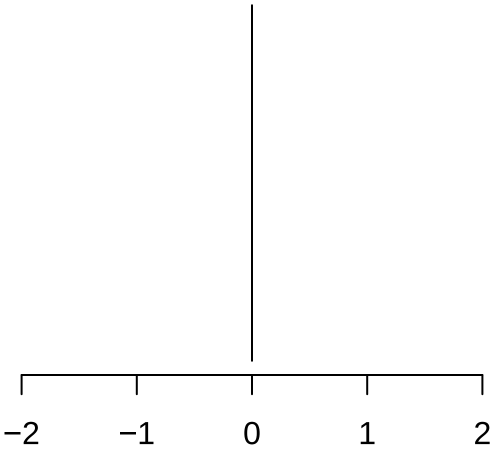
−2
−1
0
1
2

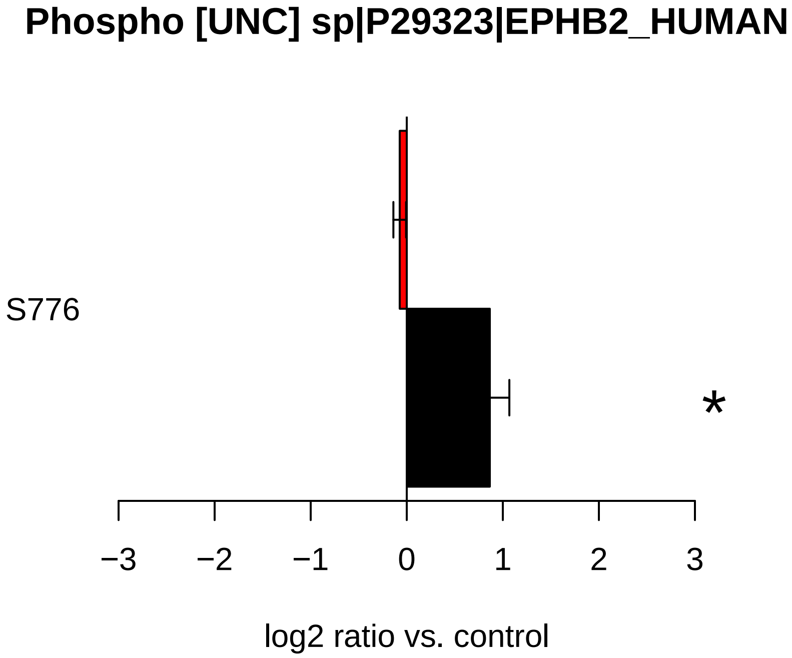
Phospho [UNC] sp|P29323|EPHB2_HUMAN
S776
*
−3
−2
−1
0
1
2
3
log2 ratio vs. control

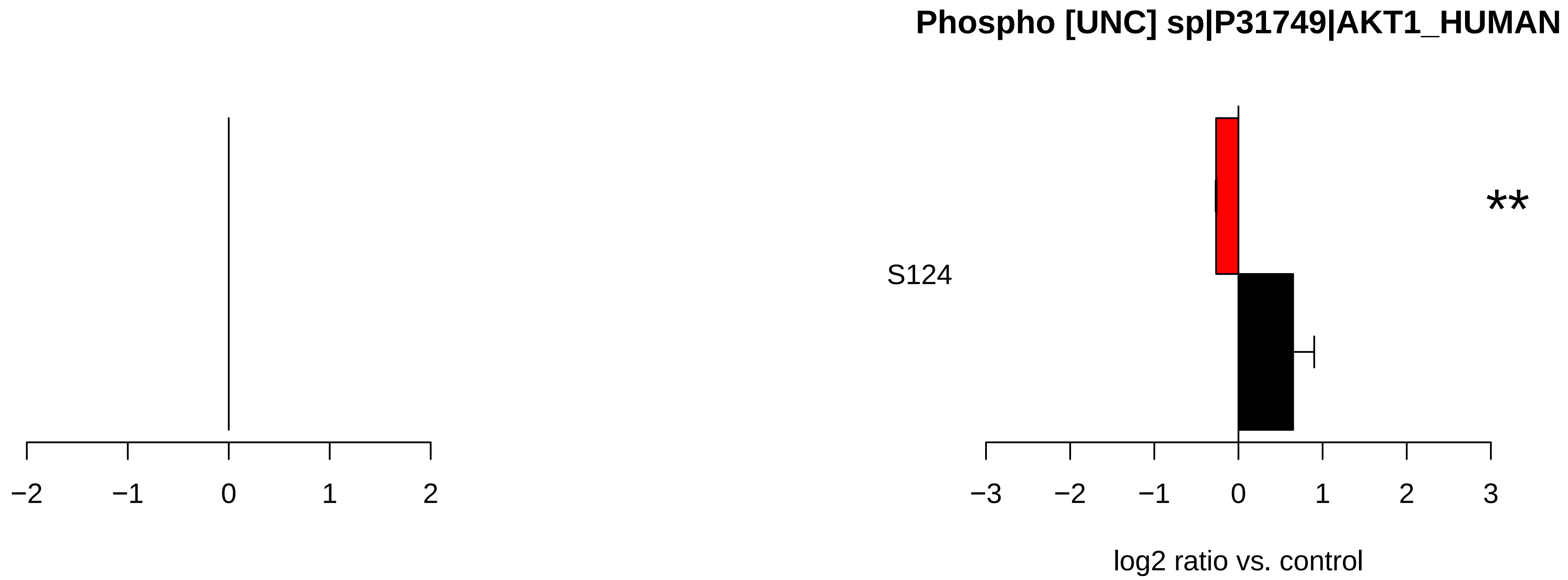
Phospho [UNC] sp|P31749|AKT1_HUMAN
**
S124
−2
−1
0
1
2
−3
−2
−1
0
1
2
3
log2 ratio vs. control

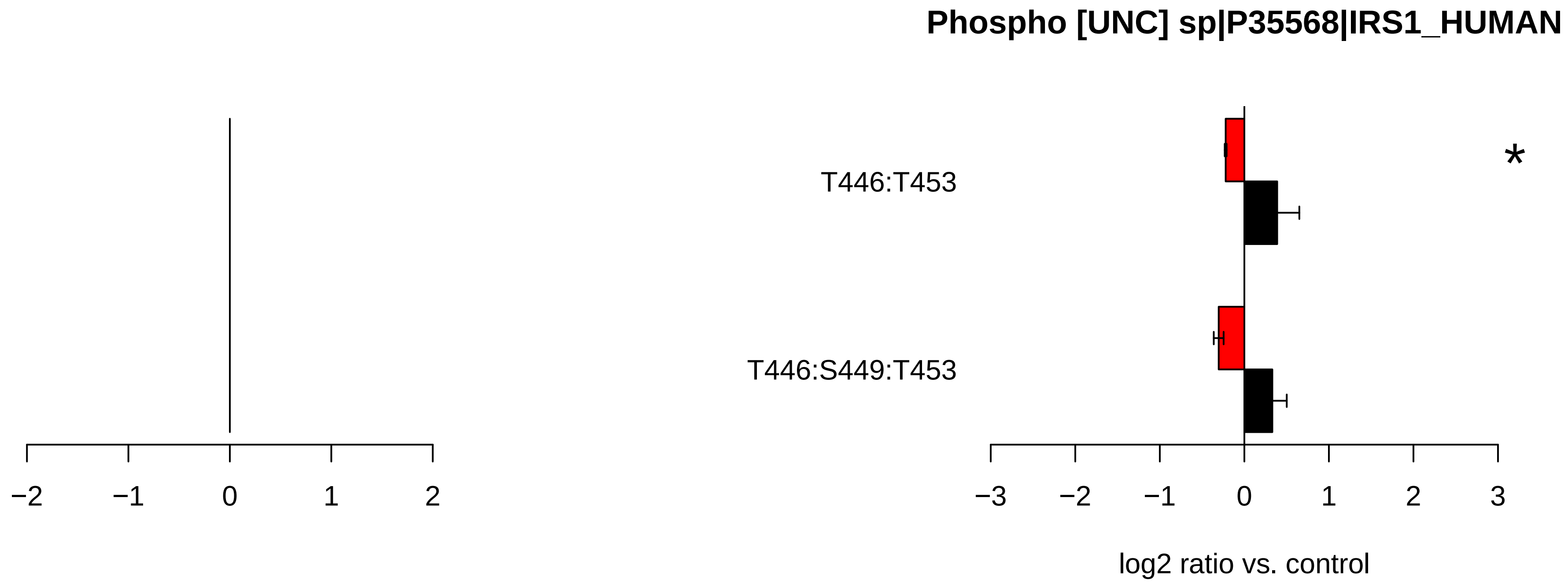

Phospho [UNC] sp|P35568|IRS1_HUMAN
T446:T453
T446:S449:T453
*
−2
−1
0
1
2
−3
−2
−1
0
1
2
3
log2 ratio vs. control

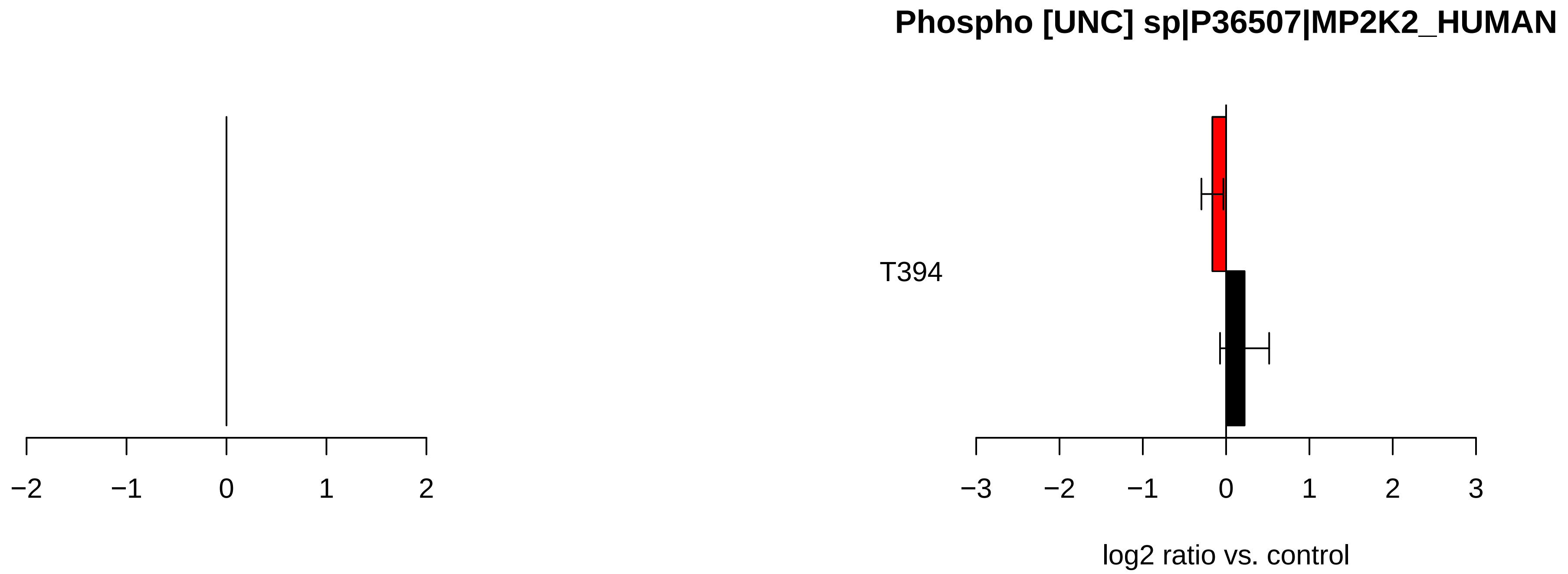
Phospho [UNC] sp|P36507|MP2K2_HUMAN
T394
−2
−1
0
1
2
−3
−2
−1
0
1
2
3
log2 ratio vs. control

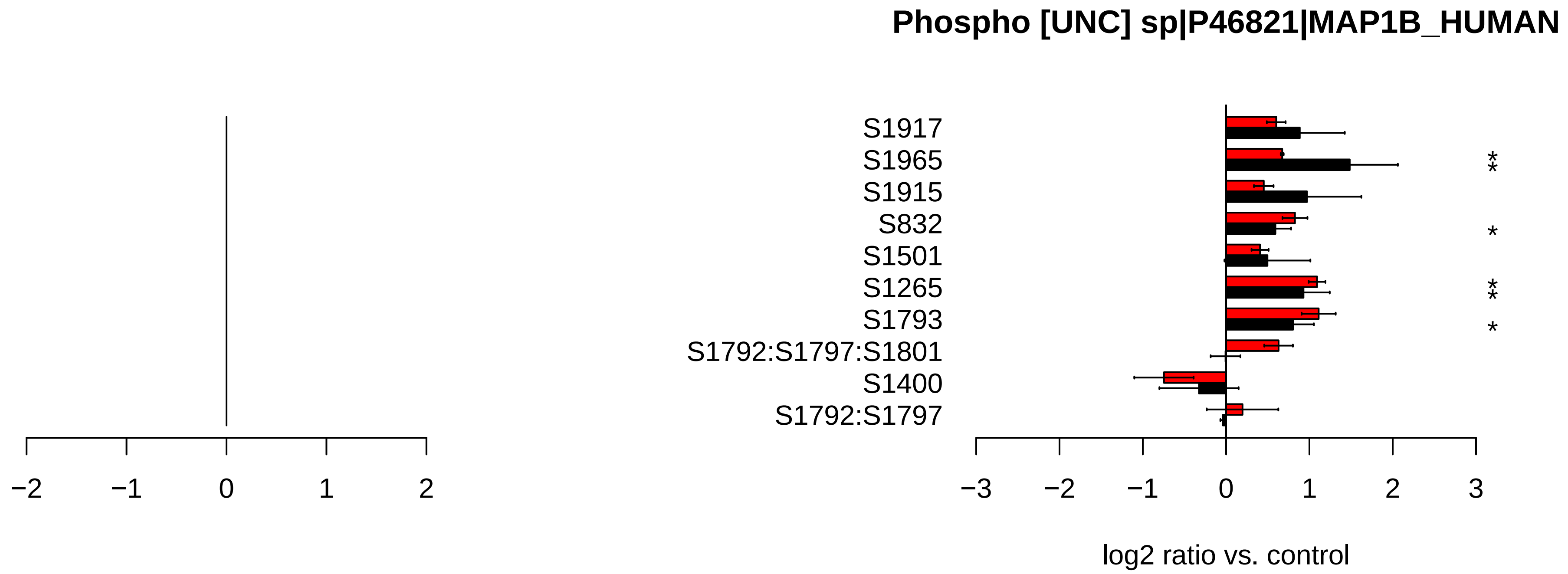

Phospho [UNC] sp|P46821|MAP1B_HUMAN
S1917
S1965
S1915
S832
S1501
S1265
S1793
S1792:S1797:S1801
S1400
S1792:S1797
−2
−1
0
1
2
−3
−2
−1
0
1
2
3
log2 ratio vs. control

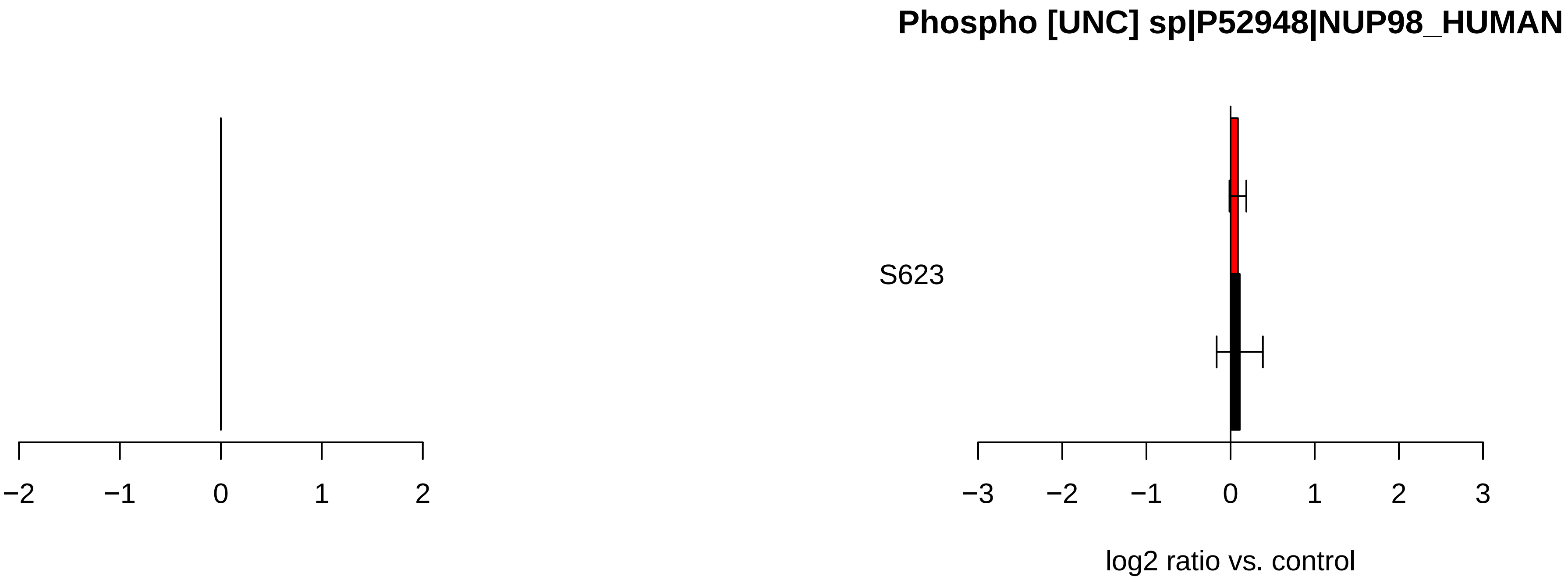

Phospho [UNC] sp|P52948|NUP98_HUMAN
S623
−2
−1
0
1
2
−3
−2
−1
0
1
2
3
log2 ratio vs. control

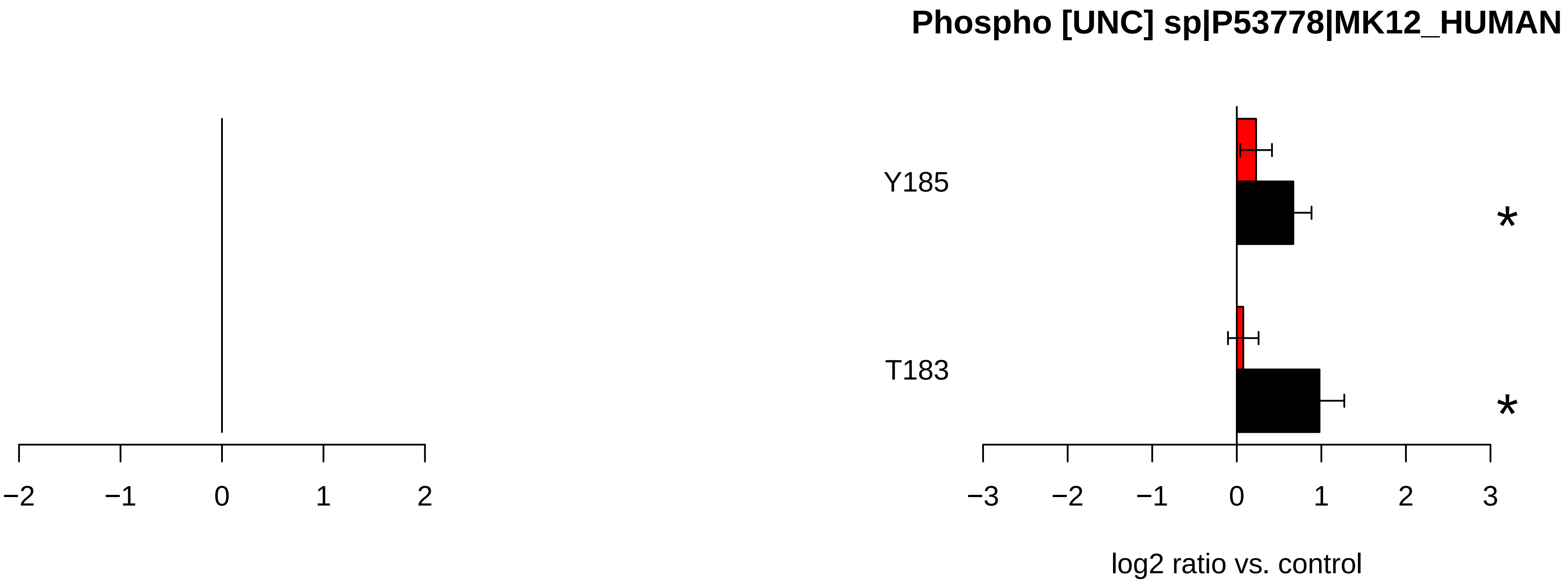

Phospho [UNC] sp|P53778|MK12_HUMAN
Y185
T183
*
*
−2
−1
0
1
2
−3
−2
−1
0
1
2
3
log2 ratio vs. control

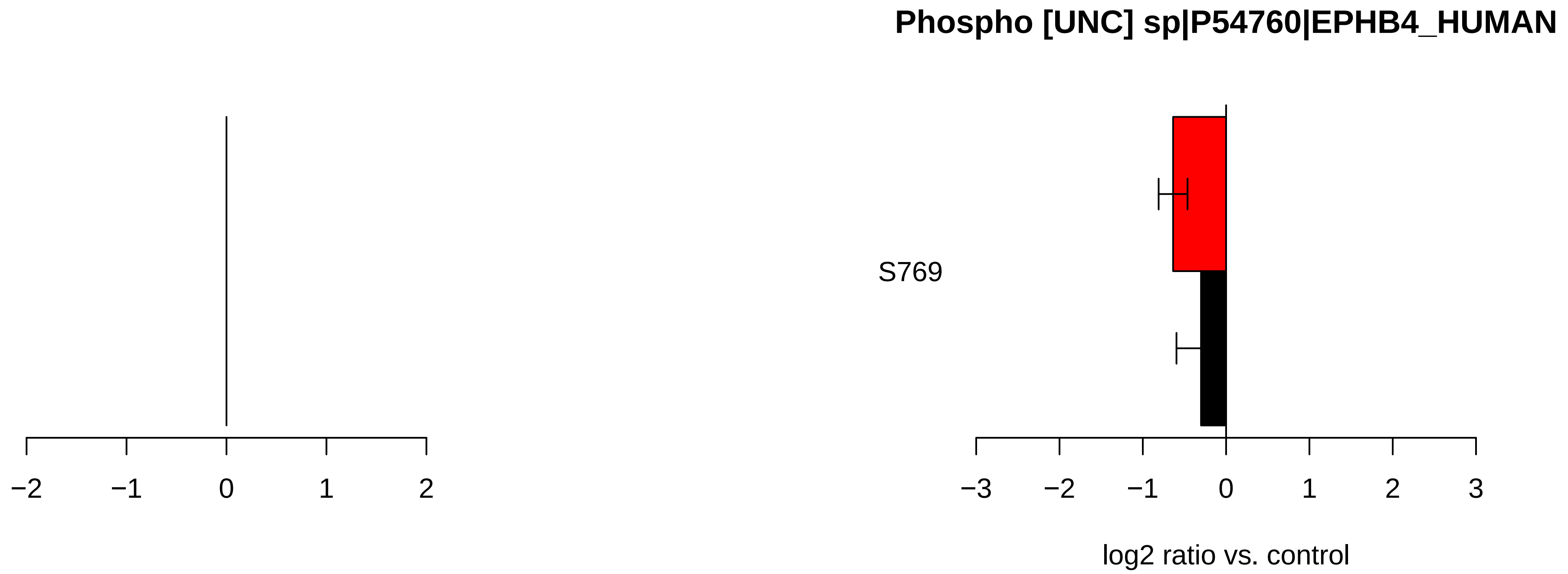

Phospho [UNC] sp|P54760|EPHB4_HUMAN
S769
−2
−1
0
1
2
−3
−2
−1
0
1
2
3
log2 ratio vs. control

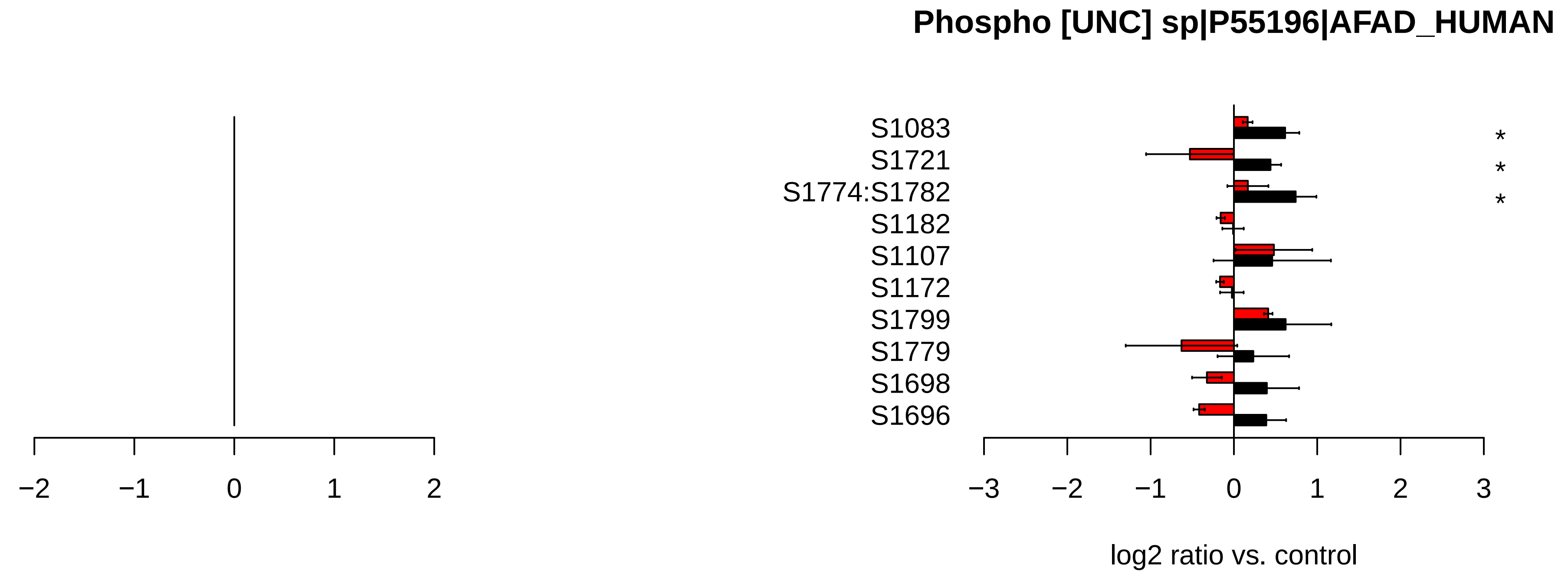

Phospho [UNC] sp|P55196|AFAD_HUMAN
S1083
S1721
S1774:S1782
S1182
S1107
S1172
S1799
S1779
S1698
S1696
*
*
*
−2
−1
0
1
2
−3
−2
−1
0
1
2
3
log2 ratio vs. control

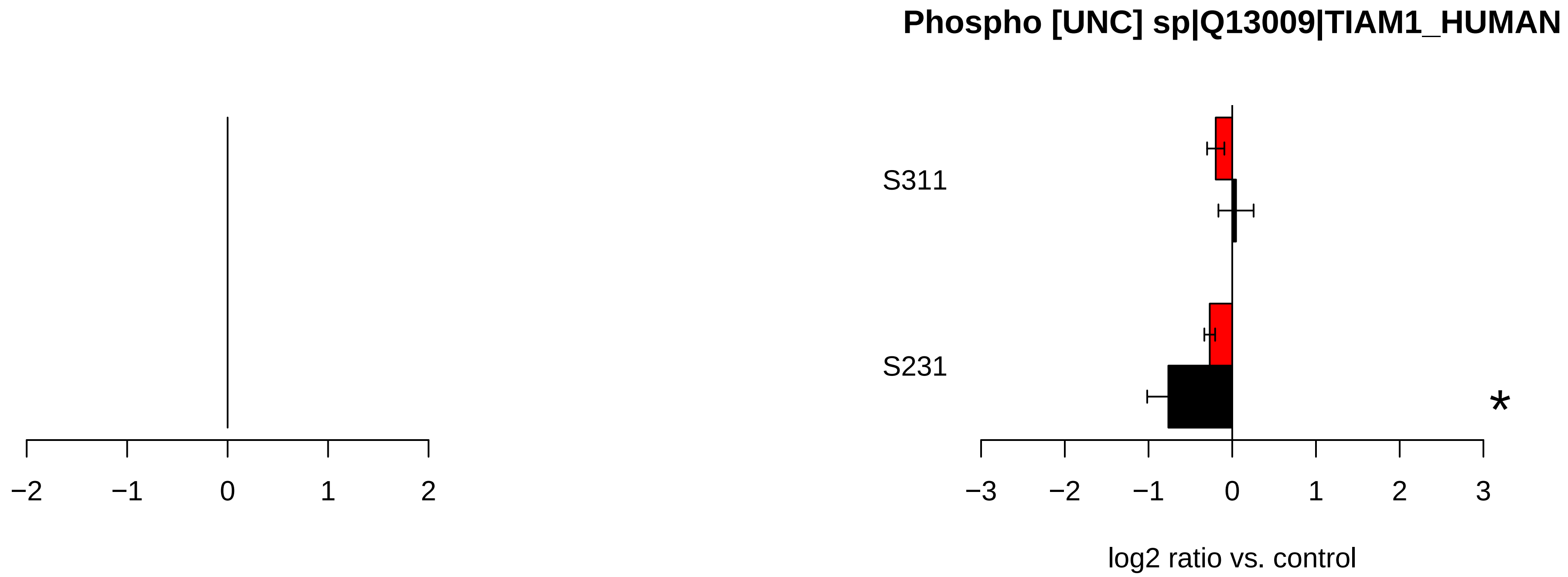
Phospho [UNC] sp|Q13009|TIAM1_HUMAN
S311
S231
*
−2
−1
0
1
2
−3
−2
−1
0
1
2
3
log2 ratio vs. control

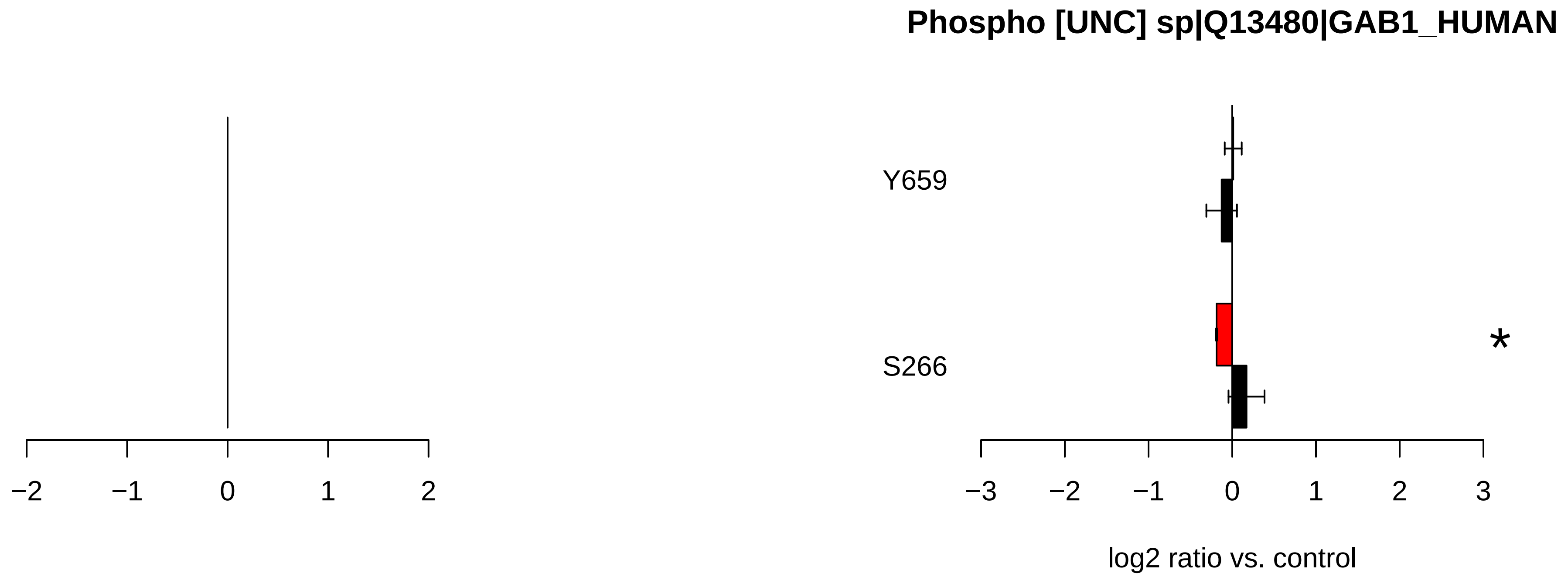
Phospho [UNC] sp|Q13480|GAB1_HUMAN
Y659
S266
*
−2
−1
0
1
2
−3
−2
−1
0
1
2
3
log2 ratio vs. control

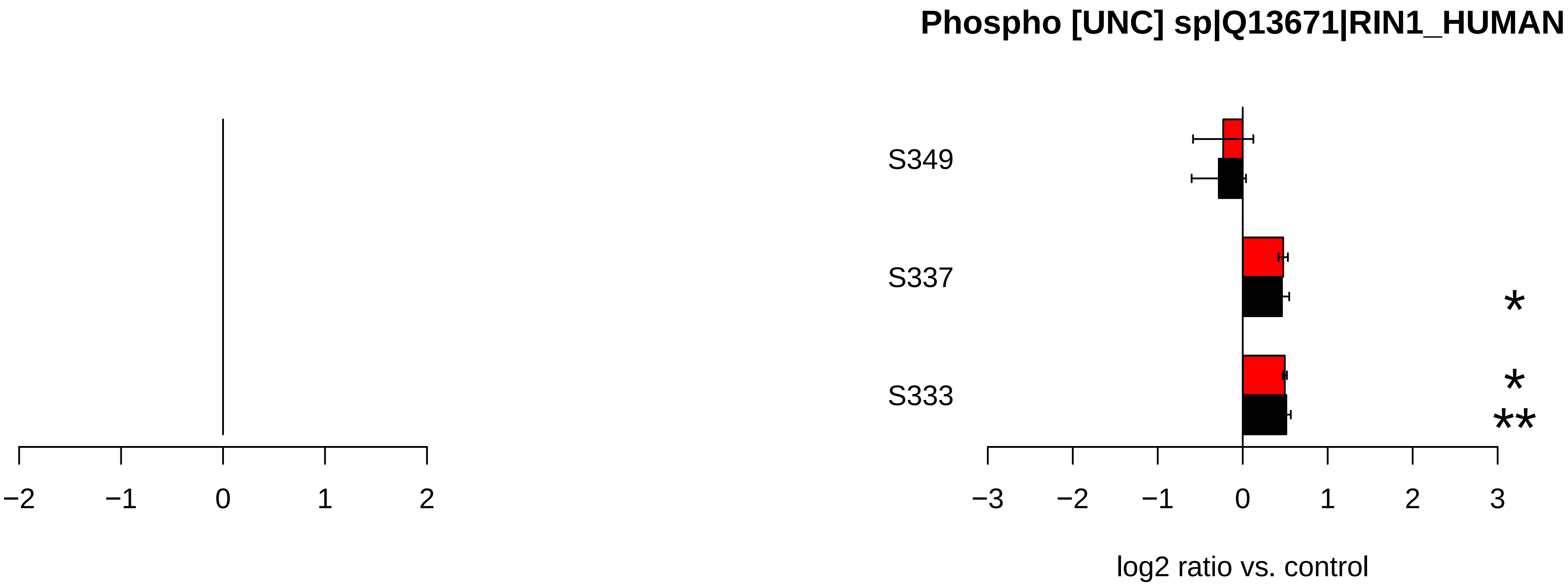

Phospho [UNC] sp|Q13671|RIN1_HUMAN
S349
S337
S333
*
*
**
−2
−1
0
1
2
−3
−2
−1
0
1
2
3
log2 ratio vs. control

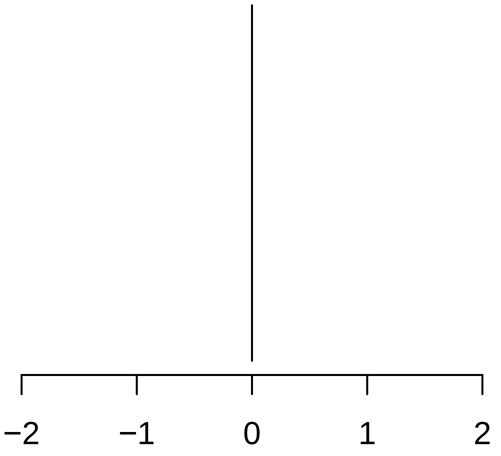

−2
−1
0
1
2


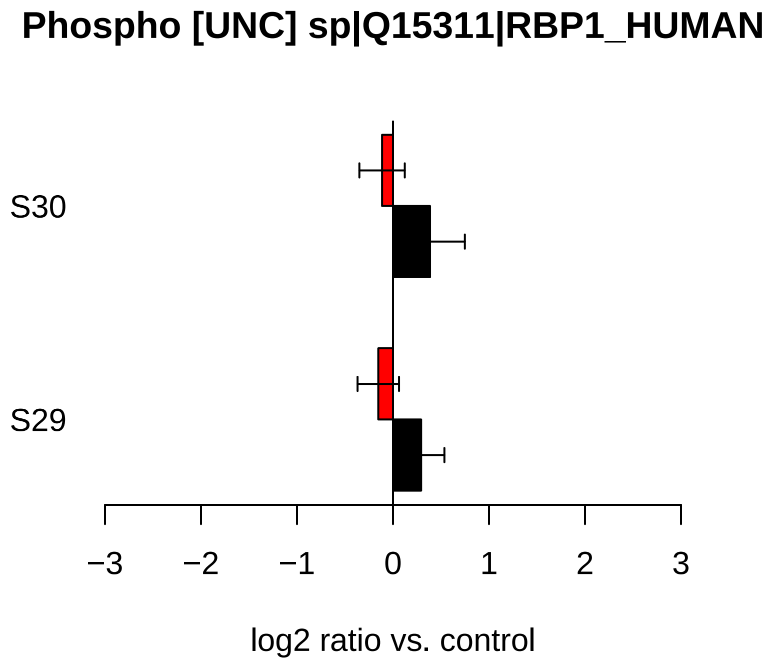

Phospho [UNC] sp|Q15311|RBP1_HUMAN
S30
S29
−3
−2
−1
0
1
2
3
log2 ratio vs. control

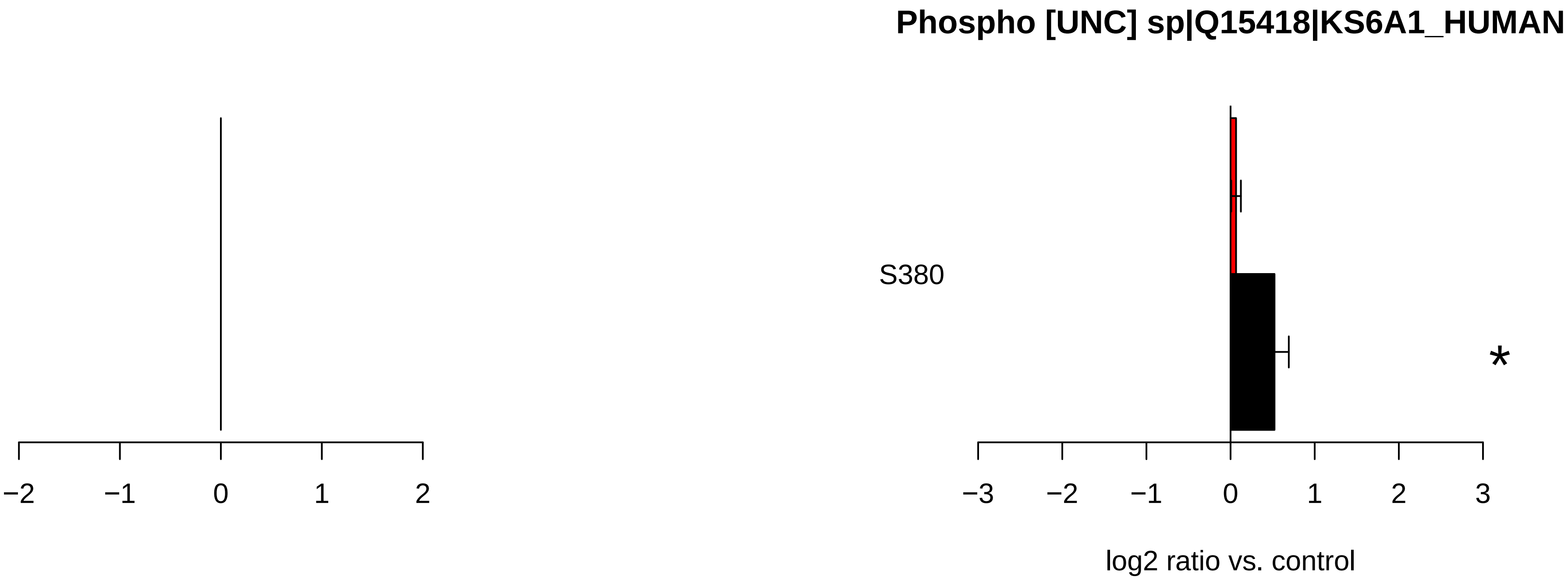

Phospho [UNC] sp|Q15418|KS6A1_HUMAN
S380
*
−2
−1
0
1
2
−3
−2
−1
0
1
2
3
log2 ratio vs. control

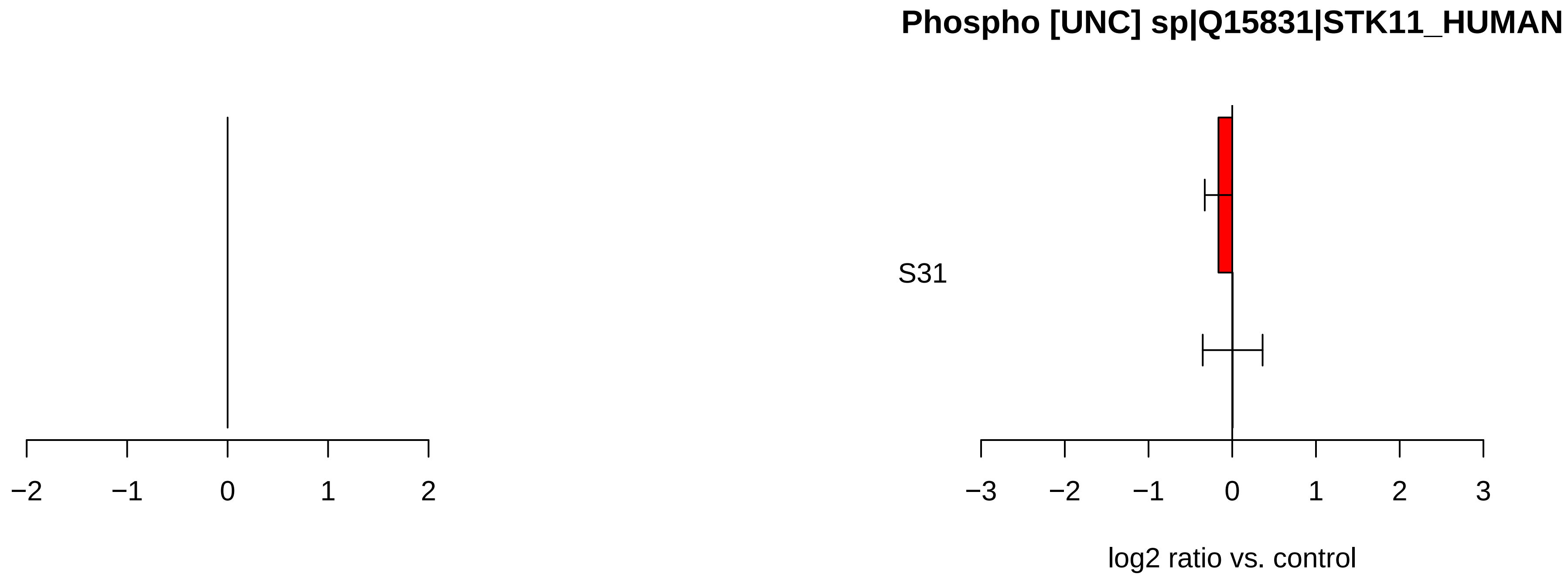
Phospho [UNC] sp|Q15831|STK11_HUMAN
S31
−2
−1
0
1
2
−3
−2
−1
0
1
2
3
log2 ratio vs. control

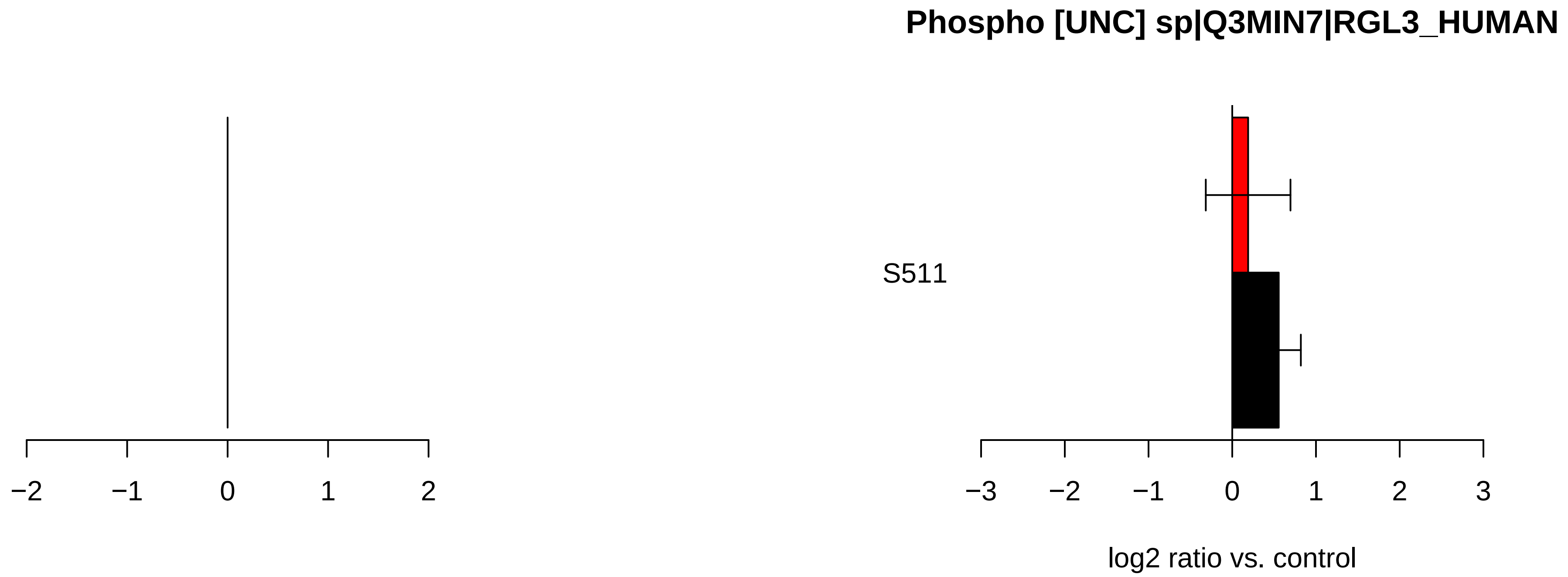
Phospho [UNC] sp|Q3MIN7|RGL3_HUMAN
S511
−2
−1
0
1
2
−3
−2
−1
0
1
2
3
log2 ratio vs. control

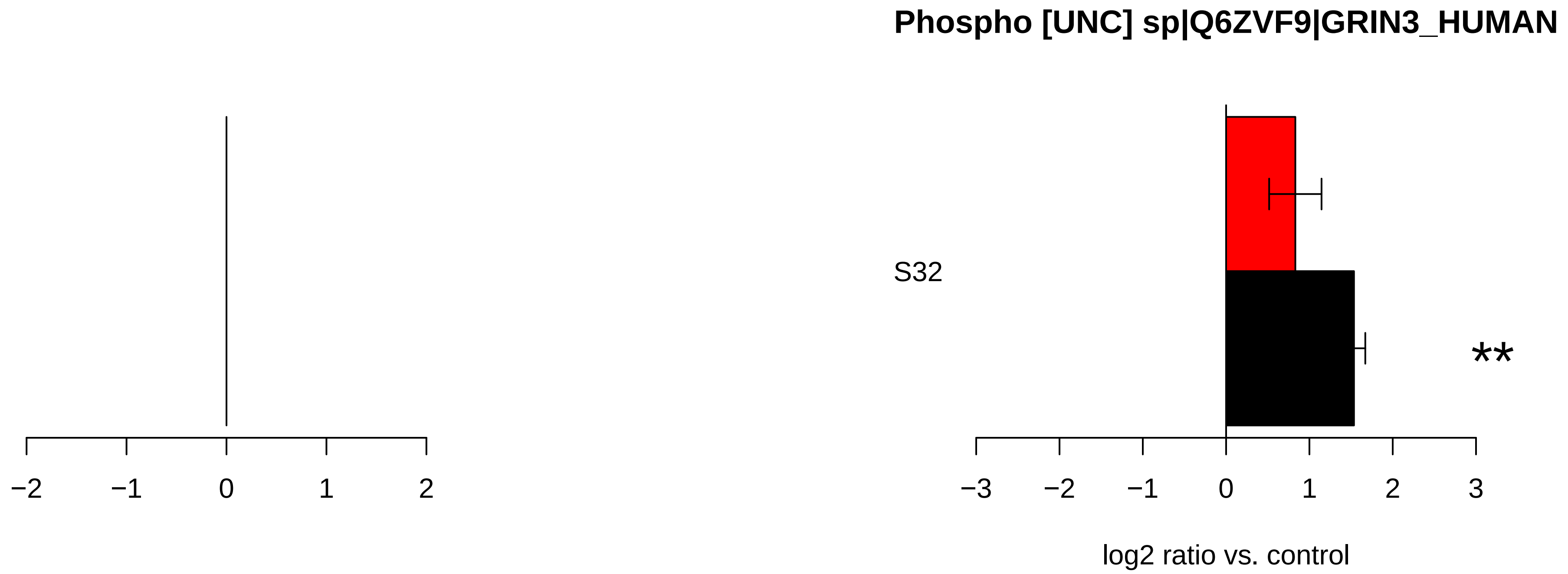

Phospho [UNC] sp|Q6ZVF9|GRIN3_HUMAN
S32
**
−2
−1
0
1
2
−3
−2
−1
0
1
2
3
log2 ratio vs. control

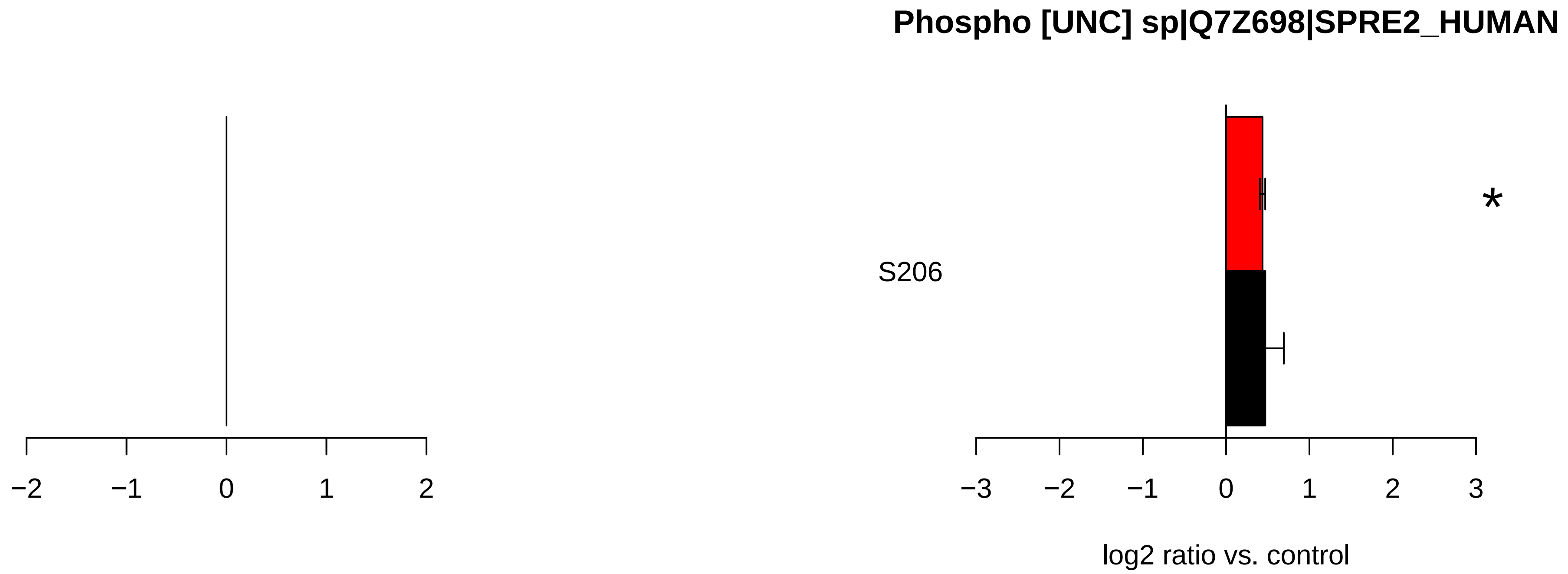

Phospho [UNC] sp|Q7Z698|SPRE2_HUMAN
S206
*
−2
−1
0
1
2
−3
−2
−1
0
1
2
3
log2 ratio vs. control

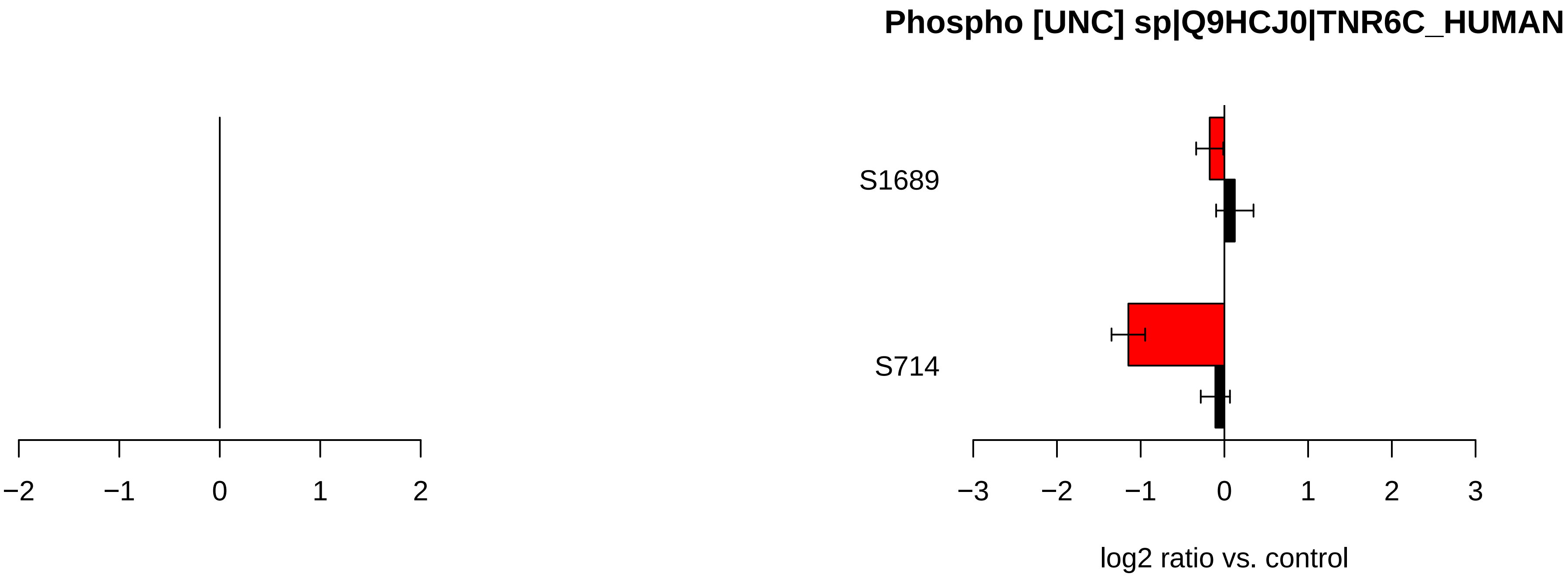
Phospho [UNC] sp|Q9HCJ0|TNR6C_HUMAN
S1689
S714
−2
−1
0
1
2
−3
−2
−1
0
1
2
3
log2 ratio vs. control

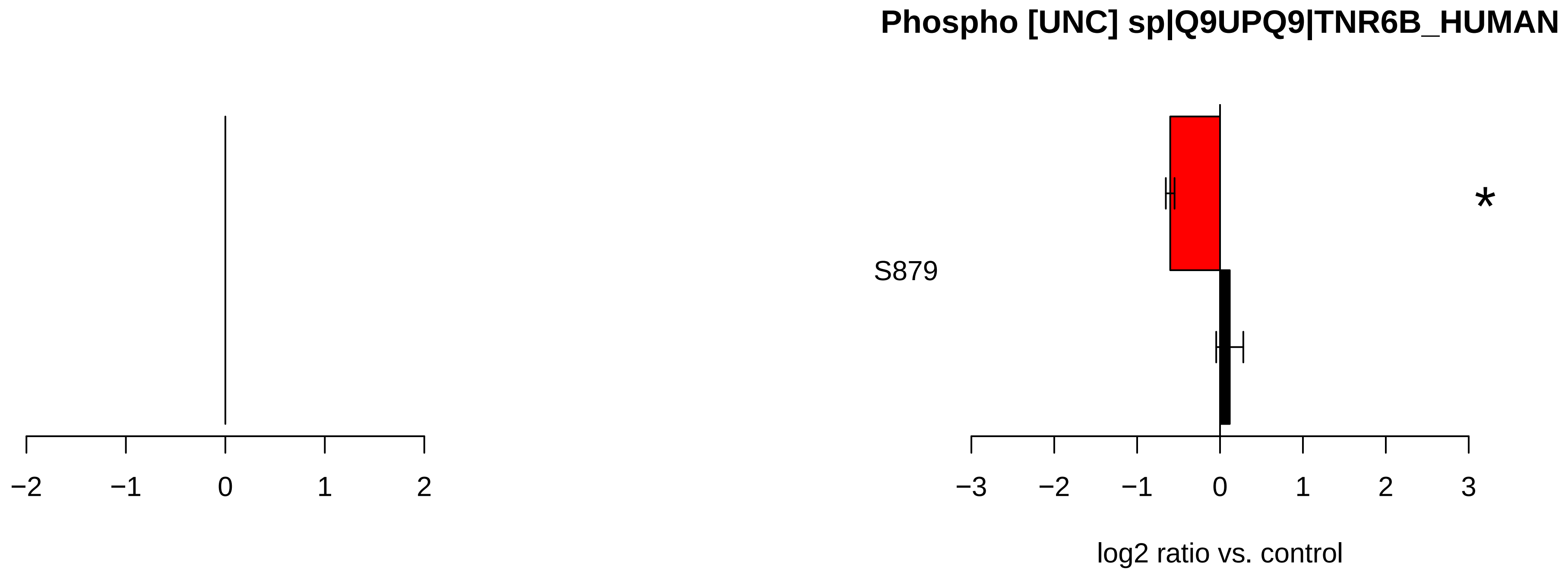
Phospho [UNC] sp|Q9UPQ9|TNR6B_HUMAN
S879
*
−2
−1
0
1
2
−3
−2
−1
0
1
2
3
log2 ratio vs. control

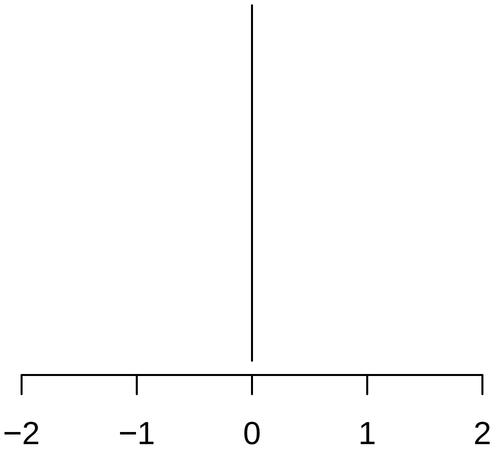
−2
−1
0
1
2

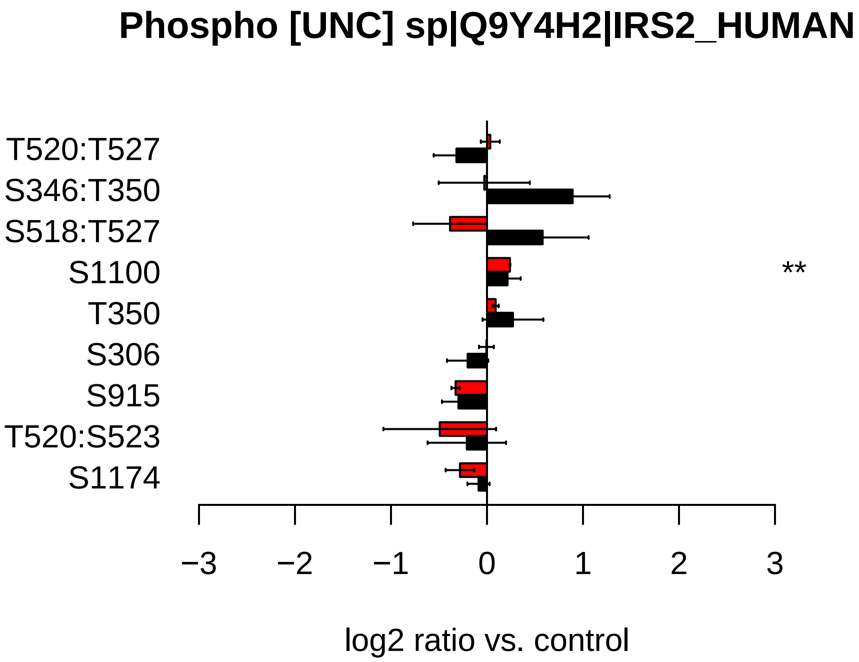
Phospho [UNC] sp|Q9Y4H2|IRS2_HUMAN
T520:T527
S346:T350
S518:T527
S1100
T350
S306
S915
T520:S523
S1174
**
−3
−2
−1
0
1
2
3
log2 ratio vs. control

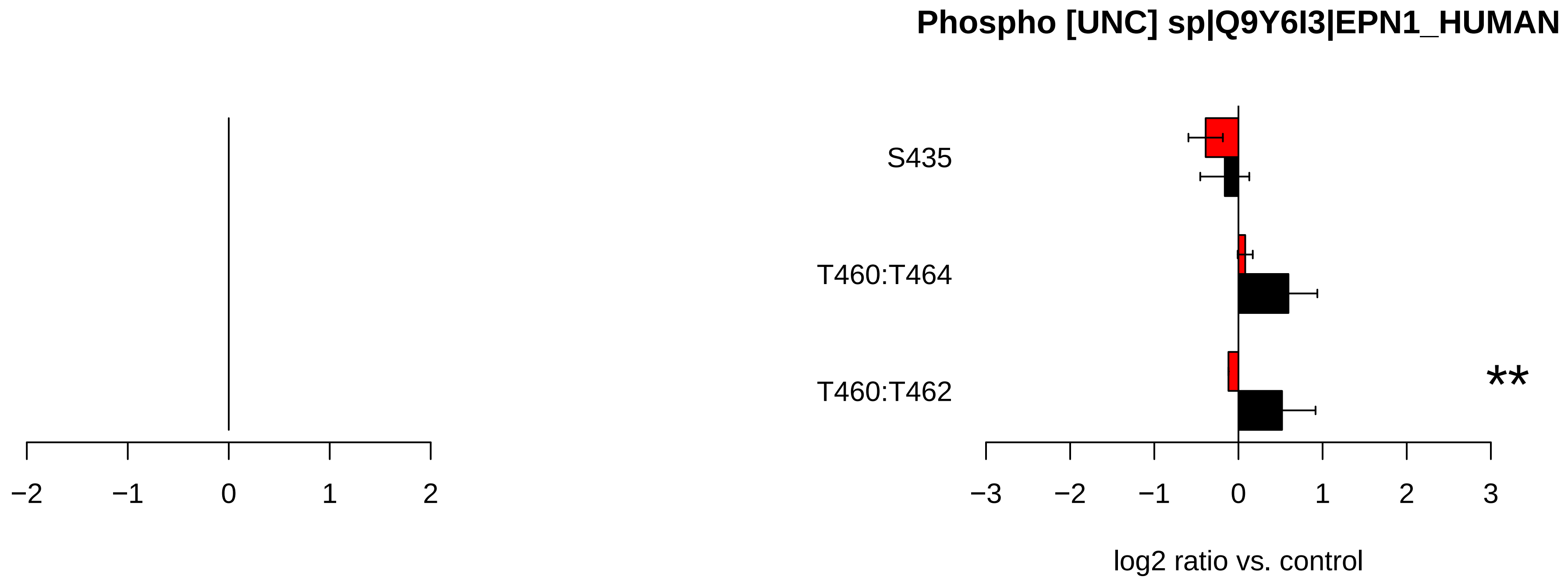
Phospho [UNC] sp|Q9Y6I3|EPN1_HUMAN
S435
T460:T464
T460:T462
**
−2
−1
0
1
2
−3
−2
−1
0
1
2
3
log2 ratio vs. control
